# The Septin Cytoskeleton is Required for Plasma Membrane Repair

**DOI:** 10.1101/2023.07.12.548547

**Authors:** M. Isabella Prislusky, Jonathan GT Lam, Viviana Ruiz Contreras, Marilynn Ng, Madeline Chamberlain, Sarika Pathak-Sharma, Madalyn Fields, Xiaoli Zhang, Amal O. Amer, Stephanie Seveau

## Abstract

Mammalian cells are frequently exposed to mechanical and biochemical stressors resulting in plasma membrane injuries. Repair mechanisms reseal the plasma membrane to restore homeostasis and prevent cell death. In the present work, a silencing RNA screen was performed to uncover plasma membrane repair mechanisms of cells exposed to a pore-forming toxin (listeriolysin O). This screen identified molecules previously known to repair the injured plasma membrane such as annexin A2 (ANXA2) as well as novel plasma membrane repair candidate proteins. Of the novel candidates, we focused on septin 7 (SEPT7) because the septins are an important family of conserved eukaryotic cytoskeletal proteins. Using diverse experimental approaches, we established for the first time that SEPT7 plays a general role in plasma membrane repair of cells perforated by pore-forming toxins and mechanical wounding. Remarkably, upon cell injury, the septin cytoskeleton is extensively redistributed in a Ca^2+^-dependent fashion, a hallmark of plasma membrane repair machineries. The septins reorganize into subplasmalemmal domains arranged as knob and loop (or ring) structures containing F-actin, myosin II, and annexin A2 (ANXA2) and protrude from the cell surface. Importantly, the formation of these domains correlates with the plasma membrane repair efficiency. Super-resolution microscopy shows that septins and actin are arranged in intertwined filaments associated with ANXA2. Silencing SEPT7 expression prevented the formation of the F-actin/myosin II/ANXA2 domains, however, silencing expression of ANXA2 had no observable effect on their formation. These results highlight the key structural role of the septins in remodeling the plasma membrane and in the recruitment of the repair molecule ANXA2. Collectively, our data support a novel model in which the septin cytoskeleton acts as a scaffold to promote the formation of plasma membrane repair domains containing contractile F-actin and annexin A2.

## INTRODUCTION

The plasma membrane of mammalian cells forms a biophysical barrier that separates the cell from its external environment. Any mechanical or biochemical perturbations that compromise the integrity of the plasma membrane can be lethal for the cell. Therefore, robust plasma membrane repair mechanisms maintain cell and tissue homeostasis [1, 2]. Excessive plasma membrane damage or defective plasma membrane repair is involved in many pathological conditions including muscular dystrophy, ischemia-reperfusion, heart failure, chronic inflammation, and neurodegenerative diseases [3]. Pathogens including parasites, bacteria, and viruses use diverse strategies to perforate the host cell plasma membrane and exploit the host cell repair responses to successfully infect their host [4–9]. In particular, the bacterial pathogen *Listeria monocytogenes* uses as a major virulence factor the pore-forming toxin listeriolysin O (LLO). LLO forms a pore complex across cholesterol-rich cytoplasmic and endosomal membranes promoting direct host cell invasion and cell-to-cell spreading [10–13]. LLO-mediated plasma membrane perforation is moderate enough that effective plasma membrane repair maintains cell viability to support the intracellular lifecycle of the pathogen [14–16]. How *L. monocytogenes*-infected cells repair their plasma membrane is not fully understood. LLO belongs to the cholesterol-dependent cytolysins (CDC) /Membrane Attack Complex/Perforin (MACPF) superfamily [17]. Several CDC members including streptolysin O (SLO) and perfringolysin O (PFO) have been successfully used to study conserved plasma membrane repair processes [18, 19]. Therefore, using LLO as a tool to inflict plasma membrane injury has the potential to uncover general repair machineries.

To uncover effectors that mediate the repair of LLO-perforated cells, we performed a siRNA screen targeting 245 protein-coding genes controlling endocytosis, exocytosis, and intracellular trafficking. The screen identified 47 candidates including previously known plasma membrane repair proteins such as calpain 1, the acid sphingomyelinase, several annexins, and components of the endosomal sorting complexes required for transport (ESCRT)-III [20–23]. The screen also identified novel plasma membrane repair candidates. Of those, we focused on septin 7 (SEPT7) because the septins are an important family of highly conserved eukaryotic GTP-binding proteins described as the fourth component of the cytoskeleton after F-actin, microtubules, and intermediate filaments [24]. Human septins include 13 members which assemble into hetero-hexamers (SEPT-2,-6,-7,-7,-6,-2) and hetero-octamers (SEPT-2,-6,-7,-9,- 9,-7,-6,-2) that further organize into filaments or rings in association with F-actin, microtubules or the plasma membrane [25, 26]. They form filaments and rings that determine the shape, curvature, and properties of the plasma membrane [25, 27]. The septins control major cellular functions including cytokinesis, cell motility, tissue morphogenesis, and host-pathogen interactions [28, 29]. However, the septin cytoskeleton had not been previously shown to be involved in plasma membrane repair. SEPT7 plays a central role in the formation of septin filaments as it is the only non-redundant septin among the 13 human septin family members [30, 31]. Our data show for the first time that SEPT7 plays a general role in plasma membrane repair of cells perforated by pore-forming toxins and mechanical wounding. Diverse microscopy methods, including super-resolution microscopy, established that the septin cytoskeleton reorganizes in injured cells to form subplasmalemmal knob and loop structures together with the contractile actomyosin cytoskeleton. Mechanistically, our data support a novel model in which the septins act as scaffolds required to promote the formation of plasma membrane repair domains containing contractile F-actin and annexin A2.

## RESULTS

### A siRNA screen identifies novel candidate genes controlling plasma membrane repair of listeriolysin O (LLO)-perforated cells

To identify the machineries involved in plasma membrane repair of LLO-perforated cells, we transfected HeLa cells with a library of siRNAs targeting 245 human genes controlling endocytosis, exocytosis, intracellular trafficking, and the cytoskeleton (Supplemental Table 1). Each gene was targeted for 72 h by a cocktail of three siRNAs with non-overlapping sequences. A fluorescence-based assay was then used to assess plasma membrane integrity [32]. In this assay, siRNA-treated HeLa H2B-GFP cells (expressing green-fluorescent histone 2B) were exposed, or not, to sub-lytic LLO concentration (Supplemental Figure (SF) 1 and movies 1, 2) for 30 min at 37°C [12, 14, 33]. The membrane-impermeant dye TO-PRO-3 was added to the cell medium, and TO-PRO-3 influx into damaged cells was measured as a readout for plasma membrane integrity. We identified 57 genes which silencing significantly affected membrane integrity, either negatively (47) or positively (10) (Table 1). Several of the 47 genes encode proteins previously shown to control plasma membrane repair such as the annexins (ANXA2, ANXA7, and ANXA11), the lysosomal enzyme acid sphingomyelinase (SMPD1), calpain 1 (CAPN1), and components of the ESCRT-III machinery (CHMP2, ANXA7) [20, 21, 23, 34, 35]. We also identified genes that encode proteins involved in membrane fusion (4 synaptotagmins and 12 SNAREs), lysosome biogenesis and functions (HPS3, HPS5, M6PRBP1, AP3S2, AP3M2, AP3S1, and Rab7) and proteins involved in exocytosis and the secretion of exosomes (EXOC1, EXOC6, Rab27A, Rab27B, and Rab11A). Finally, several genes controlling the actin cytoskeleton (IQSEC1, ABl1, GBF1, and SEPT7) were not previously known to control plasma membrane integrity. Of these candidates, we focused on SEPT7 because septins are a ubiquitous family of cytoskeletal proteins not previously known to control plasma membrane repair.

**Table 1.** Genes controlling plasma membrane integrity. The screen was conducted with HeLa cells and targeted 245 genes with a cocktail of three non-overlapping siRNAs. The assay measured TO-PRO-3 fluorescence intensity as a readout for plasma membrane integrity of cells incubated with or without 0.5 nM LLO for 30 min. Data are the statistical analyses comparing targeting siRNA-treated cells to cells treated with control siRNA. Silencing 47 genes led to a significant decrease in plasma membrane integrity and silencing 10 genes (italicized, negative values) significantly improved plasma membrane integrity.

| GENES | Estimate | st. dev. | P values | GENES | Estimate | st. dev. | P values |
| --- | --- | --- | --- | --- | --- | --- | --- |
| **Synaptotagmins** |  |  |  | **Coatmer** |  |  |  |
| SYT10 | 1.2844 | 0.1879 | <.0001 | PREB | 0.8879 | 0.2663 | 0.002 |
| SYT12 | 0.6103 | 0.2156 | 0.0076 | SAR1B | 0.8366 | 0.2742 | 0.0043 |
| SYT14 | 1.0435 | 0.1879 | <.0001 | SEC23A | 0.8038 | 0.2491 | 0.0027 |
| SYT4 | 0.9706 | 0.3385 | 0.0069 | SEC24B | 0.5802 | 0.2491 | 0.0255 |
| **SNARE** |  |  |  | **RAB** |  |  |  |
| SNAP25 | 1.0636 | 0.2779 | 0.0005 | RAB11A | 0.6593 | 0.2037 | 0.0026 |
| SNAP29 | 0.6257 | 0.2663 | 0.0244 | RAB27A | 0.6589 | 0.2037 | 0.0026 |
| VAMP3 | 0.5795 | 0.2663 | 0.0362 | RAB27B | 0.7774 | 0.2037 | 0.0005 |
| VAMP8 | 0.9043 | 0.2836 | 0.003 | RAB6C | 0.7302 | 0.213 | 0.0015 |
| STX11 | 0.8029 | 0.2836 | 0.0076 | RAB7A | 0.5283 | 0.2037 | 0.0137 |
| STX16 | 0.9561 | 0.2836 | 0.0018 | *RAB7L1* | 0.4598 | 0.2037 | 0.0302 |
| STX19 | 1.0977 | 0.1879 | <.0001 | RABL3 | 1.2823 | 0.1879 | <.0001 |
| STX1A | 0.7996 | 0.2779 | 0.0067 | RGS6 | 0.8321 | 0.2663 | 0.0035 |
| STX4 | 0.9928 | 0.2779 | 0.001 | *RAB2A* | *-0.3347* | *0.1263* | *0.0119* |
| STX8 | 0.7836 | 0.2663 | 0.0057 | *RAB40B* | *-0.9774* | *0.2851* | *0.0015* |
| STXBP1 | 0.786 | 0.2779 | 0.0076 | *RAB4A* | *-0.397* | *0.1263* | *0.0033* |
| STXBP4 | 0.9842 | 0.1879 | <.0001 | *RAB7B* | *-0.6466* | *0.1889* | *0.0016* |
| **Calpains** |  |  |  | **Intracellular trafficking** |  |  |  |
| CAPN1 | 0.5569 | 0.2466 | 0.03 | VPS13B | 0.9997 | 0.1879 | <.0001 |
| **Annexins** |  |  |  | *LMAN1* | *-1.2104* | *0.2409* | *<.0001* |
| ANXA2 | 0.59 | 0.2642 | 0.0319 | **Endocytosis** |  |  |  |
| ANXA7 | 0.7559 | 0.2642 | 0.007 | *CLTA* | *-0.5487* | *0.2026* | *0.0103* |
| ANXA11 | 0.5525 | 0.2642 | 0.0436 | *CLTB* | *-0.6981* | *0.2026* | *0.0015* |
| **ESCRT-I, -III** |  |  |  | *DNM2* | *-1.0475* | *0.2409* | *0.0001* |
| UBAP1 | 0.6887 | 0.2466 | 0.0083 | *AP2S1* | *-0.5095* | *0.2026* | *0.0165* |
| CHMP2A | 0.7421 | 0.1991 | 0.0007 | **Exocyst complex** |  |  |  |
| **Organelle biogenesis / lysosomes** |  |  |  | EXOC1 | 0.3266 | 0.1568 | 0.0444 |
| M6PRBP1 | 0.761 | 0.2491 | 0.0042 | *EXOC7* | *-0.6362* | *0.1702* | *0.0006* |
| HPS3 | 0.6083 | 0.2156 | 0.0077 | **Signaling** |  |  |  |
| SMPD1 | 0.8942 | 0.2532 | 0.0012 | IQSEC1 | 0.7637 | 0.2663 | 0.0069 |
| **Adaptor Complexes** |  |  |  | ABI1 | 0.5397 | 0.1702 | 0.0031 |
| AP3D1 | 0.7959 | 0.23 | 0.0014 | GBF1 | 0.9967 | 0.2836 | 0.0012 |
| AP3M2 | 0.6353 | 0.2491 | 0.0151 | **Septin** |  |  |  |
| AP3S2 | 0.8485 | 0.2491 | 0.0016 | SEPT7 | 0.5257 | 0.2466 | 0.0399 |
| AP4B1 | 0.7971 | 0.2491 | 0.0029 |  |  |  |  |

### The septins (SEPT6 and SEPT7) are required for plasma membrane repair

To confirm the role of the septins in plasma membrane repair, we first measured the knockdown efficiencies of each siRNA used in the screen to silence septins’ expression (SEPT2, 6, 7, and 9) (Figure 1A and SF 2A-E). We found that for each siRNA targeting SEPT7, SEPT9, and SEPT2, septin protein levels were decreased by 90-95% compared to control cells treated with non-targeting siRNA. However, SEPT6 expression was only decreased by about 35-40% for two siRNA whereas the third siRNA had no effect. As previously reported, silencing SEPT7 expression reduced SEPT2, SEPT6, and SEPT9 protein levels, whereas targeting SEPT2, SEPT6, and SEPT9 had little effect on the expression of the other tested septins (SF 2B-E) [36]. We then evaluated the effect of silencing septin expression on the plasma membrane integrity of LLO-perforated cells. Silencing SEPT7 with each siRNA resulted in a significant loss in plasma membrane integrity of LLO-treated cells (Figure 1B). This was unlikely due to an off-target effect because the three non-overlapping siRNAs were tested independently. To distinguish if the loss in plasma membrane integrity of SEPT7-deficient cells was due to an increase in plasma membrane perforation by LLO or to a defect in plasma membrane repair, we measured the integrity of cells exposed to LLO in Ca^2+^-free medium, an experimental condition preventing plasma membrane repair. In Ca^2+^-free medium, control siRNA-treated and SEPT7-deficient cells were similarly damaged leading to the conclusion that SEPT7-deficiency does not affect cell susceptibility to perforation by LLO (Figure 1C). Therefore, SEPT7 expression plays a significant role in plasma membrane repair. Furthermore, we previously showed that extracellular Ca^2+^ does not affect LLO pore formation [12]. Although the screen did not reveal a role for SEPT6, we found that SEPT6-siRNA1 and SEPT6-siRNA3, which both reduce SEPT6 expression, significantly impaired plasma membrane repair (Figure 1C, D). Importantly, SEPT6-siRNA2, which did not affect SEPT6 expression had no effect on cell integrity (data not shown). As observed in the screen, silencing SEPT2 and SEPT9 expression did not affect plasma membrane integrity (SF 2A’). We next generated a stable HeLa cell line with doxycycline-inducible expression of a short hairpin RNA (shRNA) targeting SEPT7 (D_i_SEPT7-shRNA1, Figure 1E, SF 3A). The targeting sequence was selected from the Genetic Perturbation Platform (Broad Institute) for its high specificity and was distinct from the three SEPT7 siRNA sequences used in the screen. We verified that doxycycline had no effect on LLO pore formation (SF 3B) and then repeated the repair assay comparing the ratio intensity of doxycycline-treated to non-treated D_i_SEPT7-shRNA1 and control HeLa cells in the presence or absence of extracellular Ca^2+^. Only in the presence of LLO and extracellular Ca^2+^, plasma membrane repair of SEPT7-deficient cells was significantly impaired in comparison to control cells, which confirmed the role of SEPT7 in plasma membrane repair of LLO-injured cells (Figure 1F).

**Figure 1.**
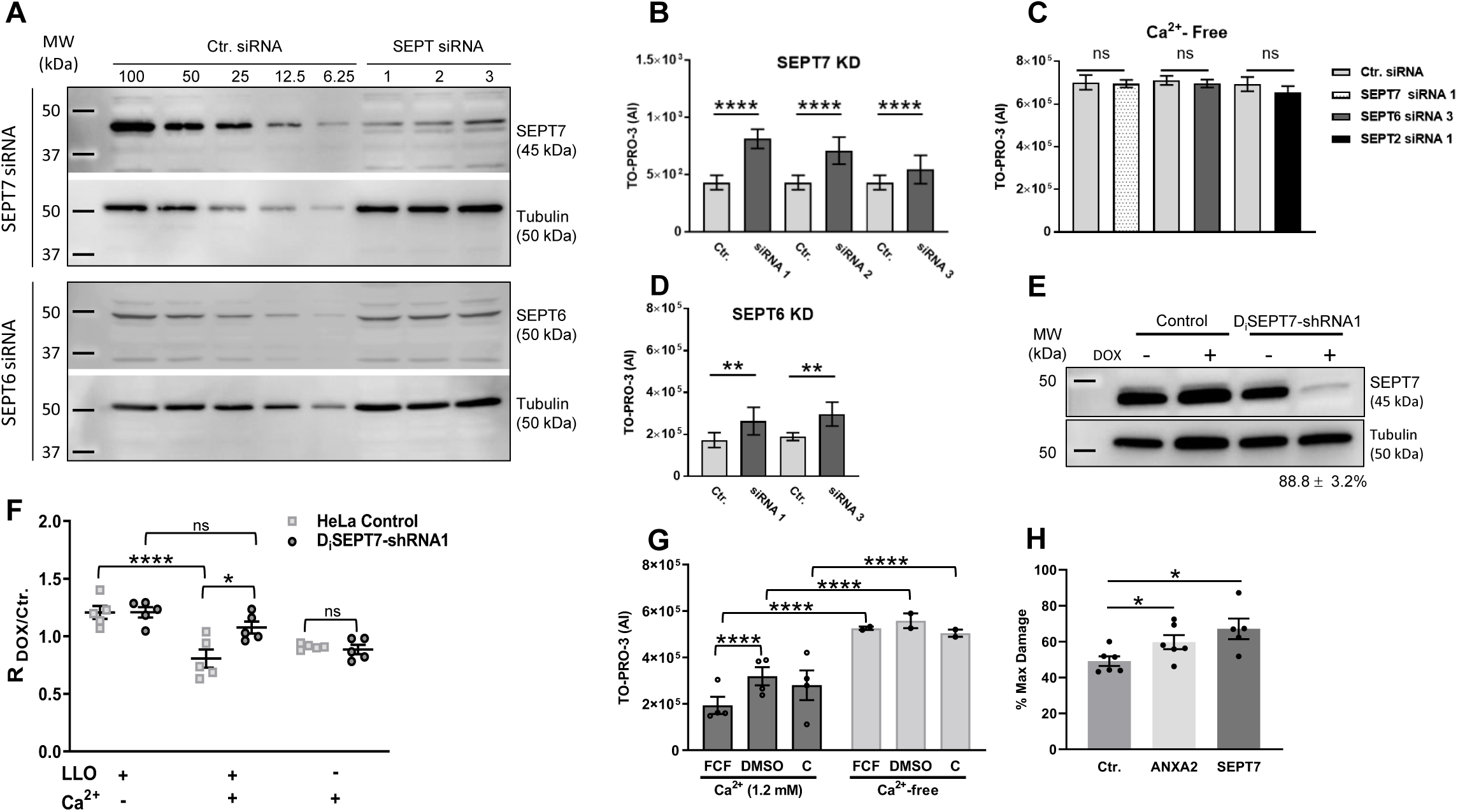
SEPT6 and SEPT7 are required for efficient plasma membrane repair. (**A**) HeLa cells were transfected with non-targeting siRNA (Ctr. siRNA) or with siRNAs targeting SEPT6 and SEPT7 (Supplemental Table 1). After 72 h, cells were lysed and analyzed by SDS-PAGE and immunoblotting. Serial dilutions of control (100% to 6.25%) and undiluted septin siRNA-treated (100%) cell lysates were loaded in the gel to facilitate the quantification of the KD efficiencies. Blots are representative of at least 5 independent experiments (N=5). KD efficiencies for SEPT6 were 39.5% ± 8.6; 3.6% ± 3.6; and 45.5% ± 12.5 for siRNA1, 2, and 3, respectively (Average ± SEM). KD efficiencies for all SEPT7 siRNAs were >90%. (**B to D**) Ctrl and SEPT-specific siRNAs treated cells were exposed to LLO (0.5 nM) for 30 min in M1 (1.2 mM Ca^2+^) (B, D) or M2 (Ca^2+^-free) (C), supplemented with TO-PRO-3. Data are the average TO-PRO-3 fluorescence intensities expressed in arbitrary units (AI) ± SEM of at least N=3 at time point 30 min. (**E**) Control HeLa cells and D_i_SEPT7-shRNA HeLa cells were cultured for 72 h in the presence (+) or in the absence (-) of Dox (50 μg/ml) and were tested for SEPT7 KD efficiency by immunoblotting. The presented blot is representative of N=8 with a KD efficiency of 88.75±3.2% (Average ± SEM). (**F**) Control HeLa cells and D_i_SEPT7-shRNA1 HeLa cells were cultured for 72 h with or without DOX and then washed to remove the DOX. Cells were exposed, or not, to 0.5 nM LLO for 30 min in M1 or M2 supplemented with TO-PRO-3. Data are expressed as the average ratio of the TO-PRO-3 fluorescence intensity of DOX-treated over TO-PRO-3 fluorescence intensity of non-treated (Ctr.) cells (R_Dox/Ctr._) ± SEM of N=5 at time point 30 min. (**G**) HeLa cells were pre-treated with 100 μM FCF (in DMSO), vehicle DMSO, or untreated (C) for 16 h. Cells were then subjected to the repair assay for 30 min with 0.5 nM LLO in M1 or M2 with TO-PRO-3. Due to FCF reversibility, FCF and DMSO were added to the buffers during the repair assay. Data are the average ± SEM of N=2 for Ca^2+^-free condition and N=4 for 1.2 mM Ca^2+^ condition at 30 min. (**H**) Ctr.-, SEPT7-siRNA 1-, and ANXA2-siRNA 3-treated cells were mechanically wounded in M1 or M2 buffers. The number of Emerald cells were counted to represent the number of damaged cells and the number of Ruby cells that were originally green were counted to represent the number of not recovered cells. The data is expressed at the number of non-recovered cells (Ruby)/ the number of Damaged cells (Emerald) which have been normalized to the Ca^2+^ -free condition. A Students Paired T Test was used to analyze the data. TO-PRO-3 intensity was measured by fluorescence microscopy (**B** and **F**) or by spectrofluorometry (**C, D** and **G**). For **B-D** and **F-G** data were log_10_ transformed and analyzed using linear mixed effects models. For **H** data were analyzed by Students Paired T-Test (*P< 0.05, **P< 0.01, *** P< 0.001, and **** P<0.0001).

As a third experimental approach to establish the role of the septin cytoskeleton in plasma membrane repair, we used the pharmacological agent forchlorfenuron (FCF), which is known to reversibly bind to and stabilize septin oligomers and/or filaments [37–39]. In the presence of Ca^2+^, cell treatment with FCF significantly decreased plasma membrane damage of LLO-treated cells (Figure 1G, SF 3C, D). In Ca^2+^-free medium, FCF treatment did not affect cell susceptibility to LLO-perforation (Figure 1G, SF 3D) and FCF had no direct effect on LLO pore formation (SF 3E). Together, these data indicate that stabilization of the septin cytoskeleton by FCF increases the plasma membrane repair efficiency of LLO-treated cells.

To ensure that the role of SEPT7 in plasma membrane repair was not specific to LLO-perforated cells, control- and SEPT7-siRNA treated cells were exposed to the pore-forming toxin pneumolysin (PLY), another CDC produced by the pathogen *Streptococcus pneumoniae* [40]. As previously observed with LLO (Figure 1B), SEPT7 expression was also required for effective plasma membrane repair of PLY-perforated cells (SF 3F, G). To determine if SEPT7 plays a general role in plasma membrane repair, we next subjected cells to mechanical wounding using glass beads. As a positive control, we silenced the expression of ANXA2 (SF 3H), which was previously known to repair mechanical wounds [41]. We found that SEPT7-deficiency, similar to ANXA2-deficiency, significantly impaired plasma membrane repair of mechanically wounded cells (Figure 1H). Collectively, our data demonstrate a novel and general role for SEPT7 in plasma membrane repair of toxin-perforated and mechanically wounded cells.

### Plasma membrane perforation remodels the septin cytoskeleton together with cortical F-actin, myosin-IIA, and annexin A2

The septin cytoskeleton is known to associate with and to regulate the actin cytoskeleton. Therefore, we studied the distribution of the septins relative to F-actin. HeLa cells were treated, or not, with LLO for 5 to 15 min at 37°C, followed by chemical fixation and fluorescent labeling of SEPT2, SEPT7, or SEPT9. We found that SEPT2, SEPT7, and SEPT9 display similar labeling patterns in all experimental conditions. This was expected since septins of different groups co-assemble to form hexamers (SEPT-2,-6,-7,-7,-6,-2) and octamers (SEPT- 2,-6,-7,-9,-9,-7,-6,-2) [25]. We observed that septin filaments predominantly colocalized with actin stress fibers in control untreated cells (Figure 2Aa, SF 4Aa, Ba). Strikingly, in LLO-treated cells, a fraction of the septin cytoskeleton clearly dissociated from the actin stress fibers (Figure 2Ab, SF 4Ab, Bb) and reorganized into knob- and loop (or ring)-like structures in association with cortical F-actin (Figure 2Ac, d, SF 4Ac, Bc). The formation of these new septin structures increased over time from 5 to 15 min post-LLO-exposure (Figure 2B, C, SF 4C). As shown in Figure 3A, the knob- and loop-like septin/F-actin structures also strongly colocalized with myosin-IIA indicating that the newly formed structures are contractile. As the septins have been proposed to regulate plasma membrane properties [25], we thought that the septins may control the organization of plasma membrane repair domains. To test this hypothesis, we investigated if the remodeled septins co-distribute with known plasma membrane repair machineries. Our screen identified that annexins and the ESCRT-III are important for the integrity of LLO-perforated cells, and the annexins and ESCRT-III have been shown to promote plasma membrane repair of perforated cells [34, 42]. We observed that the ESCRT-III ALG-2-interacting protein X (ALIX) formed larger puncta in LLO-treated cells in comparison to control cells, but rarely colocalized with the septins (SF 5A). However, in the presence of LLO, there was a clear redistribution of ANXA2 which colocalized with the septins and F-actin, as observed in cells transiently expressing ANXA2-GFP and in cells fluorescently labeled with anti-ANXA2 antibodies (Figure 3B, B’). The co-distribution of ANXA2, the septins, and F-actin was observed throughout the cell surface and within the knob and loop structures (Figure 3B’). Quantitative analysis showed that the septins display high level of colocalization with both F-actin and ANXA2 but not with ALIX (Figure 3C). In conclusion, the septin cytoskeleton, visualized via labeling of SEPT2, SEPT7, or SEPT9, is remodeled in LLO-perforated cells to colocalize with contractile actomyosin cytoskeleton and ANXA2.

**Figure 2.**
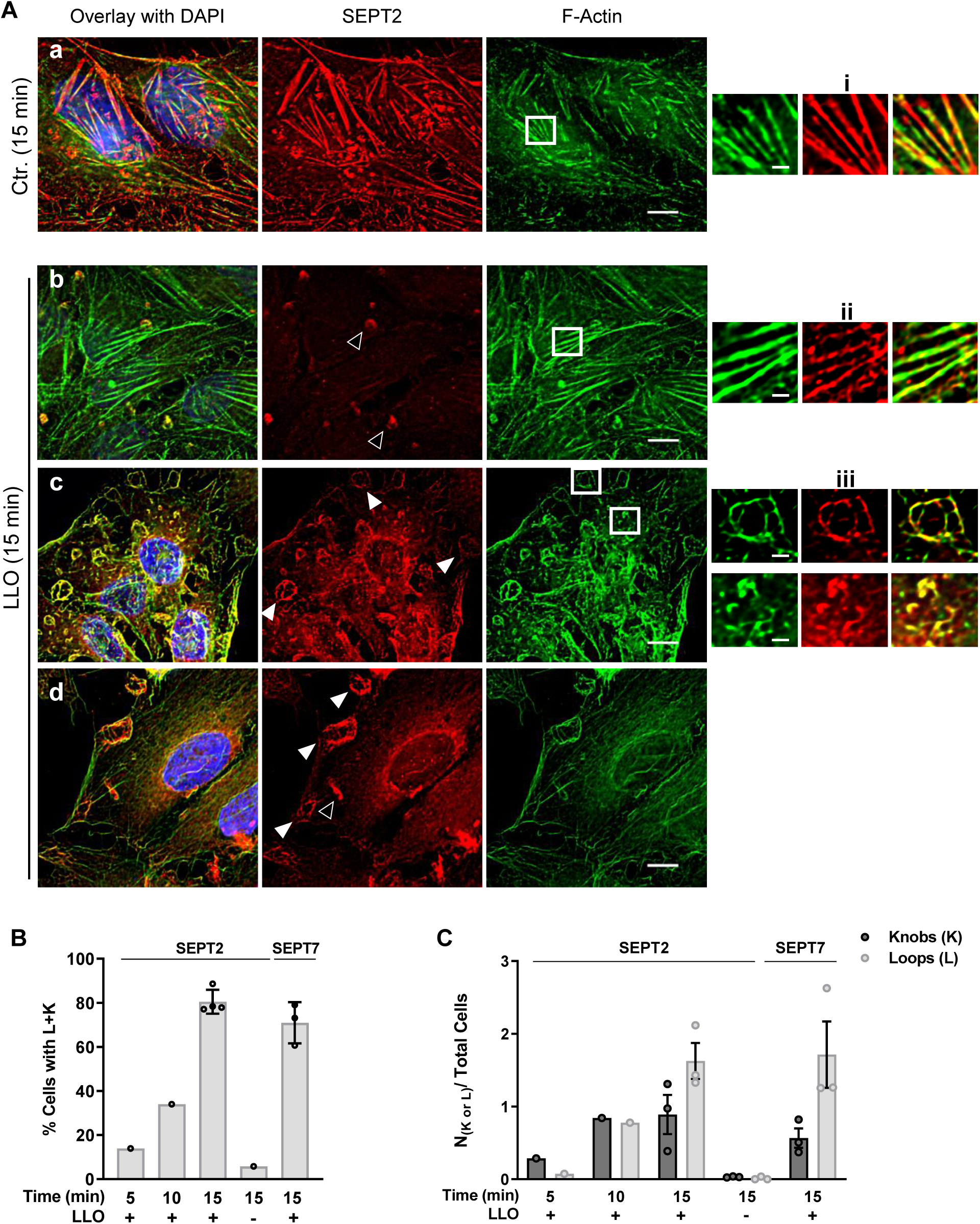
The septin cytoskeleton is remodeled in LLO-perforated cells. HeLa cells were incubated with or without (Ctr.) LLO (0.5 nM) for 5-15 min in repair permissive buffer M1 (1.2 mM Ca^2+^). Cells were fixed, permeabilized and fluorescently labeled with anti-SEPT primary Abs and Alexa Fluor 568-conjugated secondary Abs, Alexa Fluor 488- or 647-conjugated phalloidin, and DAPI. (**A**) Representative images at time point 15 min. Septin fluorescence images are presented with the same intensity scaling to visualize the loss of septin association with F-actin. To better visualize septin and F-actin, selected regions were magnified (Ai-Aiii, Bi-Biii) and the septin fluorescence display was artificially enhanced only in Aii and Bii. Scale bars are 10 μm (A) and 2 μm (Ai-Aiii, Bi-Biii). Images were acquired by z-stack widefield microscopy, deconvolved, and presented as the best focus images, except for Aa and Ab which are single planes focused on actin stress fibers. Knob and loop structures are indicated by unfilled and filled arrowheads, respectively. (**B**) The percentage of cells with septin knobs and loops (%Cells with K+L) and (**C**) the average number of knobs or loops per total cells (N _(K or L)_/Total Cell) were enumerated based on septin fluorescence. A total of 300-750 cells were analyzed in each experimental condition, N=3 at 15 min, and N=1 at 5- and 10-min time points.

**Figure 3.**
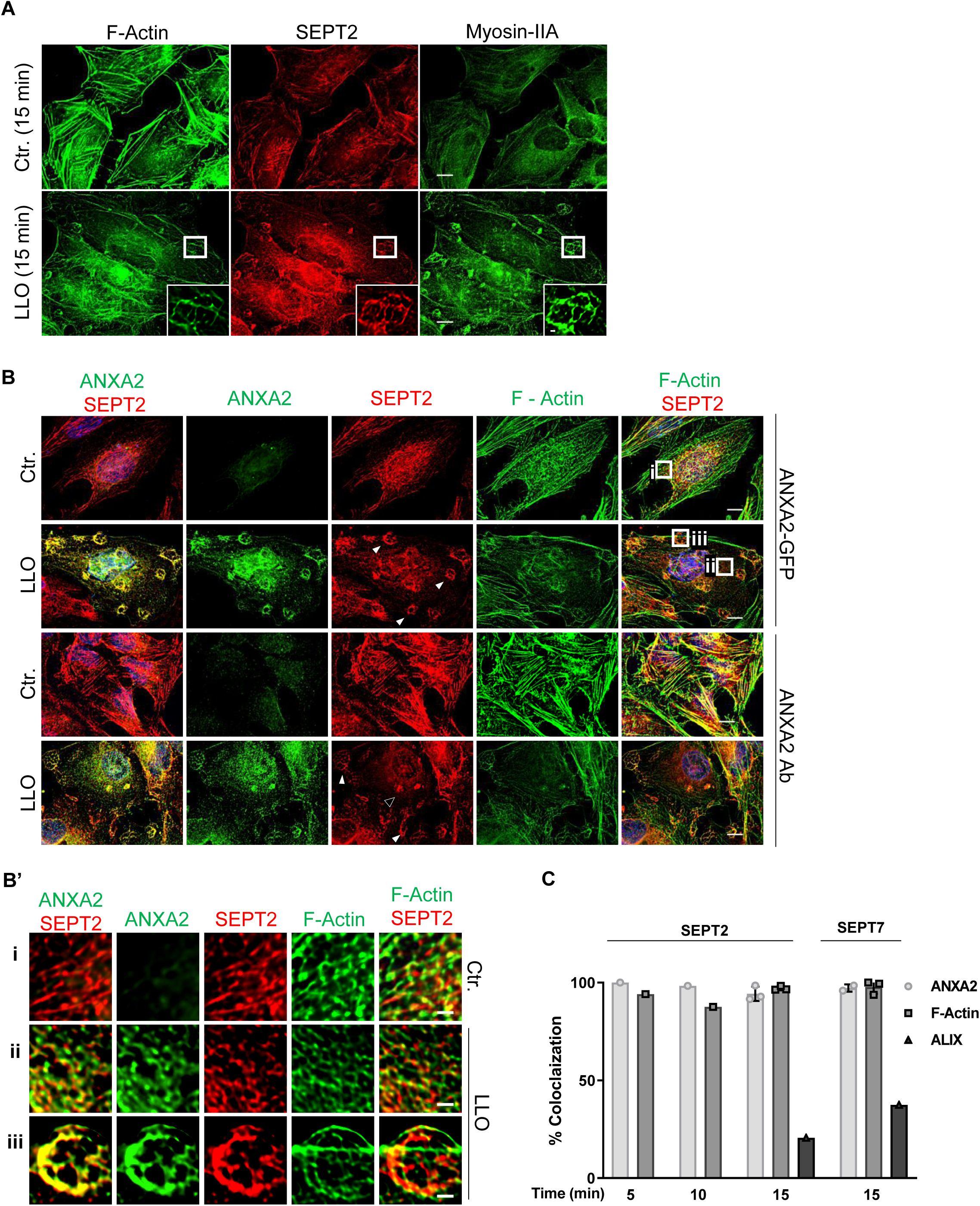
The septin cytoskeleton colocalizes with F-actin, myosin-IIA, and ANXA2 in LLO-perforated cells. HeLa cells were incubated without (Ctr.) or with 0.5 nM LLO for 5-15 min in repair permissive buffer M1 (1.2 mM Ca^2+^). Cells were fixed, permeabilized and fluorescently labeled for SEPT2 (Alexa Fluor 568), myosin IIA (Alexa Fluor 488), F-actin (Alexa Fluor 647), ANXA2 (Alexa Fluor 488), and nuclei (DAPI) for microscopy analyses. (**A**) Representative images of cells labeled for SEPT2, F-Actin, and myosin-IIA at time point 15 min. Scale bars are 10 μm or 1 μm in zoomed regions. (**B, B’**) HeLa cells were transiently transfected, or not, to express ANXA2-GFP. Representative images of cells labeled for SEPT2, F-Actin, and ANXA2 at time point 15 min. Knob and loop structures are indicated by unfilled and filled arrowheads, respectively. Scale bar: 10 μm. (B’) Enlarged regions from B to better visualize the septin colocalization with F-actin and ANXA2-GFP in both loop and non-loop regions. Fluorescence display was enhanced in B’ii. Scale bar: 2 μm. (A, B, B’) All images were acquired by z-stacks widefield microscopy, deconvolved, and presented as the best focus images. (**C**) The percentages of colocalization of the septins with ANXA2, F-actin, and ALIX were measured from a total of 300-750 cells in each experimental condition, N=3 at 15 min, except for ALIX colocalization and 5- and 10-min time points (N=1).

### Septins are redistributed at the cell surface to form circular filaments intertwined with F-actin and are associated with annexin A2 in injured cell

To better establish the subcellular localization of the remodeled septin cytoskeleton relative to the plasma membrane, HeLa cells were transiently transfected to express the fluorescent plasma membrane marker Lck-mTurquoise2 [43]. Cells were exposed to LLO for 15 min at 37°C, then fixed and labeled for F-actin and SEPT2. Images (z-stacks) were acquired by widefield microscopy and deconvolved. As shown in Figure 4A, A’, SEPT2 and F-actin knob and loop structures colocalized with the plasma membrane marker in LLO-injured cells. As a second approach, FCF-treated cells (to increase the formation of the septin structures) were exposed to LLO for 15 min at 37°C, fixed and labeled for SEPT2, F-actin, and ANXA2. Images were acquired with a spinning disk confocal microscope, deconvolved, and displayed with z-depth coding to establish the position of SEPT2 relative to the cell surface. These analyses showed that the SEPT2/F-actin/ANXA2 knob and loop structures are present on the cell surface and often protrude outward of the cell surface (Figure 4B, C, C’). Cells were also imaged by resonant scanning confocal microscopy, deconvolved, and 3D reconstructed allowing to appreciate that the SEPT2/F-actin structures are in close association with ANXA2, forming rings and spirals (Figure 5A). In accordance with the role of the septins in bending actin filaments, the average diameter of the rings and spirals ranged from 2 to 6 μm in diameters and could be as high as 4 μm (Figure 5A). Finally, we performed super resolution microscopy imaging to better resolve the relative distribution of F-actin, SEPT2, and ANXA2 (Figure 5B). HeLa cells were exposed to LLO for 15 min at 37°C, fixed and labeled for F-actin, SEPT2, and ANXA2. Images showed that SEPT2 and F-actin are closely associated forming intertwining filaments of different lengths. ANXA2 distributed as patches or elongated structures that are closely connected to SEPT2 or F-actin and which density increases towards the top of the loop structure. Together, these data show that in response to plasma membrane injury, the septin cytoskeleton is remodeled together with F-actin and ANXA2 into submembranous loops and ring-like structures. Higher resolution imaging indicates that F-actin and the septins form intertwined filaments in close connection with ANXA2.

**Figure 4.**
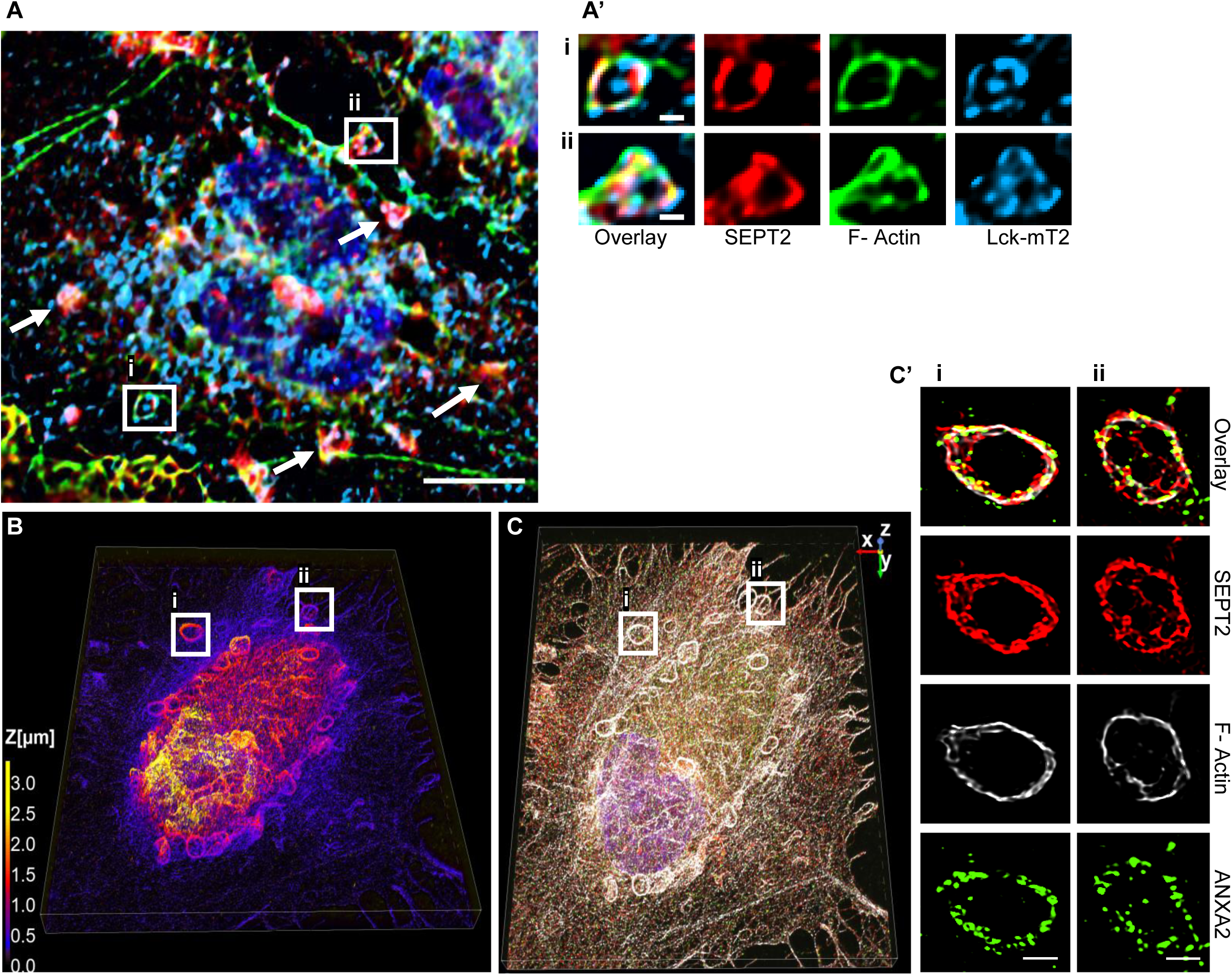
The septin cytoskeleton redistributes with F-actin and ANXA2 in close association with the plasma membrane. (**A, A’**) HeLa cells were transiently transfected to express Lck-mTurquoise2 (Lck-mT2) and incubated with 0.5 nM LLO for 15 min in M1. Cells were fixed, permeabilized, and fluorescently labeled for SEPT2 (Alexa Fluor 568) and F-actin (Alexa Fluor 647). Widefield Z-stack was deconvolved and a single plane is presented. Scale bar is 10 μm. Arrows point to additional SEPT2, F-actin, and Lck colocalization. (Ai) and (Aii) are regions enlarged from (A), scale bars are 1 μm. (**B, C, and C’**) FCF-treated cells were exposed to 0.5 nM LLO for 15 min in M1 and were fixed, permeabilized, and fluorescently labeled for SEPT2 (Alexa Fluor 568), ANXA2 (Alexa Fluor 488), and F-actin (Alexa Fluor 647). Images were acquired by super-resolution spinning disk confocal microscopy. B is a 3-D projection of all deconvolved images (44.99 μm x 56.07μm x 3.4 μm) and overlayed with a z depth code. C is a 3-D overlay of SEPT2, F-actin, and ANXA2. C’ shows the overlay and individual channels of two selected loop structures with an optical depth of 1.4 μm (scale bar, 1 μm).

**Figure 5.**
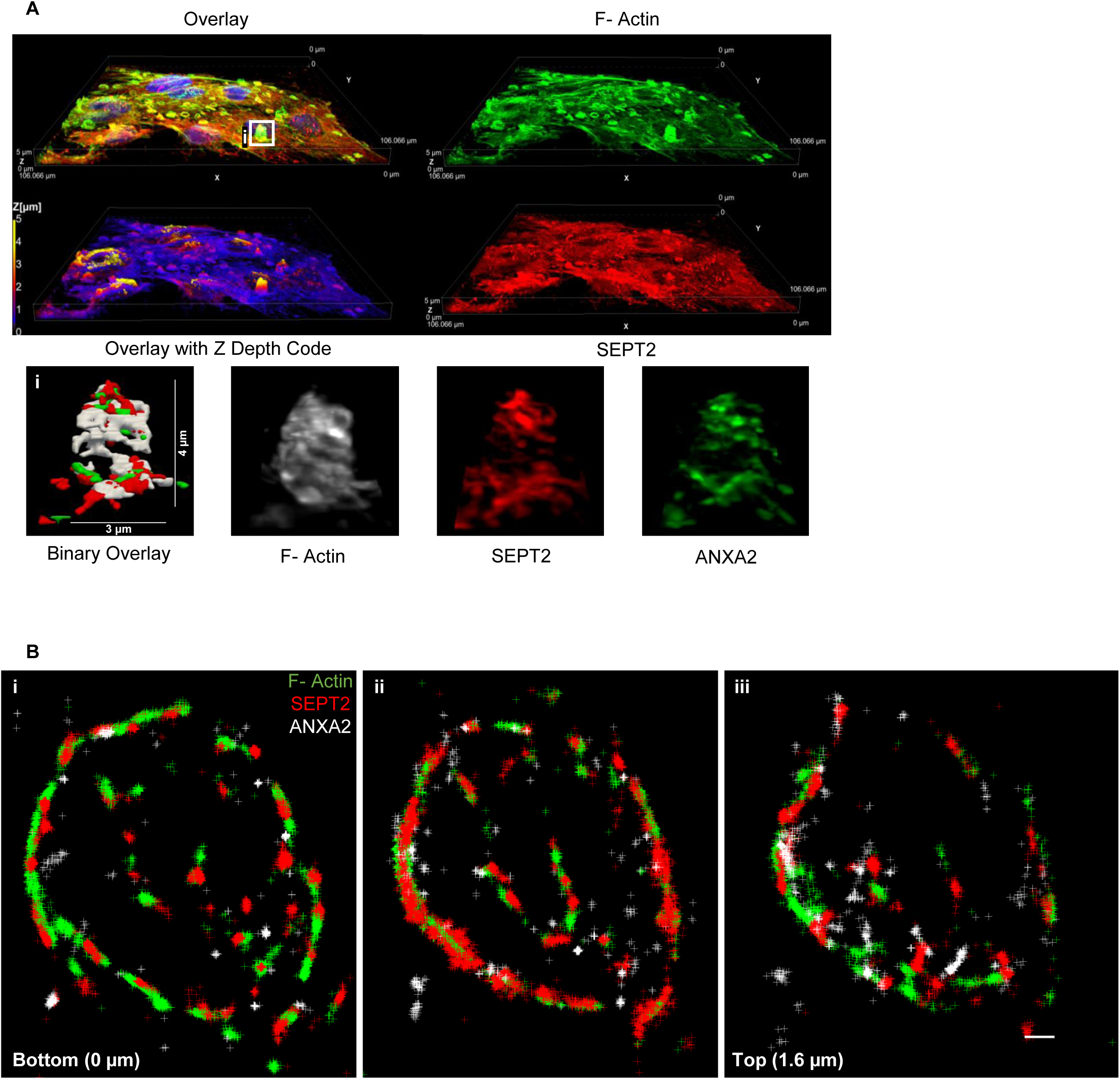
Septins form circular filaments intertwined with F-actin and connected to ANX A2. (**A**) HeLa cells, pre-treated with 100 μM FCF, were incubated with 0.5 nM LLO for 15 min in M1. Cells were fixed, permeabilized, and fluorescently labeled for SEPT2 (Alexa Fluor 568), F-actin (Alexa Fluor 647), and nuclei (DAPI). 3-D representation of individual channels and 3-D overlay with depth coding of a denoised and deconvolved Z-stack acquired by resonant scanning confocal microscopy (106 μm x 106 μm x 5 μm). (**Ai**) Enlarged 3-D representations of A and its binary overlay. (**B**) HeLa cells were incubated with o.5 nM LLO for 15 min in M1. Cells were fixed, permeabilized, and fluorescently labeled for SEPT2 (Alexa Fluor 568), ANXA2 (Alexa Fluor 647) and F-actin (ATTO 488). Images were acquired by super-resolution STORM microscopy (step size 0.2 μm). Each panel (Bi-iii) displays molecules detected within a 0.8 μm optical depth (scale bar is 1 μm).

### Septin redistribution is functionally correlated with plasma membrane repair efficiency

Our data strongly support the role of the septins in forming submembranous structures that promote plasma membrane repair. In this model, septin remodeling should correlate with the plasma membrane repair efficiency, *i.e.* septin remodeling should decrease when plasma membrane repair is prevented in Ca^2+^-free medium, and conversely, septin remodeling should increase when plasma membrane repair is enhanced in the presence of FCF. Consistent with this model, in Ca^2+^-free medium, the formation of septin/F-actin knobs and loop structures were markedly decreased in LLO-perforated cells compared to Ca^2+^-containing medium (Figure 6A, B, C). Similarly, ANXA2 redistribution was impaired in Ca^2+^-free medium (Figure 6A). Quantitative analyses confirmed that the number of septin/F-actin knobs and loops per cell and the percentage of cells presenting such structures were both significantly decreased in LLO-treated cells in Ca^2+^-free medium compared to Ca^2+^-containing medium (Figure 6B, C). Also consistent with this scenario, FCF treatment led to more pronounced formation of septin loops, which all colocalized with F-actin and ANXA2 (Figures 6D). Quantification showed that FCF treatment significantly increased the percentage of cells undergoing septin remodeling and the number of septin/F-actin knobs and loops per cell (Figure 6B, C). Together, these data establish that the redistribution of the septin cytoskeleton is functionally correlated with the plasma membrane repair efficiency, further supporting the role of the septins in plasma membrane repair.

**Figure 6.**
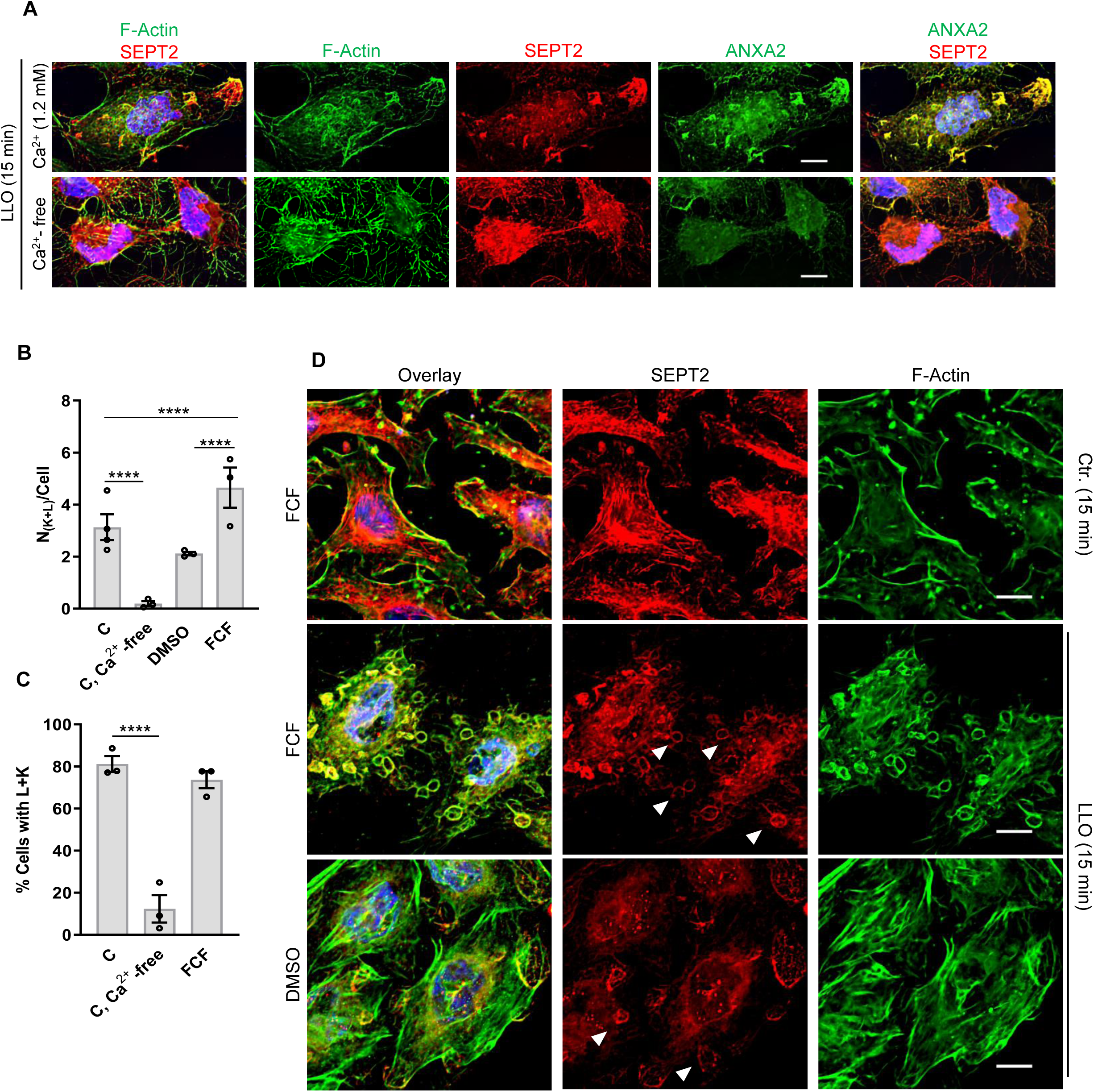
Redistribution of the septin cytoskeleton is functionally correlated with plasma membrane repair efficiency. (**A**) HeLa cells were incubated with 0.5 nM LLO for 15 min in M1 or M2 buffers. Cells were fixed, permeabilized, and fluorescently labeled for SEPT2 (Alexa Fluor 568), ANXA2 (Alexa Fluor 488), F-actin (Alexa Fluor 647), and nuclei (DAPI). Scale bars are 10 μm. (**B, C**) Cells were treated as indicated in (A) and (D). The average number of septin knobs and loops per cell presenting such structures (N_(K+ L)_/Cell) (**B**), and the percentage of cells presenting these structures (% Cells with K+L) (**C**) were enumerated based on septin wide field fluorescence images. A total of 300-750 cells were analyzed in each experimental condition from duplicate wells, N=3 at minimum. Linear mixed-effects models were used for analysis, **** P<0.0001. (**D**) Control untreated (Ctr.), FCF (100 μM) pre-incubated, and vehicle DMSO pre-incubated HeLa cells were incubated with or without 0.5 nM LLO (Ctr.) for 15 min in M1. Equivalent concentrations of FCF and DMSO were maintained during LLO exposure. Cells were fixed, permeabilized, and fluorescently labeled for SEPT2 (Alexa Fluor 568), ANXA2 (Alexa Fluor 488), F-actin (Alexa Fluor 647), and nuclei (DAPI). Images were acquired by widefield microscopy (**A**) or resonant scanning confocal microscopy (**D**) and were denoised, deconvolved, and presented as best-focus projection images. Scale bars are 10 μm.

### The septins control both the remodeling of cortical F-actin into knob and loop structures and ANXA2 recruitment into these structures

We next, silenced SEPT7 expression and labeled cells for F-actin, myosin II, and ANXA2. In SEPT7-deficient cells, the septin cytoskeleton (labeled with anti-SEPT2, 7, or 9 Abs) was markedly disrupted in accordance with the essential role of SEPT7 in the expression of several septin members and formation of septin filaments (Figure 7A, SF 5B, C). This was observed both in the presence and in the absence of LLO. In LLO-injured cells, the formation of F-actin knobs and loops associated myosin II was significantly decreased in SEPT7-deficient cells compared to cells treated with control siRNA (Figure 7A, B, SF6). This last result shows that the septins control the remodeling of cortical F-actin in knob and loop structures, in accordance with the known role of the septins in bending actin filaments [27]. Importantly, the F-actin loops still observed in SEPT7-siRNA-treated cells colocalized with low levels of septins, likely in cells that were less effectively silenced (Figure 7A, B). In SEPT7-deficient cells, ANXA2 distribution was unaffected in the absence of LLO in comparison to control siRNA-treated cells (SF 5C). However, there was a dramatic decrease in ANXA2 remodeling in SEPT7-deficient cells injured by LLO in comparison to SEPT7-proficient cells (Figure 7A). Three-dimensional quantification of ANXA2 specks formation showed that ANXA2 remodeling was dramatically and significantly decreased in LLO-injured SEPT7-deficient cells compared to control siRNA-treated cells (Figure 7C). Conversely, in ANXA2-deficient cells, septin and F-actin remodeling were unaffected in both control and LLO-injured cells (Figure 7D, SF 5D). These data show that ANXA2 remodeling in LLO-injured cells requires the formation of septin/F-actin knobs and loops. Collectively, our data support a novel model in which the septin cytoskeleton promotes plasma membrane repair by playing a key role in organizing membrane domains containing the actomyosin cytoskeleton and ANXA2.

**Figure 7.**
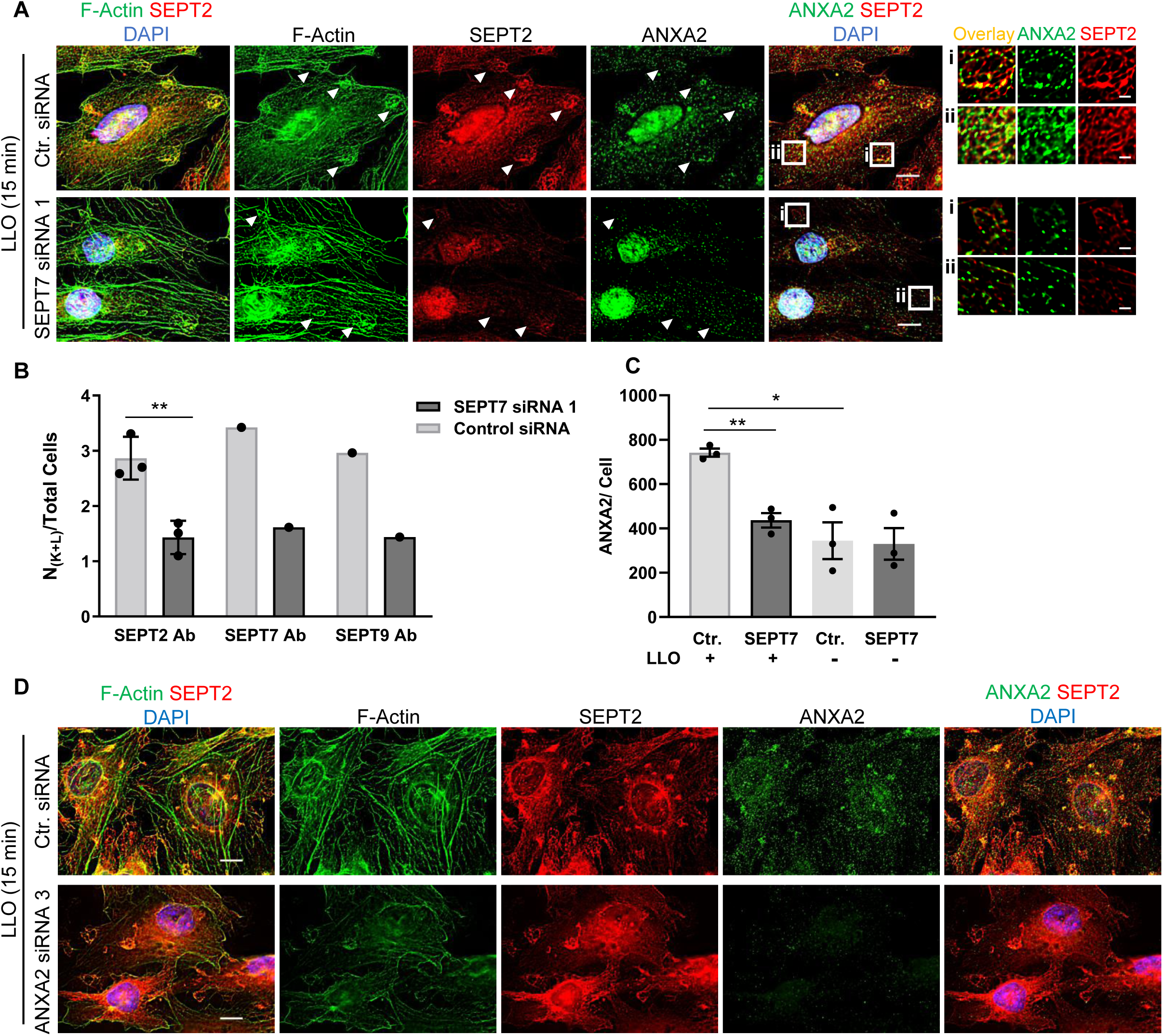
Redistribution of F-actin and ANXA2 is dependent on SEPT7 expression. **(A)** HeLa cells transfected with Ctr. siRNA or SEPT7-siRNA 1 were incubated with 0.5 nM LLO for 15 min in M1. Cells were fixed, permeabilized, and labeled for SEPT2 (Alexa Fluor 568), F-Actin (Alexa Fluor 647), ANXA2 (Alexa Fluor 488), and nuclei (DAPI). Presented images are the best-focus projections generated from deconvolved widefield Z-stacks and are displayed with the same intensity scaling for each fluorescence setting. Scale bars are 10 μm. (**Ai**) and (**Aii**) are regions enlarged from (A), scale bars are 2 μm. (**B**) Same experiment as in (A) but SEPT7 and SEPT9 were also labeled with fluorescent Abs. The number of knobs and loops per total cells (N_K+L_/ Total cells) were enumerated based on F-actin labeling in cells fluorescently labeled with anti-SEPT2, -SEPT7, and -SEPT9 antibodies. A total of 150-300 cells were analyzed from duplicate wells, per experimental condition. N=3 for SEPT2 Ab labeling and N=1 for SEPT7 and SEPT9 Ab labeling. (**C**) HeLa cells were transfected with Ctr. siRNA or with SEPT7-siRNA 1 (SEPT7) and incubated with 0.5 nM LLO (+), or not (-), for 15 min in M1. Cells were fixed, permeabilized, and labeled for SEPT2 (Alexa Fluor 568 Abs), F-Actin (Alexa Fluor 647), ANXA2 (Alexa Fluor 488-Abs), and nuclei (DAPI). Fluorescence images (z-stacks) were acquired by widefield microscopy. The ANXA2 images were deconvolved and the number of ANXA2 specks were automatically counted in 3D by the software. At least 100 cells were analyzed for each experimental condition, N=3. (B and C) Data are expressed as average ± SEM and a Students Paired T Test was used, *P<0.05 and ** P< 0.01. (**D**) HeLa cells were transfected with Ctr. siRNA or with ANXA2-siRNA 3 and incubated with 0.5 nM LLO, or not (Ctr.), for 15 min in M1. Cells were labeled indicated in (A). The presented images are the best-focus projections from deconvolved z-stacks acquired by widefield microscopy and are displayed with the same intensity scaling for each fluorescence setting. Scale bars are 10 μm.

## DISCUSSION

This work establishes that the septin cytoskeleton is required for effective plasma membrane repair of cells perforated by CDC pore-forming toxins and mechanical wounding. This is a novel role for the septins cytoskeleton. Collectively, our data support a model in which the septin cytoskeleton acts as a scaffold to promote the formation of plasma membrane repair domains containing contractile F-actin and annexin A2.

Septins are conserved GTP-binding proteins divided into four groups based on sequence homologies (SEPT 2, 3, 6, and 7), each group includes several interchangeable members except for SEPT7, the only member of its group [44]. Silencing SEPT7 and SEPT6 had a significant deleterious effect on plasma membrane repair, whereas silencing of SEPT2 or SEPT9 had no effect (Figure1, SF 2). This result can be explained by the fact that SEPT7 silencing disrupts the whole septin cytoskeleton due to: (i) the essential, non-redundant function of SEPT7 in septin hetero-oligomer and filament formation, and (ii) the degradation of numerous septin members in SEPT7-deficient cells (Figure 7, SF 2B-E, and SF 5) [29, 36]. The septin-6 group consists of septin-6, 8, 10, 11 and 14, therefore, the defect in plasma membrane repair of SEPT6-deficient cells suggests that septin 6 displays unique properties, which cannot be rescued by other members of its subgroup, during plasma membrane repair.

Due to the central role of septin 7 in the assembly of the septin cytoskeleton, septin 7 silencing added to the use of the drug forchlorfenuron (FCF) that specifically alters septin functions, and fluorescence microscopy analyses of several septins (SEPT2, 7, 9), provided a comprehensive array of experimental approaches to study the role of the septin cytoskeleton in plasma membrane repair. Our data show that the septin cytoskeleton is extensively redistributed during plasma membrane repair (Figure 2, SF4) and this redistribution requires extracellular calcium (Figure 6A, B, C), a hallmark of plasma membrane repair mechanisms [1, 45]. To date, there is no calcium-dependent mechanism known to control the organization of the septin cytoskeleton and the septins are not known to directly bind calcium ions, therefore calcium likely indirectly controls septin remodeling. In injured cells, the septins formed noticeable structures that we described as knobs and loops, proximal to the plasma membrane and that can protrude outward from the cell surface (Figures 4, 5). The knobs may be an early stage of the loop structure formation based on the timing of their respective appearance (Figure 2C, SF 4C). The size of the loops ranges from 2 to 6 µm in diameter (Figures 4, 5). The subplasmalemmal localization of the redistributed septins supports their role in plasma membrane repair. Strikingly, there was a significant increase in septin knob and loop formation in FCF-treated cells paralleled with a significant increase in plasma membrane repair efficiency, further confirming the role of the septins in plasma membrane repair (Figure 6B-D). FCF is a drug that specifically associates with the septin GTP-binding pocket, mimicking a nucleotide-bound state [46]. FCF was proposed to disrupt septins’ functions [37] but was also shown to facilitate their assemblies [38, 39]. Therefore, mechanisms by which FCF alters the septin dynamics are still not completely understood and need further elucidation. Our data clearly show that FCF treatment enhances the formation and/or stabilizes septin loops during plasma membrane repair.

During plasma membrane repair, the remodeled septins closely associate with F-actin, myosin IIA, and ANXA2 throughout the cell surface and within the septin knobs and loops (Figures 2 to 5 and SF 4 to 6). Super-resolution microscopy allowed for visualization of intertwined F-actin and septin filaments in the septin loops (Figure 5), in accordance with the role of septins in bending actin filaments [25]. The septins mostly run parallel to the F-actin and, at times, appear perpendicular as if interconnecting actin fibers (Figure 5). Collectively, our data are in accordance with recent literature that showed that in mammalian cells, septins do form filaments that preferentially associate with the contractile F-actin/myosin II cytoskeleton [44]. The same study also showed that septin filaments can anchor the actin cytoskeleton to the lipid membranes [44]. Super-resolution microscopy revealed that ANXA2 is organized in patches closely connected to both F-actin and septin filaments and are more abundant and elongated at the top of the structure (Figure 5). There is no known direct interaction between the septins and ANXA2 and it is unclear if septins and ANXA2 are directly or indirectly associated. However, similar to the septins, ANXA2 is known to interact with F-actin and phospholipids. Whether the septins and/or the ANXA2 play a role in anchoring the cortical actomyosin network to the plasma membrane during plasma membrane repair is to be determined. Silencing SEPT7 led to a strong and significant decrease in the formation of cortical F-actin/myosin II knobs and loops, whereas silencing ANXA2 had no noticeable effect (Figure 7). These results strongly support that the septins, not ANXA2, are critical in the formation of the subplasmalemmal actomyosin structures.

ANXA2 is a Ca^2+^-sensor critical for plasma membrane repair of CDC (perfringolysin O)- injured cells and was also identified in our screen as promoting plasma membrane integrity (Table 1)[18, 42]. ANXA2 was proposed to repair the plasma membrane of perfringolysin O-injured cells by facilitating the shedding of microvesicles [18]. In another model, F-actin-driven constriction was proposed to either close the membrane wounds or to promote the excision of the edges of the ruptured plasma membrane in mechanically wounded cells [41, 47]. It was also proposed that ANXA2, the protein S100A11, and cortical F-actin act together to repair mechanically ruptured cells by excision [41]. Collectively, (i) the literature, (ii) the identification of ANXA2 in the plasma membrane repair screen (Table 1), and (iii) the colocalization of ANXA2 with F-actin/myosin II and septins in the subplasmalemmal structures that form during plasma membrane repair, strongly support that the septins play a key role in organizing plasma membrane repair domains. This is further supported by the fact that silencing SEPT7 led to significant decrease in formation of ANXA2 specks in injured cells, whereas silencing ANXA2 did not affect septin/actomyosin remodeling during plasma membrane repair (Figure 7). Therefore, septins play a key structural role. Together, these findings support a model in which the septin cytoskeleton acts as a scaffold for the formation of domains containing F-actin, myosin IIA and ANXA2. Based on the known role of ANXA2 and contractile F-actin in plasma membrane repair, the septin-organized domains likely act as repair platform by sequestering and/or removing (shedding or excision) the damaged plasma membrane. Septin filaments are scaffolding proteins that interact with multiple cellular components including inositol phosphates, SNARE machinery, clathrin, actin, motor proteins, and tubulin. Through these interactions, the septins control multiple processes that encompass signaling, membrane fusion, membrane curvature control, endocytosis, and exocytosis. [25, 29, 48]. The detailed molecular mechanisms by which the septins facilitate membrane repair in coordination with the actomyosin cytoskeleton and ANXA2 remains to be elucidated.

The septins are known to organize and bend the actomyosin cytoskeleton during cytokinesis [27, 49, 50] and ANXA2 is required for cytokinesis [51]. Therefore, septin-dependent reorganization of the cortical F-actin network and ANXA2 during cytokinesis and during plasma membrane repair likely share common mechanisms.

Our screen indicates that membrane fusion, lysosome exocytosis, annexins- and ESCRT-III play a role in the repair of LLO-injured cells. Therefore, it is possible that several repair mechanisms act together. In particular, silencing CHMP2 and ANXA7 (Table 1) negatively affected plasma membrane integrity, supporting that the ESCRT-III machinery may also repair LLO-injured cells by shedding microvesicles [23, 34]. However, ALIX (SF 5), and CHMP2 (not shown) which are components of ESCRT-III machinery, poorly colocalized with the septin/actomyosin/ANXA2 structures during plasma membrane repair. These findings suggest that plasma membrane repair likely involves several mechanisms which together maintain cell integrity. Whether these repair mechanisms coexist, interact, or operate in different time scales will need to be established.

*L. monocytogenes* is a facultative intracellular pathogen, which intracellular replication is essential for pathogenesis. Paradoxically, this bacterium uses the pore-forming toxin LLO as its major virulence factor, yet infected cells remain viable to support bacterial replication [12, 52–54]. Viability of infected cells results from the combination of (i) LLO intrinsic properties that decrease its lifetime [13, 16, 55, 56], and (ii) plasma membrane repair [14, 15]. The septins were previously shown to regulate *L. monocytogenes* internalization and form a cage-like structure around cytosolic bacteria [48]. Our data highlight a novel function of the septins in the repair of LLO-perforated host cells and show that LLO redistributes the septin cytoskeleton. It will be important to establish if the roles of the septins in bacterial uptake and in their cytosolic entrapment are influenced by LLO-induced host cell perforation.

In conclusion, our findings support a general role for the septins in plasma membrane repair by showing that the repair of two distinct injuries require SEPT7 expression: (i) perforation by transmembrane toxin pores and (ii) mechanical damages (Figure 1, SF 3F, G). This opens novel research avenues on the role of the septin cytoskeleton in maintaining plasma membrane integrity relevant to many physiological and pathological conditions.

## MATERIALS and METHODS

### Cell culture and reagents

Our studies used HeLa cells which have been extensively used to identify plasma membrane repair mechanisms. HeLa cells (ATCC, # CCL-2™) and HeLa Histone 2B-GFP (HeLa H2B-GFP, EMD Millipore SCC117) were cultured in Dulbecco’s Modified Eagles Medium (DMEM, Gibco #11995) containing 10% heat-inactivated fetal bovine serum (HIFBS), 100 units/ml penicillin and 100 μg/ml streptomycin at 37°C and 5% CO_2_ atmosphere. Repair permissive buffer (M1) (Hanks Balanced Salt Solution (HBSS, Gibco #14175) supplemented with 0.5 mM MgCl_2_, 1.26 mM CaCl_2_, 25 mM glucose, and 10 mM HEPES) and repair restrictive buffer (M2) (HBSS supplemented with 0.5 mM MgCl_2,_ 25 mM glucose and 10 mM HEPES) buffers were used for the repair assays. For lentiviral production, Lenti-X™ 293T Cell Line (Takara) was cultured in 90% DMEM with high glucose (4.5 g/L), 4 mM L-glutamine, and 10% FBS; 100 units/ml penicillin G sodium, and 100 µg/ml streptomycin sulfate. When appropriate, the cell culture medium contained the Tet-system-approved FBS (Takara). Recombinant six-histidine-tagged LLO and PLY was purified from *E. coli* as previously described [12]. The hemolytic activity of LLO and PLY were measured as previously described [12]. In all of our experimental conditions, cells are viable and fully recover from LLO injury [12, 14, 33]. LLO was used at 0.5 nM in all experiments unless otherwise indicated. The stock solution of Forchlorfenuron (FCF) (Sigma, 32974) was prepared in DMSO at 50 mM. FCF is used at 100 μM and DMSO is used at a 1:500 dilution for a vehicle control. Anti-SEPT2 (Protein Tech, mouse clone 2F3B2, for Western Blot; Atlas, Rabbit polyclonal for imaging), anti-SEPT6 (Protein Tech rabbit polyclonal for Western Blot and imaging), anti-SEPT7 (Protein Tech rabbit polyclonal for Western Blot and imaging), anti-SEPT9 (Atlas, rabbit polyclonal for Western Blot and imaging), anti-annexin A2 (Invitrogen, mouse clone Z014 polyclonal for Western Blot and imaging), anti-Alix (Biolegend, clone 3A9), anti-myosin-IIA (Abnova, clone 2B3), anti-tubulin (Sigma, clone B-5-1-2), Alexa Fluor 488 conjugated Phalloidin (Invitrogen, #A12379), Alexa Fluor 647 conjugated Phalloidin (Invitrogen, #A22287), and ATTO 488 conjugated Phalloidin (ATTO-TEC) were the primary antibodies used for immunoblotting and fluorescence microscopy experiments. The secondary antibodies used were anti-mouse IgG horseradish peroxidase (HRP)-conjugated (Cell Signal Technology, #7076S), anti-rabbit IgG (HRP)-conjugated (Cell Signal Technology, #7074S), anti-mouse IgG Alexa Fluor-conjugated (Invitrogen), anti-rabbit IgG Alexa Fluor-conjugated (Invitrogen). 10 kDa lysine fixable Ruby dextran (ThermoFisher D1817) and 10 kDa lysine fixable Emerald dextran (ThermoFisher D1820) were used for the mechanical wounding assay. The vector Lck-mTurquoise2, in the backbone pEGFP-C1, was a gift from Dorus Gadella (Addgene #98822) (doi: https://doi.org/10.1101/160374). The vector encoding annexin A2-GFP, in the backbone pEGFP-N3, was a gift from Volker Gerke & Ursula Rescher (Addgene # 107196) [57]. EZ-Tet-pLKO-Puro (Addgene #85966) was a gift from Cindy Miranti and both psPaX2 (Addgene #12260), and pMD2.G (Addgene #12259) were gifts from Didier Trono [58]. Cells were transiently transfected using Lipofectamine 2000 (Invitrogen) and Opti MEM according to the manufacturer’s instruction.

### siRNA screen

A siRNA screen was performed in 96-well tissue culture-treated clear bottom black plate (Corning). HeLa H2B-GFP were plated at 7x10^3^ cells/well (DMEM, 10% HIFBS) and reverse transfected with 1 pmol/well of a combination of 3 non-overlapping 11-mer siRNAs (Ambion Silencer® Select, Thermofisher) using Lipofectamine RNAiMax transfection reagent (Invitrogen) and Opti-MEM Reduced Serum Medium (Gibco) according to the manufacturer’s instructions. For the list of targeted genes and siRNA sequences, see Supplemental Table 1. To account for potential variations in cell growth during the 72-h treatment with the various siRNAs, control cells were plated at three different concentrations (7x10^3^, 6.3 x10^3^, and 5.6 x10^3^ cells/well) in each plate and reverse transfected with 1 pmol of non-targeting Silencer Select negative control siRNA No. 1 (Invitrogen). All experimental conditions were carried out in quadruplicates. To assess membrane resealing, siRNA-treated cells were washed twice with 37°C M1 and replaced with M1 containing 1 μM TO-PRO-3 iodide (ThermoFisher) or washed once with 37°C M2 containing 5 mM ethylene glycol-bis(2-aminoethylether)-N, N, N’, N’, tetraacetic acid (EGTA), followed by a wash with M2, and replaced with M2 containing 1 μM TO-PRO-3 iodide (ThermoFisher). A SpectraMax i3X multimode detection platform equipped with a MiniMax cytometer (Molecular Devices) was used to obtain fluorescent kinetic readings and 4X magnification images (transmitted light, and green and red fluorescence; 535 nm and 638 nm excitations and 617 nm 665 nm emissions respectively). Pre-kinetic images were acquired as previously described [32]. The cell culture plate was cooled to 4°C for 5 min and ice-cold LLO and TO-PRO-3 iodide were added at 0.5 nM and 1 uM final concentrations, respectively. The plates were then placed into the spectrofluorometer for the 30 min kinetic assay at 37°C, then post-kinetic fluorescence images were acquired as described previously [32]. The total number of cells in each well was enumerated based on their nuclear fluorescence (H2B-GFP), pre- and post-kinetic. We found that the silencing of 217 proteins resulted in cell numbers ranging from 85-115% relative to control cells. Gene knockdown (KD) resulting in cell densities outside of this range were excluded (10 gene KD with cell counts >115% and 18 with cell counts <85% of control cells were excluded) (Supplemental Table 2). Importantly, cells did not detach during the assay as the average cell count ratio, defined as the cell count post-kinetic relative to the cell count pre-kinetic, was unaffected in all samples, and across all siRNA- and LLO-treatment conditions was 1.0061 ± 0.0065 (average ± SEM). Control wells with the closest cell count to the specific knockdown condition were chosen for comparison. TO-PRO-3 fluorescence intensities were compared between cells treated with specific-siRNA and control-siRNA treated cells. The average TO-PRO-3 fluorescence intensities at time point 30 min were log-transformed and then analyzed with ANOVA to establish the list of candidate genes with positive or negative effects, respectively, in plasma membrane repair.

### SDS-PAGE and Immunoblotting

HeLa cells were lysed in 150 mM NaCl, 20 mM Tris HCl, 2 mM EDTA, NP40 1% and protease inhibitors (Roche # 04693132001) pH 7.4. Samples of cells lysates were used for BCA (BioRad) protein assay. Lysates were boiled in reducing Laemmli buffer and subjected to SDS-PAGE electrophoresis followed by transfer to PVDF membranes. Membranes were labeled with the indicated antibodies (septin or annexin) and then stripped (2.35 % SDS, 73.5 mM Tris HCl pH 6.8, 134.6 mM beta-mercaptoethanol) and re-labeled with tubulin as a loading control. The knockdown efficiencies were measured using Image Lab (BioRad) software. Quantification included septin (or annexin) normalization to their respective loading controls.

### Generation of doxycycline-inducible SEPTIN7 shRNA HeLa cell line

Annealed oligonucleotides (non-overlapping with the 3 siRNAs used in the screen) were cloned into the vector EZ-Tet-pLKO-Puro using the restriction sites NheI and EcoRI [58]. The resulting construct was validated by sequencing. Oligonucleotide and primer sequences are presented in Supplemental Table 3. For lentiviral production, Lenti-X™ 293T cells were transfected with the vectors pMD2.G, psPAX2, and EZ-Tet-pLKO-Puro using Lipofectamine 2000 and Opti MEM according to the manufacturer’s instructions (Invitrogen). Lentiviruses were collected, HeLa cells were transduced and selected using 1 µg/ml puromycin. Isolated colonies were pooled and further cultured. Stable HeLa-D_i_SEPT7-shRNA1 was cultured with 50 ng/ml doxycycline (DOX) for 72 h and SEPT7 expression was measured by immunoblotting (Figure 1E, SF 3A).

### Plasma membrane repair assays

All plasma membrane repair assays (cells treated with siRNAs, shRNAs, or pharmacological agents) were based on the quantification of TO-PRO-3 fluorescence intensity. Cells were plated in 96-or 24-well plates and treated with siRNA and/or drugs (doxycycline, DMSO, or FCF) for the indicated times prior to the assay. Cells were washed twice with 37°C M1 and incubated in M1 containing 1.5 μM TO-PRO-3. For calcium-free conditions, cells were washed once with M2 (5 mM EGTA) followed by a wash with M2 at 37°C and incubated with M2 containing 1.5 μM TO-PRO-3. Cells were then incubated with or without 0.5 nM LLO or 2 nM PLY. In some experiments, FCF (100 μM) or DMSO vehicle (1/500 dilution) was added during LLO exposure for up to 30 min. TO-PRO-3 fluorescence intensity was measured in the temperature-controlled plate reader (Spectra Max i3X Multi-Mode Detection Platform, Molecular Devices) or by quantitative fluorescence microscopy as indicated in the Figure Legends.

### Mechanical Wounding Assay (adapted from [59])

Cells were cultured on 35 mm round coverglasses and transfected with siRNAs (control, SEPT7 siRNA 1, ANXA2 siRNA 3). After 72 h, cells were washed twice with M1 at 37°C, or washed once with M2 (5 mM EGTA) followed by a wash with M2 at 37°C. The coverglasses were placed on a silicone O-ring and damaged by gently rolling glass beads in the presence of 1.5 mg/ml Emerald Dextran to label the damaged cells, followed by 5 min incubation at 37°C. The Emerald Dextran and glass beads were then washed with M1 or M2 buffer. Ruby dextran (2 mg/ml) was added to the cells for 2 min at 37°C to label the “not recovered” cells. Cells were washed with M1 or M2 and fixed. Coverglasses were mounted on glass slides and immediately imaged on the NikonTi2-E microscope with the 40x air objective. 50 images were acquired per slide. Emerald-fluorescent cells were enumerated as the total number of damaged cells and cells double-fluorescent for Ruby and Emerald were counted to represent the not recovered cells. Data are expressed as the percentage of not recovered cells relative to cells incubated in Ca^2+^-free medium in each experimental condition.

### Fluorescence labeling

Cells were plated in a 24-well plate glass slides. Cells were exposed to 0.5 nM LLO or not for 5-15 min in M1 or M2 at 37°C. In some experiments, FCF (100 μM) or DMSO vehicle (1: 500) was added during LLO exposure. Cells were treated as indicated, washed twice in PHEM buffer (60 mM PIPES, 25 mM HEPES, 2 mM MgCl_2_,10 mM EGTA; pH 6.9), fixed in 4% PFA in PBS for 20 min at RT, and permeabilized for 5 min with 0.1% Triton X-100 in PBS. Blocking was performed at RT for 1 h in PBS 10% serum and cells were labeled with antibodies diluted in PBS, 10% serum for 1 h (except for septin Abs, 2 h). Due to its low expression level, SEPT6 labeling was very faint and was not carried out. Nuclei were labeled with DAPI. All primary antibodies were non fluorescent and secondary antibodies were Alexa Fluor-conjugated. F-actin was labeled with Alexa Fluor-conjugated phalloidin. Figure legends indicate the fluorochromes used in each experiment. Fluorescence images were acquired in grey scale and were color-coded. Widefield images were acquired using a 60x water objective and confocal images were acquired using a 60x oil objective.

### Super-resolution Fluorescence labeling

Cells, plated on a glass bottom 35 mm dish, were exposed to 0.5 nM LLO for 15 min in M1 at 37°C. Cells were washed twice in PHEM buffer, fixed in 3% PFA and 0.1% glutaraldehyde in PHEM for 10 min at RT and permeabilized for 15 min with 0.2% Triton X-100 in PBS. Blocking was performed by incubating cells at RT for 90 min in PBS with 10% serum and 0.05% Triton X-100, and labeling by incubating cells with antibodies diluted in PBS, 5% serum and 0.05% Triton X-100 for 1 h for all labeling. After labeling with primary Abs (anti-ANXA2, anti-SEPT2 Abs, and ATTO 488-conjugated Phalloidin) and secondary antibodies (Alexa Flour 568 and Alexa Fluor 647), cells were fixed again in 3% paraformaldehyde and 0.1% glutaraldehyde in PHEM for 10 min at RT. Protocol was modified from [60]. Cells were imaged in a STORM imaging buffer that was prepared on ice right before imaging (Buffer A: 50 mM Tris pH 8.0 with 10 mM NaCl and 10% glucose mixed with Buffer B: 0.5 mg/mL glucose oxidase and 40 µg/mL catalase in 50 mM Tris pH 8 10 mM NaCl, 1% 2-mercaptoethanol and 20 mM Cysteamine).

### Microscopes, image acquisitions, and analyses

*<u>Widefield microscopy</u> (Nikon)*. Images were acquired using a Nikon Ti2-E microscope equipped with a temperature, humidity, and CO_2_-controlled blackout enclosure. Ten excitation wavelengths (Spectra III pad 365, 440, 488, 514, 561, 594, 640, and 730 nm) and emission filter sets for DAPI (435 ≤ emission ≤ 485 nm), CFP (460 ≤ emission ≤ 500 nm), FITC (515 ≤ emission ≤ 555 nm), YFP HYQ (from 520 ≤ emission ≤ 560 nm), Cy3 (from 573 ≤ emission ≤ 648 nm) and Texas Red HYQ (573 ≤ emission ≤ 648 nm) imaging with a high-speed wheel; back-thin illuminated SCMOS camera (Orca-Fusion BT, Hamamatsu) of resolution of 5.3 Megapixels; nanopositioning Piezo Sample Scanner (Prior). The automated stage allows multi-positioning (x-y) for high-content screening involving multi-well cell culture plates. The objectives include: 10x air Plan Apo lambda (0.45NA), 20x air S Plan Fluor ELWD (0.45 NA), 40x air S Plan Fluor ELWD 40x (0.6 NA), 40x water immersion (1.25NA), 40x air Plan Apo lambda (0.95 NA), a 60x water immersion Plan Apo IR (1.27NA). The imaging system has a “smart” auto-focus. NIS Elements AR Software was used for image analysis and NIS Elements HC Software for image acquisition. *Images were randomly acquired* from duplicate wells in each experimental condition. The number of independent experiments (N) is indicated in figure legends. For each field of view, 17 to 30 planes were acquired with a step size of 0.3 nm using a 60X water immersion objective. Image deconvolution was done using the Richardson-Lucy algorithm and the best-focused projection image was created using the extended depth of focus module. *<u>Widefield microscopy</u> (Zeiss).* Images were acquired on a motorized, inverted, wide-field fluorescence microscope (Axio Observer D1, TempModule S, heating unit XL S; Zeiss) equipped with a PZ-2000 XYZ automated stage, 20x Plan Neofluar (numerical aperture [NA] = 0.5), 40x Plan Neofluar (NA = 1.3), and 63× Plan Apochromat (NA = 1.4) objectives, a high-speed Xenon fluorescence emission device (Lambda DG-4, 300 W; Sutter Instrument Company), a Lambda 10-3 optical emission filter wheel for the fluorescence imaging, a SmartShutter to control the illumination for phase-contrast and DIC imaging (Sutter Instrument Company), an ORCA-Flash 4.0 sCMOS camera (Hamamatsu). The filter sets (Chroma Technology Corporation) were as follows: DAPI (49000), Alexa Fluor 488 (49002), Alexa Fluor 568 (49005), and Cy5 (49006). Images were acquired and analyzed using MetaMorph imaging software (Molecular Devices). *<u>SoRa confocal microscope</u> (Nikon)*. Nikon Ti2-E microscope equipped with the Yokogawa CSU-W1 spinning disk with a 50 µm pinhole size (4000 rpm); a 60X Plan Apo lambda D OFN25 DIC N2 (NA 1.42) oil objective; and Quest Camera (apparent pixel size is 0.027 µm/pixel) from Hamamatsu. *<u>Resonant laser scan confocal</u> <u>A1R plus</u> (Nikon)*. Nikon Ti2-E microscope equipped with Plan Apo lambda 60x oil (NA 1.4), DU4 detector, pinhole size 39.59. For confocal images, 17-30 planes were acquired per field of view with a 0.2 nm step size. Images were denoised with denoise.ai (artificial intelligence) and deconvolved using the Richardson-Lucy algorithm (NIS Elements, Nikon). *<u>N-STORM System</u>. Stochastic Optical Reconstruction Microscopy (STORM) (Nikon).* Nikon Ti2-E microscope equipped with a SR HP Apo TRIF 100xH Oil objective (NA 1.49), an ORCA Fusion BT sCMOS camera (Hamamatsu), and a LUD-H series laser unit (405 nm, 488 nm, 561 nm, and 640 nm). The XY resolution is 20 nm and the Z resolution is 50 nm. Images (z stacks) were acquired using 3D-STORM with a 0.2 µm step size. *<u>3D ANXA2 Counting.</u>* Z stack images were acquired by widefield fluorescence microscopy (60x water objective, 0.3 um steps). Images were denoised with denoise.ai (artificial intelligence) and deconvolved using the Richardson-Lucy algorithm (NIS Elements, Nikon). The cellular area excluding the nuclei was selected. The “count 3D objects” tool was then used in the Nikon NIS software to automatically count the 3D objects which represent the total number of ANXA2 specks.

### Statistical analyses

For most of the data such as experiments with cell counts and fluorescence intensity as the measurements, log transformation was first used to reduce skewness and variation. ANOVA or linear mixed-effects models were then used for analyses depending on whether the data was independent or with repeated measurements over time. For the screening data, a p-value<0.05 without adjustment for multiple comparisons was used as the cutoff because the results were further validated in the subsequent assays. A p-value<0.05 for a single test or after adjustment for multiple comparisons with Holm’s procedure was considered significant. Additionally, GraphPad Prism software was used to perform Students Paired T -Tests and a p-value<0.05 was considered significant.

## ABBREVIATIONS

ASM: Acid sphingomyelinase
ALIX: ALG-2-interacting protein X
ANXA2: Annexin A2
Ab: Antibody
CDC: cholesterol-dependent cytolysins
DOX: Doxycycline
ESCRT: endosomal sorting complexes required for transport
EGTA: ethylene glycol-bis(2-aminoethylether)- N,N,N’,N’,tetraacetic acid
FCF: Forchlorfenuron
H2B- GFP: green-fluorescent histone 2B
K: Knobs
KD: knockdown
LLO: Listeriolysin O
L: Loops
MAC/PF: Membrane Attack Complex/Perforin
N: Number
R: Ratio
PFA: Paraformaldehyde
PI: Propidium Iodide
RT: Room Temperature
SEPT: Septins
shRNA: short hairpin RNA
siRNA: silencing RNA
SEM: Standard error of the mean
SF: Supplemental Figure
TX-100: and Triton X-100

## ACKNOWLEDGMENTS

Research reported in this publication was supported by the Institute of Allergy and Infectious Diseases of the National Institutes of Health under award number R01AI157205. The content is solely the responsibility of the authors and does not necessarily represent the official views of the National Institutes of Health. Viviana Ruiz received financial public funding from the Colombian Ministry of Science, Technology, and Innovation (Ministerio de Ciencia, Tecnología e Innovación, MinCiencias). We acknowledge resources from the Campus Microscopy and Imaging Facility (CMIF) and the OSU Comprehensive Cancer Center (OSUCCC) Microscopy Shared Resource (MSR), The Ohio State University. This facility is supported in part by grant P30 CA016058, National Cancer Institute, Bethesda, MD. We also acknowledge the precious and kind help received from Dr. Anthony Vetter for assisting with fluorescence microscopy imaging and analysis using the NIS Elements software.

**Supplemental Movies 1A, and 1B**. **HeLa cell morphology and viability post-LLO exposure**. H2B GFP HeLa cells were exposed to 0.5 nM LLO (**1A**), or not (**1B**), for 30 min 1.2 mM Ca^2+^ HBSS. After 30 min, the media were changed to DMEM with 10% HIFBS and cells were imaged (bright field and fluorescence imaging) on the microscope stage at 37°C, 5% CO_2_ every 10 min for 12 h. Scale bar is 10 μm. Four fields of view were acquired for each experimental condition and representative movies were shown. Movies show that all LLO-treated cells maintain the same morphology and behavior as non-treated cells.

**Supplemental Figure 1 Kinetic of plasma membrane integrity recovery after LLO-injury**. HeLa cells were exposed, or not, to 0.5 nM LLO in M1 or M2 for 5 min on ice to allow LLO to bind the plasma membrane. Cells were washed and warmed up to 37°C for 5, 10, 15, 30 min, 2h, and 24 h. In the last min of incubation, cells were exposed for 1 min to 100 µM propidium iodide. About 1300 cells were analyzed per experimental condition and data are expressed as the average nuclear Propidium Iodide fluorescence intensities expressed in arbitrary units (AI) ± SEM of at least N=3. N=4 for the 5-30 min conditions. *P < 0.05. Data show that plasma membrane integrity is recovered fully between 30 min to 2 h, at the whole population level.

**Supplemental Figure 2.** (**A**) HeLa cells were transfected with Ctr.-, SEPT2-, or SEPT9-siRNAs (Supplemental Table 1). After 72 h, cells were lysed and analyzed by SDS-PAGE and immunoblotting for septin and tubulin expression. Serial dilutions of control- (100% to 6.25%) and undiluted septin- (100%) siRNA treated cell lysates were loaded in the gel to facilitate the quantification of the KD efficiencies. Blots are representative of at least N=5. KD efficiencies were >90%. (**A’**) HeLa cells were treated for 72 h with ctr. or SEPT-siRNAs and were exposed to LLO (0.5 nM) for 30 min in M1 containing TO-PRO-3. Data are the average TO-PRO-3 fluorescence intensities expressed in arbitrary units (AI) ± SEM of at least N=3 at time point 30 min. (**B** to **E**) HeLa cells were transfected with Ctr.- or SEPT-siRNAs as in (A). Blots are representative of at least N=5. (**C**) Comparing control siRNA-treated cells to SEPT7-siRNA-treated cells, SEPT6 expression was reduced by 44.8% ± 24.4; 70% ± 21.2; and 52.3% ± 19.6 for siRNA1, 2, and 3, respectively, SEPT2 expression was reduced by 59.6% ± 4.3; 46.4% ± 7.7; and 40.1% ± 6.9 for siRNA1, 2, and 3, respectively. SEPT 9 expression was reduced by at least 90% for all three siRNAs. KD of SEPT2, 6, and 9 did not appear to affect the expression of the other tested septins.

**Supplemental Figure 3.** (**A**) D_i_SEPT7-shRNA1 stable HeLa cells were cultured for 48 and 72 h with DOX (50 μg/ml) or without DOX (-). Cell lysates were analyzed by SDS-PAGE and immunoblotting. The percentage of septin expression of Dox-treated cells is expressed relative to cells cultured in the absence of DOX. (**B**) The hemolytic assay showed that the presence of Dox at the indicated concentrations does not affect LLO pore formation. Note that in the repair assay, DOX-treated cells were washed three times, so only traces of DOX were present in the medium during the repair assay. (**C**, **D**) HeLa cells were pre-treated with the indicated concentrations of FCF, or corresponding DMSO dilutions, or left untreated (Ctr.) for 16 h. Equivalent concentrations of FCF and DMSO were maintained in the buffer during the repair assay. Cells were exposed to LLO for 30 min in M1 or M2 supplemented with TO-PRO-3. Data are expressed as the average TO-PRO-3 fluorescence intensities in arbitrary unit of N=2 to N=4 (each with 4 to 8 technical replicates) ± SEM at each time point. (**E**) The hemolytic assay showed that FCF at the highest concentration does not affect LLO pore formation. (**F** and **G**) HeLa cells were treated for 72 h with Ctr. siRNA or SEPT7-siRNA 1 and were exposed to PLY for 30 min in M1 (1.2 mM Ca^2+^) (F and G) or M2 (Ca^2+^-free) (G), supplemented with TO-PRO-3. Data are the average TO-PRO-3 fluorescence intensity expressed in arbitrary units (AI) ± SEM of at least N=4 at time point 30 min (F) and representative kinetic curves (G). A Students Paired T Test was used, ** P< 0.01. (**H**) HeLa cells were transfected with Ctrl siRNA or ANXA2-siRNAs (Supplemental Table 1). After 72 h, cells were lysed. Control cell lysates (100%) and annexin A2 siRNA-treated cell lysates (100%) were loaded in the gel to facilitate the quantification of the KD efficiencies. The blot is representative of 6 independent experiments (N=6) for ANXA2 siRNA 3, and N=1 for ANXA2 siRNA 1 and siRNA 2.

**Supplemental Figure 4.** HeLa cells were incubated without (Ctr.) or with 0.5 nM LLO for 5-15 min in M1. Cells were chemically fixed, permeabilized, and fluorescently labeled with anti-SEPT9, or SEPT7, or SEPT2 primary Abs and Alexa Fluor 568-conjugated secondary Abs, Alexa Fluor 488-conjugated phalloidin, and DAPI. (**A** and **B**) SEPT7 (A) and SEPT9 (B) fluorescence images are presented with the same intensity scaling showing the loss of septin association with actin stress fibers. To better visualize septin and actin filaments, selected regions were enlarged (Ai-Aiii) and the septin fluorescence display was the same for all images except for Aii which intensity was amplified. All images were acquired by z-stack widefield microscopy (0.3 μm steps), deconvolved, and presented as the best focus images, except for Aa and Ab images which are single planes focused on actin stress fibers. Septin knobs and loops are indicated by unfilled and filled arrowheads, respectively. Scale bars are 10 μm and 2 μm in Ai-Aiii. (**C**) The numbers of septin knobs or loops per cell which present such structures (N _(K or L)_/Cells) were enumerated at the indicated time points based on septin fluorescence images. A total of 300-750 cells were analyzed in each experimental condition from duplicate wells, N=3 for the 15 min time point, and N=1 for 5- and 10-min time points.

**Supplemental Figure 5.** (**A**) Cells were fluorescently labeled for SEPT2 (Alexa Fluor 568), ALIX (Alexa Fluor 488), and nuclei (DAPI) at time point 15 min. Scale bars: 10 μm. (**B - D**) HeLa cells were transfected with Ctr. siRNA, SEPT7-siRNA 1 (B and C), or ANXA2-siRNA 3 (D) for 72 h. Cells were incubated without LLO for 15 min at 37°C in M1 buffer. Cells were fixed, permeabilized and labeled with anti-SEPT primary Ab (Alexa Fluor 568-conjugated secondary Ab), Alexa Fluor 488-conjugated phalloidin (F-Actin), DAPI (nuclei), and (B-D) anti-ANXA2 primary Ab (Alexa Fluor 647-conjugated secondary Ab). Scale bars: 10 μm. (A, C, D) Z-stack images were acquired by widefield microscopy (0.3 μm steps), deconvolved, and displayed as best-focus projection images, except for B which are single plane images focused on actin stress fibers. **C** and **D** correspond to Figure 7A and 7D respectively.

**Supplemental Figure 6.** HeLa cells were transfected with Ctr. siRNA or SEPT7-siRNA 1. Cells were incubated with or without (Ctr.) 0.5 nM LLO for 15 min in M1. Cells were fixed, permeabilized, and labeled with anti-Myosin-IIA primary Ab (Alexa Fluor 647-conjugated secondary Ab), Alexa Fluor 488-conjugated phalloidin (F-Actin), DAPI (nuclei). Z-stack images were acquired by widefield microscopy (0.3 μm steps), deconvolved, and displayed as best-focus projection images. Scale bars: 10 μm.

## Literature Cited

1. Jimenez, A.J. and F. Perez, Plasma membrane repair: the adaptable cell life-insurance. Curr Opin Cell Biol, 2017. 47: p. 99–107.

2. Cooper, S.T. and P.L. McNeil, Membrane Repair: Mechanisms and Pathophysiology. Physiol Rev, 2015. 95(4): p. 1205–40.

3. Dias, C. and J. Nylandsted, Plasma membrane integrity in health and disease: significance and therapeutic potential. Cell Discov, 2021. 7(1): p. 4.

4. Bouillot, S., E. Reboud, and P. Huber, Functional Consequences of Calcium Influx Promoted by Bacterial Pore-Forming Toxins. Toxins (Basel), 2018. 10(10).

5. Banerji, R., et al., Pore-forming toxins of foodborne pathogens. Compr Rev Food Sci Food Saf, 2021. 20(3): p. 2265–2285.

6. Barisch, C., J.C.M. Holthuis, and K. Cosentino, Membrane damage and repair: a thin line between life and death. Biol Chem, 2023.

7. Andrade, L.O., Plasma membrane repair involvement in parasitic and other pathogen infections. Curr Top Membr, 2019. 84: p. 217–238.

8. Thapa, R., S. Ray, and P.A. Keyel, Interaction of Macrophages and Cholesterol-Dependent Cytolysins: The Impact on Immune Response and Cellular Survival. Toxins (Basel), 2020. 12(9).

9. Ayyar, B.V., et al., CLIC and membrane wound repair pathways enable pandemic norovirus entry and infection. Nat Commun, 2023. 14(1): p. 1148.

10. Osborne, S.E. and J.H. Brumell, Listeriolysin O: from bazooka to Swiss army knife. Philos Trans R Soc Lond B Biol Sci, 2017. 372(1726).

11. Petrisic, N., et al., The molecular mechanisms of listeriolysin O-induced lipid membrane damage. Biochim Biophys Acta Biomembr, 2021. 1863(7): p. 183604.

12. Vadia, S., et al., The pore-forming toxin listeriolysin O mediates a novel entry pathway of L. monocytogenes into human hepatocytes. PLoS Pathog, 2011. 7(11): p. e1002356.

13. Seveau, S., Multifaceted activity of listeriolysin O, the cholesterol-dependent cytolysin of Listeria monocytogenes. Subcell Biochem, 2014. 80: p. 161–95.

14. Vadia, S. and S. Seveau, Fluxes of Ca2+ and K+ are required for the listeriolysin O-dependent internalization pathway of Listeria monocytogenes. Infect Immun, 2014. 82(3): p. 1084–91.

15. Cassidy, S.K., et al., Membrane damage during Listeria monocytogenes infection triggers a caspase-7 dependent cytoprotective response. PLoS Pathog, 2012. 8(7): p. e1002628.

16. Chen, C., et al., The Listeriolysin O PEST-like Sequence Co-opts AP-2-Mediated Endocytosis to Prevent Plasma Membrane Damage during Listeria Infection. Cell Host Microbe, 2018. 23(6): p. 786–795 e5.

17. Dunstone, M.A. and R.K. Tweten, Packing a punch: the mechanism of pore formation by cholesterol dependent cytolysins and membrane attack complex/perforin-like proteins. Curr Opin Struct Biol, 2012. 22(3): p. 342–9.

18. Ray, S., R. Roth, and P.A. Keyel, Membrane repair triggered by cholesterol-dependent cytolysins is activated by mixed lineage kinases and MEK. Sci Adv, 2022. 8(11): p. eabl6367.

19. Idone, V., et al., Repair of injured plasma membrane by rapid Ca2+-dependent endocytosis. J Cell Biol, 2008. 180(5): p. 905–14.

20. Mellgren, R.L., et al., Calpain is required for the rapid, calcium-dependent repair of wounded plasma membrane. J Biol Chem, 2007. 282(4): p. 2567–75.

21. Tam, C., et al., Exocytosis of acid sphingomyelinase by wounded cells promotes endocytosis and plasma membrane repair. J Cell Biol, 2010. 189(6): p. 1027–38.

22. Draeger, A., K. Monastyrskaya, and E.B. Babiychuk, Plasma membrane repair and cellular damage control: the annexin survival kit. Biochem Pharmacol, 2011. 81(6): p. 703–12.

23. Jimenez, A.J., et al., ESCRT machinery is required for plasma membrane repair. Science, 2014. 343(6174): p. 1247136.

24. Mostowy, S. and P. Cossart, Septins: the fourth component of the cytoskeleton. Nat Rev Mol Cell Biol, 2012. 13(3): p. 183–94.

25. Benoit, B., C. Pous, and A. Baillet, Septins as membrane influencers: direct play or in association with other cytoskeleton partners. Front Cell Dev Biol, 2023. 11: p. 1112319.

26. Shuman, B. and M. Momany, Septins From Protists to People. Front Cell Dev Biol, 2021. 9: p. 824850.

27. Nakamura, M., J. Hui, and S.M. Parkhurst, Bending actin filaments: twists of fate. Fac Rev, 2023. 12: p. 7.

28. Longtine, M.S., et al., The septins: roles in cytokinesis and other processes. Curr Opin Cell Biol, 1996. 8(1): p. 106–19.

29. Spiliotis, E.T. and K. Nakos, Cellular functions of actin- and microtubule-associated septins. Curr Biol, 2021. 31(10): p. R651–R666.

30. Kinoshita, M., Assembly of mammalian septins. J Biochem, 2003. 134(4): p. 491–6.

31. Abbey, M., et al., GTPase domain driven dimerization of SEPT7 is dispensable for the critical role of septins in fibroblast cytokinesis. Sci Rep, 2016. 6: p. 20007.

32. Lam, J.G.T., C. Song, and S. Seveau, High-throughput Measurement of Plasma Membrane Resealing Efficiency in Mammalian Cells. J Vis Exp, 2019(143).

33. Lam, J.G.T., et al., Host cell perforation by listeriolysin O (LLO) activates a Ca(2+)-dependent cPKC/Rac1/Arp2/3 signaling pathway that promotes Listeria monocytogenes internalization independently of membrane resealing. Mol Biol Cell, 2018. 29(3): p. 270–284.

34. Sonder, S.L., et al., Annexin A7 is required for ESCRT III-mediated plasma membrane repair. Sci Rep, 2019. 9(1): p. 6726.

35. Satoh, H., et al., The penta-EF-hand domain of ALG-2 interacts with amino-terminal domains of both annexin VII and annexin XI in a Ca2+-dependent manner. Biochim Biophys Acta, 2002. 1600(1-2): p. 61–7.

36. Kremer, B.E., T. Haystead, and I.G. Macara, Mammalian septins regulate microtubule stability through interaction with the microtubule-binding protein MAP4. Mol Biol Cell, 2005. 16(10): p. 4648–59.

37. Iwase, M., et al., Forchlorfenuron, a phenylurea cytokinin, disturbs septin organization in Saccharomyces cerevisiae. Genes Genet Syst, 2004. 79(4): p. 199–206.

38. Hu, Q., W.J. Nelson, and E.T. Spiliotis, Forchlorfenuron alters mammalian septin assembly, organization, and dynamics. J Biol Chem, 2008. 283(43): p. 29563–71.

39. DeMay, B.S., et al., Cellular requirements for the small molecule forchlorfenuron to stabilize the septin cytoskeleton. Cytoskeleton (Hoboken), 2010. 67(6): p. 383–99.

40. Walker, J.A., et al., Molecular cloning, characterization, and complete nucleotide sequence of the gene for pneumolysin, the sulfhydryl-activated toxin of Streptococcus pneumoniae. Infect Immun, 1987. 55(5): p. 1184–9.

41. Jaiswal, J.K., et al., S100A11 is required for efficient plasma membrane repair and survival of invasive cancer cells. Nat Commun, 2014. 5: p. 3795.

42. Koerdt, S.N., A.P.K. Ashraf, and V. Gerke, Annexins and plasma membrane repair. Curr Top Membr, 2019. 84: p. 43–65.

43. Zacharias, D.A., et al., Partitioning of lipid-modified monomeric GFPs into membrane microdomains of live cells. Science, 2002. 296(5569): p. 913–6.

44. Martins, C.S., et al., Human septins organize as octamer-based filaments and mediate actin-membrane anchoring in cells. J Cell Biol, 2023. 222(3).

45. McNeil, P.L., Repairing a torn cell surface: make way, lysosomes to the rescue. J Cell Sci, 2002. 115(Pt 5): p. 873–9.

46. Angelis, D., et al., In silico docking of forchlorfenuron (FCF) to septins suggests that FCF interferes with GTP binding. PLoS One, 2014. 9(5): p. e96390.

47. Bement, W.M., C.A. Mandato, and M.N. Kirsch, Wound-induced assembly and closure of an actomyosin purse string in Xenopus oocytes. Curr Biol, 1999. 9(11): p. 579–87.

48. Robertin, S. and S. Mostowy, The history of septin biology and bacterial infection. Cell Microbiol, 2020. 22(4): p. e13173.

49. Russo, G. and M. Krauss, Septin Remodeling During Mammalian Cytokinesis. Front Cell Dev Biol, 2021. 9: p. 768309.

50. Mavrakis, M., et al., Septins promote F-actin ring formation by crosslinking actin filaments into curved bundles. Nat Cell Biol, 2014. 16(4): p. 322–34.

51. Benaud, C., et al., Annexin A2 is required for the early steps of cytokinesis. EMBO Rep, 2015. 16(4): p. 481–9.

52. Czuczman, M.A., et al., Listeria monocytogenes exploits efferocytosis to promote cell-to-cell spread. Nature, 2014. 509(7499): p. 230–4.

53. Goldfine, H., et al., Membrane permeabilization by Listeria monocytogenes phosphatidylinositol-specific phospholipase C is independent of phospholipid hydrolysis and cooperative with listeriolysin O. Proc Natl Acad Sci U S A, 1995. 92(7): p. 2979–83.

54. Goldfine, H., S.J. Wadsworth, and N.C. Johnston, Activation of host phospholipases C and D in macrophages after infection with Listeria monocytogenes. Infect Immun, 2000. 68(10): p. 5735–41.

55. Schuerch, D.W., E.M. Wilson-Kubalek, and R.K. Tweten, Molecular basis of listeriolysin O pH dependence. Proc Natl Acad Sci U S A, 2005. 102(35): p. 12537–42.

56. Rogers, S., R. Wells, and M. Rechsteiner, Amino acid sequences common to rapidly degraded proteins: the PEST hypothesis. Science, 1986. 234(4774): p. 364–8.

57. Rescher, U., N. Zobiack, and V. Gerke, Intact Ca(2+)-binding sites are required for targeting of annexin 1 to endosomal membranes in living HeLa cells. J Cell Sci, 2000. 113 (Pt 22): p. 3931–8.

58. Frank, S.B., V.V. Schulz, and C.K. Miranti, A streamlined method for the design and cloning of shRNAs into an optimized Dox-inducible lentiviral vector. BMC Biotechnol, 2017. 17(1): p. 24.

59. Defour, A., S.C. Sreetama, and J.K. Jaiswal, Imaging cell membrane injury and subcellular processes involved in repair. J Vis Exp, 2014(85).

60. Dempsey, G.T., et al., Evaluation of fluorophores for optimal performance in localization-based super-resolution imaging. Nat Methods, 2011. 8(12): p. 1027–36.

