## Supplemental Figures for "The Septin Cytoskeleton is Required for Plasma Membrane Repair"

Supplemental Figure 1

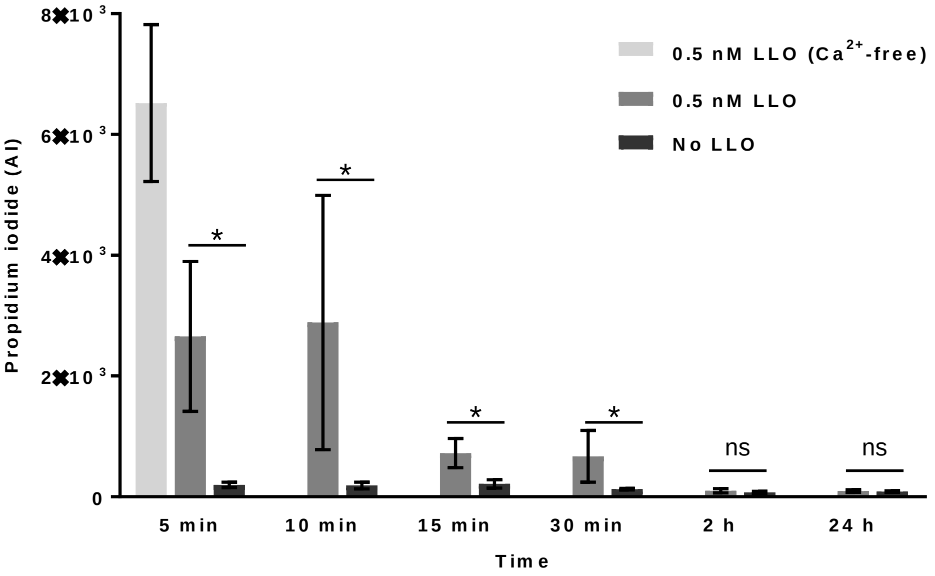

Supplemental Figure 2

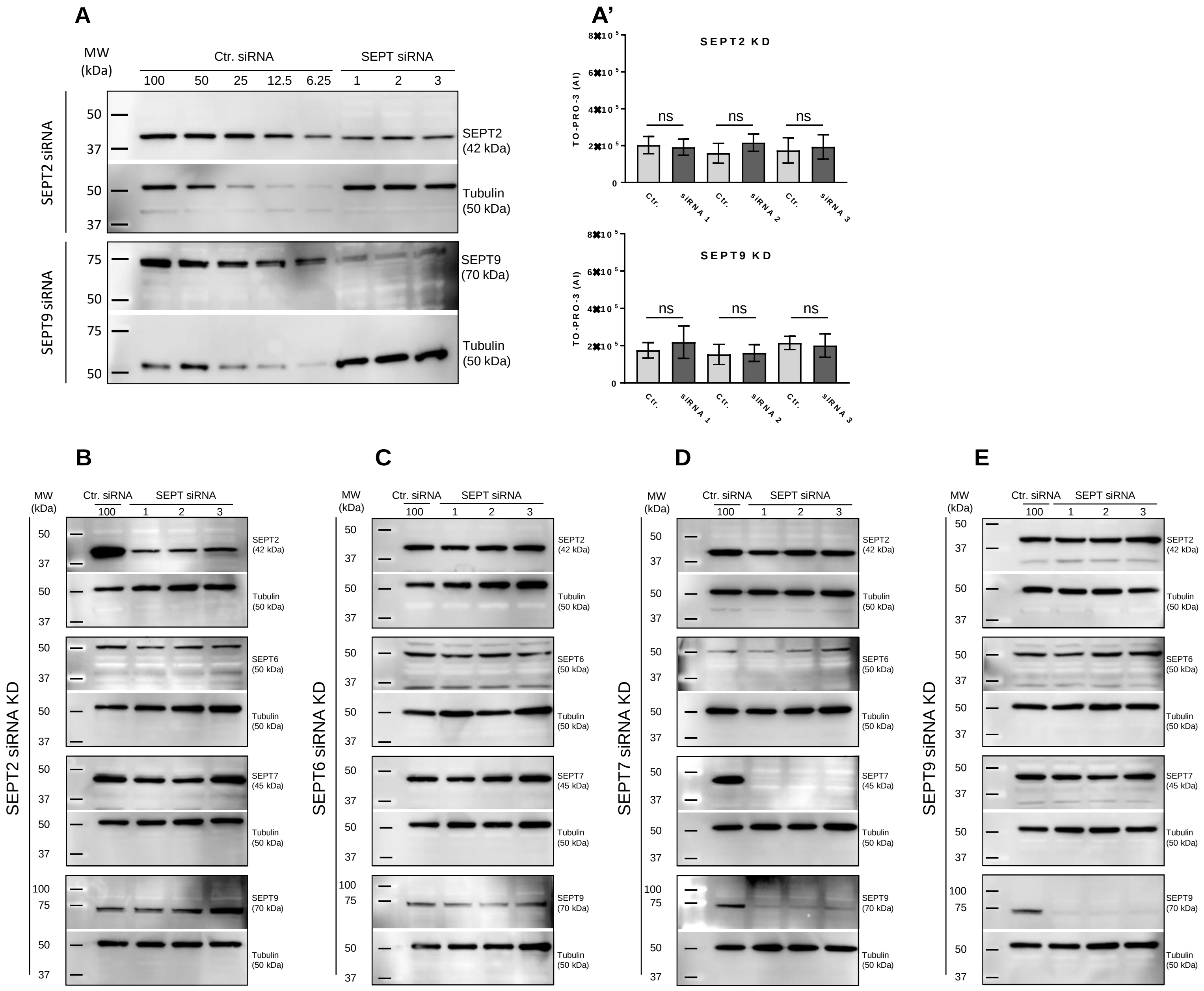

### Supplemental Figure 3

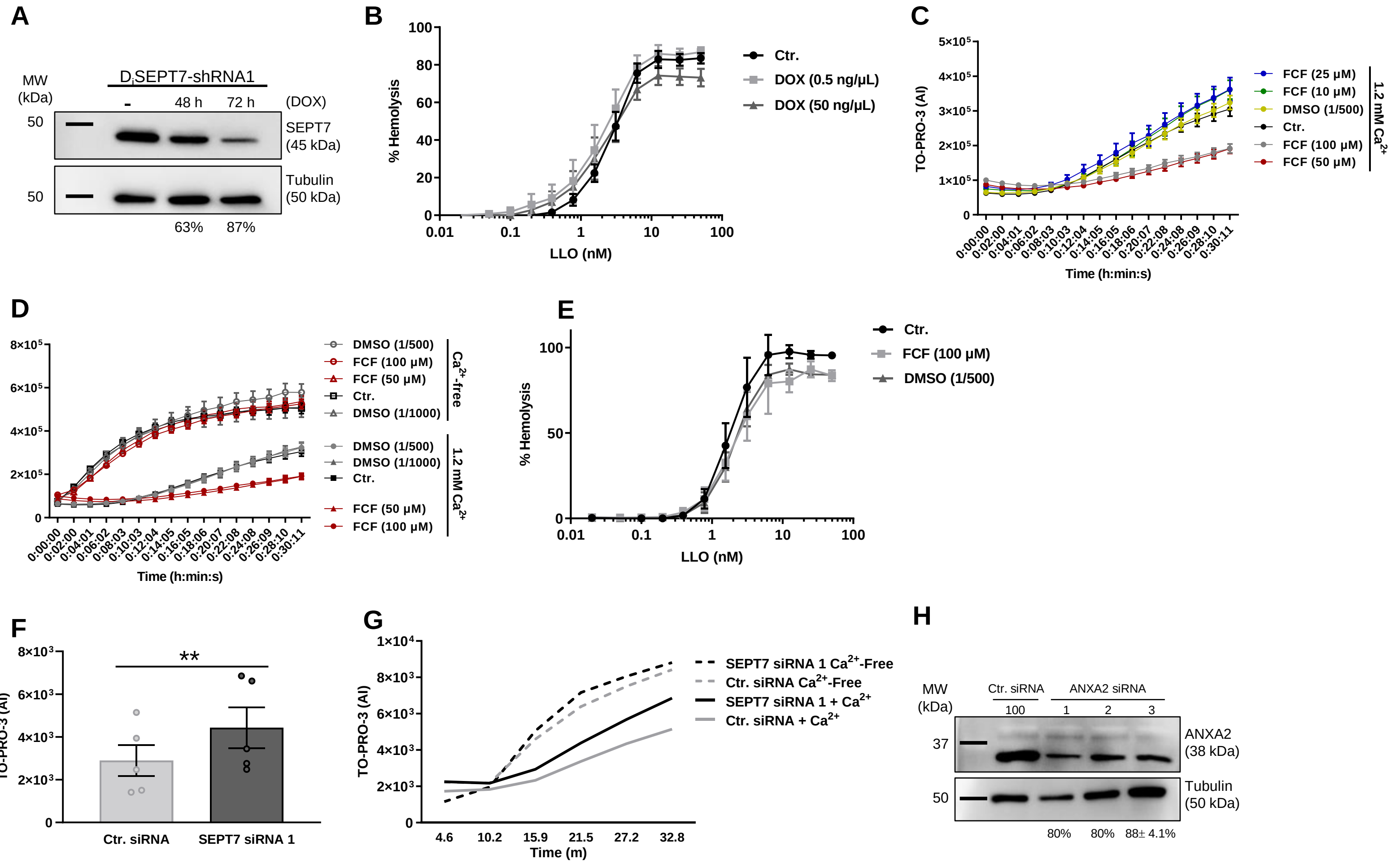

Supplemental Figure 4

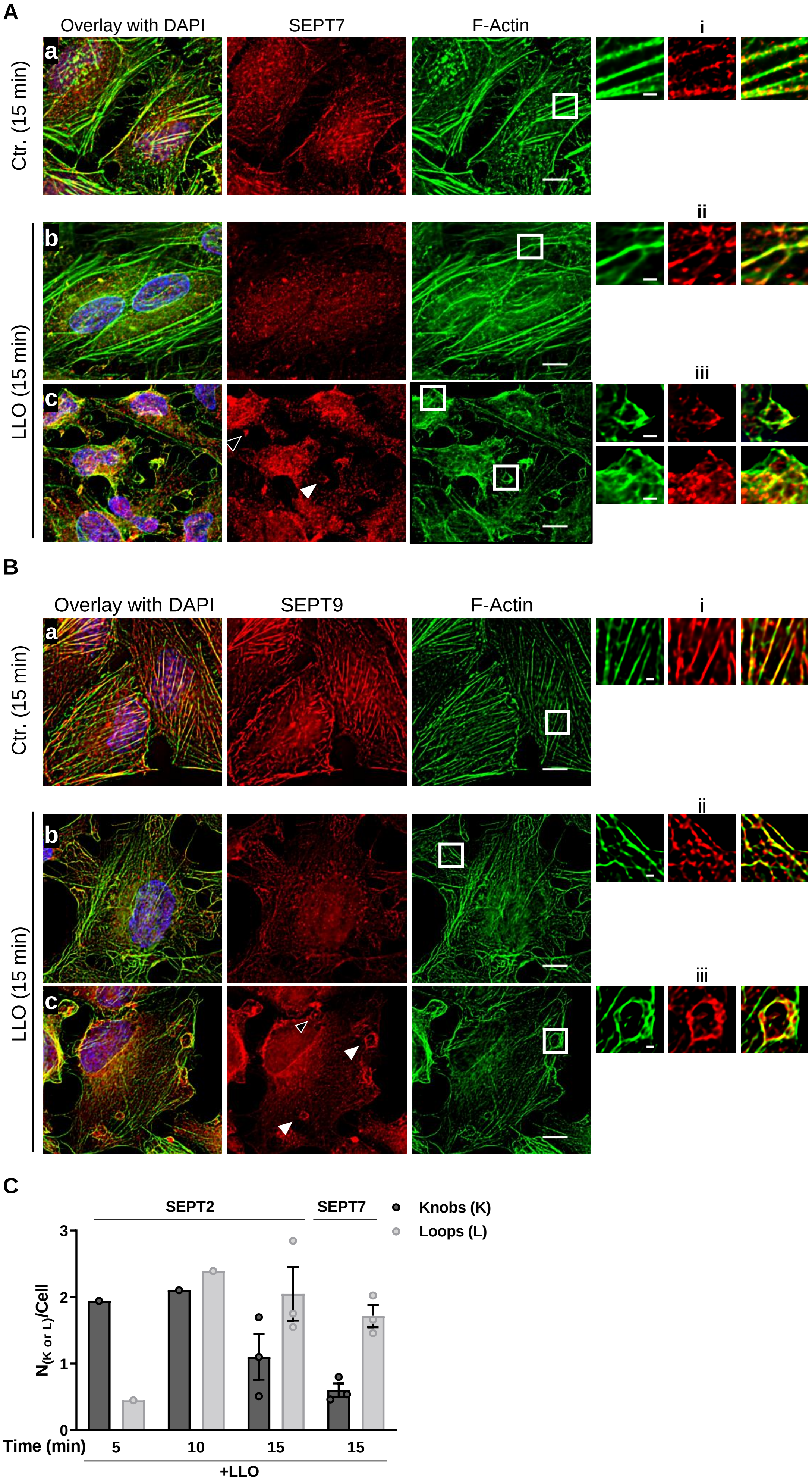

Supplemental Figure 5

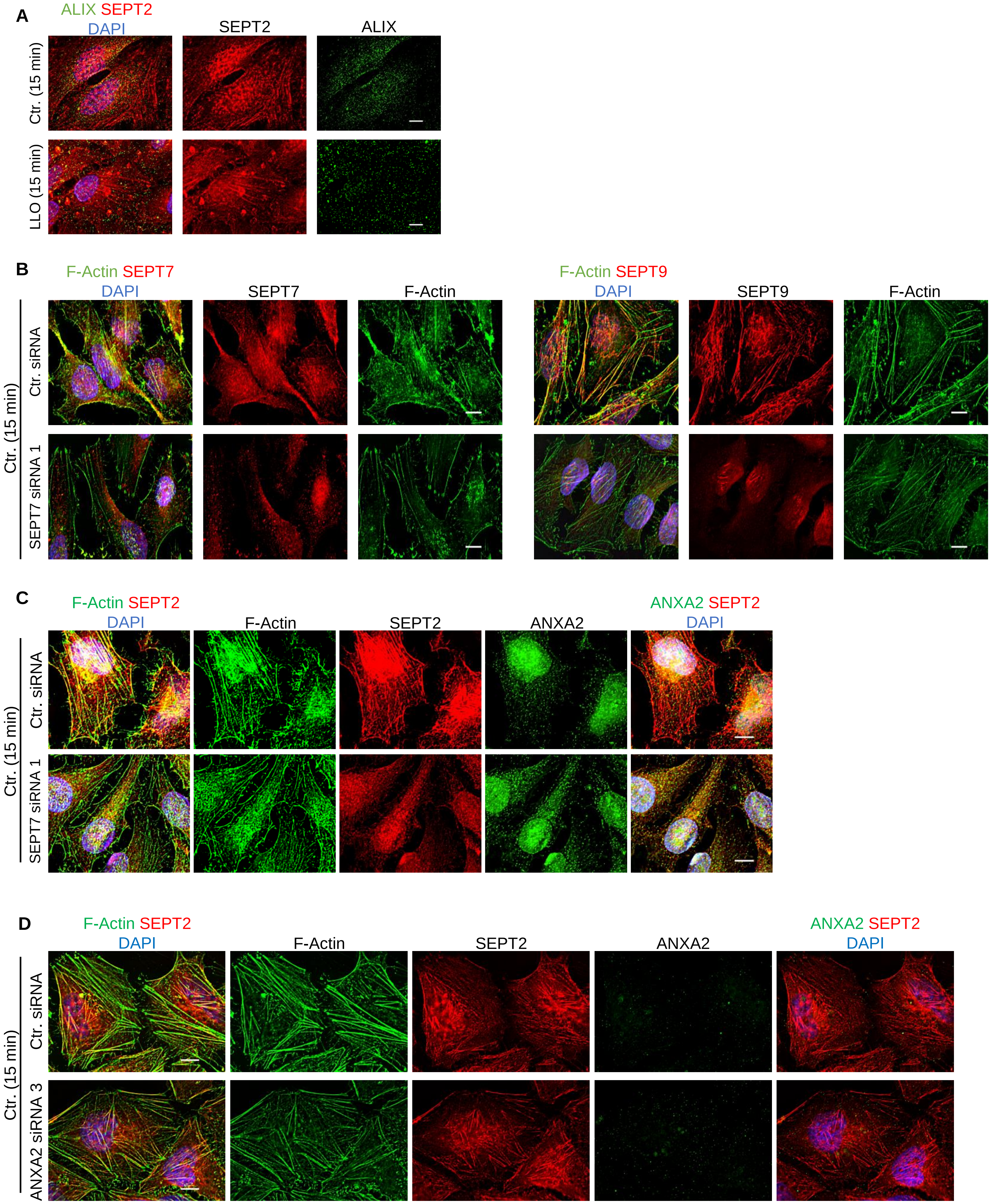

Supplemental Figure 6

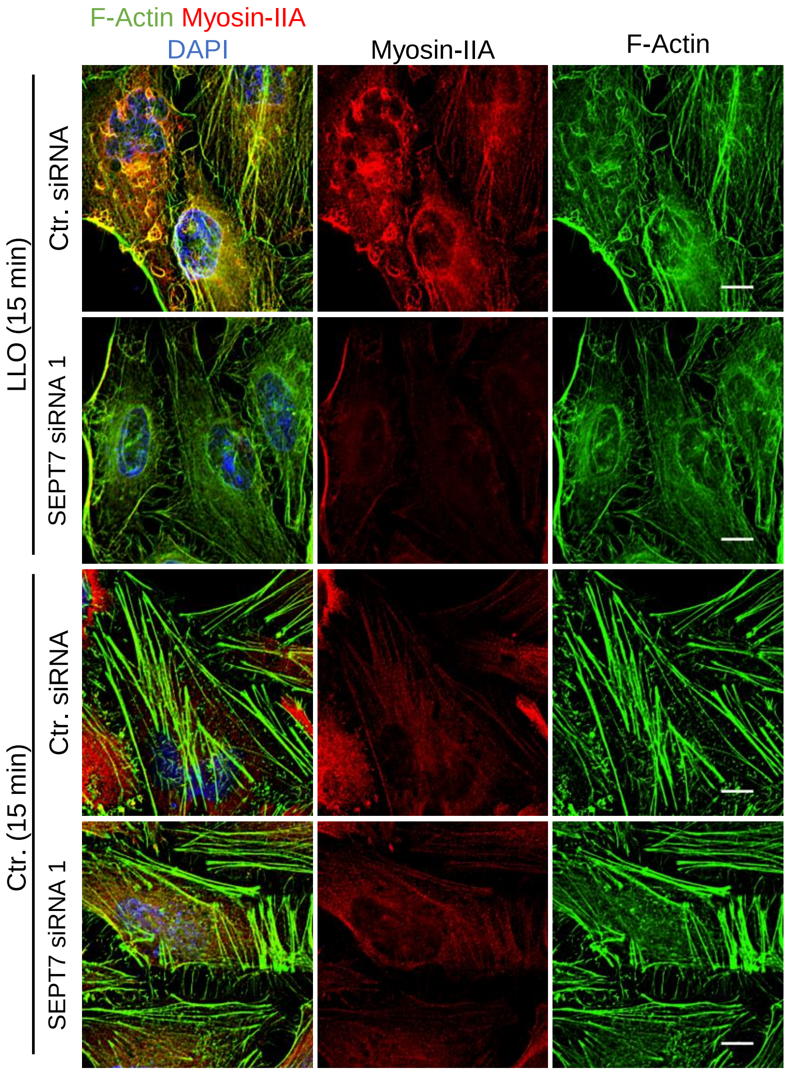
