## Supplemental Table 1 for "The Septin Cytoskeleton is Required for Plasma Membrane Repair"

**Supplemental table 1: List of genes and corresponding silencing sequences used in this work**. Gene identification (Gene ID) provides the link to the National Library of Medicine gene references. siRNA ID are the reference from the ThermoFisher catalogue.

| **Reference Sequence Accession Number** | **Gene Symbol** | **Gene ID** | **siRNA ID** | **Sense siRNA Sequence** | **Antisense siRNA Sequence** |
| --- | --- | --- | --- | --- | --- |
| **Synaptotagmins** |  |  |  |  |  |
| NM_005639 | [SYT1](http://www.ncbi.nlm.nih.gov/entrez/query.fcgi?db=gene&cmd=search&term=SYT1) | [6857](http://www.ncbi.nlm.nih.gov/entrez/query.fcgi?db=gene&cmd=Retrieve&dopt=Graphics&list_uids=6857) | s13694 | GCAAUUUACUUUCAAGGUAtt | UACCUUGAAAGUAAAUUGCtc |
| NM_005639 | [SYT1](http://www.ncbi.nlm.nih.gov/entrez/query.fcgi?db=gene&cmd=search&term=SYT1) | [6857](http://www.ncbi.nlm.nih.gov/entrez/query.fcgi?db=gene&cmd=Retrieve&dopt=Graphics&list_uids=6857) | s13695 | CCCUUAAUCCUGUCUUCAAtt | UUGAAGACAGGAUUAAGGGtt |
| NM_005639 | [SYT1](http://www.ncbi.nlm.nih.gov/entrez/query.fcgi?db=gene&cmd=search&term=SYT1) | [6857](http://www.ncbi.nlm.nih.gov/entrez/query.fcgi?db=gene&cmd=Retrieve&dopt=Graphics&list_uids=6857) | s13696 | CUUUUGAACAAAUCCAGAAtt | UUCUGGAUUUGUUCAAAAGgt |
| NM_177402 | [SYT2](http://www.ncbi.nlm.nih.gov/entrez/query.fcgi?db=gene&cmd=search&term=SYT2) | [127833](http://www.ncbi.nlm.nih.gov/entrez/query.fcgi?db=gene&cmd=Retrieve&dopt=Graphics&list_uids=127833) | s43232 | GAUAAACAAGAUUCCCUUAtt | UAAGGGAAUCUUGUUUAUCtc |
| NM_177402 | [SYT2](http://www.ncbi.nlm.nih.gov/entrez/query.fcgi?db=gene&cmd=search&term=SYT2) | [127833](http://www.ncbi.nlm.nih.gov/entrez/query.fcgi?db=gene&cmd=Retrieve&dopt=Graphics&list_uids=127833) | s43233 | AGACCAAAGUCCAUCGGAAtt | UUCCGAUGGACUUUGGUCUca |
| NM_177402 | [SYT2](http://www.ncbi.nlm.nih.gov/entrez/query.fcgi?db=gene&cmd=search&term=SYT2) | [127833](http://www.ncbi.nlm.nih.gov/entrez/query.fcgi?db=gene&cmd=Retrieve&dopt=Graphics&list_uids=127833) | s43234 | AGAUCCACCUGAUGCAGAAtt | UUCUGCAUCAGGUGGAUCUtc |
| NM_032298 | [SYT3](http://www.ncbi.nlm.nih.gov/entrez/query.fcgi?db=gene&cmd=search&term=SYT3) | [84258](http://www.ncbi.nlm.nih.gov/entrez/query.fcgi?db=gene&cmd=Retrieve&dopt=Graphics&list_uids=84258) | s38744 | CAAAGCCUCUAACCUCAAAtt | UUUGAGGUUAGAGGCUUUGat |
| NM_032298 | [SYT3](http://www.ncbi.nlm.nih.gov/entrez/query.fcgi?db=gene&cmd=search&term=SYT3) | [84258](http://www.ncbi.nlm.nih.gov/entrez/query.fcgi?db=gene&cmd=Retrieve&dopt=Graphics&list_uids=84258) | s224939 | CAGUCACAUCAGCAGGUCAtt | UGACCUGCUGAUGUGACUGgg |
| NM_032298 | [SYT3](http://www.ncbi.nlm.nih.gov/entrez/query.fcgi?db=gene&cmd=search&term=SYT3) | [84258](http://www.ncbi.nlm.nih.gov/entrez/query.fcgi?db=gene&cmd=Retrieve&dopt=Graphics&list_uids=84258) | s38743 | GGACUCCAACGGCUUCUCAtt | UGAGAAGCCGUUGGAGUCCtt |
| NM_020783 | [SYT4](http://www.ncbi.nlm.nih.gov/entrez/query.fcgi?db=gene&cmd=search&term=SYT4) | [6860](http://www.ncbi.nlm.nih.gov/entrez/query.fcgi?db=gene&cmd=Retrieve&dopt=Graphics&list_uids=6860) | s13697 | CGGACUUUCAGAUCCCUAUtt | AUAGGGAUCUGAAAGUCCGga |
| NM_020783 | [SYT4](http://www.ncbi.nlm.nih.gov/entrez/query.fcgi?db=gene&cmd=search&term=SYT4) | [6860](http://www.ncbi.nlm.nih.gov/entrez/query.fcgi?db=gene&cmd=Retrieve&dopt=Graphics&list_uids=6860) | s13698 | CAUUCUAUGGGAUACCCUAtt | UAGGGUAUCCCAUAGAAUGta |
| NM_020783 | [SYT4](http://www.ncbi.nlm.nih.gov/entrez/query.fcgi?db=gene&cmd=search&term=SYT4) | [6860](http://www.ncbi.nlm.nih.gov/entrez/query.fcgi?db=gene&cmd=Retrieve&dopt=Graphics&list_uids=6860) | s13699 | CUCUGACCCAUAUAUCAAAtt | UUUGAUAUAUGGGUCAGAGgt |
| NM_175733 | [SYT9](http://www.ncbi.nlm.nih.gov/entrez/query.fcgi?db=gene&cmd=search&term=SYT9) | [143425](http://www.ncbi.nlm.nih.gov/entrez/query.fcgi?db=gene&cmd=Retrieve&dopt=Graphics&list_uids=143425) | s44533 | GGAUAAUGAUGACGGGAGAtt | UCUCCCGUCAUCAUUAUCCaa |
| NM_175733 | [SYT9](http://www.ncbi.nlm.nih.gov/entrez/query.fcgi?db=gene&cmd=search&term=SYT9) | [143425](http://www.ncbi.nlm.nih.gov/entrez/query.fcgi?db=gene&cmd=Retrieve&dopt=Graphics&list_uids=143425) | s44532 | CCAGAGUUAUAUAAGCAGAtt | UCUGCUUAUAUAACUCUGGtt |
| NM_175733 | [SYT9](http://www.ncbi.nlm.nih.gov/entrez/query.fcgi?db=gene&cmd=search&term=SYT9) | [143425](http://www.ncbi.nlm.nih.gov/entrez/query.fcgi?db=gene&cmd=Retrieve&dopt=Graphics&list_uids=143425) | s44531 | GACUGGAAUUGGUAGAAUUtt | AAUUCUACCAAUUCCAGUCaa |
| NM_003180 | [SYT5](http://www.ncbi.nlm.nih.gov/entrez/query.fcgi?db=gene&cmd=search&term=SYT5) | [6861](http://www.ncbi.nlm.nih.gov/entrez/query.fcgi?db=gene&cmd=Retrieve&dopt=Graphics&list_uids=6861) | s13701 | GCAAAAAGGUGCGGAAGAAtt | UUCUUCCGCACCUUUUUGCcg |
| NM_003180 | [SYT5](http://www.ncbi.nlm.nih.gov/entrez/query.fcgi?db=gene&cmd=search&term=SYT5) | [6861](http://www.ncbi.nlm.nih.gov/entrez/query.fcgi?db=gene&cmd=Retrieve&dopt=Graphics&list_uids=6861) | s13702 | GGACGUAGGAGGACUGUCAtt | UGACAGUCCUCCUACGUCCat |
| NM_003180 | [SYT5](http://www.ncbi.nlm.nih.gov/entrez/query.fcgi?db=gene&cmd=search&term=SYT5) | [6861](http://www.ncbi.nlm.nih.gov/entrez/query.fcgi?db=gene&cmd=Retrieve&dopt=Graphics&list_uids=6861) | s13700 | AAACGGAGGCGGUACGAGAtt | UCUCGUACCGCCUCCGUUUgt |
| NM_205848 | [SYT6](http://www.ncbi.nlm.nih.gov/entrez/query.fcgi?db=gene&cmd=search&term=SYT6) | [148281](http://www.ncbi.nlm.nih.gov/entrez/query.fcgi?db=gene&cmd=Retrieve&dopt=Graphics&list_uids=148281) | s45163 | GGACAUCACAGGCUAUUCAtt | UGAAUAGCCUGUGAUGUCCat |
| NM_205848 | [SYT6](http://www.ncbi.nlm.nih.gov/entrez/query.fcgi?db=gene&cmd=search&term=SYT6) | [148281](http://www.ncbi.nlm.nih.gov/entrez/query.fcgi?db=gene&cmd=Retrieve&dopt=Graphics&list_uids=148281) | s45162 | GAAAACAACCAUAAAGAAAtt | UUUCUUUAUGGUUGUUUUCtt |
| NM_205848 | [SYT6](http://www.ncbi.nlm.nih.gov/entrez/query.fcgi?db=gene&cmd=search&term=SYT6) | [148281](http://www.ncbi.nlm.nih.gov/entrez/query.fcgi?db=gene&cmd=Retrieve&dopt=Graphics&list_uids=148281) | s45164 | AGACCCUGAUUGUGCGUAUtt | AUACGCACAAUCAGGGUCUcg |
| NM_004200 | [SYT7](http://www.ncbi.nlm.nih.gov/entrez/query.fcgi?db=gene&cmd=search&term=SYT7) | [9066](http://www.ncbi.nlm.nih.gov/entrez/query.fcgi?db=gene&cmd=Retrieve&dopt=Graphics&list_uids=9066) | s17291 | CCAUCAUCGUGAACAUCAUtt | AUGAUGUUCACGAUGAUGGag |
| NM_004200 | [SYT7](http://www.ncbi.nlm.nih.gov/entrez/query.fcgi?db=gene&cmd=search&term=SYT7) | [9066](http://www.ncbi.nlm.nih.gov/entrez/query.fcgi?db=gene&cmd=Retrieve&dopt=Graphics&list_uids=9066) | s17293 | AGAAGACGGUGACGAUGAAtt | UUCAUCGUCACCGUCUUCUtc |
| NM_004200 | [SYT7](http://www.ncbi.nlm.nih.gov/entrez/query.fcgi?db=gene&cmd=search&term=SYT7) | [9066](http://www.ncbi.nlm.nih.gov/entrez/query.fcgi?db=gene&cmd=Retrieve&dopt=Graphics&list_uids=9066) | s17292 | UCCAAGUCCUGGACUAUGAtt | UCAUAGUCCAGGACUUGGAgg |
| NM_138567 | [SYT8](http://www.ncbi.nlm.nih.gov/entrez/query.fcgi?db=gene&cmd=search&term=SYT8) | [90019](http://www.ncbi.nlm.nih.gov/entrez/query.fcgi?db=gene&cmd=Retrieve&dopt=Graphics&list_uids=90019) | s40230 | GGUGCAACCUGAUGUGGAUtt | AUCCACAUCAGGUUGCACCag |
| NM_138567 | [SYT8](http://www.ncbi.nlm.nih.gov/entrez/query.fcgi?db=gene&cmd=search&term=SYT8) | [90019](http://www.ncbi.nlm.nih.gov/entrez/query.fcgi?db=gene&cmd=Retrieve&dopt=Graphics&list_uids=90019) | s40231 | AGCUAUACCUGGGCUUAUUtt | AAUAAGCCCAGGUAUAGCUgt |
| NM_138567 | [SYT8](http://www.ncbi.nlm.nih.gov/entrez/query.fcgi?db=gene&cmd=search&term=SYT8) | [90019](http://www.ncbi.nlm.nih.gov/entrez/query.fcgi?db=gene&cmd=Retrieve&dopt=Graphics&list_uids=90019) | s195542 | CAGCUCAUGCUGAACCAGAtt | UCUGGUUCAGCAUGAGCUGga |
| NM_198992 | [SYT10](http://www.ncbi.nlm.nih.gov/entrez/query.fcgi?db=gene&cmd=search&term=SYT10) | [341359](http://www.ncbi.nlm.nih.gov/entrez/query.fcgi?db=gene&cmd=Retrieve&dopt=Graphics&list_uids=341359) | s50836 | GACCAGUUUUGAUAGUCAAtt | UUGACUAUCAAAACUGGUCgc |
| NM_198992 | [SYT10](http://www.ncbi.nlm.nih.gov/entrez/query.fcgi?db=gene&cmd=search&term=SYT10) | [341359](http://www.ncbi.nlm.nih.gov/entrez/query.fcgi?db=gene&cmd=Retrieve&dopt=Graphics&list_uids=341359) | s50837 | GGCCUAUCAUCGAAAACCAtt | UGGUUUUCGAUGAUAGGCCag |
| NM_198992 | [SYT10](http://www.ncbi.nlm.nih.gov/entrez/query.fcgi?db=gene&cmd=search&term=SYT10) | [341359](http://www.ncbi.nlm.nih.gov/entrez/query.fcgi?db=gene&cmd=Retrieve&dopt=Graphics&list_uids=341359) | s50835 | GCAAACCUGUGACUUCCAAtt | UUGGAAGUCACAGGUUUGCtt |
| NM_152280 | [SYT11](http://www.ncbi.nlm.nih.gov/entrez/query.fcgi?db=gene&cmd=search&term=SYT11) | [23208](http://www.ncbi.nlm.nih.gov/entrez/query.fcgi?db=gene&cmd=Retrieve&dopt=Graphics&list_uids=23208) | s23284 | GACUAUAACUUCCCGAAAAtt | UUUUCGGGAAGUUAUAGUCca |
| NM_152280 | [SYT11](http://www.ncbi.nlm.nih.gov/entrez/query.fcgi?db=gene&cmd=search&term=SYT11) | [23208](http://www.ncbi.nlm.nih.gov/entrez/query.fcgi?db=gene&cmd=Retrieve&dopt=Graphics&list_uids=23208) | s23285 | GAACCCACCAUACAAGUUUtt | AAACUUGUAUGGUGGGUUCtt |
| NM_152280 | [SYT11](http://www.ncbi.nlm.nih.gov/entrez/query.fcgi?db=gene&cmd=search&term=SYT11) | [23208](http://www.ncbi.nlm.nih.gov/entrez/query.fcgi?db=gene&cmd=Retrieve&dopt=Graphics&list_uids=23208) | s23283 | GUCUCUCAGGUAAUCCUUAtt | UAAGGAUUACCUGAGAGACcg |
| NM_177963 | [SYT12](http://www.ncbi.nlm.nih.gov/entrez/query.fcgi?db=gene&cmd=search&term=SYT12) | [91683](http://www.ncbi.nlm.nih.gov/entrez/query.fcgi?db=gene&cmd=Retrieve&dopt=Graphics&list_uids=91683) | s40740 | GGAAGAUGAGCAAAAAGAAtt | UUCUUUUUGCUCAUCUUCCtc |
| NM_177963 | [SYT12](http://www.ncbi.nlm.nih.gov/entrez/query.fcgi?db=gene&cmd=search&term=SYT12) | [91683](http://www.ncbi.nlm.nih.gov/entrez/query.fcgi?db=gene&cmd=Retrieve&dopt=Graphics&list_uids=91683) | s40738 | AGCCUGCGGUUUUCUGUAUtt | AUACAGAAAACCGCAGGCUct |
| NM_177963 | [SYT12](http://www.ncbi.nlm.nih.gov/entrez/query.fcgi?db=gene&cmd=search&term=SYT12) | [91683](http://www.ncbi.nlm.nih.gov/entrez/query.fcgi?db=gene&cmd=Retrieve&dopt=Graphics&list_uids=91683) | s40739 | CAAAGGCAGUCUCAGCAUUtt | AAUGCUGAGACUGCCUUUGcg |
| NM_020826 | [SYT13](http://www.ncbi.nlm.nih.gov/entrez/query.fcgi?db=gene&cmd=search&term=SYT13) | [57586](http://www.ncbi.nlm.nih.gov/entrez/query.fcgi?db=gene&cmd=Retrieve&dopt=Graphics&list_uids=57586) | s33380 | CAACUAUGCAGACUAUUCAtt | UGAAUAGUCUGCAUAGUUGat |
| NM_020826 | [SYT13](http://www.ncbi.nlm.nih.gov/entrez/query.fcgi?db=gene&cmd=search&term=SYT13) | [57586](http://www.ncbi.nlm.nih.gov/entrez/query.fcgi?db=gene&cmd=Retrieve&dopt=Graphics&list_uids=57586) | s33381 | GGAACGAGAUGAUCAUGUUtt | AACAUGAUCAUCUCGUUCCac |
| NM_020826 | [SYT13](http://www.ncbi.nlm.nih.gov/entrez/query.fcgi?db=gene&cmd=search&term=SYT13) | [57586](http://www.ncbi.nlm.nih.gov/entrez/query.fcgi?db=gene&cmd=Retrieve&dopt=Graphics&list_uids=57586) | s33379 | GAAGCAGACUAAACGAGCUtt | AGCUCGUUUAGUCUGCUUCtt |
| NM_153262 | [SYT14](http://www.ncbi.nlm.nih.gov/entrez/query.fcgi?db=gene&cmd=search&term=SYT14) | [255928](http://www.ncbi.nlm.nih.gov/entrez/query.fcgi?db=gene&cmd=Retrieve&dopt=Graphics&list_uids=255928) | s48718 | CCUUGUUCUUCUACCUAUAtt | UAUAGGUAGAAGAACAAGGtg |
| NM_153262 | [SYT14](http://www.ncbi.nlm.nih.gov/entrez/query.fcgi?db=gene&cmd=search&term=SYT14) | [255928](http://www.ncbi.nlm.nih.gov/entrez/query.fcgi?db=gene&cmd=Retrieve&dopt=Graphics&list_uids=255928) | s48720 | CCUGUGAUAUUGGAACCUUtt | AAGGUUCCAAUAUCACAGGca |
| NM_153262 | [SYT14](http://www.ncbi.nlm.nih.gov/entrez/query.fcgi?db=gene&cmd=search&term=SYT14) | [255928](http://www.ncbi.nlm.nih.gov/entrez/query.fcgi?db=gene&cmd=Retrieve&dopt=Graphics&list_uids=255928) | s48719 | CCAAGAUGCUCAUAUAACAtt | UGUUAUAUGAGCAUCUUGGac |
| NM_181519 | [SYT15](http://www.ncbi.nlm.nih.gov/entrez/query.fcgi?db=gene&cmd=search&term=SYT15) | [83849](http://www.ncbi.nlm.nih.gov/entrez/query.fcgi?db=gene&cmd=Retrieve&dopt=Graphics&list_uids=83849) | s38206 | CCUCCAAUCCAAGACCAAAtt | UUUGGUCUUGGAUUGGAGGaa |
| NM_181519 | [SYT15](http://www.ncbi.nlm.nih.gov/entrez/query.fcgi?db=gene&cmd=search&term=SYT15) | [83849](http://www.ncbi.nlm.nih.gov/entrez/query.fcgi?db=gene&cmd=Retrieve&dopt=Graphics&list_uids=83849) | s38208 | GUGUCUCUGAUGAACCACAtt | UGUGGUUCAUCAGAGACACtt |
| NM_181519 | [SYT15](http://www.ncbi.nlm.nih.gov/entrez/query.fcgi?db=gene&cmd=search&term=SYT15) | [83849](http://www.ncbi.nlm.nih.gov/entrez/query.fcgi?db=gene&cmd=Retrieve&dopt=Graphics&list_uids=83849) | s38207 | AGCUGUACAAGUUCCCGGAtt | UCCGGGAACUUGUACAGCUct |
| NM_031914 | [SYT16](http://www.ncbi.nlm.nih.gov/entrez/query.fcgi?db=gene&cmd=search&term=SYT16) | [83851](http://www.ncbi.nlm.nih.gov/entrez/query.fcgi?db=gene&cmd=Retrieve&dopt=Graphics&list_uids=83851) | s38212 | CGUUGAUGAUUUCCGUUUAtt | UAAACGGAAAUCAUCAACGtg |
| NM_031914 | [SYT16](http://www.ncbi.nlm.nih.gov/entrez/query.fcgi?db=gene&cmd=search&term=SYT16) | [83851](http://www.ncbi.nlm.nih.gov/entrez/query.fcgi?db=gene&cmd=Retrieve&dopt=Graphics&list_uids=83851) | s38214 | GCUUCGCUGGUUAACAUAAtt | UUAUGUUAACCAGCGAAGCag |
| NM_031914 | [SYT16](http://www.ncbi.nlm.nih.gov/entrez/query.fcgi?db=gene&cmd=search&term=SYT16) | [83851](http://www.ncbi.nlm.nih.gov/entrez/query.fcgi?db=gene&cmd=Retrieve&dopt=Graphics&list_uids=83851) | s38213 | GGAUAUCUCGGGUUUAUGAtt | UCAUAAACCCGAGAUAUCCag |
| NM_016524 | [SYT17](http://www.ncbi.nlm.nih.gov/entrez/query.fcgi?db=gene&cmd=search&term=SYT17) | [51760](http://www.ncbi.nlm.nih.gov/entrez/query.fcgi?db=gene&cmd=Retrieve&dopt=Graphics&list_uids=51760) | s28629 | GCUCUCCACUCAUCGAUAUtt | AUAUCGAUGAGUGGAGAGCtg |
| NM_016524 | [SYT17](http://www.ncbi.nlm.nih.gov/entrez/query.fcgi?db=gene&cmd=search&term=SYT17) | [51760](http://www.ncbi.nlm.nih.gov/entrez/query.fcgi?db=gene&cmd=Retrieve&dopt=Graphics&list_uids=51760) | s28627 | GCCUAGUGUUUACAGUUUUtt | AAAACUGUAAACACUAGGCtg |
| NM_016524 | [SYT17](http://www.ncbi.nlm.nih.gov/entrez/query.fcgi?db=gene&cmd=search&term=SYT17) | [51760](http://www.ncbi.nlm.nih.gov/entrez/query.fcgi?db=gene&cmd=Retrieve&dopt=Graphics&list_uids=51760) | s28628 | GCUGGUGCAUGGACUCAAAtt | UUUGAGUCCAUGCACCAGCtg |
| **SNARE** |  |  |  |  |  |
| NM_003825 | [SNAP23](http://www.ncbi.nlm.nih.gov/entrez/query.fcgi?db=gene&cmd=search&term=SNAP23) | [8773](http://www.ncbi.nlm.nih.gov/entrez/query.fcgi?db=gene&cmd=Retrieve&dopt=Graphics&list_uids=8773) | s16709 | GGAACAACUAAACCGCAUAtt | UAUGCGGUUUAGUUGUUCCtt |
| NM_003825 | [SNAP23](http://www.ncbi.nlm.nih.gov/entrez/query.fcgi?db=gene&cmd=search&term=SNAP23) | [8773](http://www.ncbi.nlm.nih.gov/entrez/query.fcgi?db=gene&cmd=Retrieve&dopt=Graphics&list_uids=8773) | s16708 | CAGAGAUCGUAUUGAUAUUtt | AAUAUCAAUACGAUCUCUGtt |
| NM_003825 | [SNAP23](http://www.ncbi.nlm.nih.gov/entrez/query.fcgi?db=gene&cmd=search&term=SNAP23) | [8773](http://www.ncbi.nlm.nih.gov/entrez/query.fcgi?db=gene&cmd=Retrieve&dopt=Graphics&list_uids=8773) | s16710 | UCACUAUGCUGGAUGAACAtt | UGUUCAUCCAGCAUAGUGAtg |
| NM_003081 | [SNAP25](http://www.ncbi.nlm.nih.gov/entrez/query.fcgi?db=gene&cmd=search&term=SNAP25) | [6616](http://www.ncbi.nlm.nih.gov/entrez/query.fcgi?db=gene&cmd=Retrieve&dopt=Graphics&list_uids=6616) | s13189 | GGACUUUGGUUAUGUUGGAtt | UCCAACAUAACCAAAGUCCtg |
| NM_003081 | [SNAP25](http://www.ncbi.nlm.nih.gov/entrez/query.fcgi?db=gene&cmd=search&term=SNAP25) | [6616](http://www.ncbi.nlm.nih.gov/entrez/query.fcgi?db=gene&cmd=Retrieve&dopt=Graphics&list_uids=6616) | s13188 | UGAAAUGGAUGAAAACCUAtt | UAGGUUUUCAUCCAUUUCAtt |
| NM_003081 | [SNAP25](http://www.ncbi.nlm.nih.gov/entrez/query.fcgi?db=gene&cmd=search&term=SNAP25) | [6616](http://www.ncbi.nlm.nih.gov/entrez/query.fcgi?db=gene&cmd=Retrieve&dopt=Graphics&list_uids=6616) | s13190 | GGUUGAAGAGAGUAAAGAUtt | AUCUUUACUCUCUUCAACCag |
| NM_004782 | [SNAP29](http://www.ncbi.nlm.nih.gov/entrez/query.fcgi?db=gene&cmd=search&term=SNAP29) | [9342](http://www.ncbi.nlm.nih.gov/entrez/query.fcgi?db=gene&cmd=Retrieve&dopt=Graphics&list_uids=9342) | s17859 | CAACCAAAGUGGACAAGUUtt | AACUUGUCCACUUUGGUUGtc |
| NM_004782 | [SNAP29](http://www.ncbi.nlm.nih.gov/entrez/query.fcgi?db=gene&cmd=search&term=SNAP29) | [9342](http://www.ncbi.nlm.nih.gov/entrez/query.fcgi?db=gene&cmd=Retrieve&dopt=Graphics&list_uids=9342) | s17860 | GUACUGAUGCUUACCCAAAtt | UUUGGGUAAGCAUCAGUACtc |
| NM_004782 | [SNAP29](http://www.ncbi.nlm.nih.gov/entrez/query.fcgi?db=gene&cmd=search&term=SNAP29) | [9342](http://www.ncbi.nlm.nih.gov/entrez/query.fcgi?db=gene&cmd=Retrieve&dopt=Graphics&list_uids=9342) | s17861 | AAGCUAUAAGUACAAGUAAtt | UUACUUGUACUUAUAGCUUct |
| NM_199245 | [VAMP1](http://www.ncbi.nlm.nih.gov/entrez/query.fcgi?db=gene&cmd=search&term=VAMP1) | [6843](http://www.ncbi.nlm.nih.gov/entrez/query.fcgi?db=gene&cmd=Retrieve&dopt=Graphics&list_uids=6843) | s13665 | AGGAAGUAUUGGUGGAAAAtt | UUUUCCACCAAUACUUCCUct |
| NM_199245 | [VAMP1](http://www.ncbi.nlm.nih.gov/entrez/query.fcgi?db=gene&cmd=search&term=VAMP1) | [6843](http://www.ncbi.nlm.nih.gov/entrez/query.fcgi?db=gene&cmd=Retrieve&dopt=Graphics&list_uids=6843) | s13664 | CUCCUAACAUGACCAGUAAtt | UUACUGGUCAUGUUAGGAGga |
| NM_199245 | [VAMP1](http://www.ncbi.nlm.nih.gov/entrez/query.fcgi?db=gene&cmd=search&term=VAMP1) | [6843](http://www.ncbi.nlm.nih.gov/entrez/query.fcgi?db=gene&cmd=Retrieve&dopt=Graphics&list_uids=6843) | s13666 | GCAUCACAAUUUGAGAGCAtt | UGCUCUCAAAUUGUGAUGCtc |
| NM_014232 | [VAMP2](http://www.ncbi.nlm.nih.gov/entrez/query.fcgi?db=gene&cmd=search&term=VAMP2) | [6844](http://www.ncbi.nlm.nih.gov/entrez/query.fcgi?db=gene&cmd=Retrieve&dopt=Graphics&list_uids=6844) | s13667 | UCAUGAGGGUGAACGUGGAtt | UCCACGUUCACCCUCAUGAtg |
| NM_014232 | [VAMP2](http://www.ncbi.nlm.nih.gov/entrez/query.fcgi?db=gene&cmd=search&term=VAMP2) | [6844](http://www.ncbi.nlm.nih.gov/entrez/query.fcgi?db=gene&cmd=Retrieve&dopt=Graphics&list_uids=6844) | s13668 | CCAGUAACAGGAGACUGCAtt | UGCAGUCUCCUGUUACUGGtg |
| NM_014232 | [VAMP2](http://www.ncbi.nlm.nih.gov/entrez/query.fcgi?db=gene&cmd=search&term=VAMP2) | [6844](http://www.ncbi.nlm.nih.gov/entrez/query.fcgi?db=gene&cmd=Retrieve&dopt=Graphics&list_uids=6844) | s13669 | GGAAAAACCUCAAGAUGAUtt | AUCAUCUUGAGGUUUUUCCac |
| NM_004781 | [VAMP3](http://www.ncbi.nlm.nih.gov/entrez/query.fcgi?db=gene&cmd=search&term=VAMP3) | [9341](http://www.ncbi.nlm.nih.gov/entrez/query.fcgi?db=gene&cmd=Retrieve&dopt=Graphics&list_uids=9341) | s17857 | AAAUAUUGGUGGAAGAAUUtt | AAUUCUUCCACCAAUAUUUcc |
| NM_004781 | [VAMP3](http://www.ncbi.nlm.nih.gov/entrez/query.fcgi?db=gene&cmd=search&term=VAMP3) | [9341](http://www.ncbi.nlm.nih.gov/entrez/query.fcgi?db=gene&cmd=Retrieve&dopt=Graphics&list_uids=9341) | s17856 | CGGGAUUACUGUUCUGGUUtt | AACCAGAACAGUAAUCCCGat |
| NM_004781 | [VAMP3](http://www.ncbi.nlm.nih.gov/entrez/query.fcgi?db=gene&cmd=search&term=VAMP3) | [9341](http://www.ncbi.nlm.nih.gov/entrez/query.fcgi?db=gene&cmd=Retrieve&dopt=Graphics&list_uids=9341) | s228492 | GCUUCUCAAUUUGAAACGAtt | UCGUUUCAAAUUGAGAAGCgc |
| NM_003762 | [VAMP4](http://www.ncbi.nlm.nih.gov/entrez/query.fcgi?db=gene&cmd=search&term=VAMP4) | [8674](http://www.ncbi.nlm.nih.gov/entrez/query.fcgi?db=gene&cmd=Retrieve&dopt=Graphics&list_uids=8674) | s16525 | CAAACAACUUCGAAGGCAAtt | UUGCCUUCGAAGUUGUUUGga |
| NM_003762 | [VAMP4](http://www.ncbi.nlm.nih.gov/entrez/query.fcgi?db=gene&cmd=search&term=VAMP4) | [8674](http://www.ncbi.nlm.nih.gov/entrez/query.fcgi?db=gene&cmd=Retrieve&dopt=Graphics&list_uids=8674) | s16526 | GAUUUGGACCUAGAAAUGAtt | UCAUUUCUAGGUCCAAAUCtt |
| NM_003762 | [VAMP4](http://www.ncbi.nlm.nih.gov/entrez/query.fcgi?db=gene&cmd=search&term=VAMP4) | [8674](http://www.ncbi.nlm.nih.gov/entrez/query.fcgi?db=gene&cmd=Retrieve&dopt=Graphics&list_uids=8674) | s16527 | AGCUUAUCGGAUAAUGCAAtt | UUGCAUUAUCCGAUAAGCUtt |
| NM_005638 | [VAMP7](http://www.ncbi.nlm.nih.gov/entrez/query.fcgi?db=gene&cmd=search&term=VAMP7) | [6845](http://www.ncbi.nlm.nih.gov/entrez/query.fcgi?db=gene&cmd=Retrieve&dopt=Graphics&list_uids=6845) | s13672 | GAUUGGAAUUAUUGAUUGAtt | UCAAUCAAUAAUUCCAAUCtt |
| NM_005638 | [VAMP7](http://www.ncbi.nlm.nih.gov/entrez/query.fcgi?db=gene&cmd=search&term=VAMP7) | [6845](http://www.ncbi.nlm.nih.gov/entrez/query.fcgi?db=gene&cmd=Retrieve&dopt=Graphics&list_uids=6845) | s13671 | GACUACUUACGGUUCAAGAtt | UCUUGAACCGUAAGUAGUCtg |
| NM_005638 | [VAMP7](http://www.ncbi.nlm.nih.gov/entrez/query.fcgi?db=gene&cmd=search&term=VAMP7) | [6845](http://www.ncbi.nlm.nih.gov/entrez/query.fcgi?db=gene&cmd=Retrieve&dopt=Graphics&list_uids=6845) | s13670 | UCAUCAUCGUAUCAAUUGUtt | ACAAUUGAUACGAUGAUGAtg |
| NM_003761 | [VAMP8](http://www.ncbi.nlm.nih.gov/entrez/query.fcgi?db=gene&cmd=search&term=VAMP8) | [8673](http://www.ncbi.nlm.nih.gov/entrez/query.fcgi?db=gene&cmd=Retrieve&dopt=Graphics&list_uids=8673) | s16523 | UGAAGAUGAUUGUCCUUAUtt | AUAAGGACAAUCAUCUUCAcg |
| NM_003761 | [VAMP8](http://www.ncbi.nlm.nih.gov/entrez/query.fcgi?db=gene&cmd=search&term=VAMP8) | [8673](http://www.ncbi.nlm.nih.gov/entrez/query.fcgi?db=gene&cmd=Retrieve&dopt=Graphics&list_uids=8673) | s16524 | GAACAUCUCCGCAACAAGAtt | UCUUGUUGCGGAGAUGUUCca |
| NM_003761 | [VAMP8](http://www.ncbi.nlm.nih.gov/entrez/query.fcgi?db=gene&cmd=search&term=VAMP8) | [8673](http://www.ncbi.nlm.nih.gov/entrez/query.fcgi?db=gene&cmd=Retrieve&dopt=Graphics&list_uids=8673) | s16522 | GGAGUUAAGAAUAUUAUGAtt | UCAUAAUAUUCUUAACUCCct |
| NM_004738 | [VAPB](http://www.ncbi.nlm.nih.gov/entrez/query.fcgi?db=gene&cmd=search&term=VAPB) | [9217](http://www.ncbi.nlm.nih.gov/entrez/query.fcgi?db=gene&cmd=Retrieve&dopt=Graphics&list_uids=9217) | s17623 | GCAAAACCGGAAGACCUUAtt | UAAGGUCUUCCGGUUUUGCct |
| NM_004738 | [VAPB](http://www.ncbi.nlm.nih.gov/entrez/query.fcgi?db=gene&cmd=search&term=VAPB) | [9217](http://www.ncbi.nlm.nih.gov/entrez/query.fcgi?db=gene&cmd=Retrieve&dopt=Graphics&list_uids=9217) | s17625 | AAACACCAAUAGUGUCUAAtt | UUAGACACUAUUGGUGUUUct |
| NM_004738 | [VAPB](http://www.ncbi.nlm.nih.gov/entrez/query.fcgi?db=gene&cmd=search&term=VAPB) | [9217](http://www.ncbi.nlm.nih.gov/entrez/query.fcgi?db=gene&cmd=Retrieve&dopt=Graphics&list_uids=9217) | s17624 | UAAUGUAUCUGUGAUGUUAtt | UAACAUCACAGAUACAUUAat |
| NM_004738 | [VAPB](http://www.ncbi.nlm.nih.gov/entrez/query.fcgi?db=gene&cmd=search&term=VAPB) | [9217](http://www.ncbi.nlm.nih.gov/entrez/query.fcgi?db=gene&cmd=Retrieve&dopt=Graphics&list_uids=9217) | s17623 | GCAAAACCGGAAGACCUUAtt | UAAGGUCUUCCGGUUUUGCct |
| NM_004738 | [VAPB](http://www.ncbi.nlm.nih.gov/entrez/query.fcgi?db=gene&cmd=search&term=VAPB) | [9217](http://www.ncbi.nlm.nih.gov/entrez/query.fcgi?db=gene&cmd=Retrieve&dopt=Graphics&list_uids=9217) | s17625 | AAACACCAAUAGUGUCUAAtt | UUAGACACUAUUGGUGUUUct |
| NM_004738 | [VAPB](http://www.ncbi.nlm.nih.gov/entrez/query.fcgi?db=gene&cmd=search&term=VAPB) | [9217](http://www.ncbi.nlm.nih.gov/entrez/query.fcgi?db=gene&cmd=Retrieve&dopt=Graphics&list_uids=9217) | s17624 | UAAUGUAUCUGUGAUGUUAtt | UAACAUCACAGAUACAUUAat |
| NM_003765 | [STX10](http://www.ncbi.nlm.nih.gov/entrez/query.fcgi?db=gene&cmd=search&term=STX10) | [8677](http://www.ncbi.nlm.nih.gov/entrez/query.fcgi?db=gene&cmd=Retrieve&dopt=Graphics&list_uids=8677) | s16536 | AGACCAUCGGUAUAGUGGAtt | UCCACUAUACCGAUGGUCUct |
| NM_003765 | [STX10](http://www.ncbi.nlm.nih.gov/entrez/query.fcgi?db=gene&cmd=search&term=STX10) | [8677](http://www.ncbi.nlm.nih.gov/entrez/query.fcgi?db=gene&cmd=Retrieve&dopt=Graphics&list_uids=8677) | s16535 | CAACAGCCGUAGCAUUUUUtt | AAAAAUGCUACGGCUGUUGgg |
| NM_003765 | [STX10](http://www.ncbi.nlm.nih.gov/entrez/query.fcgi?db=gene&cmd=search&term=STX10) | [8677](http://www.ncbi.nlm.nih.gov/entrez/query.fcgi?db=gene&cmd=Retrieve&dopt=Graphics&list_uids=8677) | s225020 | GAGAGAUACUCGCAGGCAAtt | UUGCCUGCGAGUAUCUCUCtg |
| NM_003764 | [STX11](http://www.ncbi.nlm.nih.gov/entrez/query.fcgi?db=gene&cmd=search&term=STX11) | [8676](http://www.ncbi.nlm.nih.gov/entrez/query.fcgi?db=gene&cmd=Retrieve&dopt=Graphics&list_uids=8676) | s16532 | GAGCUCAACGUACAAAAGAtt | UCUUUUGUACGUUGAGCUCga |
| NM_003764 | [STX11](http://www.ncbi.nlm.nih.gov/entrez/query.fcgi?db=gene&cmd=search&term=STX11) | [8676](http://www.ncbi.nlm.nih.gov/entrez/query.fcgi?db=gene&cmd=Retrieve&dopt=Graphics&list_uids=8676) | s16533 | AGUGGGACGUGUUUUCCGAtt | UCGGAAAACACGUCCCACUta |
| NM_003764 | [STX11](http://www.ncbi.nlm.nih.gov/entrez/query.fcgi?db=gene&cmd=search&term=STX11) | [8676](http://www.ncbi.nlm.nih.gov/entrez/query.fcgi?db=gene&cmd=Retrieve&dopt=Graphics&list_uids=8676) | s16531 | AAAUCACACCAAACACUUAtt | UAAGUGUUUGGUGUGAUUUta |
| NM_177424 | [STX12](http://www.ncbi.nlm.nih.gov/entrez/query.fcgi?db=gene&cmd=search&term=STX12) | [23673](http://www.ncbi.nlm.nih.gov/entrez/query.fcgi?db=gene&cmd=Retrieve&dopt=Graphics&list_uids=23673) | s24309 | GCAGGACUCAAGCAAGCUAtt | UAGCUUGCUUGAGUCCUGCtt |
| NM_177424 | [STX12](http://www.ncbi.nlm.nih.gov/entrez/query.fcgi?db=gene&cmd=search&term=STX12) | [23673](http://www.ncbi.nlm.nih.gov/entrez/query.fcgi?db=gene&cmd=Retrieve&dopt=Graphics&list_uids=23673) | s24307 | GGAAAAUCUGCAACAGUUAtt | UAACUGUUGCAGAUUUUCCtg |
| NM_177424 | [STX12](http://www.ncbi.nlm.nih.gov/entrez/query.fcgi?db=gene&cmd=search&term=STX12) | [23673](http://www.ncbi.nlm.nih.gov/entrez/query.fcgi?db=gene&cmd=Retrieve&dopt=Graphics&list_uids=23673) | s24308 | GGAUUUGGAACUUAUUAAAtt | UUUAAUAAGUUCCAAAUCCtg |
| NM_001001433 | [STX16](http://www.ncbi.nlm.nih.gov/entrez/query.fcgi?db=gene&cmd=search&term=STX16) | [8675](http://www.ncbi.nlm.nih.gov/entrez/query.fcgi?db=gene&cmd=Retrieve&dopt=Graphics&list_uids=8675) | s16530 | CAGCGAUUGGUGUGACAAAtt | UUUGUCACACCAAUCGCUGct |
| NM_001001433 | [STX16](http://www.ncbi.nlm.nih.gov/entrez/query.fcgi?db=gene&cmd=search&term=STX16) | [8675](http://www.ncbi.nlm.nih.gov/entrez/query.fcgi?db=gene&cmd=Retrieve&dopt=Graphics&list_uids=8675) | s16528 | GCAUCAGCUUAGAUCCAGAtt | UCUGGAUCUAAGCUGAUGCct |
| NM_001001433 | [STX16](http://www.ncbi.nlm.nih.gov/entrez/query.fcgi?db=gene&cmd=search&term=STX16) | [8675](http://www.ncbi.nlm.nih.gov/entrez/query.fcgi?db=gene&cmd=Retrieve&dopt=Graphics&list_uids=8675) | s16529 | CGGAAUAAUUCCAUCCAAAtt | UUUGGAUGGAAUUAUUCCGca |
| NM_017919 | [STX17](http://www.ncbi.nlm.nih.gov/entrez/query.fcgi?db=gene&cmd=search&term=STX17) | [55014](http://www.ncbi.nlm.nih.gov/entrez/query.fcgi?db=gene&cmd=Retrieve&dopt=Graphics&list_uids=55014) | s224334 | CCAAUAUCCGAGAAAUUGAtt | UCAAUUUCUCGGAUAUUGGat |
| NM_017919 | [STX17](http://www.ncbi.nlm.nih.gov/entrez/query.fcgi?db=gene&cmd=search&term=STX17) | [55014](http://www.ncbi.nlm.nih.gov/entrez/query.fcgi?db=gene&cmd=Retrieve&dopt=Graphics&list_uids=55014) | s29994 | GAGUUUGACUCAGAUAUAUtt | AUAUAUCUGAGUCAAACUCtg |
| NM_017919 | [STX17](http://www.ncbi.nlm.nih.gov/entrez/query.fcgi?db=gene&cmd=search&term=STX17) | [55014](http://www.ncbi.nlm.nih.gov/entrez/query.fcgi?db=gene&cmd=Retrieve&dopt=Graphics&list_uids=55014) | s224335 | GGAGAAGAUUGACAGCAUUtt | AAUGCUGUCAAUCUUCUCCtg |
| NM_016930 | [STX18](http://www.ncbi.nlm.nih.gov/entrez/query.fcgi?db=gene&cmd=search&term=STX18) | [53407](http://www.ncbi.nlm.nih.gov/entrez/query.fcgi?db=gene&cmd=Retrieve&dopt=Graphics&list_uids=53407) | s28733 | GCAAUUCAGCAACUACGAAtt | UUCGUAGUUGCUGAAUUGCtt |
| NM_016930 | [STX18](http://www.ncbi.nlm.nih.gov/entrez/query.fcgi?db=gene&cmd=search&term=STX18) | [53407](http://www.ncbi.nlm.nih.gov/entrez/query.fcgi?db=gene&cmd=Retrieve&dopt=Graphics&list_uids=53407) | s28734 | GCAACGAAGACAUAAGAGAtt | UCUCUUAUGUCUUCGUUGCct |
| NM_016930 | [STX18](http://www.ncbi.nlm.nih.gov/entrez/query.fcgi?db=gene&cmd=search&term=STX18) | [53407](http://www.ncbi.nlm.nih.gov/entrez/query.fcgi?db=gene&cmd=Retrieve&dopt=Graphics&list_uids=53407) | s28735 | GGACCGCUGUUUUGGAUUUtt | AAAUCCAAAACAGCGGUCCtg |
| NM_001001850 | [STX19](http://www.ncbi.nlm.nih.gov/entrez/query.fcgi?db=gene&cmd=search&term=STX19) | [415117](http://www.ncbi.nlm.nih.gov/entrez/query.fcgi?db=gene&cmd=Retrieve&dopt=Graphics&list_uids=415117) | s53949 | GAAAUGACAGUGAAUAGUAtt | UACUAUUCACUGUCAUUUCaa |
| NM_001001850 | [STX19](http://www.ncbi.nlm.nih.gov/entrez/query.fcgi?db=gene&cmd=search&term=STX19) | [415117](http://www.ncbi.nlm.nih.gov/entrez/query.fcgi?db=gene&cmd=Retrieve&dopt=Graphics&list_uids=415117) | s53950 | CAAGGAUACUUAAAUCUCAtt | UGAGAUUUAAGUAUCCUUGtg |
| NM_001001850 | [STX19](http://www.ncbi.nlm.nih.gov/entrez/query.fcgi?db=gene&cmd=search&term=STX19) | [415117](http://www.ncbi.nlm.nih.gov/entrez/query.fcgi?db=gene&cmd=Retrieve&dopt=Graphics&list_uids=415117) | s53951 | GCUUACUUACAGAAAUCAAtt | UUGAUUUCUGUAAGUAAGCtt |
| NM_004603 | [STX1A](http://www.ncbi.nlm.nih.gov/entrez/query.fcgi?db=gene&cmd=search&term=STX1A) | [6804](http://www.ncbi.nlm.nih.gov/entrez/query.fcgi?db=gene&cmd=Retrieve&dopt=Graphics&list_uids=6804) | s13590 | GAACUCAUGUCCGACAUAAtt | UUAUGUCGGACAUGAGUUCtt |
| NM_004603 | [STX1A](http://www.ncbi.nlm.nih.gov/entrez/query.fcgi?db=gene&cmd=search&term=STX1A) | [6804](http://www.ncbi.nlm.nih.gov/entrez/query.fcgi?db=gene&cmd=Retrieve&dopt=Graphics&list_uids=6804) | s13591 | AGAUCGCAGAGAACGUGGAtt | UCCACGUUCUCUGCGAUCUtg |
| NM_004603 | [STX1A](http://www.ncbi.nlm.nih.gov/entrez/query.fcgi?db=gene&cmd=search&term=STX1A) | [6804](http://www.ncbi.nlm.nih.gov/entrez/query.fcgi?db=gene&cmd=Retrieve&dopt=Graphics&list_uids=6804) | s13589 | CAAACAAAGUUCGUUCCAAtt | UUGGAACGAACUUUGUUUGct |
| NM_052874 | [STX1B](http://www.ncbi.nlm.nih.gov/entrez/query.fcgi?db=gene&cmd=search&term=STX1B) | [112755](http://www.ncbi.nlm.nih.gov/entrez/query.fcgi?db=gene&cmd=Retrieve&dopt=Graphics&list_uids=112755) | s223220 | CAGAUGACAUCAAAAUGGAtt | UCCAUUUUGAUGUCAUCUGtg |
| NM_052874 | [STX1B](http://www.ncbi.nlm.nih.gov/entrez/query.fcgi?db=gene&cmd=search&term=STX1B) | [112755](http://www.ncbi.nlm.nih.gov/entrez/query.fcgi?db=gene&cmd=Retrieve&dopt=Graphics&list_uids=112755) | s41369 | CCACCAACGAAGAACUGGAtt | UCCAGUUCUUCGUUGGUGGtg |
| NM_052874 | [STX1B](http://www.ncbi.nlm.nih.gov/entrez/query.fcgi?db=gene&cmd=search&term=STX1B) | [112755](http://www.ncbi.nlm.nih.gov/entrez/query.fcgi?db=gene&cmd=Retrieve&dopt=Graphics&list_uids=112755) | s41368 | GGUUCGGUCCAAAUUGAAAtt | UUUCAAUUUGGACCGAACCtt |
| NM_001980 | [STX2](http://www.ncbi.nlm.nih.gov/entrez/query.fcgi?db=gene&cmd=search&term=STX2) | [2054](http://www.ncbi.nlm.nih.gov/entrez/query.fcgi?db=gene&cmd=Retrieve&dopt=Graphics&list_uids=2054) | s4756 | CAGUUGUUGUGGUUGAGAAtt | UUCUCAACCACAACAACUGtg |
| NM_001980 | [STX2](http://www.ncbi.nlm.nih.gov/entrez/query.fcgi?db=gene&cmd=search&term=STX2) | [2054](http://www.ncbi.nlm.nih.gov/entrez/query.fcgi?db=gene&cmd=Retrieve&dopt=Graphics&list_uids=2054) | s4755 | CAAGAAAACUGCGAAUAAAtt | UUUAUUCGCAGUUUUCUUGat |
| NM_001980 | [STX2](http://www.ncbi.nlm.nih.gov/entrez/query.fcgi?db=gene&cmd=search&term=STX2) | [2054](http://www.ncbi.nlm.nih.gov/entrez/query.fcgi?db=gene&cmd=Retrieve&dopt=Graphics&list_uids=2054) | s4757 | CUAUGUUUGUGGAGACUCAtt | UGAGUCUCCACAAACAUAGcc |
| NM_004177 | [STX3](http://www.ncbi.nlm.nih.gov/entrez/query.fcgi?db=gene&cmd=search&term=STX3) | [6809](http://www.ncbi.nlm.nih.gov/entrez/query.fcgi?db=gene&cmd=Retrieve&dopt=Graphics&list_uids=6809) | s13592 | GGCACGAGAUGAAACGAAAtt | UUUCGUUUCAUCUCGUGCCtt |
| NM_004177 | [STX3](http://www.ncbi.nlm.nih.gov/entrez/query.fcgi?db=gene&cmd=search&term=STX3) | [6809](http://www.ncbi.nlm.nih.gov/entrez/query.fcgi?db=gene&cmd=Retrieve&dopt=Graphics&list_uids=6809) | s13593 | CGGCUUUUAUGGACGAGUUtt | AACUCGUCCAUAAAAGCCGtg |
| NM_004177 | [STX3](http://www.ncbi.nlm.nih.gov/entrez/query.fcgi?db=gene&cmd=search&term=STX3) | [6809](http://www.ncbi.nlm.nih.gov/entrez/query.fcgi?db=gene&cmd=Retrieve&dopt=Graphics&list_uids=6809) | s13594 | GGUGAGAUGUUAGAUAACAtt | UGUUAUCUAACAUCUCACCct |
| NM_004604 | [STX4](http://www.ncbi.nlm.nih.gov/entrez/query.fcgi?db=gene&cmd=search&term=STX4) | [6810](http://www.ncbi.nlm.nih.gov/entrez/query.fcgi?db=gene&cmd=Retrieve&dopt=Graphics&list_uids=6810) | s13595 | GGACAAUUCGGCAGACUAUtt | AUAGUCUGCCGAAUUGUCCgg |
| NM_004604 | [STX4](http://www.ncbi.nlm.nih.gov/entrez/query.fcgi?db=gene&cmd=search&term=STX4) | [6810](http://www.ncbi.nlm.nih.gov/entrez/query.fcgi?db=gene&cmd=Retrieve&dopt=Graphics&list_uids=6810) | s13596 | CAGCAAUUCGUGGAGCUCAtt | UGAGCUCCACGAAUUGCUGgg |
| NM_004604 | [STX4](http://www.ncbi.nlm.nih.gov/entrez/query.fcgi?db=gene&cmd=search&term=STX4) | [6810](http://www.ncbi.nlm.nih.gov/entrez/query.fcgi?db=gene&cmd=Retrieve&dopt=Graphics&list_uids=6810) | s13597 | UGAUCAAUCGGAUUGAGAAtt | UUCUCAAUCCGAUUGAUCAtc |
| NM_003164 | [STX5](http://www.ncbi.nlm.nih.gov/entrez/query.fcgi?db=gene&cmd=search&term=STX5) | [6811](http://www.ncbi.nlm.nih.gov/entrez/query.fcgi?db=gene&cmd=Retrieve&dopt=Graphics&list_uids=6811) | s13599 | GAACAUUGAGUCGACAAUUtt | AAUUGUCGACUCAAUGUUCtg |
| NM_003164 | [STX5](http://www.ncbi.nlm.nih.gov/entrez/query.fcgi?db=gene&cmd=search&term=STX5) | [6811](http://www.ncbi.nlm.nih.gov/entrez/query.fcgi?db=gene&cmd=Retrieve&dopt=Graphics&list_uids=6811) | s13598 | GAGCUAACAUAUAUCAUCAtt | UGAUGAUAUAUGUUAGCUCtt |
| NM_003164 | [STX5](http://www.ncbi.nlm.nih.gov/entrez/query.fcgi?db=gene&cmd=search&term=STX5) | [6811](http://www.ncbi.nlm.nih.gov/entrez/query.fcgi?db=gene&cmd=Retrieve&dopt=Graphics&list_uids=6811) | s13600 | GCCCAUUCAGAGAUCCUCAtt | UGAGGAUCUCUGAAUGGGCgg |
| NM_005819 | [STX6](http://www.ncbi.nlm.nih.gov/entrez/query.fcgi?db=gene&cmd=search&term=STX6) | [10228](http://www.ncbi.nlm.nih.gov/entrez/query.fcgi?db=gene&cmd=Retrieve&dopt=Graphics&list_uids=10228) | s19959 | GCAACUGAAUUGAGUAUAAtt | UUAUACUCAAUUCAGUUGCat |
| NM_005819 | [STX6](http://www.ncbi.nlm.nih.gov/entrez/query.fcgi?db=gene&cmd=search&term=STX6) | [10228](http://www.ncbi.nlm.nih.gov/entrez/query.fcgi?db=gene&cmd=Retrieve&dopt=Graphics&list_uids=10228) | s19960 | GCAGUUAUGUUGGAAGAUUtt | AAUCUUCCAACAUAACUGCct |
| NM_005819 | [STX6](http://www.ncbi.nlm.nih.gov/entrez/query.fcgi?db=gene&cmd=search&term=STX6) | [10228](http://www.ncbi.nlm.nih.gov/entrez/query.fcgi?db=gene&cmd=Retrieve&dopt=Graphics&list_uids=10228) | s19958 | CCAACGAGCUGAGAAAUAAtt | UUAUUUCUCAGCUCGUUGGtg |
| NM_003569 | [STX7](http://www.ncbi.nlm.nih.gov/entrez/query.fcgi?db=gene&cmd=search&term=STX7) | [8417](http://www.ncbi.nlm.nih.gov/entrez/query.fcgi?db=gene&cmd=Retrieve&dopt=Graphics&list_uids=8417) | s15979 | GAAUGAUGAUUCAUGAACAtt | UGUUCAUGAAUCAUCAUUCcc |
| NM_003569 | [STX7](http://www.ncbi.nlm.nih.gov/entrez/query.fcgi?db=gene&cmd=search&term=STX7) | [8417](http://www.ncbi.nlm.nih.gov/entrez/query.fcgi?db=gene&cmd=Retrieve&dopt=Graphics&list_uids=8417) | s15978 | CAUCAUCAUUCUUAUCCUUtt | AAGGAUAAGAAUGAUGAUGca |
| NM_003569 | [STX7](http://www.ncbi.nlm.nih.gov/entrez/query.fcgi?db=gene&cmd=search&term=STX7) | [8417](http://www.ncbi.nlm.nih.gov/entrez/query.fcgi?db=gene&cmd=Retrieve&dopt=Graphics&list_uids=8417) | s15980 | GAGUUGCGAUUAUCAGUCUtt | AGACUGAUAAUCGCAACUCca |
| NM_004853 | [STX8](http://www.ncbi.nlm.nih.gov/entrez/query.fcgi?db=gene&cmd=search&term=STX8) | [9482](http://www.ncbi.nlm.nih.gov/entrez/query.fcgi?db=gene&cmd=Retrieve&dopt=Graphics&list_uids=9482) | s18183 | GGAUGAUCUUGUAACUCGAtt | UCGAGUUACAAGAUCAUCCaa |
| NM_004853 | [STX8](http://www.ncbi.nlm.nih.gov/entrez/query.fcgi?db=gene&cmd=search&term=STX8) | [9482](http://www.ncbi.nlm.nih.gov/entrez/query.fcgi?db=gene&cmd=Retrieve&dopt=Graphics&list_uids=9482) | s18184 | GCACCAAAGCUUACCGUGAtt | UCACGGUAAGCUUUGGUGCct |
| NM_004853 | [STX8](http://www.ncbi.nlm.nih.gov/entrez/query.fcgi?db=gene&cmd=search&term=STX8) | [9482](http://www.ncbi.nlm.nih.gov/entrez/query.fcgi?db=gene&cmd=Retrieve&dopt=Graphics&list_uids=9482) | s18182 | CGAAAUCAAUAUGAACGAAtt | UUCGUUCAUAUUGAUUUCGtt |
| NM_003165 | [STXBP1](http://www.ncbi.nlm.nih.gov/entrez/query.fcgi?db=gene&cmd=search&term=STXBP1) | [6812](http://www.ncbi.nlm.nih.gov/entrez/query.fcgi?db=gene&cmd=Retrieve&dopt=Graphics&list_uids=6812) | s13601 | CCUUAUAUCUCUACCCGUUtt | AACGGGUAGAGAUAUAAGGgt |
| NM_003165 | [STXBP1](http://www.ncbi.nlm.nih.gov/entrez/query.fcgi?db=gene&cmd=search&term=STXBP1) | [6812](http://www.ncbi.nlm.nih.gov/entrez/query.fcgi?db=gene&cmd=Retrieve&dopt=Graphics&list_uids=6812) | s13602 | GGUGGACUCCGAUUAUCAAtt | UUGAUAAUCGGAGUCCACCgt |
| NM_003165 | [STXBP1](http://www.ncbi.nlm.nih.gov/entrez/query.fcgi?db=gene&cmd=search&term=STXBP1) | [6812](http://www.ncbi.nlm.nih.gov/entrez/query.fcgi?db=gene&cmd=Retrieve&dopt=Graphics&list_uids=6812) | s13603 | GCAUAACGAUUGUGGAAGAtt | UCUUCCACAAUCGUUAUGCcc |
| NM_006949 | [STXBP2](http://www.ncbi.nlm.nih.gov/entrez/query.fcgi?db=gene&cmd=search&term=STXBP2) | [6813](http://www.ncbi.nlm.nih.gov/entrez/query.fcgi?db=gene&cmd=Retrieve&dopt=Graphics&list_uids=6813) | s227459 | CGGACAAGGCGAACAUCAAtt | UUGAUGUUCGCCUUGUCCGtg |
| NM_006949 | [STXBP2](http://www.ncbi.nlm.nih.gov/entrez/query.fcgi?db=gene&cmd=search&term=STXBP2) | [6813](http://www.ncbi.nlm.nih.gov/entrez/query.fcgi?db=gene&cmd=Retrieve&dopt=Graphics&list_uids=6813) | s13604 | GGAGCUGAAUAAGUAUUCUtt | AGAAUACUUAUUCAGCUCCtt |
| NM_006949 | [STXBP2](http://www.ncbi.nlm.nih.gov/entrez/query.fcgi?db=gene&cmd=search&term=STXBP2) | [6813](http://www.ncbi.nlm.nih.gov/entrez/query.fcgi?db=gene&cmd=Retrieve&dopt=Graphics&list_uids=6813) | s227460 | GGUUCAGGCCCUGAUCAAAtt | UUUGAUCAGGGCCUGAACCga |
| NM_007269 | [STXBP3](http://www.ncbi.nlm.nih.gov/entrez/query.fcgi?db=gene&cmd=search&term=STXBP3) | [6814](http://www.ncbi.nlm.nih.gov/entrez/query.fcgi?db=gene&cmd=Retrieve&dopt=Graphics&list_uids=6814) | s13608 | GAGAGUGACAUGAUUCGUAtt | UACGAAUCAUGUCACUCUCat |
| NM_007269 | [STXBP3](http://www.ncbi.nlm.nih.gov/entrez/query.fcgi?db=gene&cmd=search&term=STXBP3) | [6814](http://www.ncbi.nlm.nih.gov/entrez/query.fcgi?db=gene&cmd=Retrieve&dopt=Graphics&list_uids=6814) | s13607 | GCAUAUGAUCUACUACCAAtt | UUGGUAGUAGAUCAUAUGCca |
| NM_007269 | [STXBP3](http://www.ncbi.nlm.nih.gov/entrez/query.fcgi?db=gene&cmd=search&term=STXBP3) | [6814](http://www.ncbi.nlm.nih.gov/entrez/query.fcgi?db=gene&cmd=Retrieve&dopt=Graphics&list_uids=6814) | s13609 | GUAAAUCGGAGAACAAGUAtt | UACUUGUUCUCCGAUUUACtt |
| NM_178509 | [STXBP4](http://www.ncbi.nlm.nih.gov/entrez/query.fcgi?db=gene&cmd=search&term=STXBP4) | [252983](http://www.ncbi.nlm.nih.gov/entrez/query.fcgi?db=gene&cmd=Retrieve&dopt=Graphics&list_uids=252983) | s48386 | GUGCCUUACUUGAUAUGGAtt | UCCAUAUCAAGUAAGGCACga |
| NM_178509 | [STXBP4](http://www.ncbi.nlm.nih.gov/entrez/query.fcgi?db=gene&cmd=search&term=STXBP4) | [252983](http://www.ncbi.nlm.nih.gov/entrez/query.fcgi?db=gene&cmd=Retrieve&dopt=Graphics&list_uids=252983) | s48385 | CAACUUGUCUCAGUCAACAtt | UGUUGACUGAGACAAGUUGat |
| NM_178509 | [STXBP4](http://www.ncbi.nlm.nih.gov/entrez/query.fcgi?db=gene&cmd=search&term=STXBP4) | [252983](http://www.ncbi.nlm.nih.gov/entrez/query.fcgi?db=gene&cmd=Retrieve&dopt=Graphics&list_uids=252983) | s48384 | GAAGGCCCAUUGGUAUAUAtt | UAUAUACCAAUGGGCCUUCat |
| NM_139244 | [STXBP5](http://www.ncbi.nlm.nih.gov/entrez/query.fcgi?db=gene&cmd=search&term=STXBP5) | [134957](http://www.ncbi.nlm.nih.gov/entrez/query.fcgi?db=gene&cmd=Retrieve&dopt=Graphics&list_uids=134957) | s43906 | CGAUGCUUGAAGUUCGAUUtt | AAUCGAACUUCAAGCAUCGga |
| NM_139244 | [STXBP5](http://www.ncbi.nlm.nih.gov/entrez/query.fcgi?db=gene&cmd=search&term=STXBP5) | [134957](http://www.ncbi.nlm.nih.gov/entrez/query.fcgi?db=gene&cmd=Retrieve&dopt=Graphics&list_uids=134957) | s225638 | GAGACUUACUUAUAGUCAAtt | UUGACUAUAAGUAAGUCUCtg |
| NM_139244 | [STXBP5](http://www.ncbi.nlm.nih.gov/entrez/query.fcgi?db=gene&cmd=search&term=STXBP5) | [134957](http://www.ncbi.nlm.nih.gov/entrez/query.fcgi?db=gene&cmd=Retrieve&dopt=Graphics&list_uids=134957) | s43905 | GGAUAACUCCUUUAGCCGAtt | UCGGCUAAAGGAGUUAUCCtt |
| NM_014178 | [STXBP6](http://www.ncbi.nlm.nih.gov/entrez/query.fcgi?db=gene&cmd=search&term=STXBP6) | [29091](http://www.ncbi.nlm.nih.gov/entrez/query.fcgi?db=gene&cmd=Retrieve&dopt=Graphics&list_uids=29091) | s26467 | CCAAAAUUAUGGGAGGAAAtt | UUUCCUCCCAUAAUUUUGGat |
| NM_014178 | [STXBP6](http://www.ncbi.nlm.nih.gov/entrez/query.fcgi?db=gene&cmd=search&term=STXBP6) | [29091](http://www.ncbi.nlm.nih.gov/entrez/query.fcgi?db=gene&cmd=Retrieve&dopt=Graphics&list_uids=29091) | s26465 | AAGGCGAAUAUUUAACUUAtt | UAAGUUAAAUAUUCGCCUUga |
| NM_014178 | [STXBP6](http://www.ncbi.nlm.nih.gov/entrez/query.fcgi?db=gene&cmd=search&term=STXBP6) | [29091](http://www.ncbi.nlm.nih.gov/entrez/query.fcgi?db=gene&cmd=Retrieve&dopt=Graphics&list_uids=29091) | s26466 | UCACAAAGGUCAAACAGUUtt | AACUGUUUGACCUUUGUGAtg |
| NM_145003 | [TSNARE1](http://www.ncbi.nlm.nih.gov/entrez/query.fcgi?db=gene&cmd=search&term=TSNARE1) | [203062](http://www.ncbi.nlm.nih.gov/entrez/query.fcgi?db=gene&cmd=Retrieve&dopt=Graphics&list_uids=203062) | s47464 | AGCUGUUGAUAGUAUUGAAtt | UUCAAUACUAUCAACAGCUtc |
| NM_145003 | [TSNARE1](http://www.ncbi.nlm.nih.gov/entrez/query.fcgi?db=gene&cmd=search&term=TSNARE1) | [203062](http://www.ncbi.nlm.nih.gov/entrez/query.fcgi?db=gene&cmd=Retrieve&dopt=Graphics&list_uids=203062) | s47465 | GGAGACCAACAAGACCAUUtt | AAUGGUCUUGUUGGUCUCCtg |
| NM_145003 | [TSNARE1](http://www.ncbi.nlm.nih.gov/entrez/query.fcgi?db=gene&cmd=search&term=TSNARE1) | [203062](http://www.ncbi.nlm.nih.gov/entrez/query.fcgi?db=gene&cmd=Retrieve&dopt=Graphics&list_uids=203062) | s47463 | GCUAUGGAGUGGUGCAGAAtt | UUCUGCACCACUCCAUAGCac |
| NM_006370 | [VTI1B](http://www.ncbi.nlm.nih.gov/entrez/query.fcgi?db=gene&cmd=search&term=VTI1B) | [10490](http://www.ncbi.nlm.nih.gov/entrez/query.fcgi?db=gene&cmd=Retrieve&dopt=Graphics&list_uids=10490) | s20556 | CGAGACCAGUUAGAACGUAtt | UACGUUCUAACUGGUCUCGtt |
| NM_006370 | [VTI1B](http://www.ncbi.nlm.nih.gov/entrez/query.fcgi?db=gene&cmd=search&term=VTI1B) | [10490](http://www.ncbi.nlm.nih.gov/entrez/query.fcgi?db=gene&cmd=Retrieve&dopt=Graphics&list_uids=10490) | s20557 | GGACCUUGCUAAACUCCAUtt | AUGGAGUUUAGCAAGGUCCtt |
| NM_006370 | [VTI1B](http://www.ncbi.nlm.nih.gov/entrez/query.fcgi?db=gene&cmd=search&term=VTI1B) | [10490](http://www.ncbi.nlm.nih.gov/entrez/query.fcgi?db=gene&cmd=Retrieve&dopt=Graphics&list_uids=10490) | s20558 | GCUCAGAAAUCAUAGAAGAtt | UCUUCUAUGAUUUCUGAGCca |
| **Calpains** |  |  |  |  |  |
| NM_005186 | [CAPN1](http://www.ncbi.nlm.nih.gov/entrez/query.fcgi?db=gene&cmd=search&term=CAPN1) | [823](http://www.ncbi.nlm.nih.gov/entrez/query.fcgi?db=gene&cmd=Retrieve&dopt=Graphics&list_uids=823) | s490 | GGAAUUACCUGUCCAUCUUtt | AAGAUGGACAGGUAAUUCCgg |
| NM_005186 | [CAPN1](http://www.ncbi.nlm.nih.gov/entrez/query.fcgi?db=gene&cmd=search&term=CAPN1) | [823](http://www.ncbi.nlm.nih.gov/entrez/query.fcgi?db=gene&cmd=Retrieve&dopt=Graphics&list_uids=823) | s491 | GGAACAACGUGGACCCAUAtt | UAUGGGUCCACGUUGUUCCac |
| NM_005186 | [CAPN1](http://www.ncbi.nlm.nih.gov/entrez/query.fcgi?db=gene&cmd=search&term=CAPN1) | [823](http://www.ncbi.nlm.nih.gov/entrez/query.fcgi?db=gene&cmd=Retrieve&dopt=Graphics&list_uids=823) | s492 | CGUGUUCCAUAUAGAGGAAtt | UUCCUCUAUAUGGAACACGgg |
| NM_001748 | [CAPN2](http://www.ncbi.nlm.nih.gov/entrez/query.fcgi?db=gene&cmd=search&term=CAPN2) | [824](http://www.ncbi.nlm.nih.gov/entrez/query.fcgi?db=gene&cmd=Retrieve&dopt=Graphics&list_uids=824) | s320 | CCAGCGAUACCUACAAGAAtt | UUCUUGUAGGUAUCGCUGGtg |
| NM_001748 | [CAPN2](http://www.ncbi.nlm.nih.gov/entrez/query.fcgi?db=gene&cmd=search&term=CAPN2) | [824](http://www.ncbi.nlm.nih.gov/entrez/query.fcgi?db=gene&cmd=Retrieve&dopt=Graphics&list_uids=824) | s321 | CGAGAAUACUGGAACAAUAtt | UAUUGUUCCAGUAUUCUCGgg |
| NM_001748 | [CAPN2](http://www.ncbi.nlm.nih.gov/entrez/query.fcgi?db=gene&cmd=search&term=CAPN2) | [824](http://www.ncbi.nlm.nih.gov/entrez/query.fcgi?db=gene&cmd=Retrieve&dopt=Graphics&list_uids=824) | s319 | GAUCAACGGAUGCUAUGAAtt | UUCAUAGCAUCCGUUGAUCtt |
| NM_173090 | [CAPN3](http://www.ncbi.nlm.nih.gov/entrez/query.fcgi?db=gene&cmd=search&term=CAPN3) | [825](http://www.ncbi.nlm.nih.gov/entrez/query.fcgi?db=gene&cmd=Retrieve&dopt=Graphics&list_uids=825) | s2383 | CCACCUCUGGAACAAGAUUtt | AAUCUUGUUCCAGAGGUGGtg |
| NM_173090 | [CAPN3](http://www.ncbi.nlm.nih.gov/entrez/query.fcgi?db=gene&cmd=search&term=CAPN3) | [825](http://www.ncbi.nlm.nih.gov/entrez/query.fcgi?db=gene&cmd=Retrieve&dopt=Graphics&list_uids=825) | s2384 | GCAUGAUUGCGCUCAUGGAtt | UCCAUGAGCGCAAUCAUGCta |
| NM_173090 | [CAPN3](http://www.ncbi.nlm.nih.gov/entrez/query.fcgi?db=gene&cmd=search&term=CAPN3) | [825](http://www.ncbi.nlm.nih.gov/entrez/query.fcgi?db=gene&cmd=Retrieve&dopt=Graphics&list_uids=825) | s2382 | ACAGCUACGAGAUGCGAAAtt | UUUCGCAUCUCGUAGCUGUtg |
| **Annexins** |  |  |  |  |  |
| NM_000700 | [ANXA1](http://www.ncbi.nlm.nih.gov/entrez/query.fcgi?db=gene&cmd=search&term=ANXA1) | [301](http://www.ncbi.nlm.nih.gov/entrez/query.fcgi?db=gene&cmd=Retrieve&dopt=Graphics&list_uids=301) | s1382 | GACGUAAACGUGUUCAAUAtt | UAUUGAACACGUUUACGUCtg |
| NM_000700 | [ANXA1](http://www.ncbi.nlm.nih.gov/entrez/query.fcgi?db=gene&cmd=search&term=ANXA1) | [301](http://www.ncbi.nlm.nih.gov/entrez/query.fcgi?db=gene&cmd=Retrieve&dopt=Graphics&list_uids=301) | s1381 | GCCUCACAGCUAUCGUGAAtt | UUCACGAUAGCUGUGAGGCat |
| NM_000700 | [ANXA1](http://www.ncbi.nlm.nih.gov/entrez/query.fcgi?db=gene&cmd=search&term=ANXA1) | [301](http://www.ncbi.nlm.nih.gov/entrez/query.fcgi?db=gene&cmd=Retrieve&dopt=Graphics&list_uids=301) | s1380 | GGACUUUGGUGUGAAUGAAtt | UUCAUUCACACCAAAGUCCtc |
| NM_145869 | [ANXA11](http://www.ncbi.nlm.nih.gov/entrez/query.fcgi?db=gene&cmd=search&term=ANXA11) | [311](http://www.ncbi.nlm.nih.gov/entrez/query.fcgi?db=gene&cmd=Retrieve&dopt=Graphics&list_uids=311) | s1401 | GAGCACAUCCGAGAAUUAAtt | UUAAUUCUCGGAUGUGCUCat |
| NM_145869 | [ANXA11](http://www.ncbi.nlm.nih.gov/entrez/query.fcgi?db=gene&cmd=search&term=ANXA11) | [311](http://www.ncbi.nlm.nih.gov/entrez/query.fcgi?db=gene&cmd=Retrieve&dopt=Graphics&list_uids=311) | s1403 | CAGUCCUCUUUGACAUUUAtt | UAAAUGUCAAAGAGGACUGgg |
| NM_145869 | [ANXA11](http://www.ncbi.nlm.nih.gov/entrez/query.fcgi?db=gene&cmd=search&term=ANXA11) | [311](http://www.ncbi.nlm.nih.gov/entrez/query.fcgi?db=gene&cmd=Retrieve&dopt=Graphics&list_uids=311) | s1402 | UCAAAGAUCUGAAAUCUGAtt | UCAGAUUUCAGAUCUUUGAtc |
| NM_004039 | [ANXA2](http://www.ncbi.nlm.nih.gov/entrez/query.fcgi?db=gene&cmd=search&term=ANXA2)  (siRNA 2) | [302](http://www.ncbi.nlm.nih.gov/entrez/query.fcgi?db=gene&cmd=Retrieve&dopt=Graphics&list_uids=302) | s1384 | GCAAGUCCCUGUACUAUUAtt | UAAUAGUACAGGGACUUGCcg |
| NM_004039 | [ANXA2](http://www.ncbi.nlm.nih.gov/entrez/query.fcgi?db=gene&cmd=search&term=ANXA2)  (siRNA 3) | [302](http://www.ncbi.nlm.nih.gov/entrez/query.fcgi?db=gene&cmd=Retrieve&dopt=Graphics&list_uids=302) | s1385 | GAACUUGCAUCAGCACUGAtt | UCAGUGCUGAUGCAAGUUCct |
| NM_004039 | [ANXA2](http://www.ncbi.nlm.nih.gov/entrez/query.fcgi?db=gene&cmd=search&term=ANXA2)  (siRNA 1) | [302](http://www.ncbi.nlm.nih.gov/entrez/query.fcgi?db=gene&cmd=Retrieve&dopt=Graphics&list_uids=302) | s1383 | GGCAAAGGGUAGAAGAGCAtt | UGCUCUUCUACCCUUUGCCag |
| NM_005139 | [ANXA3](http://www.ncbi.nlm.nih.gov/entrez/query.fcgi?db=gene&cmd=search&term=ANXA3) | [306](http://www.ncbi.nlm.nih.gov/entrez/query.fcgi?db=gene&cmd=Retrieve&dopt=Graphics&list_uids=306) | s1386 | CAUCGAGCCUUGAAGGGUAtt | UACCCUUCAAGGCUCGAUGca |
| NM_005139 | [ANXA3](http://www.ncbi.nlm.nih.gov/entrez/query.fcgi?db=gene&cmd=search&term=ANXA3) | [306](http://www.ncbi.nlm.nih.gov/entrez/query.fcgi?db=gene&cmd=Retrieve&dopt=Graphics&list_uids=306) | s1388 | GCAAUUAAAUCGGAUACUUtt | AAGUAUCCGAUUUAAUUGCtg |
| NM_005139 | [ANXA3](http://www.ncbi.nlm.nih.gov/entrez/query.fcgi?db=gene&cmd=search&term=ANXA3) | [306](http://www.ncbi.nlm.nih.gov/entrez/query.fcgi?db=gene&cmd=Retrieve&dopt=Graphics&list_uids=306) | s1387 | GAAGAUGCCUUGAUUGAAAtt | UUUCAAUCAAGGCAUCUUCgt |
| NM_001153 | [ANXA4](http://www.ncbi.nlm.nih.gov/entrez/query.fcgi?db=gene&cmd=search&term=ANXA4) | [307](http://www.ncbi.nlm.nih.gov/entrez/query.fcgi?db=gene&cmd=Retrieve&dopt=Graphics&list_uids=307) | s1389 | GCAGGGACUUGAUAGACGAtt | UCGUCUAUCAAGUCCCUGCcg |
| NM_001153 | [ANXA4](http://www.ncbi.nlm.nih.gov/entrez/query.fcgi?db=gene&cmd=search&term=ANXA4) | [307](http://www.ncbi.nlm.nih.gov/entrez/query.fcgi?db=gene&cmd=Retrieve&dopt=Graphics&list_uids=307) | s1390 | GGAUAUCACAGAAGGAUAUtt | AUAUCCUUCUGUGAUAUCCtt |
| NM_001153 | [ANXA4](http://www.ncbi.nlm.nih.gov/entrez/query.fcgi?db=gene&cmd=search&term=ANXA4) | [307](http://www.ncbi.nlm.nih.gov/entrez/query.fcgi?db=gene&cmd=Retrieve&dopt=Graphics&list_uids=307) | s1391 | GAAAUUGACAUGUUGGAUAtt | UAUCCAACAUGUCAAUUUCtg |
| NM_001154 | [ANXA5](http://www.ncbi.nlm.nih.gov/entrez/query.fcgi?db=gene&cmd=search&term=ANXA5) | [308](http://www.ncbi.nlm.nih.gov/entrez/query.fcgi?db=gene&cmd=Retrieve&dopt=Graphics&list_uids=308) | s1392 | GUACAUGACUAUAUCAGGAtt | UCCUGAUAUAGUCAUGUACtt |
| NM_001154 | [ANXA5](http://www.ncbi.nlm.nih.gov/entrez/query.fcgi?db=gene&cmd=search&term=ANXA5) | [308](http://www.ncbi.nlm.nih.gov/entrez/query.fcgi?db=gene&cmd=Retrieve&dopt=Graphics&list_uids=308) | s1393 | GACCUGAAAUCAGAACUAAtt | UUAGUUCUGAUUUCAGGUCat |
| NM_001154 | [ANXA5](http://www.ncbi.nlm.nih.gov/entrez/query.fcgi?db=gene&cmd=search&term=ANXA5) | [308](http://www.ncbi.nlm.nih.gov/entrez/query.fcgi?db=gene&cmd=Retrieve&dopt=Graphics&list_uids=308) | s1394 | GGAGUGAGAUUGAUCUGUUtt | AACAGAUCAAUCUCACUCCtg |
| NM_001155 | [ANXA6](http://www.ncbi.nlm.nih.gov/entrez/query.fcgi?db=gene&cmd=search&term=ANXA6) | [309](http://www.ncbi.nlm.nih.gov/entrez/query.fcgi?db=gene&cmd=Retrieve&dopt=Graphics&list_uids=309) | s1397 | GUAGUGAAGUGUAUCCGGAtt | UCCGGAUACACUUCACUACgg |
| NM_001155 | [ANXA6](http://www.ncbi.nlm.nih.gov/entrez/query.fcgi?db=gene&cmd=search&term=ANXA6) | [309](http://www.ncbi.nlm.nih.gov/entrez/query.fcgi?db=gene&cmd=Retrieve&dopt=Graphics&list_uids=309) | s1396 | GGUUGGUGUUCGAUGAGUAtt | UACUCAUCGAACACCAACCga |
| NM_001155 | [ANXA6](http://www.ncbi.nlm.nih.gov/entrez/query.fcgi?db=gene&cmd=search&term=ANXA6) | [309](http://www.ncbi.nlm.nih.gov/entrez/query.fcgi?db=gene&cmd=Retrieve&dopt=Graphics&list_uids=309) | s1395 | GGCUCUUCAAGGCUAUGAAtt | UUCAUAGCCUUGAAGAGCCtt |
| NM_004034 | [ANXA7](http://www.ncbi.nlm.nih.gov/entrez/query.fcgi?db=gene&cmd=search&term=ANXA7) | [310](http://www.ncbi.nlm.nih.gov/entrez/query.fcgi?db=gene&cmd=Retrieve&dopt=Graphics&list_uids=310) | s1400 | GGCUAAUCGAGACUUGUUAtt | UAACAAGUCUCGAUUAGCCat |
| NM_004034 | [ANXA7](http://www.ncbi.nlm.nih.gov/entrez/query.fcgi?db=gene&cmd=search&term=ANXA7) | [310](http://www.ncbi.nlm.nih.gov/entrez/query.fcgi?db=gene&cmd=Retrieve&dopt=Graphics&list_uids=310) | s1398 | GAAUCUUGCUUUAACAUGAtt | UCAUGUUAAAGCAAGAUUCat |
| NM_004034 | [ANXA7](http://www.ncbi.nlm.nih.gov/entrez/query.fcgi?db=gene&cmd=search&term=ANXA7) | [310](http://www.ncbi.nlm.nih.gov/entrez/query.fcgi?db=gene&cmd=Retrieve&dopt=Graphics&list_uids=310) | s1399 | GAUCUCAAAUCAGAGUUAAtt | UUAACUCUGAUUUGAGAUCtt |
| **Endocytosis** |  |  |  |  |  |
| NM_004408 | [DNM1](http://www.ncbi.nlm.nih.gov/entrez/query.fcgi?db=gene&cmd=search&term=DNM1) | [1759](http://www.ncbi.nlm.nih.gov/entrez/query.fcgi?db=gene&cmd=Retrieve&dopt=Graphics&list_uids=1759) | s144 | GCAGUUCGCCGUAGACUUUtt | AAAGUCUACGGCGAACUGCtg |
| NM_004408 | [DNM1](http://www.ncbi.nlm.nih.gov/entrez/query.fcgi?db=gene&cmd=search&term=DNM1) | [1759](http://www.ncbi.nlm.nih.gov/entrez/query.fcgi?db=gene&cmd=Retrieve&dopt=Graphics&list_uids=1759) | s145 | GCUAUGCUAUCAAGAAUAUtt | AUAUUCUUGAUAGCAUAGCtg |
| NM_004408 | [DNM1](http://www.ncbi.nlm.nih.gov/entrez/query.fcgi?db=gene&cmd=search&term=DNM1) | [1759](http://www.ncbi.nlm.nih.gov/entrez/query.fcgi?db=gene&cmd=Retrieve&dopt=Graphics&list_uids=1759) | s146 | GCACUGCAAGGGAAAGAAAtt | UUUCUUUCCCUUGCAGUGCag |
| NM_001005360 | [DNM2](http://www.ncbi.nlm.nih.gov/entrez/query.fcgi?db=gene&cmd=search&term=DNM2) | [1785](http://www.ncbi.nlm.nih.gov/entrez/query.fcgi?db=gene&cmd=Retrieve&dopt=Graphics&list_uids=1785) | s4212 | ACAUCAACACGAACCAUGAtt | UCAUGGUUCGUGUUGAUGUag |
| NM_001005360 | [DNM2](http://www.ncbi.nlm.nih.gov/entrez/query.fcgi?db=gene&cmd=search&term=DNM2) | [1785](http://www.ncbi.nlm.nih.gov/entrez/query.fcgi?db=gene&cmd=Retrieve&dopt=Graphics&list_uids=1785) | s4213 | AGUCCUACAUCAACACGAAtt | UUCGUGUUGAUGUAGGACUgc |
| NM_001005360 | [DNM2](http://www.ncbi.nlm.nih.gov/entrez/query.fcgi?db=gene&cmd=search&term=DNM2) | [1785](http://www.ncbi.nlm.nih.gov/entrez/query.fcgi?db=gene&cmd=Retrieve&dopt=Graphics&list_uids=1785) | s4214 | AGCGAAUCGUCACCACUUAtt | UAAGUGGUGACGAUUCGCUct |
| NM_015569 | [DNM3](http://www.ncbi.nlm.nih.gov/entrez/query.fcgi?db=gene&cmd=search&term=DNM3) | [26052](http://www.ncbi.nlm.nih.gov/entrez/query.fcgi?db=gene&cmd=Retrieve&dopt=Graphics&list_uids=26052) | s25019 | GUGUGGAUCUGGUAAUACAtt | UGUAUUACCAGAUCCACACtc |
| NM_015569 | [DNM3](http://www.ncbi.nlm.nih.gov/entrez/query.fcgi?db=gene&cmd=search&term=DNM3) | [26052](http://www.ncbi.nlm.nih.gov/entrez/query.fcgi?db=gene&cmd=Retrieve&dopt=Graphics&list_uids=26052) | s25017 | GCAUUUGAAGCGAUAGUCAtt | UGACUAUCGCUUCAAAUGCca |
| NM_015569 | [DNM3](http://www.ncbi.nlm.nih.gov/entrez/query.fcgi?db=gene&cmd=search&term=DNM3) | [26052](http://www.ncbi.nlm.nih.gov/entrez/query.fcgi?db=gene&cmd=Retrieve&dopt=Graphics&list_uids=26052) | s25018 | GACUCCUACAUGUCCAUUAtt | UAAUGGACAUGUAGGAGUCta |
| NM_001076677 | [CLTA](http://www.ncbi.nlm.nih.gov/entrez/query.fcgi?db=gene&cmd=search&term=CLTA) | [1211](http://www.ncbi.nlm.nih.gov/entrez/query.fcgi?db=gene&cmd=Retrieve&dopt=Graphics&list_uids=1211) | s3190 | GGACGAGCAGCUACAGAAAtt | UUUCUGUAGCUGCUCGUCCtg |
| NM_001076677 | [CLTA](http://www.ncbi.nlm.nih.gov/entrez/query.fcgi?db=gene&cmd=search&term=CLTA) | [1211](http://www.ncbi.nlm.nih.gov/entrez/query.fcgi?db=gene&cmd=Retrieve&dopt=Graphics&list_uids=1211) | s3189 | AGCCUGAAAGUAUCCGUAAtt | UUACGGAUACUUUCAGGCUct |
| NM_001076677 | [CLTA](http://www.ncbi.nlm.nih.gov/entrez/query.fcgi?db=gene&cmd=search&term=CLTA) | [1211](http://www.ncbi.nlm.nih.gov/entrez/query.fcgi?db=gene&cmd=Retrieve&dopt=Graphics&list_uids=1211) | s3188 | AGACAGUUAUGCAGCUAUUtt | AAUAGCUGCAUAACUGUCUgt |
| NM_001834 | [CLTB](http://www.ncbi.nlm.nih.gov/entrez/query.fcgi?db=gene&cmd=search&term=CLTB) | [1212](http://www.ncbi.nlm.nih.gov/entrez/query.fcgi?db=gene&cmd=Retrieve&dopt=Graphics&list_uids=1212) | s3191 | GCCCAGCUAUGUGACUUCAtt | UGAAGUCACAUAGCUGGGCca |
| NM_001834 | [CLTB](http://www.ncbi.nlm.nih.gov/entrez/query.fcgi?db=gene&cmd=search&term=CLTB) | [1212](http://www.ncbi.nlm.nih.gov/entrez/query.fcgi?db=gene&cmd=Retrieve&dopt=Graphics&list_uids=1212) | s3193 | GAACAAGUAGAGAAGAACAtt | UGUUCUUCUCUACUUGUUCac |
| NM_001834 | [CLTB](http://www.ncbi.nlm.nih.gov/entrez/query.fcgi?db=gene&cmd=search&term=CLTB) | [1212](http://www.ncbi.nlm.nih.gov/entrez/query.fcgi?db=gene&cmd=Retrieve&dopt=Graphics&list_uids=1212) | s3192 | CCUCCUCUCAGUCUACUCAtt | UGAGUAGACUGAGAGGAGGcg |
| NM_004859 | [CLTC](http://www.ncbi.nlm.nih.gov/entrez/query.fcgi?db=gene&cmd=search&term=CLTC) | [1213](http://www.ncbi.nlm.nih.gov/entrez/query.fcgi?db=gene&cmd=Retrieve&dopt=Graphics&list_uids=1213) | s475 | GGUUGCUCUUGUUACGGAUtt | AUCCGUAACAAGAGCAACCgt |
| NM_004859 | [CLTC](http://www.ncbi.nlm.nih.gov/entrez/query.fcgi?db=gene&cmd=search&term=CLTC) | [1213](http://www.ncbi.nlm.nih.gov/entrez/query.fcgi?db=gene&cmd=Retrieve&dopt=Graphics&list_uids=1213) | s477 | GGGAAUUCUUCGUACUCCAtt | UGGAGUACGAAGAAUUCCCtt |
| NM_004859 | [CLTC](http://www.ncbi.nlm.nih.gov/entrez/query.fcgi?db=gene&cmd=search&term=CLTC) | [1213](http://www.ncbi.nlm.nih.gov/entrez/query.fcgi?db=gene&cmd=Retrieve&dopt=Graphics&list_uids=1213) | s223262 | GAUUAUCAAUUACCGUACAtt | UGUACGGUAAUUGAUAAUCtg |
| NM_001753 | [CAV1](http://www.ncbi.nlm.nih.gov/entrez/query.fcgi?db=gene&cmd=search&term=CAV1) | [857](http://www.ncbi.nlm.nih.gov/entrez/query.fcgi?db=gene&cmd=Retrieve&dopt=Graphics&list_uids=857) | s2448 | CCUUCACUGUGACGAAAUAtt | UAUUUCGUCACAGUGAAGGtg |
| NM_001753 | [CAV1](http://www.ncbi.nlm.nih.gov/entrez/query.fcgi?db=gene&cmd=search&term=CAV1) | [857](http://www.ncbi.nlm.nih.gov/entrez/query.fcgi?db=gene&cmd=Retrieve&dopt=Graphics&list_uids=857) | s2446 | GCUUCCUGAUUGAGAUUCAtt | UGAAUCUCAAUCAGGAAGCtc |
| NM_001753 | [CAV1](http://www.ncbi.nlm.nih.gov/entrez/query.fcgi?db=gene&cmd=search&term=CAV1) | [857](http://www.ncbi.nlm.nih.gov/entrez/query.fcgi?db=gene&cmd=Retrieve&dopt=Graphics&list_uids=857) | s2447 | GCCGUGUCUAUUCCAUCUAtt | UAGAUGGAAUAGACACGGCtg |
| NM_001233 | [CAV2](http://www.ncbi.nlm.nih.gov/entrez/query.fcgi?db=gene&cmd=search&term=CAV2) | [858](http://www.ncbi.nlm.nih.gov/entrez/query.fcgi?db=gene&cmd=Retrieve&dopt=Graphics&list_uids=858) | s2450 | CGGCUCAACUCGCAUCUCAtt | UGAGAUGCGAGUUGAGCCGgt |
| NM_001233 | [CAV2](http://www.ncbi.nlm.nih.gov/entrez/query.fcgi?db=gene&cmd=search&term=CAV2) | [858](http://www.ncbi.nlm.nih.gov/entrez/query.fcgi?db=gene&cmd=Retrieve&dopt=Graphics&list_uids=858) | s2449 | AUGUUAUCAUUGCUCCAUUtt | AAUGGAGCAAUGAUAACAUct |
| NM_001233 | [CAV2](http://www.ncbi.nlm.nih.gov/entrez/query.fcgi?db=gene&cmd=search&term=CAV2) | [858](http://www.ncbi.nlm.nih.gov/entrez/query.fcgi?db=gene&cmd=Retrieve&dopt=Graphics&list_uids=858) | s2451 | AGACCUGCCUAAUGGUUCUtt | AGAACCAUUAGGCAGGUCUtt |
| NM_001234 | [CAV3](http://www.ncbi.nlm.nih.gov/entrez/query.fcgi?db=gene&cmd=search&term=CAV3) | [859](http://www.ncbi.nlm.nih.gov/entrez/query.fcgi?db=gene&cmd=Retrieve&dopt=Graphics&list_uids=859) | s2453 | AGAUCGUCAAGGAUAUCCAtt | UGGAUAUCCUUGACGAUCUgg |
| NM_001234 | [CAV3](http://www.ncbi.nlm.nih.gov/entrez/query.fcgi?db=gene&cmd=search&term=CAV3) | [859](http://www.ncbi.nlm.nih.gov/entrez/query.fcgi?db=gene&cmd=Retrieve&dopt=Graphics&list_uids=859) | s2452 | GAACAUUAACGAGGACAUAtt | UAUGUCCUCGUUAAUGUUCtt |
| NM_001234 | [CAV3](http://www.ncbi.nlm.nih.gov/entrez/query.fcgi?db=gene&cmd=search&term=CAV3) | [859](http://www.ncbi.nlm.nih.gov/entrez/query.fcgi?db=gene&cmd=Retrieve&dopt=Graphics&list_uids=859) | s225003 | GGAACAAGGUCUAAUUUGUtt | ACAAAUUAGACCUUGUUCCtt |
| NM_005803 | [FLOT1](http://www.ncbi.nlm.nih.gov/entrez/query.fcgi?db=gene&cmd=search&term=FLOT1) | [10211](http://www.ncbi.nlm.nih.gov/entrez/query.fcgi?db=gene&cmd=Retrieve&dopt=Graphics&list_uids=10211) | s19913 | GCAGAGAAGUCCCAACUAAtt | UUAGUUGGGACUUCUCUGCct |
| NM_005803 | [FLOT1](http://www.ncbi.nlm.nih.gov/entrez/query.fcgi?db=gene&cmd=search&term=FLOT1) | [10211](http://www.ncbi.nlm.nih.gov/entrez/query.fcgi?db=gene&cmd=Retrieve&dopt=Graphics&list_uids=10211) | s19914 | GCAUCAGUGUGGUUAGCUAtt | UAGCUAACCACACUGAUGCcc |
| NM_005803 | [FLOT1](http://www.ncbi.nlm.nih.gov/entrez/query.fcgi?db=gene&cmd=search&term=FLOT1) | [10211](http://www.ncbi.nlm.nih.gov/entrez/query.fcgi?db=gene&cmd=Retrieve&dopt=Graphics&list_uids=10211) | s19915 | AGAGAGAUUACGAACUGAAtt | UUCAGUUCGUAAUCUCUCUgt |
| NM_004475 | [FLOT2](http://www.ncbi.nlm.nih.gov/entrez/query.fcgi?db=gene&cmd=search&term=FLOT2) | [2319](http://www.ncbi.nlm.nih.gov/entrez/query.fcgi?db=gene&cmd=Retrieve&dopt=Graphics&list_uids=2319) | s5284 | CCAAGAUUGCUGACUCUAAtt | UUAGAGUCAGCAAUCUUGGtg |
| NM_004475 | [FLOT2](http://www.ncbi.nlm.nih.gov/entrez/query.fcgi?db=gene&cmd=search&term=FLOT2) | [2319](http://www.ncbi.nlm.nih.gov/entrez/query.fcgi?db=gene&cmd=Retrieve&dopt=Graphics&list_uids=2319) | s5286 | GACUAUAAACAGUACGUGUtt | ACACGUACUGUUUAUAGUCgg |
| NM_004475 | [FLOT2](http://www.ncbi.nlm.nih.gov/entrez/query.fcgi?db=gene&cmd=search&term=FLOT2) | [2319](http://www.ncbi.nlm.nih.gov/entrez/query.fcgi?db=gene&cmd=Retrieve&dopt=Graphics&list_uids=2319) | s5285 | ACAGUAAGGUCACAUCAGAtt | UCUGAUGUGACCUUACUGUtg |
| **Adaptor complexes** |  |  |  |  |  |
| NM_001127 | [AP1B1](http://www.ncbi.nlm.nih.gov/entrez/query.fcgi?db=gene&cmd=search&term=AP1B1) | [162](http://www.ncbi.nlm.nih.gov/entrez/query.fcgi?db=gene&cmd=Retrieve&dopt=Graphics&list_uids=162) | s1141 | GAGCUGAAAGAGUACGCAAtt | UUGCGUACUCUUUCAGCUCtg |
| NM_001127 | [AP1B1](http://www.ncbi.nlm.nih.gov/entrez/query.fcgi?db=gene&cmd=search&term=AP1B1) | [162](http://www.ncbi.nlm.nih.gov/entrez/query.fcgi?db=gene&cmd=Retrieve&dopt=Graphics&list_uids=162) | s1139 | GCAACCUGCUCGAUCUGAAtt | UUCAGAUCGAGCAGGUUGCtg |
| NM_001127 | [AP1B1](http://www.ncbi.nlm.nih.gov/entrez/query.fcgi?db=gene&cmd=search&term=AP1B1) | [162](http://www.ncbi.nlm.nih.gov/entrez/query.fcgi?db=gene&cmd=Retrieve&dopt=Graphics&list_uids=162) | s1140 | CCAUCUCCAUGGACCUGCAtt | UGCAGGUCCAUGGAGAUGGag |
| NM_001128 | [AP1G1](http://www.ncbi.nlm.nih.gov/entrez/query.fcgi?db=gene&cmd=search&term=AP1G1) | [164](http://www.ncbi.nlm.nih.gov/entrez/query.fcgi?db=gene&cmd=Retrieve&dopt=Graphics&list_uids=164) | s1144 | GGUACGAAUUUUGCGGUUAtt | UAACCGCAAAAUUCGUACCtg |
| NM_001128 | [AP1G1](http://www.ncbi.nlm.nih.gov/entrez/query.fcgi?db=gene&cmd=search&term=AP1G1) | [164](http://www.ncbi.nlm.nih.gov/entrez/query.fcgi?db=gene&cmd=Retrieve&dopt=Graphics&list_uids=164) | s1143 | GGGAGGAAAUGACAUAACAtt | UGUUAUGUCAUUUCCUCCCaa |
| NM_001128 | [AP1G1](http://www.ncbi.nlm.nih.gov/entrez/query.fcgi?db=gene&cmd=search&term=AP1G1) | [164](http://www.ncbi.nlm.nih.gov/entrez/query.fcgi?db=gene&cmd=Retrieve&dopt=Graphics&list_uids=164) | s1142 | GAUAUUCACCAGAACAUGAtt | UCAUGUUCUGGUGAAUAUCcg |
| NM_003917 | [AP1G2](http://www.ncbi.nlm.nih.gov/entrez/query.fcgi?db=gene&cmd=search&term=AP1G2) | [8906](http://www.ncbi.nlm.nih.gov/entrez/query.fcgi?db=gene&cmd=Retrieve&dopt=Graphics&list_uids=8906) | s17030 | GCAUAUCUGGAGUCAGCGAtt | UCGCUGACUCCAGAUAUGCtg |
| NM_003917 | [AP1G2](http://www.ncbi.nlm.nih.gov/entrez/query.fcgi?db=gene&cmd=search&term=AP1G2) | [8906](http://www.ncbi.nlm.nih.gov/entrez/query.fcgi?db=gene&cmd=Retrieve&dopt=Graphics&list_uids=8906) | s17031 | GGUUCUAGCUGUCAACAUUtt | AAUGUUGACAGCUAGAACCcg |
| NM_003917 | [AP1G2](http://www.ncbi.nlm.nih.gov/entrez/query.fcgi?db=gene&cmd=search&term=AP1G2) | [8906](http://www.ncbi.nlm.nih.gov/entrez/query.fcgi?db=gene&cmd=Retrieve&dopt=Graphics&list_uids=8906) | s17029 | GGUCCUGUUUGAGACAGUAtt | UACUGUCUCAAACAGGACCgc |
| NM_080550 | [AP1GBP1](http://www.ncbi.nlm.nih.gov/entrez/query.fcgi?db=gene&cmd=search&term=AP1GBP1) | [11276](http://www.ncbi.nlm.nih.gov/entrez/query.fcgi?db=gene&cmd=Retrieve&dopt=Graphics&list_uids=11276) | s22267 | GGCCUUAGCUAAUCGAACUtt | AGUUCGAUUAGCUAAGGCCca |
| NM_080550 | [AP1GBP1](http://www.ncbi.nlm.nih.gov/entrez/query.fcgi?db=gene&cmd=search&term=AP1GBP1) | [11276](http://www.ncbi.nlm.nih.gov/entrez/query.fcgi?db=gene&cmd=Retrieve&dopt=Graphics&list_uids=11276) | s22266 | CAACCUACCCUGCAAAUCAtt | UGAUUUGCAGGGUAGGUUGgt |
| NM_080550 | [AP1GBP1](http://www.ncbi.nlm.nih.gov/entrez/query.fcgi?db=gene&cmd=search&term=AP1GBP1) | [11276](http://www.ncbi.nlm.nih.gov/entrez/query.fcgi?db=gene&cmd=Retrieve&dopt=Graphics&list_uids=11276) | s22265 | GACUCCAACUGGAAUAGAUtt | AUCUAUUCCAGUUGGAGUCat |
| NM_032493 | [AP1M1](http://www.ncbi.nlm.nih.gov/entrez/query.fcgi?db=gene&cmd=search&term=AP1M1) | [8907](http://www.ncbi.nlm.nih.gov/entrez/query.fcgi?db=gene&cmd=Retrieve&dopt=Graphics&list_uids=8907) | s17032 | ACAACUUUGUUAUCAUCUAtt | UAGAUGAUAACAAAGUUGUcc |
| NM_032493 | [AP1M1](http://www.ncbi.nlm.nih.gov/entrez/query.fcgi?db=gene&cmd=search&term=AP1M1) | [8907](http://www.ncbi.nlm.nih.gov/entrez/query.fcgi?db=gene&cmd=Retrieve&dopt=Graphics&list_uids=8907) | s225055 | GGCUAUCACGCUUCGAGAAtt | UUCUCGAAGCGUGAUAGCCgc |
| NM_032493 | [AP1M1](http://www.ncbi.nlm.nih.gov/entrez/query.fcgi?db=gene&cmd=search&term=AP1M1) | [8907](http://www.ncbi.nlm.nih.gov/entrez/query.fcgi?db=gene&cmd=Retrieve&dopt=Graphics&list_uids=8907) | s17033 | UCAAGUUCGAGAUCCCUUAtt | UAAGGGAUCUCGAACUUGAca |
| NM_005498 | [AP1M2](http://www.ncbi.nlm.nih.gov/entrez/query.fcgi?db=gene&cmd=search&term=AP1M2) | [10053](http://www.ncbi.nlm.nih.gov/entrez/query.fcgi?db=gene&cmd=Retrieve&dopt=Graphics&list_uids=10053) | s19546 | GGAGAUAUCUGUGCCUGUAtt | UACAGGCACAGAUAUCUCCac |
| NM_005498 | [AP1M2](http://www.ncbi.nlm.nih.gov/entrez/query.fcgi?db=gene&cmd=search&term=AP1M2) | [10053](http://www.ncbi.nlm.nih.gov/entrez/query.fcgi?db=gene&cmd=Retrieve&dopt=Graphics&list_uids=10053) | s19545 | GCUCCGAGGGUAUCAAGUAtt | UACUUGAUACCCUCGGAGCgc |
| NM_005498 | [AP1M2](http://www.ncbi.nlm.nih.gov/entrez/query.fcgi?db=gene&cmd=search&term=AP1M2) | [10053](http://www.ncbi.nlm.nih.gov/entrez/query.fcgi?db=gene&cmd=Retrieve&dopt=Graphics&list_uids=10053) | s19544 | GUCGUGAUUUGGAGUAUUAtt | UAAUACUCCAAAUCACGACgt |
| NM_001283 | [AP1S1](http://www.ncbi.nlm.nih.gov/entrez/query.fcgi?db=gene&cmd=search&term=AP1S1) | [1174](http://www.ncbi.nlm.nih.gov/entrez/query.fcgi?db=gene&cmd=Retrieve&dopt=Graphics&list_uids=1174) | s3116 | GGACCUCAAAGUUGUCUAUtt | AUAGACAACUUUGAGGUCCct |
| NM_001283 | [AP1S1](http://www.ncbi.nlm.nih.gov/entrez/query.fcgi?db=gene&cmd=search&term=AP1S1) | [1174](http://www.ncbi.nlm.nih.gov/entrez/query.fcgi?db=gene&cmd=Retrieve&dopt=Graphics&list_uids=1174) | s3115 | GGAAAACUGCGGCUGCAAAtt | UUUGCAGCCGCAGUUUUCCct |
| NM_001283 | [AP1S1](http://www.ncbi.nlm.nih.gov/entrez/query.fcgi?db=gene&cmd=search&term=AP1S1) | [1174](http://www.ncbi.nlm.nih.gov/entrez/query.fcgi?db=gene&cmd=Retrieve&dopt=Graphics&list_uids=1174) | s3117 | ACGUGGAGCUCUUAGACAAtt | UUGUCUAAGAGCUCCACGUat |
| NM_003916 | [AP1S2](http://www.ncbi.nlm.nih.gov/entrez/query.fcgi?db=gene&cmd=search&term=AP1S2) | [8905](http://www.ncbi.nlm.nih.gov/entrez/query.fcgi?db=gene&cmd=Retrieve&dopt=Graphics&list_uids=8905) | s225054 | AAUUCAUCGUUAUGUGGAAtt | UUCCACAUAACGAUGAAUUat |
| NM_003916 | [AP1S2](http://www.ncbi.nlm.nih.gov/entrez/query.fcgi?db=gene&cmd=search&term=AP1S2) | [8905](http://www.ncbi.nlm.nih.gov/entrez/query.fcgi?db=gene&cmd=Retrieve&dopt=Graphics&list_uids=8905) | s17028 | AGACCGUUUUAGCACGGAAtt | UUCCGUGCUAAAACGGUCUga |
| NM_003916 | [AP1S2](http://www.ncbi.nlm.nih.gov/entrez/query.fcgi?db=gene&cmd=search&term=AP1S2) | [8905](http://www.ncbi.nlm.nih.gov/entrez/query.fcgi?db=gene&cmd=Retrieve&dopt=Graphics&list_uids=8905) | s17027 | AGAUCACAAGAGAACUUGUtt | ACAAGUUCUCUUGUGAUCUtt |
| NM_001039569 | [AP1S3](http://www.ncbi.nlm.nih.gov/entrez/query.fcgi?db=gene&cmd=search&term=AP1S3) | [130340](http://www.ncbi.nlm.nih.gov/entrez/query.fcgi?db=gene&cmd=Retrieve&dopt=Graphics&list_uids=130340) | s43489 | GGAAAUUACGGCUACAGAAtt | UUCUGUAGCCGUAAUUUCCct |
| NM_001039569 | [AP1S3](http://www.ncbi.nlm.nih.gov/entrez/query.fcgi?db=gene&cmd=search&term=AP1S3) | [130340](http://www.ncbi.nlm.nih.gov/entrez/query.fcgi?db=gene&cmd=Retrieve&dopt=Graphics&list_uids=130340) | s43488 | GUUUAUAAAAGGUAUGCUAtt | UAGCAUACCUUUUAUAAACaa |
| NM_001039569 | [AP1S3](http://www.ncbi.nlm.nih.gov/entrez/query.fcgi?db=gene&cmd=search&term=AP1S3) | [130340](http://www.ncbi.nlm.nih.gov/entrez/query.fcgi?db=gene&cmd=Retrieve&dopt=Graphics&list_uids=130340) | s43490 | CCUGGACGAGUUUAUAAUAtt | UAUUAUAAACUCGUCCAGGat |
| NM_018569 | [AP1AR](http://www.ncbi.nlm.nih.gov/entrez/query.fcgi?db=gene&cmd=search&term=C4orf16) | [55435](http://www.ncbi.nlm.nih.gov/entrez/query.fcgi?db=gene&cmd=Retrieve&dopt=Graphics&list_uids=55435) | s30872 | GACUGGAAGUAAUCCUACAtt | UGUAGGAUUACUUCCAGUCtt |
| NM_018569 | [AP1AR](http://www.ncbi.nlm.nih.gov/entrez/query.fcgi?db=gene&cmd=search&term=C4orf16) | [55435](http://www.ncbi.nlm.nih.gov/entrez/query.fcgi?db=gene&cmd=Retrieve&dopt=Graphics&list_uids=55435) | s30870 | GAAGUUUUUCGAAGUAGUAtt | UACUACUUCGAAAAACUUCat |
| NM_018569 | [AP1AR](http://www.ncbi.nlm.nih.gov/entrez/query.fcgi?db=gene&cmd=search&term=C4orf16) | [55435](http://www.ncbi.nlm.nih.gov/entrez/query.fcgi?db=gene&cmd=Retrieve&dopt=Graphics&list_uids=55435) | s30871 | GAAAGCAGUUGUGAUUUAAtt | UUAAAUCACAACUGCUUUCtg |
| NM_130787 | [AP2A1](http://www.ncbi.nlm.nih.gov/entrez/query.fcgi?db=gene&cmd=search&term=AP2A1) | [160](http://www.ncbi.nlm.nih.gov/entrez/query.fcgi?db=gene&cmd=Retrieve&dopt=Graphics&list_uids=160) | s183 | GCAGAUCGGAGUCAAGUCAtt | UGACUUGACUCCGAUCUGCag |
| NM_130787 | [AP2A1](http://www.ncbi.nlm.nih.gov/entrez/query.fcgi?db=gene&cmd=search&term=AP2A1) | [160](http://www.ncbi.nlm.nih.gov/entrez/query.fcgi?db=gene&cmd=Retrieve&dopt=Graphics&list_uids=160) | s184 | GCCGAUGAGUUGCUGAAUAtt | UAUUCAGCAACUCAUCGGCtt |
| NM_130787 | [AP2A1](http://www.ncbi.nlm.nih.gov/entrez/query.fcgi?db=gene&cmd=search&term=AP2A1) | [160](http://www.ncbi.nlm.nih.gov/entrez/query.fcgi?db=gene&cmd=Retrieve&dopt=Graphics&list_uids=160) | s185 | CAUCCGCUCCAAGUUCAAAtt | UUUGAACUUGGAGCGGAUGtt |
| NM_012305 | [AP2A2](http://www.ncbi.nlm.nih.gov/entrez/query.fcgi?db=gene&cmd=search&term=AP2A2) | [161](http://www.ncbi.nlm.nih.gov/entrez/query.fcgi?db=gene&cmd=Retrieve&dopt=Graphics&list_uids=161) | s1137 | GCUGCGCACAAGUAAGGAAtt | UUCCUUACUUGUGCGCAGCgt |
| NM_012305 | [AP2A2](http://www.ncbi.nlm.nih.gov/entrez/query.fcgi?db=gene&cmd=search&term=AP2A2) | [161](http://www.ncbi.nlm.nih.gov/entrez/query.fcgi?db=gene&cmd=Retrieve&dopt=Graphics&list_uids=161) | s1138 | GGCAAAUAUCAGAUCAAAAtt | UUUUGAUCUGAUAUUUGCCag |
| NM_012305 | [AP2A2](http://www.ncbi.nlm.nih.gov/entrez/query.fcgi?db=gene&cmd=search&term=AP2A2) | [161](http://www.ncbi.nlm.nih.gov/entrez/query.fcgi?db=gene&cmd=Retrieve&dopt=Graphics&list_uids=161) | s1136 | CCUGCUGAGUUCAAACAGAtt | UCUGUUUGAACUCAGCAGGtt |
| NM_001282 | [AP2B1](http://www.ncbi.nlm.nih.gov/entrez/query.fcgi?db=gene&cmd=search&term=AP2B1) | [163](http://www.ncbi.nlm.nih.gov/entrez/query.fcgi?db=gene&cmd=Retrieve&dopt=Graphics&list_uids=163) | s36 | GGAUCCCUAUGUUCGGAAAtt | UUUCCGAACAUAGGGAUCCtc |
| NM_001282 | [AP2B1](http://www.ncbi.nlm.nih.gov/entrez/query.fcgi?db=gene&cmd=search&term=AP2B1) | [163](http://www.ncbi.nlm.nih.gov/entrez/query.fcgi?db=gene&cmd=Retrieve&dopt=Graphics&list_uids=163) | s37 | GGAGGGUUUUCACGAUGAAtt | UUCAUCGUGAAAACCCUCCag |
| NM_001282 | [AP2B1](http://www.ncbi.nlm.nih.gov/entrez/query.fcgi?db=gene&cmd=search&term=AP2B1) | [163](http://www.ncbi.nlm.nih.gov/entrez/query.fcgi?db=gene&cmd=Retrieve&dopt=Graphics&list_uids=163) | s38 | GGCUUGGAGAUUUCCGGAAtt | UUCCGGAAAUCUCCAAGCCtt |
| NM_004068 | [AP2M1](http://www.ncbi.nlm.nih.gov/entrez/query.fcgi?db=gene&cmd=search&term=AP2M1) | [1173](http://www.ncbi.nlm.nih.gov/entrez/query.fcgi?db=gene&cmd=Retrieve&dopt=Graphics&list_uids=1173) | s3113 | GGCGAGAGGGUAUCAAGUAtt | UACUUGAUACCCUCUCGCCgc |
| NM_004068 | [AP2M1](http://www.ncbi.nlm.nih.gov/entrez/query.fcgi?db=gene&cmd=search&term=AP2M1) | [1173](http://www.ncbi.nlm.nih.gov/entrez/query.fcgi?db=gene&cmd=Retrieve&dopt=Graphics&list_uids=1173) | s3112 | GGAUGAGAUUCUAGACUUUtt | AAAGUCUAGAAUCUCAUCCag |
| NM_004068 | [AP2M1](http://www.ncbi.nlm.nih.gov/entrez/query.fcgi?db=gene&cmd=search&term=AP2M1) | [1173](http://www.ncbi.nlm.nih.gov/entrez/query.fcgi?db=gene&cmd=Retrieve&dopt=Graphics&list_uids=1173) | s3114 | GAAGAGCAGUCACAGAUCAtt | UGAUCUGUGACUGCUCUUCtt |
| NM_021575 | [AP2S1](http://www.ncbi.nlm.nih.gov/entrez/query.fcgi?db=gene&cmd=search&term=AP2S1) | [1175](http://www.ncbi.nlm.nih.gov/entrez/query.fcgi?db=gene&cmd=Retrieve&dopt=Graphics&list_uids=1175) | s3118 | AGGUUUACACGGUCGUGGAtt | UCCACGACCGUGUAAACCUtg |
| NM_021575 | [AP2S1](http://www.ncbi.nlm.nih.gov/entrez/query.fcgi?db=gene&cmd=search&term=AP2S1) | [1175](http://www.ncbi.nlm.nih.gov/entrez/query.fcgi?db=gene&cmd=Retrieve&dopt=Graphics&list_uids=1175) | s3119 | GGCGAAAUCCGAGAGACCAtt | UGGUCUCUCGGAUUUCGCCag |
| NM_021575 | [AP2S1](http://www.ncbi.nlm.nih.gov/entrez/query.fcgi?db=gene&cmd=search&term=AP2S1) | [1175](http://www.ncbi.nlm.nih.gov/entrez/query.fcgi?db=gene&cmd=Retrieve&dopt=Graphics&list_uids=1175) | s194335 | AGACGAAGGUGCUGAAACAtt | UGUUUCAGCACCUUCGUCUgg |
| NM_003664 | [AP3B1](http://www.ncbi.nlm.nih.gov/entrez/query.fcgi?db=gene&cmd=search&term=AP3B1) | [8546](http://www.ncbi.nlm.nih.gov/entrez/query.fcgi?db=gene&cmd=Retrieve&dopt=Graphics&list_uids=8546) | s16268 | GCUUAUUGUUCCGAAUGUAtt | UACAUUCGGAACAAUAAGCtg |
| NM_003664 | [AP3B1](http://www.ncbi.nlm.nih.gov/entrez/query.fcgi?db=gene&cmd=search&term=AP3B1) | [8546](http://www.ncbi.nlm.nih.gov/entrez/query.fcgi?db=gene&cmd=Retrieve&dopt=Graphics&list_uids=8546) | s16267 | CUUGCACUCCUGUCCAUAAtt | UUAUGGACAGGAGUGCAAGat |
| NM_003664 | [AP3B1](http://www.ncbi.nlm.nih.gov/entrez/query.fcgi?db=gene&cmd=search&term=AP3B1) | [8546](http://www.ncbi.nlm.nih.gov/entrez/query.fcgi?db=gene&cmd=Retrieve&dopt=Graphics&list_uids=8546) | s16266 | CCGCAAGCUAUGUAACUUAtt | UAAGUUACAUAGCUUGCGGta |
| NM_004644 | [AP3B2](http://www.ncbi.nlm.nih.gov/entrez/query.fcgi?db=gene&cmd=search&term=AP3B2) | [8120](http://www.ncbi.nlm.nih.gov/entrez/query.fcgi?db=gene&cmd=Retrieve&dopt=Graphics&list_uids=8120) | s15638 | CCAUCAUGAUGCUAGCUAUtt | AUAGCUAGCAUCAUGAUGGgc |
| NM_004644 | [AP3B2](http://www.ncbi.nlm.nih.gov/entrez/query.fcgi?db=gene&cmd=search&term=AP3B2) | [8120](http://www.ncbi.nlm.nih.gov/entrez/query.fcgi?db=gene&cmd=Retrieve&dopt=Graphics&list_uids=8120) | s15640 | CGAUUCCAGCAGUAGCUCAtt | UGAGCUACUGCUGGAAUCGct |
| NM_004644 | [AP3B2](http://www.ncbi.nlm.nih.gov/entrez/query.fcgi?db=gene&cmd=search&term=AP3B2) | [8120](http://www.ncbi.nlm.nih.gov/entrez/query.fcgi?db=gene&cmd=Retrieve&dopt=Graphics&list_uids=8120) | s15639 | GCAUCGACCUGAUUCACAAtt | UUGUGAAUCAGGUCGAUGCgc |
| NM_001077523 | [AP3D1](http://www.ncbi.nlm.nih.gov/entrez/query.fcgi?db=gene&cmd=search&term=AP3D1) | [8943](http://www.ncbi.nlm.nih.gov/entrez/query.fcgi?db=gene&cmd=Retrieve&dopt=Graphics&list_uids=8943) | s17106 | GCCUAUGUCAGAUCAGUAUtt | AUACUGAUCUGACAUAGGCag |
| NM_001077523 | [AP3D1](http://www.ncbi.nlm.nih.gov/entrez/query.fcgi?db=gene&cmd=search&term=AP3D1) | [8943](http://www.ncbi.nlm.nih.gov/entrez/query.fcgi?db=gene&cmd=Retrieve&dopt=Graphics&list_uids=8943) | s17108 | CGCUGAAAAUUCCUAUGUUtt | AACAUAGGAAUUUUCAGCGag |
| NM_001077523 | [AP3D1](http://www.ncbi.nlm.nih.gov/entrez/query.fcgi?db=gene&cmd=search&term=AP3D1) | [8943](http://www.ncbi.nlm.nih.gov/entrez/query.fcgi?db=gene&cmd=Retrieve&dopt=Graphics&list_uids=8943) | s17107 | ACAGGGCUCUGGAUAUUGAtt | UCAAUAUCCAGAGCCCUGUag |
| NM_207012 | [AP3M1](http://www.ncbi.nlm.nih.gov/entrez/query.fcgi?db=gene&cmd=search&term=AP3M1) | [26985](http://www.ncbi.nlm.nih.gov/entrez/query.fcgi?db=gene&cmd=Retrieve&dopt=Graphics&list_uids=26985) | s25662 | GGACUACUUUGGUGAGUGUtt | ACACUCACCAAAGUAGUCCtg |
| NM_207012 | [AP3M1](http://www.ncbi.nlm.nih.gov/entrez/query.fcgi?db=gene&cmd=search&term=AP3M1) | [26985](http://www.ncbi.nlm.nih.gov/entrez/query.fcgi?db=gene&cmd=Retrieve&dopt=Graphics&list_uids=26985) | s25663 | CCAAGGUACUAACAUGGGAtt | UCCCAUGUUAGUACCUUGGtg |
| NM_207012 | [AP3M1](http://www.ncbi.nlm.nih.gov/entrez/query.fcgi?db=gene&cmd=search&term=AP3M1) | [26985](http://www.ncbi.nlm.nih.gov/entrez/query.fcgi?db=gene&cmd=Retrieve&dopt=Graphics&list_uids=26985) | s25661 | GAAUUACAGUGACAGUUCAtt | UGAACUGUCACUGUAAUUCct |
| NM_006803 | [AP3M2](http://www.ncbi.nlm.nih.gov/entrez/query.fcgi?db=gene&cmd=search&term=AP3M2) | [10947](http://www.ncbi.nlm.nih.gov/entrez/query.fcgi?db=gene&cmd=Retrieve&dopt=Graphics&list_uids=10947) | s21537 | CAGUGUAUGUCAAACAUAAtt | UUAUGUUUGACAUACACUGgg |
| NM_006803 | [AP3M2](http://www.ncbi.nlm.nih.gov/entrez/query.fcgi?db=gene&cmd=search&term=AP3M2) | [10947](http://www.ncbi.nlm.nih.gov/entrez/query.fcgi?db=gene&cmd=Retrieve&dopt=Graphics&list_uids=10947) | s21539 | CAAUUAACCUGCAGUUUAAtt | UUAAACUGCAGGUUAAUUGtg |
| NM_006803 | [AP3M2](http://www.ncbi.nlm.nih.gov/entrez/query.fcgi?db=gene&cmd=search&term=AP3M2) | [10947](http://www.ncbi.nlm.nih.gov/entrez/query.fcgi?db=gene&cmd=Retrieve&dopt=Graphics&list_uids=10947) | s21538 | CGAACAUUCUUAAAGAACUtt | AGUUCUUUAAGAAUGUUCGac |
| NM_001284 | [AP3S1](http://www.ncbi.nlm.nih.gov/entrez/query.fcgi?db=gene&cmd=search&term=AP3S1) | [1176](http://www.ncbi.nlm.nih.gov/entrez/query.fcgi?db=gene&cmd=Retrieve&dopt=Graphics&list_uids=1176) | s3122 | GGAUCUGACAACAAACUGAtt | UCAGUUUGUUGUCAGAUCCtc |
| NM_001284 | [AP3S1](http://www.ncbi.nlm.nih.gov/entrez/query.fcgi?db=gene&cmd=search&term=AP3S1) | [1176](http://www.ncbi.nlm.nih.gov/entrez/query.fcgi?db=gene&cmd=Retrieve&dopt=Graphics&list_uids=1176) | s3121 | GUAUCUAAGAGAGAUGAAAtt | UUUCAUCUCUCUUAGAUACca |
| NM_001284 | [AP3S1](http://www.ncbi.nlm.nih.gov/entrez/query.fcgi?db=gene&cmd=search&term=AP3S1) | [1176](http://www.ncbi.nlm.nih.gov/entrez/query.fcgi?db=gene&cmd=Retrieve&dopt=Graphics&list_uids=1176) | s3120 | CCCAAGAAAUAUUAACAUUtt | AAUGUUAAUAUUUCUUGGGat |
| NM_005829 | [AP3S2](http://www.ncbi.nlm.nih.gov/entrez/query.fcgi?db=gene&cmd=search&term=AP3S2) | [10239](http://www.ncbi.nlm.nih.gov/entrez/query.fcgi?db=gene&cmd=Retrieve&dopt=Graphics&list_uids=10239) | s19988 | CUGGAUAAGUGUUUCGAAAtt | UUUCGAAACACUUAUCCAGag |
| NM_005829 | [AP3S2](http://www.ncbi.nlm.nih.gov/entrez/query.fcgi?db=gene&cmd=search&term=AP3S2) | [10239](http://www.ncbi.nlm.nih.gov/entrez/query.fcgi?db=gene&cmd=Retrieve&dopt=Graphics&list_uids=10239) | s19989 | CUAUGCUACCCUCUACUUUtt | AAAGUAGAGGGUAGCAUAGtg |
| NM_005829 | [AP3S2](http://www.ncbi.nlm.nih.gov/entrez/query.fcgi?db=gene&cmd=search&term=AP3S2) | [10239](http://www.ncbi.nlm.nih.gov/entrez/query.fcgi?db=gene&cmd=Retrieve&dopt=Graphics&list_uids=10239) | s19990 | GGCUCUGACUACAAACUGAtt | UCAGUUUGUAGUCAGAGCCac |
| NM_006594 | [AP4B1](http://www.ncbi.nlm.nih.gov/entrez/query.fcgi?db=gene&cmd=search&term=AP4B1) | [10717](http://www.ncbi.nlm.nih.gov/entrez/query.fcgi?db=gene&cmd=Retrieve&dopt=Graphics&list_uids=10717) | s21051 | GAUCUUGCAUAGUUUACCAtt | UGGUAAACUAUGCAAGAUCtg |
| NM_006594 | [AP4B1](http://www.ncbi.nlm.nih.gov/entrez/query.fcgi?db=gene&cmd=search&term=AP4B1) | [10717](http://www.ncbi.nlm.nih.gov/entrez/query.fcgi?db=gene&cmd=Retrieve&dopt=Graphics&list_uids=10717) | s21052 | CGAGUGAUUAGGUACAUGAtt | UCAUGUACCUAAUCACUCGct |
| NM_006594 | [AP4B1](http://www.ncbi.nlm.nih.gov/entrez/query.fcgi?db=gene&cmd=search&term=AP4B1) | [10717](http://www.ncbi.nlm.nih.gov/entrez/query.fcgi?db=gene&cmd=Retrieve&dopt=Graphics&list_uids=10717) | s21050 | CGUACAAACUGAUGUCCUUtt | AAGGACAUCAGUUUGUACGtg |
| NM_007347 | [AP4E1](http://www.ncbi.nlm.nih.gov/entrez/query.fcgi?db=gene&cmd=search&term=AP4E1) | [23431](http://www.ncbi.nlm.nih.gov/entrez/query.fcgi?db=gene&cmd=Retrieve&dopt=Graphics&list_uids=23431) | s23818 | GAACUGCCCUUGGUUGAGAtt | UCUCAACCAAGGGCAGUUCtg |
| NM_007347 | [AP4E1](http://www.ncbi.nlm.nih.gov/entrez/query.fcgi?db=gene&cmd=search&term=AP4E1) | [23431](http://www.ncbi.nlm.nih.gov/entrez/query.fcgi?db=gene&cmd=Retrieve&dopt=Graphics&list_uids=23431) | s23819 | CCUAAUCAAGUACAACAUAtt | UAUGUUGUACUUGAUUAGGag |
| NM_007347 | [AP4E1](http://www.ncbi.nlm.nih.gov/entrez/query.fcgi?db=gene&cmd=search&term=AP4E1) | [23431](http://www.ncbi.nlm.nih.gov/entrez/query.fcgi?db=gene&cmd=Retrieve&dopt=Graphics&list_uids=23431) | s23817 | GUCUAUUCUAUUUAUCCUAtt | UAGGAUAAAUAGAAUAGACtg |
| NM_004722 | [AP4M1](http://www.ncbi.nlm.nih.gov/entrez/query.fcgi?db=gene&cmd=search&term=AP4M1) | [9179](http://www.ncbi.nlm.nih.gov/entrez/query.fcgi?db=gene&cmd=Retrieve&dopt=Graphics&list_uids=9179) | s17541 | GAGAUUGUCUGUACUGAUAtt | UAUCAGUACAGACAAUCUCtc |
| NM_004722 | [AP4M1](http://www.ncbi.nlm.nih.gov/entrez/query.fcgi?db=gene&cmd=search&term=AP4M1) | [9179](http://www.ncbi.nlm.nih.gov/entrez/query.fcgi?db=gene&cmd=Retrieve&dopt=Graphics&list_uids=9179) | s17539 | GGCUCCAGGUUUAUCUAAAtt | UUUAGAUAAACCUGGAGCCgg |
| NM_004722 | [AP4M1](http://www.ncbi.nlm.nih.gov/entrez/query.fcgi?db=gene&cmd=search&term=AP4M1) | [9179](http://www.ncbi.nlm.nih.gov/entrez/query.fcgi?db=gene&cmd=Retrieve&dopt=Graphics&list_uids=9179) | s17540 | GAAUUUGAGUCUCAUCGAAtt | UUCGAUGAGACUCAAAUUCgt |
| NM_007077 | [AP4S1](http://www.ncbi.nlm.nih.gov/entrez/query.fcgi?db=gene&cmd=search&term=AP4S1) | [11154](http://www.ncbi.nlm.nih.gov/entrez/query.fcgi?db=gene&cmd=Retrieve&dopt=Graphics&list_uids=11154) | s22002 | UCAUUGUGGUUGGAGUUAAtt | UUAACUCCAACCACAAUGAag |
| NM_001128126 | [AP4S1](http://www.ncbi.nlm.nih.gov/entrez/query.fcgi?db=gene&cmd=search&term=AP4S1) | [11154](http://www.ncbi.nlm.nih.gov/entrez/query.fcgi?db=gene&cmd=Retrieve&dopt=Graphics&list_uids=11154) | s230357 | GGCUAUUUAUGAAUUCAUUtt | AAUGAAUUCAUAAAUAGCCat |
| NM_001128126 | [AP4S1](http://www.ncbi.nlm.nih.gov/entrez/query.fcgi?db=gene&cmd=search&term=AP4S1) | [11154](http://www.ncbi.nlm.nih.gov/entrez/query.fcgi?db=gene&cmd=Retrieve&dopt=Graphics&list_uids=11154) | s230358 | GCCGAGUGAGUGAAUUAGAtt | UCUAAUUCACUCACUCGGCtg |
| NM_001658 | [ARF1](http://www.ncbi.nlm.nih.gov/entrez/query.fcgi?db=gene&cmd=search&term=ARF1) | [375](http://www.ncbi.nlm.nih.gov/entrez/query.fcgi?db=gene&cmd=Retrieve&dopt=Graphics&list_uids=375) | s1551 | CCAUAGGCUUCAACGUGGAtt | UCCACGUUGAAGCCUAUGGtg |
| NM_001658 | [ARF1](http://www.ncbi.nlm.nih.gov/entrez/query.fcgi?db=gene&cmd=search&term=ARF1) | [375](http://www.ncbi.nlm.nih.gov/entrez/query.fcgi?db=gene&cmd=Retrieve&dopt=Graphics&list_uids=375) | s1550 | CCAUUCCCACCAUAGGCUUtt | AAGCCUAUGGUGGGAAUGGtg |
| NM_001658 | [ARF1](http://www.ncbi.nlm.nih.gov/entrez/query.fcgi?db=gene&cmd=search&term=ARF1) | [375](http://www.ncbi.nlm.nih.gov/entrez/query.fcgi?db=gene&cmd=Retrieve&dopt=Graphics&list_uids=375) | s1552 | UCUUCGUGGUGGACAGCAAtt | UUGCUGUCCACCACGAAGAtc |
| **Coatamer proteins** |  |  |  |  |  |
| NM_004371 | [COPA](http://www.ncbi.nlm.nih.gov/entrez/query.fcgi?db=gene&cmd=search&term=COPA) | [1314](http://www.ncbi.nlm.nih.gov/entrez/query.fcgi?db=gene&cmd=Retrieve&dopt=Graphics&list_uids=1314) | s3369 | GGAUGAGAAAACUCGCUUUtt | AAAGCGAGUUUUCUCAUCCtt |
| NM_004371 | [COPA](http://www.ncbi.nlm.nih.gov/entrez/query.fcgi?db=gene&cmd=search&term=COPA) | [1314](http://www.ncbi.nlm.nih.gov/entrez/query.fcgi?db=gene&cmd=Retrieve&dopt=Graphics&list_uids=1314) | s3370 | GCAGAGACCAUGAUCGUUUtt | AAACGAUCAUGGUCUCUGCgg |
| NM_004371 | [COPA](http://www.ncbi.nlm.nih.gov/entrez/query.fcgi?db=gene&cmd=search&term=COPA) | [1314](http://www.ncbi.nlm.nih.gov/entrez/query.fcgi?db=gene&cmd=Retrieve&dopt=Graphics&list_uids=1314) | s3368 | GGAUGUGCACUCUCAUUGAtt | UCAAUGAGAGUGCACAUCCga |
| NM_016451 | [COPB1](http://www.ncbi.nlm.nih.gov/entrez/query.fcgi?db=gene&cmd=search&term=COPB1) | [1315](http://www.ncbi.nlm.nih.gov/entrez/query.fcgi?db=gene&cmd=Retrieve&dopt=Graphics&list_uids=1315) | s3373 | CCUCAUGACUUCGCAAAUAtt | UAUUUGCGAAGUCAUGAGGag |
| NM_016451 | [COPB1](http://www.ncbi.nlm.nih.gov/entrez/query.fcgi?db=gene&cmd=search&term=COPB1) | [1315](http://www.ncbi.nlm.nih.gov/entrez/query.fcgi?db=gene&cmd=Retrieve&dopt=Graphics&list_uids=1315) | s3371 | GGUCUGUCAUGCUAAUCCAtt | UGGAUUAGCAUGACAGACCtt |
| NM_016451 | [COPB1](http://www.ncbi.nlm.nih.gov/entrez/query.fcgi?db=gene&cmd=search&term=COPB1) | [1315](http://www.ncbi.nlm.nih.gov/entrez/query.fcgi?db=gene&cmd=Retrieve&dopt=Graphics&list_uids=1315) | s3372 | GGAGAUGUAAAGUCAAAGAtt | UCUUUGACUUUACAUCUCCtt |
| NM_004766 | [COPB2](http://www.ncbi.nlm.nih.gov/entrez/query.fcgi?db=gene&cmd=search&term=COPB2) | [9276](http://www.ncbi.nlm.nih.gov/entrez/query.fcgi?db=gene&cmd=Retrieve&dopt=Graphics&list_uids=9276) | s17736 | GGUCAAACAAUGUCGCUUUtt | AAAGCGACAUUGUUUGACCct |
| NM_004766 | [COPB2](http://www.ncbi.nlm.nih.gov/entrez/query.fcgi?db=gene&cmd=search&term=COPB2) | [9276](http://www.ncbi.nlm.nih.gov/entrez/query.fcgi?db=gene&cmd=Retrieve&dopt=Graphics&list_uids=9276) | s17738 | CGAUGUAUCUCCUAGGCUAtt | UAGCCUAGGAGAUACAUCGtc |
| NM_004766 | [COPB2](http://www.ncbi.nlm.nih.gov/entrez/query.fcgi?db=gene&cmd=search&term=COPB2) | [9276](http://www.ncbi.nlm.nih.gov/entrez/query.fcgi?db=gene&cmd=Retrieve&dopt=Graphics&list_uids=9276) | s17737 | CAGUAUCCACAGAUCCUGAtt | UCAGGAUCUGUGGAUACUGta |
| NM_199444 | [COPE](http://www.ncbi.nlm.nih.gov/entrez/query.fcgi?db=gene&cmd=search&term=COPE) | [11316](http://www.ncbi.nlm.nih.gov/entrez/query.fcgi?db=gene&cmd=Retrieve&dopt=Graphics&list_uids=11316) | s22308 | AGCUGUUCGACGUAAAGAAtt | UUCUUUACGUCGAACAGCUcg |
| NM_199444 | [COPE](http://www.ncbi.nlm.nih.gov/entrez/query.fcgi?db=gene&cmd=search&term=COPE) | [11316](http://www.ncbi.nlm.nih.gov/entrez/query.fcgi?db=gene&cmd=Retrieve&dopt=Graphics&list_uids=11316) | s223223 | CCUUCUACAUCGGCAGCUAtt | UAGCUGCCGAUGUAGAAGGcg |
| NM_199444 | [COPE](http://www.ncbi.nlm.nih.gov/entrez/query.fcgi?db=gene&cmd=search&term=COPE) | [11316](http://www.ncbi.nlm.nih.gov/entrez/query.fcgi?db=gene&cmd=Retrieve&dopt=Graphics&list_uids=11316) | s22309 | GUGACAAACCGAUACCUGUtt | ACAGGUAUCGGUUUGUCACct |
| NM_016128 | [COPG1](http://www.ncbi.nlm.nih.gov/entrez/query.fcgi?db=gene&cmd=search&term=COPG1) | [22820](http://www.ncbi.nlm.nih.gov/entrez/query.fcgi?db=gene&cmd=Retrieve&dopt=Graphics&list_uids=22820) | s22432 | GCGUAAGAAUGACCGCCUAtt | UAGGCGGUCAUUCUUACGCac |
| NM_016128 | [COPG1](http://www.ncbi.nlm.nih.gov/entrez/query.fcgi?db=gene&cmd=search&term=COPG1) | [22820](http://www.ncbi.nlm.nih.gov/entrez/query.fcgi?db=gene&cmd=Retrieve&dopt=Graphics&list_uids=22820) | s22430 | CUGUUUGACUUCAUCGAGAtt | UCUCGAUGAAGUCAAACAGtg |
| NM_016128 | [COPG1](http://www.ncbi.nlm.nih.gov/entrez/query.fcgi?db=gene&cmd=search&term=COPG1) | [22820](http://www.ncbi.nlm.nih.gov/entrez/query.fcgi?db=gene&cmd=Retrieve&dopt=Graphics&list_uids=22820) | s22431 | CGUCGGAUGUGCUACUUGAtt | UCAAGUAGCACAUCCGACGga |
| NM_012133 | [COPG2](http://www.ncbi.nlm.nih.gov/entrez/query.fcgi?db=gene&cmd=search&term=COPG2) | [26958](http://www.ncbi.nlm.nih.gov/entrez/query.fcgi?db=gene&cmd=Retrieve&dopt=Graphics&list_uids=26958) | s223713 | GGAUCACCGAUGGAACAAUtt | AUUGUUCCAUCGGUGAUCCtg |
| NM_012133 | [COPG2](http://www.ncbi.nlm.nih.gov/entrez/query.fcgi?db=gene&cmd=search&term=COPG2) | [26958](http://www.ncbi.nlm.nih.gov/entrez/query.fcgi?db=gene&cmd=Retrieve&dopt=Graphics&list_uids=26958) | s223714 | GGAUGGGCAUGAAAGUCCAtt | UGGACUUUCAUGCCCAUCCtc |
| NM_012133 | [COPG2](http://www.ncbi.nlm.nih.gov/entrez/query.fcgi?db=gene&cmd=search&term=COPG2) | [26958](http://www.ncbi.nlm.nih.gov/entrez/query.fcgi?db=gene&cmd=Retrieve&dopt=Graphics&list_uids=26958) | s25645 | GGGUGAACACUUUGGAACAtt | UGUUCCAAAGUGUUCACCCtg |
| NM_016057 | [COPZ1](http://www.ncbi.nlm.nih.gov/entrez/query.fcgi?db=gene&cmd=search&term=COPZ1) | [22818](http://www.ncbi.nlm.nih.gov/entrez/query.fcgi?db=gene&cmd=Retrieve&dopt=Graphics&list_uids=22818) | s22428 | GCCUGACAGUGGUAUACAAtt | UUGUAUACCACUGUCAGGCct |
| NM_016057 | [COPZ1](http://www.ncbi.nlm.nih.gov/entrez/query.fcgi?db=gene&cmd=search&term=COPZ1) | [22818](http://www.ncbi.nlm.nih.gov/entrez/query.fcgi?db=gene&cmd=Retrieve&dopt=Graphics&list_uids=22818) | s22429 | CCAUCGGACUGACAGUGAAtt | UUCACUGUCAGUCCGAUGGgt |
| NM_016057 | [COPZ1](http://www.ncbi.nlm.nih.gov/entrez/query.fcgi?db=gene&cmd=search&term=COPZ1) | [22818](http://www.ncbi.nlm.nih.gov/entrez/query.fcgi?db=gene&cmd=Retrieve&dopt=Graphics&list_uids=22818) | s22427 | GUAUAGAUCUCUAUUUCUAtt | UAGAAAUAGAGAUCUAUACtg |
| NM_016429 | [COPZ2](http://www.ncbi.nlm.nih.gov/entrez/query.fcgi?db=gene&cmd=search&term=COPZ2) | [51226](http://www.ncbi.nlm.nih.gov/entrez/query.fcgi?db=gene&cmd=Retrieve&dopt=Graphics&list_uids=51226) | s27713 | AGCAGAUGGUUUUCGAGAAtt | UUCUCGAAAACCAUCUGCUcc |
| NM_016429 | [COPZ2](http://www.ncbi.nlm.nih.gov/entrez/query.fcgi?db=gene&cmd=search&term=COPZ2) | [51226](http://www.ncbi.nlm.nih.gov/entrez/query.fcgi?db=gene&cmd=Retrieve&dopt=Graphics&list_uids=51226) | s27712 | UGACCAUCGUCUACAAGAAtt | UUCUUGUAGACGAUGGUCAta |
| NM_016429 | [COPZ2](http://www.ncbi.nlm.nih.gov/entrez/query.fcgi?db=gene&cmd=search&term=COPZ2) | [51226](http://www.ncbi.nlm.nih.gov/entrez/query.fcgi?db=gene&cmd=Retrieve&dopt=Graphics&list_uids=51226) | s27714 | CCUUCCCUCUACACCAUCAtt | UGAUGGUGUAGAGGGAAGGtt |
| NM_006364 | [SEC23A](http://www.ncbi.nlm.nih.gov/entrez/query.fcgi?db=gene&cmd=search&term=SEC23A) | [10484](http://www.ncbi.nlm.nih.gov/entrez/query.fcgi?db=gene&cmd=Retrieve&dopt=Graphics&list_uids=10484) | s20538 | GGGUGAUUCUUUCAAUACUtt | AGUAUUGAAAGAAUCACCCat |
| NM_006364 | [SEC23A](http://www.ncbi.nlm.nih.gov/entrez/query.fcgi?db=gene&cmd=search&term=SEC23A) | [10484](http://www.ncbi.nlm.nih.gov/entrez/query.fcgi?db=gene&cmd=Retrieve&dopt=Graphics&list_uids=10484) | s20539 | GGAUAUGCCUGAGUAUGAAtt | UUCAUACUCAGGCAUAUCCtg |
| NM_006364 | [SEC23A](http://www.ncbi.nlm.nih.gov/entrez/query.fcgi?db=gene&cmd=search&term=SEC23A) | [10484](http://www.ncbi.nlm.nih.gov/entrez/query.fcgi?db=gene&cmd=Retrieve&dopt=Graphics&list_uids=10484) | s20540 | GCAUUCUUGCAGAUCGUAUtt | AUACGAUCUGCAAGAAUGCta |
| NM_006323 | [SEC24B](http://www.ncbi.nlm.nih.gov/entrez/query.fcgi?db=gene&cmd=search&term=SEC24B) | [10427](http://www.ncbi.nlm.nih.gov/entrez/query.fcgi?db=gene&cmd=Retrieve&dopt=Graphics&list_uids=10427) | s20398 | GGUUCCAUCUGGAUAUGGAtt | UCCAUAUCCAGAUGGAACCtg |
| NM_006323 | [SEC24B](http://www.ncbi.nlm.nih.gov/entrez/query.fcgi?db=gene&cmd=search&term=SEC24B) | [10427](http://www.ncbi.nlm.nih.gov/entrez/query.fcgi?db=gene&cmd=Retrieve&dopt=Graphics&list_uids=10427) | s20396 | GCCCUAUUAUAUACAUCAAtt | UUGAUGUAUAUAAUAGGGCtg |
| NM_006323 | [SEC24B](http://www.ncbi.nlm.nih.gov/entrez/query.fcgi?db=gene&cmd=search&term=SEC24B) | [10427](http://www.ncbi.nlm.nih.gov/entrez/query.fcgi?db=gene&cmd=Retrieve&dopt=Graphics&list_uids=10427) | s20397 | GCAAUUACCAGUGAUAACAtt | UGUUAUCACUGGUAAUUGCgt |
| NM_001077206 | [SEC31A](http://www.ncbi.nlm.nih.gov/entrez/query.fcgi?db=gene&cmd=search&term=SEC31A) | [22872](http://www.ncbi.nlm.nih.gov/entrez/query.fcgi?db=gene&cmd=Retrieve&dopt=Graphics&list_uids=22872) | s22551 | GGAGCAUUGAAACUCGAAAtt | UUUCGAGUUUCAAUGCUCCtt |
| NM_001077206 | [SEC31A](http://www.ncbi.nlm.nih.gov/entrez/query.fcgi?db=gene&cmd=search&term=SEC31A) | [22872](http://www.ncbi.nlm.nih.gov/entrez/query.fcgi?db=gene&cmd=Retrieve&dopt=Graphics&list_uids=22872) | s22550 | CAAUCACCAGUGGUUUACAtt | UGUAAACCACUGGUGAUUGtt |
| NM_001077206 | [SEC31A](http://www.ncbi.nlm.nih.gov/entrez/query.fcgi?db=gene&cmd=search&term=SEC31A) | [22872](http://www.ncbi.nlm.nih.gov/entrez/query.fcgi?db=gene&cmd=Retrieve&dopt=Graphics&list_uids=22872) | s22552 | GAUCCAUCCUUGGAUAUGAtt | UCAUAUCCAAGGAUGGAUCag |
| NM_020150 | [SAR1A](http://www.ncbi.nlm.nih.gov/entrez/query.fcgi?db=gene&cmd=search&term=SAR1A) | [56681](http://www.ncbi.nlm.nih.gov/entrez/query.fcgi?db=gene&cmd=Retrieve&dopt=Graphics&list_uids=56681) | s32254 | GGAAUGACCUUUACAACUUtt | AAGUUGUAAAGGUCAUUCCag |
| NM_020150 | [SAR1A](http://www.ncbi.nlm.nih.gov/entrez/query.fcgi?db=gene&cmd=search&term=SAR1A) | [56681](http://www.ncbi.nlm.nih.gov/entrez/query.fcgi?db=gene&cmd=Retrieve&dopt=Graphics&list_uids=56681) | s32253 | GAUGCAAUCAGUGAAGAAAtt | UUUCUUCACUGAUUGCAUCtg |
| NM_020150 | [SAR1A](http://www.ncbi.nlm.nih.gov/entrez/query.fcgi?db=gene&cmd=search&term=SAR1A) | [56681](http://www.ncbi.nlm.nih.gov/entrez/query.fcgi?db=gene&cmd=Retrieve&dopt=Graphics&list_uids=56681) | s32255 | AACUUGUAUUCUUAGGUUUtt | AAACCUAAGAAUACAAGUUtt |
| NM_001033503 | [SAR1B](http://www.ncbi.nlm.nih.gov/entrez/query.fcgi?db=gene&cmd=search&term=SAR1B) | [51128](http://www.ncbi.nlm.nih.gov/entrez/query.fcgi?db=gene&cmd=Retrieve&dopt=Graphics&list_uids=51128) | s224190 | CCAUUGCUAAUGUGCCUAUtt | AUAGGCACAUUAGCAAUGGtt |
| NM_001033503 | [SAR1B](http://www.ncbi.nlm.nih.gov/entrez/query.fcgi?db=gene&cmd=search&term=SAR1B) | [51128](http://www.ncbi.nlm.nih.gov/entrez/query.fcgi?db=gene&cmd=Retrieve&dopt=Graphics&list_uids=51128) | s27508 | CUGGUAAACUGGUAUUUCUtt | AGAAAUACCAGUUUACCAGtt |
| NM_001033503 | [SAR1B](http://www.ncbi.nlm.nih.gov/entrez/query.fcgi?db=gene&cmd=search&term=SAR1B) | [51128](http://www.ncbi.nlm.nih.gov/entrez/query.fcgi?db=gene&cmd=Retrieve&dopt=Graphics&list_uids=51128) | s27507 | GAGUAUAUCUCUGAAAGAAtt | UUCUUUCAGAGAUAUACUCcc |
| NM_013388 | [PREB](http://www.ncbi.nlm.nih.gov/entrez/query.fcgi?db=gene&cmd=search&term=PREB) | [10113](http://www.ncbi.nlm.nih.gov/entrez/query.fcgi?db=gene&cmd=Retrieve&dopt=Graphics&list_uids=10113) | s19679 | GGCUUAUUAUUGUGACCAUtt | AUGGUCACAAUAAUAAGCCcg |
| NM_013388 | [PREB](http://www.ncbi.nlm.nih.gov/entrez/query.fcgi?db=gene&cmd=search&term=PREB) | [10113](http://www.ncbi.nlm.nih.gov/entrez/query.fcgi?db=gene&cmd=Retrieve&dopt=Graphics&list_uids=10113) | s19680 | CAGUGCCUCUACUACGUGAtt | UCACGUAGUAGAGGCACUGga |
| NM_013388 | [PREB](http://www.ncbi.nlm.nih.gov/entrez/query.fcgi?db=gene&cmd=search&term=PREB) | [10113](http://www.ncbi.nlm.nih.gov/entrez/query.fcgi?db=gene&cmd=Retrieve&dopt=Graphics&list_uids=10113) | s19681 | CCAGCACACCUUACCGCUAtt | UAGCGGUAAGGUGUGCUGGaa |
| **Intracellular trafficking** |  |  |  |  |  |
| NM_005570 | [LMAN1](http://www.ncbi.nlm.nih.gov/entrez/query.fcgi?db=gene&cmd=search&term=LMAN1) | [3998](http://www.ncbi.nlm.nih.gov/entrez/query.fcgi?db=gene&cmd=Retrieve&dopt=Graphics&list_uids=3998) | s8218 | GAGCAAAGAUUACCUAUUAtt | UAAUAGGUAAUCUUUGCUCgg |
| NM_005570 | [LMAN1](http://www.ncbi.nlm.nih.gov/entrez/query.fcgi?db=gene&cmd=search&term=LMAN1) | [3998](http://www.ncbi.nlm.nih.gov/entrez/query.fcgi?db=gene&cmd=Retrieve&dopt=Graphics&list_uids=3998) | s8220 | GAGGCUCAGUGUGGACAAAtt | UUUGUCCACACUGAGCCUCtt |
| NM_005570 | [LMAN1](http://www.ncbi.nlm.nih.gov/entrez/query.fcgi?db=gene&cmd=search&term=LMAN1) | [3998](http://www.ncbi.nlm.nih.gov/entrez/query.fcgi?db=gene&cmd=Retrieve&dopt=Graphics&list_uids=3998) | s8219 | GCAGAUUACUCAACAAGAAtt | UUCUUGUUGAGUAAUCUGCcc |
| NM_181661 | [VPS13B](http://www.ncbi.nlm.nih.gov/entrez/query.fcgi?db=gene&cmd=search&term=VPS13B) | [157680](http://www.ncbi.nlm.nih.gov/entrez/query.fcgi?db=gene&cmd=Retrieve&dopt=Graphics&list_uids=157680) | s45977 | CUGCCUAUGUUUAUUCGUAtt | UACGAAUAAACAUAGGCAGtt |
| NM_181661 | [VPS13B](http://www.ncbi.nlm.nih.gov/entrez/query.fcgi?db=gene&cmd=search&term=VPS13B) | [157680](http://www.ncbi.nlm.nih.gov/entrez/query.fcgi?db=gene&cmd=Retrieve&dopt=Graphics&list_uids=157680) | s45978 | CAGGAUCCUUUAUUAUACAtt | UGUAUAAUAAAGGAUCCUGgt |
| NM_181661 | [VPS13B](http://www.ncbi.nlm.nih.gov/entrez/query.fcgi?db=gene&cmd=search&term=VPS13B) | [157680](http://www.ncbi.nlm.nih.gov/entrez/query.fcgi?db=gene&cmd=Retrieve&dopt=Graphics&list_uids=157680) | s45979 | GACUUUGAAGGAUCCUAUUtt | AAUAGGAUCCUUCAAAGUCtg |
| NM_018668 | [VPS33B](http://www.ncbi.nlm.nih.gov/entrez/query.fcgi?db=gene&cmd=search&term=VPS33B) | [26276](http://www.ncbi.nlm.nih.gov/entrez/query.fcgi?db=gene&cmd=Retrieve&dopt=Graphics&list_uids=26276) | s25354 | GGAUCAACACUGUAGCUCAtt | UGAGCUACAGUGUUGAUCCaa |
| NM_018668 | [VPS33B](http://www.ncbi.nlm.nih.gov/entrez/query.fcgi?db=gene&cmd=search&term=VPS33B) | [26276](http://www.ncbi.nlm.nih.gov/entrez/query.fcgi?db=gene&cmd=Retrieve&dopt=Graphics&list_uids=26276) | s25353 | CGGCGAGUAUGAUCUGAAAtt | UUUCAGAUCAUACUCGCCGtc |
| NM_018668 | [VPS33B](http://www.ncbi.nlm.nih.gov/entrez/query.fcgi?db=gene&cmd=search&term=VPS33B) | [26276](http://www.ncbi.nlm.nih.gov/entrez/query.fcgi?db=gene&cmd=Retrieve&dopt=Graphics&list_uids=26276) | s25352 | GCCGUGGAGAGUAAAGUGAtt | UCACUUUACUCUCCACGGCtg |
| **Calcium-binding contact ER PM** |  |  |  |  |  |
| NM_020728 | [FAM62B](http://www.ncbi.nlm.nih.gov/entrez/query.fcgi?db=gene&cmd=search&term=FAM62B) | [57488](http://www.ncbi.nlm.nih.gov/entrez/query.fcgi?db=gene&cmd=Retrieve&dopt=Graphics&list_uids=57488) | s33137 | GCACCUUUAGUUUCACGAAtt | UUCGUGAAACUAAAGGUGCta |
| NM_020728 | [FAM62B](http://www.ncbi.nlm.nih.gov/entrez/query.fcgi?db=gene&cmd=search&term=FAM62B) | [57488](http://www.ncbi.nlm.nih.gov/entrez/query.fcgi?db=gene&cmd=Retrieve&dopt=Graphics&list_uids=57488) | s33136 | GGAUCAAUGGUGUUAAGGUtt | ACCUUAACACCAUUGAUCCtg |
| NM_020728 | [FAM62B](http://www.ncbi.nlm.nih.gov/entrez/query.fcgi?db=gene&cmd=search&term=FAM62B) | [57488](http://www.ncbi.nlm.nih.gov/entrez/query.fcgi?db=gene&cmd=Retrieve&dopt=Graphics&list_uids=57488) | s33138 | GAGAAGUUGUUUCGAGAAAtt | UUUCUCGAAACAACUUCUCta |
| NM_031913 | [FAM62C](http://www.ncbi.nlm.nih.gov/entrez/query.fcgi?db=gene&cmd=search&term=FAM62C) | [83850](http://www.ncbi.nlm.nih.gov/entrez/query.fcgi?db=gene&cmd=Retrieve&dopt=Graphics&list_uids=83850) | s38210 | CGAAUUCCUUGACAAUGAAtt | UUCAUUGUCAAGGAAUUCGaa |
| NM_031913 | [FAM62C](http://www.ncbi.nlm.nih.gov/entrez/query.fcgi?db=gene&cmd=search&term=FAM62C) | [83850](http://www.ncbi.nlm.nih.gov/entrez/query.fcgi?db=gene&cmd=Retrieve&dopt=Graphics&list_uids=83850) | s38211 | CCAACAGAGUGGUGGAUGAtt | UCAUCCACCACUCUGUUGGtc |
| NM_031913 | [FAM62C](http://www.ncbi.nlm.nih.gov/entrez/query.fcgi?db=gene&cmd=search&term=FAM62C) | [83850](http://www.ncbi.nlm.nih.gov/entrez/query.fcgi?db=gene&cmd=Retrieve&dopt=Graphics&list_uids=83850) | s38209 | CUGAGGACCUUUACCUUUAtt | UAAAGGUAAAGGUCCUCAGgt |
| **Sorting nexins** |  |  |  |  |  |
| NM_052948 | [SNX26](http://www.ncbi.nlm.nih.gov/entrez/query.fcgi?db=gene&cmd=search&term=SNX26) | [115703](http://www.ncbi.nlm.nih.gov/entrez/query.fcgi?db=gene&cmd=Retrieve&dopt=Graphics&list_uids=115703) | s41853 | CCCAUGUGAUCAAACGGUAtt | UACCGUUUGAUCACAUGGGcg |
| NM_052948 | [SNX26](http://www.ncbi.nlm.nih.gov/entrez/query.fcgi?db=gene&cmd=search&term=SNX26) | [115703](http://www.ncbi.nlm.nih.gov/entrez/query.fcgi?db=gene&cmd=Retrieve&dopt=Graphics&list_uids=115703) | s41851 | GGCACGAGUUUGACAGUGAtt | UCACUGUCAAACUCGUGCCga |
| NM_052948 | [SNX26](http://www.ncbi.nlm.nih.gov/entrez/query.fcgi?db=gene&cmd=search&term=SNX26) | [115703](http://www.ncbi.nlm.nih.gov/entrez/query.fcgi?db=gene&cmd=Retrieve&dopt=Graphics&list_uids=115703) | s41852 | CCGGAGUUACGAUGACUUUtt | AAAGUCAUCGUAACUCCGGag |
| **RAB** |  |  |  |  |  |
| NM_016131 | [RAB10](http://www.ncbi.nlm.nih.gov/entrez/query.fcgi?db=gene&cmd=search&term=RAB10) | [10890](http://www.ncbi.nlm.nih.gov/entrez/query.fcgi?db=gene&cmd=Retrieve&dopt=Graphics&list_uids=10890) | s21391 | GGAUGAUGCCUUCAAUACUtt | AGUAUUGAAGGCAUCAUCCga |
| NM_016131 | [RAB10](http://www.ncbi.nlm.nih.gov/entrez/query.fcgi?db=gene&cmd=search&term=RAB10) | [10890](http://www.ncbi.nlm.nih.gov/entrez/query.fcgi?db=gene&cmd=Retrieve&dopt=Graphics&list_uids=10890) | s21390 | GGACGACAAAAGAGUUGUAtt | UACAACUCUUUUGUCGUCCat |
| NM_016131 | [RAB10](http://www.ncbi.nlm.nih.gov/entrez/query.fcgi?db=gene&cmd=search&term=RAB10) | [10890](http://www.ncbi.nlm.nih.gov/entrez/query.fcgi?db=gene&cmd=Retrieve&dopt=Graphics&list_uids=10890) | s21392 | GGAAUAGACUUCAAGAUCAtt | UGAUCUUGAAGUCUAUUCCta |
| NM_004663 | [RAB11A](http://www.ncbi.nlm.nih.gov/entrez/query.fcgi?db=gene&cmd=search&term=RAB11A) | [8766](http://www.ncbi.nlm.nih.gov/entrez/query.fcgi?db=gene&cmd=Retrieve&dopt=Graphics&list_uids=8766) | s16704 | GGAGUAGAGUUUGCAACAAtt | UUGUUGCAAACUCUACUCCaa |
| NM_004663 | [RAB11A](http://www.ncbi.nlm.nih.gov/entrez/query.fcgi?db=gene&cmd=search&term=RAB11A) | [8766](http://www.ncbi.nlm.nih.gov/entrez/query.fcgi?db=gene&cmd=Retrieve&dopt=Graphics&list_uids=8766) | s16703 | GAGAUUUACCGCAUUGUUUtt | AAACAAUGCGGUAAAUCUCtg |
| NM_004663 | [RAB11A](http://www.ncbi.nlm.nih.gov/entrez/query.fcgi?db=gene&cmd=search&term=RAB11A) | [8766](http://www.ncbi.nlm.nih.gov/entrez/query.fcgi?db=gene&cmd=Retrieve&dopt=Graphics&list_uids=8766) | s16702 | CAACAAUGUGGUUCCUAUUtt | AAUAGGAACCACAUUGUUGct |
| NM_004218 | [RAB11B](http://www.ncbi.nlm.nih.gov/entrez/query.fcgi?db=gene&cmd=search&term=RAB11B) | [9230](http://www.ncbi.nlm.nih.gov/entrez/query.fcgi?db=gene&cmd=Retrieve&dopt=Graphics&list_uids=9230) | s17647 | CUAACGUAGAGGAAGCAUUtt | AAUGCUUCCUCUACGUUAGtg |
| NM_004218 | [RAB11B](http://www.ncbi.nlm.nih.gov/entrez/query.fcgi?db=gene&cmd=search&term=RAB11B) | [9230](http://www.ncbi.nlm.nih.gov/entrez/query.fcgi?db=gene&cmd=Retrieve&dopt=Graphics&list_uids=9230) | s17649 | GCAACGAGUUCAACCUGGAtt | UCCAGGUUGAACUCGUUGCgg |
| NM_004218 | [RAB11B](http://www.ncbi.nlm.nih.gov/entrez/query.fcgi?db=gene&cmd=search&term=RAB11B) | [9230](http://www.ncbi.nlm.nih.gov/entrez/query.fcgi?db=gene&cmd=Retrieve&dopt=Graphics&list_uids=9230) | s17648 | GCAUUCAAGAACAUCCUCAtt | UGAGGAUGUUCUUGAAUGCtt |
| NM_001025300 | [RAB12](http://www.ncbi.nlm.nih.gov/entrez/query.fcgi?db=gene&cmd=search&term=RAB12) | [201475](http://www.ncbi.nlm.nih.gov/entrez/query.fcgi?db=gene&cmd=Retrieve&dopt=Graphics&list_uids=201475) | s223483 | CGUGGGUGUUGACUUCAAAtt | UUUGAAGUCAACACCCACGgt |
| NM_001025300 | [RAB12](http://www.ncbi.nlm.nih.gov/entrez/query.fcgi?db=gene&cmd=search&term=RAB12) | [201475](http://www.ncbi.nlm.nih.gov/entrez/query.fcgi?db=gene&cmd=Retrieve&dopt=Graphics&list_uids=201475) | s47369 | GAUGCCUCUGGAUAUUUUAtt | UAAAAUAUCCAGAGGCAUCtt |
| NM_001025300 | [RAB12](http://www.ncbi.nlm.nih.gov/entrez/query.fcgi?db=gene&cmd=search&term=RAB12) | [201475](http://www.ncbi.nlm.nih.gov/entrez/query.fcgi?db=gene&cmd=Retrieve&dopt=Graphics&list_uids=201475) | s47370 | UGAUAAGUAUGCUUCAGAAtt | UUCUGAAGCAUACUUAUCAat |
| NM_002870 | [RAB13](http://www.ncbi.nlm.nih.gov/entrez/query.fcgi?db=gene&cmd=search&term=RAB13) | [5872](http://www.ncbi.nlm.nih.gov/entrez/query.fcgi?db=gene&cmd=Retrieve&dopt=Graphics&list_uids=5872) | s11691 | GGAAUCCGAUUUUUCGAAAtt | UUUCGAAAAAUCGGAUUCCat |
| NM_002870 | [RAB13](http://www.ncbi.nlm.nih.gov/entrez/query.fcgi?db=gene&cmd=search&term=RAB13) | [5872](http://www.ncbi.nlm.nih.gov/entrez/query.fcgi?db=gene&cmd=Retrieve&dopt=Graphics&list_uids=5872) | s11690 | GAGCGGUUCAAGACAAUAAtt | UUAUUGUCUUGAACCGCUCtt |
| NM_002870 | [RAB13](http://www.ncbi.nlm.nih.gov/entrez/query.fcgi?db=gene&cmd=search&term=RAB13) | [5872](http://www.ncbi.nlm.nih.gov/entrez/query.fcgi?db=gene&cmd=Retrieve&dopt=Graphics&list_uids=5872) | s11692 | GAUCCGCACUGUGGAUAUAtt | UAUAUCCACAGUGCGGAUCtt |
| NM_016322 | [RAB14](http://www.ncbi.nlm.nih.gov/entrez/query.fcgi?db=gene&cmd=search&term=RAB14) | [51552](http://www.ncbi.nlm.nih.gov/entrez/query.fcgi?db=gene&cmd=Retrieve&dopt=Graphics&list_uids=51552) | s28312 | GGUCUAUGAUAUCACUAGAtt | UCUAGUGAUAUCAUAGACCat |
| NM_016322 | [RAB14](http://www.ncbi.nlm.nih.gov/entrez/query.fcgi?db=gene&cmd=search&term=RAB14) | [51552](http://www.ncbi.nlm.nih.gov/entrez/query.fcgi?db=gene&cmd=Retrieve&dopt=Graphics&list_uids=51552) | s28311 | GAAAAUCUAUCAGAACAUUtt | AAUGUUCUGAUAGAUUUUCtt |
| NM_016322 | [RAB14](http://www.ncbi.nlm.nih.gov/entrez/query.fcgi?db=gene&cmd=search&term=RAB14) | [51552](http://www.ncbi.nlm.nih.gov/entrez/query.fcgi?db=gene&cmd=Retrieve&dopt=Graphics&list_uids=51552) | s28313 | AAGUACAUAUAACCACUUAtt | UAAGUGGUUAUAUGUACUUct |
| NM_198686 | [RAB15](http://www.ncbi.nlm.nih.gov/entrez/query.fcgi?db=gene&cmd=search&term=RAB15) | [376267](http://www.ncbi.nlm.nih.gov/entrez/query.fcgi?db=gene&cmd=Retrieve&dopt=Graphics&list_uids=376267) | s51762 | GAGAGAUACCAGACCAUCAtt | UGAUGGUCUGGUAUCUCUCct |
| NM_198686 | [RAB15](http://www.ncbi.nlm.nih.gov/entrez/query.fcgi?db=gene&cmd=search&term=RAB15) | [376267](http://www.ncbi.nlm.nih.gov/entrez/query.fcgi?db=gene&cmd=Retrieve&dopt=Graphics&list_uids=376267) | s51761 | CCAUCGGUGUUGACUUUAAtt | UUAAAGUCAACACCGAUGGtg |
| NM_198686 | [RAB15](http://www.ncbi.nlm.nih.gov/entrez/query.fcgi?db=gene&cmd=search&term=RAB15) | [376267](http://www.ncbi.nlm.nih.gov/entrez/query.fcgi?db=gene&cmd=Retrieve&dopt=Graphics&list_uids=376267) | s51763 | GCAUCAAAGUGCGGAUACAtt | UGUAUCCGCACUUUGAUGCcg |
| NM_022449 | [RAB17](http://www.ncbi.nlm.nih.gov/entrez/query.fcgi?db=gene&cmd=search&term=RAB17) | [64284](http://www.ncbi.nlm.nih.gov/entrez/query.fcgi?db=gene&cmd=Retrieve&dopt=Graphics&list_uids=64284) | s34610 | GCUGUGCGUUCUUCACAAAtt | UUUGUGAAGAACGCACAGCcc |
| NM_022449 | [RAB17](http://www.ncbi.nlm.nih.gov/entrez/query.fcgi?db=gene&cmd=search&term=RAB17) | [64284](http://www.ncbi.nlm.nih.gov/entrez/query.fcgi?db=gene&cmd=Retrieve&dopt=Graphics&list_uids=64284) | s34608 | GGAGGUGACCUUCCAGGAAtt | UUCCUGGAAGGUCACCUCCcg |
| NM_022449 | [RAB17](http://www.ncbi.nlm.nih.gov/entrez/query.fcgi?db=gene&cmd=search&term=RAB17) | [64284](http://www.ncbi.nlm.nih.gov/entrez/query.fcgi?db=gene&cmd=Retrieve&dopt=Graphics&list_uids=64284) | s34609 | GCGCUUCUGGUGUACGACAtt | UGUCGUACACCAGAAGCGCag |
| NM_021252 | [RAB18](http://www.ncbi.nlm.nih.gov/entrez/query.fcgi?db=gene&cmd=search&term=RAB18) | [22931](http://www.ncbi.nlm.nih.gov/entrez/query.fcgi?db=gene&cmd=Retrieve&dopt=Graphics&list_uids=22931) | s22704 | GGUUCACAGAUGAUACGUUtt | AACGUAUCAUCUGUGAACCtc |
| NM_021252 | [RAB18](http://www.ncbi.nlm.nih.gov/entrez/query.fcgi?db=gene&cmd=search&term=RAB18) | [22931](http://www.ncbi.nlm.nih.gov/entrez/query.fcgi?db=gene&cmd=Retrieve&dopt=Graphics&list_uids=22931) | s22705 | GGAAAAUCGUGAAGUCGAUtt | AUCGACUUCACGAUUUUCCtt |
| NM_021252 | [RAB18](http://www.ncbi.nlm.nih.gov/entrez/query.fcgi?db=gene&cmd=search&term=RAB18) | [22931](http://www.ncbi.nlm.nih.gov/entrez/query.fcgi?db=gene&cmd=Retrieve&dopt=Graphics&list_uids=22931) | s22703 | GGAUGGAAAUAAGGCUAAAtt | UUUAGCCUUAUUUCCAUCCac |
| NM_001008749 | [RAB19](http://www.ncbi.nlm.nih.gov/entrez/query.fcgi?db=gene&cmd=search&term=RAB19) | [401409](http://www.ncbi.nlm.nih.gov/entrez/query.fcgi?db=gene&cmd=Retrieve&dopt=Graphics&list_uids=401409) | s53589 | GGAUUCAUGAGAUAGAGAAtt | UUCUCUAUCUCAUGAAUCCag |
| NM_001008749 | [RAB19](http://www.ncbi.nlm.nih.gov/entrez/query.fcgi?db=gene&cmd=search&term=RAB19) | [401409](http://www.ncbi.nlm.nih.gov/entrez/query.fcgi?db=gene&cmd=Retrieve&dopt=Graphics&list_uids=401409) | s53588 | GGACCAAGAGUUCCCAUUAtt | UAAUGGGAACUCUUGGUCCag |
| NM_001008749 | [RAB19](http://www.ncbi.nlm.nih.gov/entrez/query.fcgi?db=gene&cmd=search&term=RAB19) | [401409](http://www.ncbi.nlm.nih.gov/entrez/query.fcgi?db=gene&cmd=Retrieve&dopt=Graphics&list_uids=401409) | s53590 | GACUAUUUGUUCAAGAUUAtt | UAAUCUUGAACAAAUAGUCaa |
| NM_004161 | [RAB1A](http://www.ncbi.nlm.nih.gov/entrez/query.fcgi?db=gene&cmd=search&term=RAB1A) | [5861](http://www.ncbi.nlm.nih.gov/entrez/query.fcgi?db=gene&cmd=Retrieve&dopt=Graphics&list_uids=5861) | s11658 | GGGAACAAAUGUGAUCUGAtt | UCAGAUCACAUUUGUUCCCta |
| NM_004161 | [RAB1A](http://www.ncbi.nlm.nih.gov/entrez/query.fcgi?db=gene&cmd=search&term=RAB1A) | [5861](http://www.ncbi.nlm.nih.gov/entrez/query.fcgi?db=gene&cmd=Retrieve&dopt=Graphics&list_uids=5861) | s229381 | CAAAGAAAGUAGUAGACUAtt | UAGUCUACUACUUUCUUUGtg |
| NM_015543 | [RAB1A](http://www.ncbi.nlm.nih.gov/entrez/query.fcgi?db=gene&cmd=search&term=RAB1A) | [5861](http://www.ncbi.nlm.nih.gov/entrez/query.fcgi?db=gene&cmd=Retrieve&dopt=Graphics&list_uids=5861) | s229382 | GAGUUAGACGGGAAAACAAtt | UUGUUUUCCCGUCUAACUCta |
| NM_030981 | [RAB1B](http://www.ncbi.nlm.nih.gov/entrez/query.fcgi?db=gene&cmd=search&term=RAB1B) | [81876](http://www.ncbi.nlm.nih.gov/entrez/query.fcgi?db=gene&cmd=Retrieve&dopt=Graphics&list_uids=81876) | s117 | GCUGAAAUCAAAAAGCGGAtt | UCCGCUUUUUGAUUUCAGCag |
| NM_030981 | [RAB1B](http://www.ncbi.nlm.nih.gov/entrez/query.fcgi?db=gene&cmd=search&term=RAB1B) | [81876](http://www.ncbi.nlm.nih.gov/entrez/query.fcgi?db=gene&cmd=Retrieve&dopt=Graphics&list_uids=81876) | s119 | GCACCAGCCUUAACCCUCAtt | UGAGGGUUAAGGCUGGUGCcc |
| NM_030981 | [RAB1B](http://www.ncbi.nlm.nih.gov/entrez/query.fcgi?db=gene&cmd=search&term=RAB1B) | [81876](http://www.ncbi.nlm.nih.gov/entrez/query.fcgi?db=gene&cmd=Retrieve&dopt=Graphics&list_uids=81876) | s118 | AGAGCGACCUCACCACCAAtt | UUGGUGGUGAGGUCGCUCUtg |
| NM_017817 | [RAB20](http://www.ncbi.nlm.nih.gov/entrez/query.fcgi?db=gene&cmd=search&term=RAB20) | [55647](http://www.ncbi.nlm.nih.gov/entrez/query.fcgi?db=gene&cmd=Retrieve&dopt=Graphics&list_uids=55647) | s31158 | CAGUGGAUAUAUCCAGUCAtt | UGACUGGAUAUAUCCACUGtg |
| NM_017817 | [RAB20](http://www.ncbi.nlm.nih.gov/entrez/query.fcgi?db=gene&cmd=search&term=RAB20) | [55647](http://www.ncbi.nlm.nih.gov/entrez/query.fcgi?db=gene&cmd=Retrieve&dopt=Graphics&list_uids=55647) | s31160 | AGAUCCUCAAGUACAAGAUtt | AUCUUGUACUUGAGGAUCUtt |
| NM_017817 | [RAB20](http://www.ncbi.nlm.nih.gov/entrez/query.fcgi?db=gene&cmd=search&term=RAB20) | [55647](http://www.ncbi.nlm.nih.gov/entrez/query.fcgi?db=gene&cmd=Retrieve&dopt=Graphics&list_uids=55647) | s31159 | CUCUCGACGUGUUCAUUUAtt | UAAAUGAACACGUCGAGAGtc |
| NM_014999 | [RAB21](http://www.ncbi.nlm.nih.gov/entrez/query.fcgi?db=gene&cmd=search&term=RAB21) | [23011](http://www.ncbi.nlm.nih.gov/entrez/query.fcgi?db=gene&cmd=Retrieve&dopt=Graphics&list_uids=23011) | s22824 | GGGUCAAAGAAUUACGGAAtt | UUCCGUAAUUCUUUGACCCag |
| NM_014999 | [RAB21](http://www.ncbi.nlm.nih.gov/entrez/query.fcgi?db=gene&cmd=search&term=RAB21) | [23011](http://www.ncbi.nlm.nih.gov/entrez/query.fcgi?db=gene&cmd=Retrieve&dopt=Graphics&list_uids=23011) | s22825 | GAGCGAUUUUAGUUUAUGAtt | UCAUAAACUAAAAUCGCUCca |
| NM_014999 | [RAB21](http://www.ncbi.nlm.nih.gov/entrez/query.fcgi?db=gene&cmd=search&term=RAB21) | [23011](http://www.ncbi.nlm.nih.gov/entrez/query.fcgi?db=gene&cmd=Retrieve&dopt=Graphics&list_uids=23011) | s22823 | GGGUCCAAUUUACUACAGAtt | UCUGUAGUAAAUUGGACCCaa |
| NM_020673 | [RAB22A](http://www.ncbi.nlm.nih.gov/entrez/query.fcgi?db=gene&cmd=search&term=RAB22A) | [57403](http://www.ncbi.nlm.nih.gov/entrez/query.fcgi?db=gene&cmd=Retrieve&dopt=Graphics&list_uids=57403) | s32994 | CGCCGACUCUAUUCAUGCAtt | UGCAUGAAUAGAGUCGGCGta |
| NM_020673 | [RAB22A](http://www.ncbi.nlm.nih.gov/entrez/query.fcgi?db=gene&cmd=search&term=RAB22A) | [57403](http://www.ncbi.nlm.nih.gov/entrez/query.fcgi?db=gene&cmd=Retrieve&dopt=Graphics&list_uids=57403) | s32993 | CAGCUAUAAUCGUUUAUGAtt | UCAUAAACGAUUAUAGCUGca |
| NM_020673 | [RAB22A](http://www.ncbi.nlm.nih.gov/entrez/query.fcgi?db=gene&cmd=search&term=RAB22A) | [57403](http://www.ncbi.nlm.nih.gov/entrez/query.fcgi?db=gene&cmd=Retrieve&dopt=Graphics&list_uids=57403) | s32992 | UGAGCUACAUAAAUUCCUAtt | UAGGAAUUUAUGUAGCUCAtt |
| NM_016277 | [RAB23](http://www.ncbi.nlm.nih.gov/entrez/query.fcgi?db=gene&cmd=search&term=RAB23) | [51715](http://www.ncbi.nlm.nih.gov/entrez/query.fcgi?db=gene&cmd=Retrieve&dopt=Graphics&list_uids=51715) | s28568 | CAAUCUUAGACCCAACAAAtt | UUUGUUGGGUCUAAGAUUGat |
| NM_016277 | [RAB23](http://www.ncbi.nlm.nih.gov/entrez/query.fcgi?db=gene&cmd=search&term=RAB23) | [51715](http://www.ncbi.nlm.nih.gov/entrez/query.fcgi?db=gene&cmd=Retrieve&dopt=Graphics&list_uids=51715) | s28569 | GAUUCAGCGAUAUUGCAAAtt | UUUGCAAUAUCGCUGAAUCat |
| NM_016277 | [RAB23](http://www.ncbi.nlm.nih.gov/entrez/query.fcgi?db=gene&cmd=search&term=RAB23) | [51715](http://www.ncbi.nlm.nih.gov/entrez/query.fcgi?db=gene&cmd=Retrieve&dopt=Graphics&list_uids=51715) | s28567 | CAGUGAAAGAAGAUCUAAAtt | UUUAGAUCUUCUUUCACUGat |
| NM_130781 | [RAB24](http://www.ncbi.nlm.nih.gov/entrez/query.fcgi?db=gene&cmd=search&term=RAB24) | [53917](http://www.ncbi.nlm.nih.gov/entrez/query.fcgi?db=gene&cmd=Retrieve&dopt=Graphics&list_uids=53917) | s224289 | AGGUGAUGUCGGUCGGAGAtt | UCUCCGACCGACAUCACCUtg |
| NM_130781 | [RAB24](http://www.ncbi.nlm.nih.gov/entrez/query.fcgi?db=gene&cmd=search&term=RAB24) | [53917](http://www.ncbi.nlm.nih.gov/entrez/query.fcgi?db=gene&cmd=Retrieve&dopt=Graphics&list_uids=53917) | s28803 | CGAGCAAAGUUCUGGGUGAtt | UCACCCAGAACUUUGCUCGct |
| NM_130781 | [RAB24](http://www.ncbi.nlm.nih.gov/entrez/query.fcgi?db=gene&cmd=search&term=RAB24) | [53917](http://www.ncbi.nlm.nih.gov/entrez/query.fcgi?db=gene&cmd=Retrieve&dopt=Graphics&list_uids=53917) | s28805 | GCUCAGCUCUUUGAAACAUtt | AUGUUUCAAAGAGCUGAGCtt |
| NM_020387 | [RAB25](http://www.ncbi.nlm.nih.gov/entrez/query.fcgi?db=gene&cmd=search&term=RAB25) | [57111](http://www.ncbi.nlm.nih.gov/entrez/query.fcgi?db=gene&cmd=Retrieve&dopt=Graphics&list_uids=57111) | s32701 | CAUGCUCGUGGGUAACAAAtt | UUUGUUACCCACGAGCAUGac |
| NM_020387 | [RAB25](http://www.ncbi.nlm.nih.gov/entrez/query.fcgi?db=gene&cmd=search&term=RAB25) | [57111](http://www.ncbi.nlm.nih.gov/entrez/query.fcgi?db=gene&cmd=Retrieve&dopt=Graphics&list_uids=57111) | s32702 | GAAUGUUCGCUGAAAACAAtt | UUGUUUUCAGCGAACAUUCgg |
| NM_020387 | [RAB25](http://www.ncbi.nlm.nih.gov/entrez/query.fcgi?db=gene&cmd=search&term=RAB25) | [57111](http://www.ncbi.nlm.nih.gov/entrez/query.fcgi?db=gene&cmd=Retrieve&dopt=Graphics&list_uids=57111) | s224452 | GACCUAACCAAGCACCAGAtt | UCUGGUGCUUGGUUAGGUCaa |
| NM_014353 | [RAB26](http://www.ncbi.nlm.nih.gov/entrez/query.fcgi?db=gene&cmd=search&term=RAB26) | [25837](http://www.ncbi.nlm.nih.gov/entrez/query.fcgi?db=gene&cmd=Retrieve&dopt=Graphics&list_uids=25837) | s223662 | CGCACAGUGCCUAACCCUAtt | UAGGGUUAGGCACUGUGCGtg |
| NM_014353 | [RAB26](http://www.ncbi.nlm.nih.gov/entrez/query.fcgi?db=gene&cmd=search&term=RAB26) | [25837](http://www.ncbi.nlm.nih.gov/entrez/query.fcgi?db=gene&cmd=Retrieve&dopt=Graphics&list_uids=25837) | s24591 | CGGCUGCAUGAUUACGUUAtt | UAACGUAAUCAUGCAGCCGga |
| NM_014353 | [RAB26](http://www.ncbi.nlm.nih.gov/entrez/query.fcgi?db=gene&cmd=search&term=RAB26) | [25837](http://www.ncbi.nlm.nih.gov/entrez/query.fcgi?db=gene&cmd=Retrieve&dopt=Graphics&list_uids=25837) | s24590 | GUUCCGCAGUGUUACCCAUtt | AUGGGUAACACUGCGGAACcg |
| NM_004580 | [RAB27A](http://www.ncbi.nlm.nih.gov/entrez/query.fcgi?db=gene&cmd=search&term=RAB27A) | [5873](http://www.ncbi.nlm.nih.gov/entrez/query.fcgi?db=gene&cmd=Retrieve&dopt=Graphics&list_uids=5873) | s11694 | GGAGAGGUUUCGUAGCUUAtt | UAAGCUACGAAACCUCUCCtg |
| NM_004580 | [RAB27A](http://www.ncbi.nlm.nih.gov/entrez/query.fcgi?db=gene&cmd=search&term=RAB27A) | [5873](http://www.ncbi.nlm.nih.gov/entrez/query.fcgi?db=gene&cmd=Retrieve&dopt=Graphics&list_uids=5873) | s11693 | GCCUCUACGGAUCAGUUAAtt | UUAACUGAUCCGUAGAGGCat |
| NM_004580 | [RAB27A](http://www.ncbi.nlm.nih.gov/entrez/query.fcgi?db=gene&cmd=search&term=RAB27A) | [5873](http://www.ncbi.nlm.nih.gov/entrez/query.fcgi?db=gene&cmd=Retrieve&dopt=Graphics&list_uids=5873) | s11695 | GGAAGACCAGUGUACUUUAtt | UAAAGUACACUGGUCUUCCct |
| NM_004163 | [RAB27B](http://www.ncbi.nlm.nih.gov/entrez/query.fcgi?db=gene&cmd=search&term=RAB27B) | [5874](http://www.ncbi.nlm.nih.gov/entrez/query.fcgi?db=gene&cmd=Retrieve&dopt=Graphics&list_uids=5874) | s11696 | GACUUAAUCAUGAAGCGAAtt | UUCGCUUCAUGAUUAAGUCca |
| NM_004163 | [RAB27B](http://www.ncbi.nlm.nih.gov/entrez/query.fcgi?db=gene&cmd=search&term=RAB27B) | [5874](http://www.ncbi.nlm.nih.gov/entrez/query.fcgi?db=gene&cmd=Retrieve&dopt=Graphics&list_uids=5874) | s11697 | GGAAUAGACUUUCGGGAAAtt | UUUCCCGAAAGUCUAUUCCta |
| NM_004163 | [RAB27B](http://www.ncbi.nlm.nih.gov/entrez/query.fcgi?db=gene&cmd=search&term=RAB27B) | [5874](http://www.ncbi.nlm.nih.gov/entrez/query.fcgi?db=gene&cmd=Retrieve&dopt=Graphics&list_uids=5874) | s224511 | UGCUUAUUGUGAAAAUCCAtt | UGGAUUUUCACAAUAAGCAtt |
| NM_004249 | [RAB28](http://www.ncbi.nlm.nih.gov/entrez/query.fcgi?db=gene&cmd=search&term=RAB28) | [9364](http://www.ncbi.nlm.nih.gov/entrez/query.fcgi?db=gene&cmd=Retrieve&dopt=Graphics&list_uids=9364) | s17910 | GGAGCAUAUGCGAACAAUAtt | UAUUGUUCGCAUAUGCUCCaa |
| NM_004249 | [RAB28](http://www.ncbi.nlm.nih.gov/entrez/query.fcgi?db=gene&cmd=search&term=RAB28) | [9364](http://www.ncbi.nlm.nih.gov/entrez/query.fcgi?db=gene&cmd=Retrieve&dopt=Graphics&list_uids=9364) | s17911 | CUAUAGGACUGGAUUUCUUtt | AAGAAAUCCAGUCCUAUAGtt |
| NM_004249 | [RAB28](http://www.ncbi.nlm.nih.gov/entrez/query.fcgi?db=gene&cmd=search&term=RAB28) | [9364](http://www.ncbi.nlm.nih.gov/entrez/query.fcgi?db=gene&cmd=Retrieve&dopt=Graphics&list_uids=9364) | s17912 | AAACUUUUGGGAAACAGUAtt | UACUGUUUCCCAAAAGUUUct |
| NM_002865 | [RAB2A](http://www.ncbi.nlm.nih.gov/entrez/query.fcgi?db=gene&cmd=search&term=RAB2A) | [5862](http://www.ncbi.nlm.nih.gov/entrez/query.fcgi?db=gene&cmd=Retrieve&dopt=Graphics&list_uids=5862) | s11660 | GAAGGAGUCUUUGACAUUAtt | UAAUGUCAAAGACUCCUUCtt |
| NM_002865 | [RAB2A](http://www.ncbi.nlm.nih.gov/entrez/query.fcgi?db=gene&cmd=search&term=RAB2A) | [5862](http://www.ncbi.nlm.nih.gov/entrez/query.fcgi?db=gene&cmd=Retrieve&dopt=Graphics&list_uids=5862) | s11661 | GAGCUUUACUAGUUUACGAtt | UCGUAAACUAGUAAAGCUCct |
| NM_002865 | [RAB2A](http://www.ncbi.nlm.nih.gov/entrez/query.fcgi?db=gene&cmd=search&term=RAB2A) | [5862](http://www.ncbi.nlm.nih.gov/entrez/query.fcgi?db=gene&cmd=Retrieve&dopt=Graphics&list_uids=5862) | s11662 | GCAGGAGCUUUACUAGUUUtt | AAACUAGUAAAGCUCCUGCtg |
| NM_032846 | [RAB2B](http://www.ncbi.nlm.nih.gov/entrez/query.fcgi?db=gene&cmd=search&term=RAB2B) | [84932](http://www.ncbi.nlm.nih.gov/entrez/query.fcgi?db=gene&cmd=Retrieve&dopt=Graphics&list_uids=84932) | s39689 | GAAUCCUUCCGUUCUAUCAtt | UGAUAGAACGGAAGGAUUCtt |
| NM_032846 | [RAB2B](http://www.ncbi.nlm.nih.gov/entrez/query.fcgi?db=gene&cmd=search&term=RAB2B) | [84932](http://www.ncbi.nlm.nih.gov/entrez/query.fcgi?db=gene&cmd=Retrieve&dopt=Graphics&list_uids=84932) | s39690 | CGACAUUACAAGGCGUGAAtt | UUCACGCCUUGUAAUGUCGta |
| NM_032846 | [RAB2B](http://www.ncbi.nlm.nih.gov/entrez/query.fcgi?db=gene&cmd=search&term=RAB2B) | [84932](http://www.ncbi.nlm.nih.gov/entrez/query.fcgi?db=gene&cmd=Retrieve&dopt=Graphics&list_uids=84932) | s39688 | UCAUGCUCAUUGGGAAUAAtt | UUAUUCCCAAUGAGCAUGAta |
| NM_014488 | [RAB30](http://www.ncbi.nlm.nih.gov/entrez/query.fcgi?db=gene&cmd=search&term=RAB30) | [27314](http://www.ncbi.nlm.nih.gov/entrez/query.fcgi?db=gene&cmd=Retrieve&dopt=Graphics&list_uids=27314) | s26138 | GAGAGAUUUCGGUCCAUUAtt | UAAUGGACCGAAAUCUCUCtt |
| NM_014488 | [RAB30](http://www.ncbi.nlm.nih.gov/entrez/query.fcgi?db=gene&cmd=search&term=RAB30) | [27314](http://www.ncbi.nlm.nih.gov/entrez/query.fcgi?db=gene&cmd=Retrieve&dopt=Graphics&list_uids=27314) | s26137 | CAAUGUAUCCUCACCCUUAtt | UAAGGGUGAGGAUACAUUGtt |
| NM_014488 | [RAB30](http://www.ncbi.nlm.nih.gov/entrez/query.fcgi?db=gene&cmd=search&term=RAB30) | [27314](http://www.ncbi.nlm.nih.gov/entrez/query.fcgi?db=gene&cmd=Retrieve&dopt=Graphics&list_uids=27314) | s26136 | GGAGUUGAUUUUAUGAUUAtt | UAAUCAUAAAAUCAACUCCaa |
| NM_006868 | [RAB31](http://www.ncbi.nlm.nih.gov/entrez/query.fcgi?db=gene&cmd=search&term=RAB31) | [11031](http://www.ncbi.nlm.nih.gov/entrez/query.fcgi?db=gene&cmd=Retrieve&dopt=Graphics&list_uids=11031) | s21731 | GGAAUACGCUGAAUCCAUAtt | UAUGGAUUCAGCGUAUUCCtt |
| NM_006868 | [RAB31](http://www.ncbi.nlm.nih.gov/entrez/query.fcgi?db=gene&cmd=search&term=RAB31) | [11031](http://www.ncbi.nlm.nih.gov/entrez/query.fcgi?db=gene&cmd=Retrieve&dopt=Graphics&list_uids=11031) | s21732 | CAAUGGAACAAUCAAAGUUtt | AACUUUGAUUGUUCCAUUGtt |
| NM_006868 | [RAB31](http://www.ncbi.nlm.nih.gov/entrez/query.fcgi?db=gene&cmd=search&term=RAB31) | [11031](http://www.ncbi.nlm.nih.gov/entrez/query.fcgi?db=gene&cmd=Retrieve&dopt=Graphics&list_uids=11031) | s21733 | GAACUUCACAAGUUCCUCAtt | UGAGGAACUUGUGAAGUUCat |
| NM_006834 | [RAB32](http://www.ncbi.nlm.nih.gov/entrez/query.fcgi?db=gene&cmd=search&term=RAB32) | [10981](http://www.ncbi.nlm.nih.gov/entrez/query.fcgi?db=gene&cmd=Retrieve&dopt=Graphics&list_uids=10981) | s21620 | CCAAAGCUUUCCUAAUGAAtt | UUCAUUAGGAAAGCUUUGGtg |
| NM_006834 | [RAB32](http://www.ncbi.nlm.nih.gov/entrez/query.fcgi?db=gene&cmd=search&term=RAB32) | [10981](http://www.ncbi.nlm.nih.gov/entrez/query.fcgi?db=gene&cmd=Retrieve&dopt=Graphics&list_uids=10981) | s21619 | CCGAGUAUACUACAAGGAAtt | UUCCUUGUAGUAUACUCGGgt |
| NM_006834 | [RAB32](http://www.ncbi.nlm.nih.gov/entrez/query.fcgi?db=gene&cmd=search&term=RAB32) | [10981](http://www.ncbi.nlm.nih.gov/entrez/query.fcgi?db=gene&cmd=Retrieve&dopt=Graphics&list_uids=10981) | s223173 | GAGGCAGUCUUAAAAUGGAtt | UCCAUUUUAAGACUGCCUCaa |
| NM_004794 | [RAB33A](http://www.ncbi.nlm.nih.gov/entrez/query.fcgi?db=gene&cmd=search&term=RAB33A) | [9363](http://www.ncbi.nlm.nih.gov/entrez/query.fcgi?db=gene&cmd=Retrieve&dopt=Graphics&list_uids=9363) | s225118 | CCCUCCAACUUAGCCCUGAtt | UCAGGGCUAAGUUGGAGGGca |
| NM_004794 | [RAB33A](http://www.ncbi.nlm.nih.gov/entrez/query.fcgi?db=gene&cmd=search&term=RAB33A) | [9363](http://www.ncbi.nlm.nih.gov/entrez/query.fcgi?db=gene&cmd=Retrieve&dopt=Graphics&list_uids=9363) | s17908 | GAUUCGCAUCUUCAAAAUAtt | UAUUUUGAAGAUGCGAAUCtg |
| NM_004794 | [RAB33A](http://www.ncbi.nlm.nih.gov/entrez/query.fcgi?db=gene&cmd=search&term=RAB33A) | [9363](http://www.ncbi.nlm.nih.gov/entrez/query.fcgi?db=gene&cmd=Retrieve&dopt=Graphics&list_uids=9363) | s17907 | GCAUUACUACCGCAACGUAtt | UACGUUGCGGUAGUAAUGCtc |
| NM_031296 | [RAB33B](http://www.ncbi.nlm.nih.gov/entrez/query.fcgi?db=gene&cmd=search&term=RAB33B) | [83452](http://www.ncbi.nlm.nih.gov/entrez/query.fcgi?db=gene&cmd=Retrieve&dopt=Graphics&list_uids=83452) | s37937 | CCUACCAUCUUGGAUAGAAtt | UUCUAUCCAAGAUGGUAGGct |
| NM_031296 | [RAB33B](http://www.ncbi.nlm.nih.gov/entrez/query.fcgi?db=gene&cmd=search&term=RAB33B) | [83452](http://www.ncbi.nlm.nih.gov/entrez/query.fcgi?db=gene&cmd=Retrieve&dopt=Graphics&list_uids=83452) | s37939 | CACGGAUUCUUGUUGGAAAtt | UUUCCAACAAGAAUCCGUGgt |
| NM_031296 | [RAB33B](http://www.ncbi.nlm.nih.gov/entrez/query.fcgi?db=gene&cmd=search&term=RAB33B) | [83452](http://www.ncbi.nlm.nih.gov/entrez/query.fcgi?db=gene&cmd=Retrieve&dopt=Graphics&list_uids=83452) | s37938 | GAACGAUUCAGAAAGAGCAtt | UGCUCUUUCUGAAUCGUUCtt |
| NM_031934 | [RAB34](http://www.ncbi.nlm.nih.gov/entrez/query.fcgi?db=gene&cmd=search&term=RAB34) | [83871](http://www.ncbi.nlm.nih.gov/entrez/query.fcgi?db=gene&cmd=Retrieve&dopt=Graphics&list_uids=83871) | s38246 | CCAUUGGAGUGGACUUCGAtt | UCGAAGUCCACUCCAAUGGtg |
| NM_031934 | [RAB34](http://www.ncbi.nlm.nih.gov/entrez/query.fcgi?db=gene&cmd=search&term=RAB34) | [83871](http://www.ncbi.nlm.nih.gov/entrez/query.fcgi?db=gene&cmd=Retrieve&dopt=Graphics&list_uids=83871) | s38245 | ACACCUUUGAUAAGAAUUAtt | UAAUUCUUAUCAAAGGUGUct |
| NM_031934 | [RAB34](http://www.ncbi.nlm.nih.gov/entrez/query.fcgi?db=gene&cmd=search&term=RAB34) | [83871](http://www.ncbi.nlm.nih.gov/entrez/query.fcgi?db=gene&cmd=Retrieve&dopt=Graphics&list_uids=83871) | s38244 | GGAGAGGUUCAAAUGCAUUtt | AAUGCAUUUGAACCUCUCCtg |
| NM_006861 | [RAB35](http://www.ncbi.nlm.nih.gov/entrez/query.fcgi?db=gene&cmd=search&term=RAB35) | [11021](http://www.ncbi.nlm.nih.gov/entrez/query.fcgi?db=gene&cmd=Retrieve&dopt=Graphics&list_uids=11021) | s21709 | GGCAUCCAGUUGUUCGAGAtt | UCUCGAACAACUGGAUGCCca |
| NM_006861 | [RAB35](http://www.ncbi.nlm.nih.gov/entrez/query.fcgi?db=gene&cmd=search&term=RAB35) | [11021](http://www.ncbi.nlm.nih.gov/entrez/query.fcgi?db=gene&cmd=Retrieve&dopt=Graphics&list_uids=11021) | s21708 | GAAGAGAUGUUCAACUGCAtt | UGCAGUUGAACAUCUCUUCca |
| NM_006861 | [RAB35](http://www.ncbi.nlm.nih.gov/entrez/query.fcgi?db=gene&cmd=search&term=RAB35) | [11021](http://www.ncbi.nlm.nih.gov/entrez/query.fcgi?db=gene&cmd=Retrieve&dopt=Graphics&list_uids=11021) | s21707 | GCAGUUUACUGUUGCGUUUtt | AAACGCAACAGUAAACUGCtc |
| NM_004914 | [RAB36](http://www.ncbi.nlm.nih.gov/entrez/query.fcgi?db=gene&cmd=search&term=RAB36) | [9609](http://www.ncbi.nlm.nih.gov/entrez/query.fcgi?db=gene&cmd=Retrieve&dopt=Graphics&list_uids=9609) | s18461 | AGAAUGUUUUUGAUCGAGAtt | UCUCGAUCAAAAACAUUCUtg |
| NM_004914 | [RAB36](http://www.ncbi.nlm.nih.gov/entrez/query.fcgi?db=gene&cmd=search&term=RAB36) | [9609](http://www.ncbi.nlm.nih.gov/entrez/query.fcgi?db=gene&cmd=Retrieve&dopt=Graphics&list_uids=9609) | s18463 | AGAUUGCUGGGAUUCCCUAtt | UAGGGAAUCCCAGCAAUCUca |
| NM_004914 | [RAB36](http://www.ncbi.nlm.nih.gov/entrez/query.fcgi?db=gene&cmd=search&term=RAB36) | [9609](http://www.ncbi.nlm.nih.gov/entrez/query.fcgi?db=gene&cmd=Retrieve&dopt=Graphics&list_uids=9609) | s225166 | AGUGCAUCGCAUCUGCCUAtt | UAGGCAGAUGCGAUGCACUtg |
| NM_001006638 | [RAB37](http://www.ncbi.nlm.nih.gov/entrez/query.fcgi?db=gene&cmd=search&term=RAB37) | [326624](http://www.ncbi.nlm.nih.gov/entrez/query.fcgi?db=gene&cmd=Retrieve&dopt=Graphics&list_uids=326624) | s50263 | CCAGCUUCCAGAUCCGAGAtt | UCUCGGAUCUGGAAGCUGGgc |
| NM_001006638 | [RAB37](http://www.ncbi.nlm.nih.gov/entrez/query.fcgi?db=gene&cmd=search&term=RAB37) | [326624](http://www.ncbi.nlm.nih.gov/entrez/query.fcgi?db=gene&cmd=Retrieve&dopt=Graphics&list_uids=326624) | s223875 | GGAUAUGAGCAGCGAAAGAtt | UCUUUCGCUGCUCAUAUCCgc |
| NM_001006638 | [RAB37](http://www.ncbi.nlm.nih.gov/entrez/query.fcgi?db=gene&cmd=search&term=RAB37) | [326624](http://www.ncbi.nlm.nih.gov/entrez/query.fcgi?db=gene&cmd=Retrieve&dopt=Graphics&list_uids=326624) | s223876 | UGAUCAUGCUGCUAGGCAAtt | UUGCCUAGCAGCAUGAUCAcc |
| NM_022337 | [RAB38](http://www.ncbi.nlm.nih.gov/entrez/query.fcgi?db=gene&cmd=search&term=RAB38) | [23682](http://www.ncbi.nlm.nih.gov/entrez/query.fcgi?db=gene&cmd=Retrieve&dopt=Graphics&list_uids=23682) | s24320 | GCAAAUGAGUGUGACCUAAtt | UUAGGUCACACUCAUUUGCaa |
| NM_022337 | [RAB38](http://www.ncbi.nlm.nih.gov/entrez/query.fcgi?db=gene&cmd=search&term=RAB38) | [23682](http://www.ncbi.nlm.nih.gov/entrez/query.fcgi?db=gene&cmd=Retrieve&dopt=Graphics&list_uids=23682) | s24319 | CCAGAUGCCUGGUGAAACAtt | UGUUUCACCAGGCAUCUGGag |
| NM_022337 | [RAB38](http://www.ncbi.nlm.nih.gov/entrez/query.fcgi?db=gene&cmd=search&term=RAB38) | [23682](http://www.ncbi.nlm.nih.gov/entrez/query.fcgi?db=gene&cmd=Retrieve&dopt=Graphics&list_uids=23682) | s24321 | ACAUGACGAGGGUCUAUUAtt | UAAUAGACCCUCGUCAUGUtt |
| NM_017516 | [RAB39A](http://www.ncbi.nlm.nih.gov/entrez/query.fcgi?db=gene&cmd=search&term=RAB39A) | [54734](http://www.ncbi.nlm.nih.gov/entrez/query.fcgi?db=gene&cmd=Retrieve&dopt=Graphics&list_uids=54734) | s29360 | GGAGCGGUUCAGAUCAAUAtt | UAUUGAUCUGAACCGCUCCtg |
| NM_017516 | [RAB39A](http://www.ncbi.nlm.nih.gov/entrez/query.fcgi?db=gene&cmd=search&term=RAB39A) | [54734](http://www.ncbi.nlm.nih.gov/entrez/query.fcgi?db=gene&cmd=Retrieve&dopt=Graphics&list_uids=54734) | s29361 | GCAACUCAGUUGGUGGAUUtt | AAUCCACCAACUGAGUUGCgg |
| NM_017516 | [RAB39A](http://www.ncbi.nlm.nih.gov/entrez/query.fcgi?db=gene&cmd=search&term=RAB39A) | [54734](http://www.ncbi.nlm.nih.gov/entrez/query.fcgi?db=gene&cmd=Retrieve&dopt=Graphics&list_uids=54734) | s29359 | GACUGUGGAAUGAAGUAUAtt | UAUACUUCAUUCCACAGUCtg |
| NM_171998 | [RAB39B](http://www.ncbi.nlm.nih.gov/entrez/query.fcgi?db=gene&cmd=search&term=RAB39B) | [116442](http://www.ncbi.nlm.nih.gov/entrez/query.fcgi?db=gene&cmd=Retrieve&dopt=Graphics&list_uids=116442) | s42010 | AGACCAAAGUACACGUUCAtt | UGAACGUGUACUUUGGUCUct |
| NM_171998 | [RAB39B](http://www.ncbi.nlm.nih.gov/entrez/query.fcgi?db=gene&cmd=search&term=RAB39B) | [116442](http://www.ncbi.nlm.nih.gov/entrez/query.fcgi?db=gene&cmd=Retrieve&dopt=Graphics&list_uids=116442) | s42008 | ACCUGACAAGAGACAUAUAtt | UAUAUGUCUCUUGUCAGGUct |
| NM_171998 | [RAB39B](http://www.ncbi.nlm.nih.gov/entrez/query.fcgi?db=gene&cmd=search&term=RAB39B) | [116442](http://www.ncbi.nlm.nih.gov/entrez/query.fcgi?db=gene&cmd=Retrieve&dopt=Graphics&list_uids=116442) | s42009 | AGAGUGGAUUUGUACCAAAtt | UUUGGUACAAAUCCACUCUtc |
| NM_002867 | [RAB3B](http://www.ncbi.nlm.nih.gov/entrez/query.fcgi?db=gene&cmd=search&term=RAB3B) | [5865](http://www.ncbi.nlm.nih.gov/entrez/query.fcgi?db=gene&cmd=Retrieve&dopt=Graphics&list_uids=5865) | s11669 | GCUUCAUUCUGAUGUAUGAtt | UCAUACAUCAGAAUGAAGCcc |
| NM_002867 | [RAB3B](http://www.ncbi.nlm.nih.gov/entrez/query.fcgi?db=gene&cmd=search&term=RAB3B) | [5865](http://www.ncbi.nlm.nih.gov/entrez/query.fcgi?db=gene&cmd=Retrieve&dopt=Graphics&list_uids=5865) | s11671 | GCUAUGCUGAUGACACGUUtt | AACGUGUCAUCAGCAUAGCgg |
| NM_002867 | [RAB3B](http://www.ncbi.nlm.nih.gov/entrez/query.fcgi?db=gene&cmd=search&term=RAB3B) | [5865](http://www.ncbi.nlm.nih.gov/entrez/query.fcgi?db=gene&cmd=Retrieve&dopt=Graphics&list_uids=5865) | s11670 | CGACUUCAAGGUGAAGACAtt | UGUCUUCACCUUGAAGUCGat |
| NM_138453 | [RAB3C](http://www.ncbi.nlm.nih.gov/entrez/query.fcgi?db=gene&cmd=search&term=RAB3C) | [115827](http://www.ncbi.nlm.nih.gov/entrez/query.fcgi?db=gene&cmd=Retrieve&dopt=Graphics&list_uids=115827) | s41884 | GCACAGUUGGGAUCGAUUUtt | AAAUCGAUCCCAACUGUGCtg |
| NM_138453 | [RAB3C](http://www.ncbi.nlm.nih.gov/entrez/query.fcgi?db=gene&cmd=search&term=RAB3C) | [115827](http://www.ncbi.nlm.nih.gov/entrez/query.fcgi?db=gene&cmd=Retrieve&dopt=Graphics&list_uids=115827) | s41885 | CAAGAUUGGUCAACUCAAAtt | UUUGAGUUGACCAAUCUUGta |
| NM_138453 | [RAB3C](http://www.ncbi.nlm.nih.gov/entrez/query.fcgi?db=gene&cmd=search&term=RAB3C) | [115827](http://www.ncbi.nlm.nih.gov/entrez/query.fcgi?db=gene&cmd=Retrieve&dopt=Graphics&list_uids=115827) | s41886 | GAAAGAUACAGGACUAUCAtt | UGAUAGUCCUGUAUCUUUCct |
| NM_004283 | [RAB3D](http://www.ncbi.nlm.nih.gov/entrez/query.fcgi?db=gene&cmd=search&term=RAB3D) | [9545](http://www.ncbi.nlm.nih.gov/entrez/query.fcgi?db=gene&cmd=Retrieve&dopt=Graphics&list_uids=9545) | s18328 | GGGAUGCAGCAGAUCAGAAtt | UUCUGAUCUGCUGCAUCCCgt |
| NM_004283 | [RAB3D](http://www.ncbi.nlm.nih.gov/entrez/query.fcgi?db=gene&cmd=search&term=RAB3D) | [9545](http://www.ncbi.nlm.nih.gov/entrez/query.fcgi?db=gene&cmd=Retrieve&dopt=Graphics&list_uids=9545) | s18326 | AGGAGAACAUCAAUGUGAAtt | UUCACAUUGAUGUUCUCCUtg |
| NM_004283 | [RAB3D](http://www.ncbi.nlm.nih.gov/entrez/query.fcgi?db=gene&cmd=search&term=RAB3D) | [9545](http://www.ncbi.nlm.nih.gov/entrez/query.fcgi?db=gene&cmd=Retrieve&dopt=Graphics&list_uids=9545) | s18327 | AACUGCUACUGAUAGGCAAtt | UUGCCUAUCAGUAGCAGUUtg |
| NM_080879 | [RAB40A](http://www.ncbi.nlm.nih.gov/entrez/query.fcgi?db=gene&cmd=search&term=RAB40A) | [142684](http://www.ncbi.nlm.nih.gov/entrez/query.fcgi?db=gene&cmd=Retrieve&dopt=Graphics&list_uids=142684) | s44481 | AGAUGGAAGUCGCAUUGUAtt | UACAAUGCGACUUCCAUCUtc |
| NM_080879 | [RAB40A](http://www.ncbi.nlm.nih.gov/entrez/query.fcgi?db=gene&cmd=search&term=RAB40A) | [142684](http://www.ncbi.nlm.nih.gov/entrez/query.fcgi?db=gene&cmd=Retrieve&dopt=Graphics&list_uids=142684) | s223341 | AGGAUGAAUUGGCUCGGGAtt | UCCCGAGCCAAUUCAUCCUgt |
| NM_080879 | [RAB40A](http://www.ncbi.nlm.nih.gov/entrez/query.fcgi?db=gene&cmd=search&term=RAB40A) | [142684](http://www.ncbi.nlm.nih.gov/entrez/query.fcgi?db=gene&cmd=Retrieve&dopt=Graphics&list_uids=142684) | s223342 | GGAAUGUGUGUGAUGCCUUtt | AAGGCAUCACACACAUUCCca |
| NM_001031834 | [RAB40AL](http://www.ncbi.nlm.nih.gov/entrez/query.fcgi?db=gene&cmd=search&term=RAB40AL) | [282808](http://www.ncbi.nlm.nih.gov/entrez/query.fcgi?db=gene&cmd=Retrieve&dopt=Graphics&list_uids=282808) | s50182 | GGUAUGGAUCGAUGGAUUAtt | UAAUCCAUCGAUCCAUACCct |
| NM_001031834 | [RAB40AL](http://www.ncbi.nlm.nih.gov/entrez/query.fcgi?db=gene&cmd=search&term=RAB40AL) | [282808](http://www.ncbi.nlm.nih.gov/entrez/query.fcgi?db=gene&cmd=Retrieve&dopt=Graphics&list_uids=282808) | s50184 | UCAUAGAGUCUUUCACGGAtt | UCCGUGAAAGACUCUAUGAtg |
| NM_001031834 | [RAB40AL](http://www.ncbi.nlm.nih.gov/entrez/query.fcgi?db=gene&cmd=search&term=RAB40AL) | [282808](http://www.ncbi.nlm.nih.gov/entrez/query.fcgi?db=gene&cmd=Retrieve&dopt=Graphics&list_uids=282808) | s50183 | CUACUCUCGUGGUGCACAAtt | UUGUGCACCACGAGAGUAGga |
| NM_006822 | [RAB40B](http://www.ncbi.nlm.nih.gov/entrez/query.fcgi?db=gene&cmd=search&term=RAB40B) | [10966](http://www.ncbi.nlm.nih.gov/entrez/query.fcgi?db=gene&cmd=Retrieve&dopt=Graphics&list_uids=10966) | s223170 | GAAUGUUACUUUAUCAUGAtt | UCAUGAUAAAGUAACAUUCag |
| NM_006822 | [RAB40B](http://www.ncbi.nlm.nih.gov/entrez/query.fcgi?db=gene&cmd=search&term=RAB40B) | [10966](http://www.ncbi.nlm.nih.gov/entrez/query.fcgi?db=gene&cmd=Retrieve&dopt=Graphics&list_uids=10966) | s21590 | GGCAUUGAUCGAUGGAUUAtt | UAAUCCAUCGAUCAAUGCCgt |
| NM_006822 | [RAB40B](http://www.ncbi.nlm.nih.gov/entrez/query.fcgi?db=gene&cmd=search&term=RAB40B) | [10966](http://www.ncbi.nlm.nih.gov/entrez/query.fcgi?db=gene&cmd=Retrieve&dopt=Graphics&list_uids=10966) | s21589 | UCACAGAGUCGUUCACGGAtt | UCCGUGAACGACUCUGUGAtg |
| NM_021168 | [RAB40C](http://www.ncbi.nlm.nih.gov/entrez/query.fcgi?db=gene&cmd=search&term=RAB40C) | [57799](http://www.ncbi.nlm.nih.gov/entrez/query.fcgi?db=gene&cmd=Retrieve&dopt=Graphics&list_uids=57799) | s33731 | GCAUGAACGCGGUCAUGAUtt | AUCAUGACCGCGUUCAUGCcg |
| NM_021168 | [RAB40C](http://www.ncbi.nlm.nih.gov/entrez/query.fcgi?db=gene&cmd=search&term=RAB40C) | [57799](http://www.ncbi.nlm.nih.gov/entrez/query.fcgi?db=gene&cmd=Retrieve&dopt=Graphics&list_uids=57799) | s33730 | ACAGUAACGGGAUCGACUAtt | UAGUCGAUCCCGUUACUGUag |
| NM_021168 | [RAB40C](http://www.ncbi.nlm.nih.gov/entrez/query.fcgi?db=gene&cmd=search&term=RAB40C) | [57799](http://www.ncbi.nlm.nih.gov/entrez/query.fcgi?db=gene&cmd=Retrieve&dopt=Graphics&list_uids=57799) | s33729 | CACCAACCGCUGGUCCUUUtt | AAAGGACCAGCGGUUGGUGat |
| NM_001032726 | [RAB41](http://www.ncbi.nlm.nih.gov/entrez/query.fcgi?db=gene&cmd=search&term=RAB41) | [347517](http://www.ncbi.nlm.nih.gov/entrez/query.fcgi?db=gene&cmd=Retrieve&dopt=Graphics&list_uids=347517) | s51269 | GAAUUGACUUCUUGUCUAAtt | UUAGACAAGAAGUCAAUUCca |
| NM_001032726 | [RAB41](http://www.ncbi.nlm.nih.gov/entrez/query.fcgi?db=gene&cmd=search&term=RAB41) | [347517](http://www.ncbi.nlm.nih.gov/entrez/query.fcgi?db=gene&cmd=Retrieve&dopt=Graphics&list_uids=347517) | s51268 | CGGUUGAAAUCGAACUGGAtt | UCCAGUUCGAUUUCAACCGtc |
| NM_001032726 | [RAB41](http://www.ncbi.nlm.nih.gov/entrez/query.fcgi?db=gene&cmd=search&term=RAB41) | [347517](http://www.ncbi.nlm.nih.gov/entrez/query.fcgi?db=gene&cmd=Retrieve&dopt=Graphics&list_uids=347517) | s51270 | GUGGUUGUCUAUGACAUUAtt | UAAUGUCAUAGACAACCACtg |
| NM_152304 | [RAB42](http://www.ncbi.nlm.nih.gov/entrez/query.fcgi?db=gene&cmd=search&term=RAB42) | [115273](http://www.ncbi.nlm.nih.gov/entrez/query.fcgi?db=gene&cmd=Retrieve&dopt=Graphics&list_uids=115273) | s41783 | AGACCUCGGUUAAAAACAAtt | UUGUUUUUAACCGAGGUCUcc |
| NM_152304 | [RAB42](http://www.ncbi.nlm.nih.gov/entrez/query.fcgi?db=gene&cmd=search&term=RAB42) | [115273](http://www.ncbi.nlm.nih.gov/entrez/query.fcgi?db=gene&cmd=Retrieve&dopt=Graphics&list_uids=115273) | s41782 | CAGUUACUCCAGAGGCUAAtt | UUAGCCUCUGGAGUAACUGgg |
| NM_152304 | [RAB42](http://www.ncbi.nlm.nih.gov/entrez/query.fcgi?db=gene&cmd=search&term=RAB42) | [115273](http://www.ncbi.nlm.nih.gov/entrez/query.fcgi?db=gene&cmd=Retrieve&dopt=Graphics&list_uids=115273) | s41781 | UCGAGGCUUUCUACAGAAAtt | UUUCUGUAGAAAGCCUCGAct |
| NM_198490 | [RAB43](http://www.ncbi.nlm.nih.gov/entrez/query.fcgi?db=gene&cmd=search&term=RAB43) | [339122](http://www.ncbi.nlm.nih.gov/entrez/query.fcgi?db=gene&cmd=Retrieve&dopt=Graphics&list_uids=339122) | s50453 | GCUGAUCGGGAACAAGUCAtt | UGACUUGUUCCCGAUCAGCag |
| NM_198490 | [RAB43](http://www.ncbi.nlm.nih.gov/entrez/query.fcgi?db=gene&cmd=search&term=RAB43) | [339122](http://www.ncbi.nlm.nih.gov/entrez/query.fcgi?db=gene&cmd=Retrieve&dopt=Graphics&list_uids=339122) | s50452 | CCACAGACUUGGUGCCUGAtt | UCAGGCACCAAGUCUGUGGgt |
| NM_198490 | [RAB43](http://www.ncbi.nlm.nih.gov/entrez/query.fcgi?db=gene&cmd=search&term=RAB43) | [339122](http://www.ncbi.nlm.nih.gov/entrez/query.fcgi?db=gene&cmd=Retrieve&dopt=Graphics&list_uids=339122) | s50454 | GCCUUAUCAGGACCUCUGAtt | UCAGAGGUCCUGAUAAGGCag |
| NM_004578 | [RAB4A](http://www.ncbi.nlm.nih.gov/entrez/query.fcgi?db=gene&cmd=search&term=RAB4A) | [5867](http://www.ncbi.nlm.nih.gov/entrez/query.fcgi?db=gene&cmd=Retrieve&dopt=Graphics&list_uids=5867) | s11677 | CCUACAAUGCGCUUACUAAtt | UUAGUAAGCGCAUUGUAGGtt |
| NM_004578 | [RAB4A](http://www.ncbi.nlm.nih.gov/entrez/query.fcgi?db=gene&cmd=search&term=RAB4A) | [5867](http://www.ncbi.nlm.nih.gov/entrez/query.fcgi?db=gene&cmd=Retrieve&dopt=Graphics&list_uids=5867) | s11676 | GAACGAUUCAGGUCCGUGAtt | UCACGGACCUGAAUCGUUCtt |
| NM_004578 | [RAB4A](http://www.ncbi.nlm.nih.gov/entrez/query.fcgi?db=gene&cmd=search&term=RAB4A) | [5867](http://www.ncbi.nlm.nih.gov/entrez/query.fcgi?db=gene&cmd=Retrieve&dopt=Graphics&list_uids=5867) | s11675 | GGUCCGUGACGAGAAGUUAtt | UAACUUCUCGUCACGGACCtg |
| NM_016154 | [RAB4B](http://www.ncbi.nlm.nih.gov/entrez/query.fcgi?db=gene&cmd=search&term=RAB4B) | [53916](http://www.ncbi.nlm.nih.gov/entrez/query.fcgi?db=gene&cmd=Retrieve&dopt=Graphics&list_uids=53916) | s28800 | GGAAGACUGUGAAGCUACAtt | UGUAGCUUCACAGUCUUCCca |
| NM_016154 | [RAB4B](http://www.ncbi.nlm.nih.gov/entrez/query.fcgi?db=gene&cmd=search&term=RAB4B) | [53916](http://www.ncbi.nlm.nih.gov/entrez/query.fcgi?db=gene&cmd=Retrieve&dopt=Graphics&list_uids=53916) | s28802 | GCACUAUCCUCAACAAGAUtt | AUCUUGUUGAGGAUAGUGCgg |
| NM_016154 | [RAB4B](http://www.ncbi.nlm.nih.gov/entrez/query.fcgi?db=gene&cmd=search&term=RAB4B) | [53916](http://www.ncbi.nlm.nih.gov/entrez/query.fcgi?db=gene&cmd=Retrieve&dopt=Graphics&list_uids=53916) | s28801 | AGAAUAAGUUCAAACAGGAtt | UCCUGUUUGAACUUAUUCUca |
| NM_207498 | [RAB44](http://www.ncbi.nlm.nih.gov/entrez/query.fcgi?db=gene&cmd=search&term=) | [401258](http://www.ncbi.nlm.nih.gov/entrez/query.fcgi?db=gene&cmd=Retrieve&dopt=Graphics&list_uids=401258) | s53537 | GAAUUCAGCUCUGGACUCAtt | UGAGUCCAGAGCUGAAUUCtt |
| NM_207498 | [RAB44](http://www.ncbi.nlm.nih.gov/entrez/query.fcgi?db=gene&cmd=search&term=) | [401258](http://www.ncbi.nlm.nih.gov/entrez/query.fcgi?db=gene&cmd=Retrieve&dopt=Graphics&list_uids=401258) | s53536 | AUUCGGCUGUUGCUCCUGAtt | UCAGGAGCAACAGCCGAAUct |
| NM_207498 | [RAB44](http://www.ncbi.nlm.nih.gov/entrez/query.fcgi?db=gene&cmd=search&term=) | [401258](http://www.ncbi.nlm.nih.gov/entrez/query.fcgi?db=gene&cmd=Retrieve&dopt=Graphics&list_uids=401258) | s53538 | UCAGGACUCGGAUCCCAGAtt | UCUGGGAUCCGAGUCCUGAgt |
| NM_004162 | [RAB5A](http://www.ncbi.nlm.nih.gov/entrez/query.fcgi?db=gene&cmd=search&term=RAB5A) | [5868](http://www.ncbi.nlm.nih.gov/entrez/query.fcgi?db=gene&cmd=Retrieve&dopt=Graphics&list_uids=5868) | s11679 | CAAGCCUAGUGCUUCGUUUtt | AAACGAAGCACUAGGCUUGat |
| NM_004162 | [RAB5A](http://www.ncbi.nlm.nih.gov/entrez/query.fcgi?db=gene&cmd=search&term=RAB5A) | [5868](http://www.ncbi.nlm.nih.gov/entrez/query.fcgi?db=gene&cmd=Retrieve&dopt=Graphics&list_uids=5868) | s11680 | GCAAGCAAGUCCUAACAUUtt | AAUGUUAGGACUUGCUUGCct |
| NM_004162 | [RAB5A](http://www.ncbi.nlm.nih.gov/entrez/query.fcgi?db=gene&cmd=search&term=RAB5A) | [5868](http://www.ncbi.nlm.nih.gov/entrez/query.fcgi?db=gene&cmd=Retrieve&dopt=Graphics&list_uids=5868) | s11678 | GGAAGAGGAGUAGACCUUAtt | UAAGGUCUACUCCUCUUCCtc |
| NM_002868 | [RAB5B](http://www.ncbi.nlm.nih.gov/entrez/query.fcgi?db=gene&cmd=search&term=RAB5B) | [5869](http://www.ncbi.nlm.nih.gov/entrez/query.fcgi?db=gene&cmd=Retrieve&dopt=Graphics&list_uids=5869) | s11682 | GGAGCGAUAUCACAGCUUAtt | UAAGCUGUGAUAUCGCUCCtg |
| NM_002868 | [RAB5B](http://www.ncbi.nlm.nih.gov/entrez/query.fcgi?db=gene&cmd=search&term=RAB5B) | [5869](http://www.ncbi.nlm.nih.gov/entrez/query.fcgi?db=gene&cmd=Retrieve&dopt=Graphics&list_uids=5869) | s11683 | GGUAUUACGUUUUGUCAAAtt | UUUGACAAAACGUAAUACCag |
| NM_002868 | [RAB5B](http://www.ncbi.nlm.nih.gov/entrez/query.fcgi?db=gene&cmd=search&term=RAB5B) | [5869](http://www.ncbi.nlm.nih.gov/entrez/query.fcgi?db=gene&cmd=Retrieve&dopt=Graphics&list_uids=5869) | s11681 | CGACAUUACUAAUCAGGAAtt | UUCCUGAUUAGUAAUGUCGta |
| NM_004583 | [RAB5C](http://www.ncbi.nlm.nih.gov/entrez/query.fcgi?db=gene&cmd=search&term=RAB5C) | [5878](http://www.ncbi.nlm.nih.gov/entrez/query.fcgi?db=gene&cmd=Retrieve&dopt=Graphics&list_uids=5878) | s11710 | GCAAUGAACGUGAACGAAAtt | UUUCGUUCACGUUCAUUGCag |
| NM_004583 | [RAB5C](http://www.ncbi.nlm.nih.gov/entrez/query.fcgi?db=gene&cmd=search&term=RAB5C) | [5878](http://www.ncbi.nlm.nih.gov/entrez/query.fcgi?db=gene&cmd=Retrieve&dopt=Graphics&list_uids=5878) | s11708 | GGACAGGAGCGGUAUCACAtt | UGUGAUACCGCUCCUGUCCag |
| NM_004583 | [RAB5C](http://www.ncbi.nlm.nih.gov/entrez/query.fcgi?db=gene&cmd=search&term=RAB5C) | [5878](http://www.ncbi.nlm.nih.gov/entrez/query.fcgi?db=gene&cmd=Retrieve&dopt=Graphics&list_uids=5878) | s224512 | AGAGAGCCGUGGAAUUCCAtt | UGGAAUUCCACGGCUCUCUtg |
| NM_198896 | [RAB6A](http://www.ncbi.nlm.nih.gov/entrez/query.fcgi?db=gene&cmd=search&term=RAB6A) | [5870](http://www.ncbi.nlm.nih.gov/entrez/query.fcgi?db=gene&cmd=Retrieve&dopt=Graphics&list_uids=5870) | s11685 | GAGCUGAAUGUUAUGUUUAtt | UAAACAUAACAUUCAGCUCtt |
| NM_198896 | [RAB6A](http://www.ncbi.nlm.nih.gov/entrez/query.fcgi?db=gene&cmd=search&term=RAB6A) | [5870](http://www.ncbi.nlm.nih.gov/entrez/query.fcgi?db=gene&cmd=Retrieve&dopt=Graphics&list_uids=5870) | s11686 | AAUUCUUCACGGUAAGAAAtt | UUUCUUACCGUGAAGAAUUaa |
| NM_198896 | [RAB6A](http://www.ncbi.nlm.nih.gov/entrez/query.fcgi?db=gene&cmd=search&term=RAB6A) | [5870](http://www.ncbi.nlm.nih.gov/entrez/query.fcgi?db=gene&cmd=Retrieve&dopt=Graphics&list_uids=5870) | s11684 | CCCUAUUACCCUAUUCUUAtt | UAAGAAUAGGGUAAUAGGGaa |
| NM_016577 | [RAB6B](http://www.ncbi.nlm.nih.gov/entrez/query.fcgi?db=gene&cmd=search&term=RAB6B) | [51560](http://www.ncbi.nlm.nih.gov/entrez/query.fcgi?db=gene&cmd=Retrieve&dopt=Graphics&list_uids=51560) | s28327 | CCAUUGGGAUUGACUUCUUtt | AAGAAGUCAAUCCCAAUGGtt |
| NM_016577 | [RAB6B](http://www.ncbi.nlm.nih.gov/entrez/query.fcgi?db=gene&cmd=search&term=RAB6B) | [51560](http://www.ncbi.nlm.nih.gov/entrez/query.fcgi?db=gene&cmd=Retrieve&dopt=Graphics&list_uids=51560) | s28328 | UGAUUACGAGGUUCAUGUAtt | UACAUGAACCUCGUAAUCAga |
| NM_016577 | [RAB6B](http://www.ncbi.nlm.nih.gov/entrez/query.fcgi?db=gene&cmd=search&term=RAB6B) | [51560](http://www.ncbi.nlm.nih.gov/entrez/query.fcgi?db=gene&cmd=Retrieve&dopt=Graphics&list_uids=51560) | s28326 | AGGCAACCAUUGGGAUUGAtt | UCAAUCCCAAUGGUUGCCUgg |
| NM_032144 | [RAB6C](http://www.ncbi.nlm.nih.gov/entrez/query.fcgi?db=gene&cmd=search&term=RAB6C) | [84084](http://www.ncbi.nlm.nih.gov/entrez/query.fcgi?db=gene&cmd=Retrieve&dopt=Graphics&list_uids=84084) | s38490 | GCAUAUUCCGGUGAUAACUtt | AGUUAUCACCGGAAUAUGCtg |
| NM_032144 | [RAB6C](http://www.ncbi.nlm.nih.gov/entrez/query.fcgi?db=gene&cmd=search&term=RAB6C) | [84084](http://www.ncbi.nlm.nih.gov/entrez/query.fcgi?db=gene&cmd=Retrieve&dopt=Graphics&list_uids=84084) | s38489 | GCACUGAUGUAAAUUACUUtt | AAGUAAUUUACAUCAGUGCtc |
| NM_032144 | [RAB6C](http://www.ncbi.nlm.nih.gov/entrez/query.fcgi?db=gene&cmd=search&term=RAB6C) | [84084](http://www.ncbi.nlm.nih.gov/entrez/query.fcgi?db=gene&cmd=Retrieve&dopt=Graphics&list_uids=84084) | s195453 | UCGUGGAGGUGAUCUAUUAtt | UAAUAGAUCACCUCCACGAga |
| NM_004637 | [RAB7A](http://www.ncbi.nlm.nih.gov/entrez/query.fcgi?db=gene&cmd=search&term=RAB7A) | [7879](http://www.ncbi.nlm.nih.gov/entrez/query.fcgi?db=gene&cmd=Retrieve&dopt=Graphics&list_uids=7879) | s15443 | GCUAGUCACAAUGCAGAUAtt | UAUCUGCAUUGUGACUAGCct |
| NM_004637 | [RAB7A](http://www.ncbi.nlm.nih.gov/entrez/query.fcgi?db=gene&cmd=search&term=RAB7A) | [7879](http://www.ncbi.nlm.nih.gov/entrez/query.fcgi?db=gene&cmd=Retrieve&dopt=Graphics&list_uids=7879) | s15444 | GAGCUGACUUUCUGACCAAtt | UUGGUCAGAAAGUCAGCUCct |
| NM_004637 | [RAB7A](http://www.ncbi.nlm.nih.gov/entrez/query.fcgi?db=gene&cmd=search&term=RAB7A) | [7879](http://www.ncbi.nlm.nih.gov/entrez/query.fcgi?db=gene&cmd=Retrieve&dopt=Graphics&list_uids=7879) | s15442 | GCUGCGUUCUGGUAUUUGAtt | UCAAAUACCAGAACGCAGCag |
| NM_177403 | [RAB7B](http://www.ncbi.nlm.nih.gov/entrez/query.fcgi?db=gene&cmd=search&term=RAB7B) | [338382](http://www.ncbi.nlm.nih.gov/entrez/query.fcgi?db=gene&cmd=Retrieve&dopt=Graphics&list_uids=338382) | s50335 | GAAAGAUAUUCCUUACUUUtt | AAAGUAAGGAAUAUCUUUCtc |
| NM_177403 | [RAB7B](http://www.ncbi.nlm.nih.gov/entrez/query.fcgi?db=gene&cmd=search&term=RAB7B) | [338382](http://www.ncbi.nlm.nih.gov/entrez/query.fcgi?db=gene&cmd=Retrieve&dopt=Graphics&list_uids=338382) | s50334 | AGAUUAUCAUAUUGGGUGAtt | UCACCCAAUAUGAUAAUCUtg |
| NM_177403 | [RAB7B](http://www.ncbi.nlm.nih.gov/entrez/query.fcgi?db=gene&cmd=search&term=RAB7B) | [338382](http://www.ncbi.nlm.nih.gov/entrez/query.fcgi?db=gene&cmd=Retrieve&dopt=Graphics&list_uids=338382) | s50333 | GCUCUGUCGAGGUACCAGAtt | UCUGGUACCUCGACAGAGCcc |
| NM_003929 | [RAB7L1](http://www.ncbi.nlm.nih.gov/entrez/query.fcgi?db=gene&cmd=search&term=RAB7L1) | [8934](http://www.ncbi.nlm.nih.gov/entrez/query.fcgi?db=gene&cmd=Retrieve&dopt=Graphics&list_uids=8934) | s17084 | CGGUUUCACAGGUUGGACAtt | UGUCCAACCUGUGAAACCGtt |
| NM_003929 | [RAB7L1](http://www.ncbi.nlm.nih.gov/entrez/query.fcgi?db=gene&cmd=search&term=RAB7L1) | [8934](http://www.ncbi.nlm.nih.gov/entrez/query.fcgi?db=gene&cmd=Retrieve&dopt=Graphics&list_uids=8934) | s17082 | GAUUGACCGGUUCAGUAAAtt | UUUACUGAACCGGUCAAUCtg |
| NM_003929 | [RAB7L1](http://www.ncbi.nlm.nih.gov/entrez/query.fcgi?db=gene&cmd=search&term=RAB7L1) | [8934](http://www.ncbi.nlm.nih.gov/entrez/query.fcgi?db=gene&cmd=Retrieve&dopt=Graphics&list_uids=8934) | s17083 | UCACCUCUAUGACACGAUUtt | AAUCGUGUCAUAGAGGUGAag |
| NM_005370 | [RAB8A](http://www.ncbi.nlm.nih.gov/entrez/query.fcgi?db=gene&cmd=search&term=RAB8A) | [4218](http://www.ncbi.nlm.nih.gov/entrez/query.fcgi?db=gene&cmd=Retrieve&dopt=Graphics&list_uids=4218) | s8680 | GAGUCAAAAUCACACCGGAtt | UCCGGUGUGAUUUUGACUCcc |
| NM_005370 | [RAB8A](http://www.ncbi.nlm.nih.gov/entrez/query.fcgi?db=gene&cmd=search&term=RAB8A) | [4218](http://www.ncbi.nlm.nih.gov/entrez/query.fcgi?db=gene&cmd=Retrieve&dopt=Graphics&list_uids=4218) | s8679 | GCAAGAGAAUUAAACUGCAtt | UGCAGUUUAAUUCUCUUGCca |
| NM_005370 | [RAB8A](http://www.ncbi.nlm.nih.gov/entrez/query.fcgi?db=gene&cmd=search&term=RAB8A) | [4218](http://www.ncbi.nlm.nih.gov/entrez/query.fcgi?db=gene&cmd=Retrieve&dopt=Graphics&list_uids=4218) | s8681 | CUUUAAAAUUAGGACCAUAtt | UAUGGUCCUAAUUUUAAAGtc |
| NM_016530 | [RAB8B](http://www.ncbi.nlm.nih.gov/entrez/query.fcgi?db=gene&cmd=search&term=RAB8B) | [51762](http://www.ncbi.nlm.nih.gov/entrez/query.fcgi?db=gene&cmd=Retrieve&dopt=Graphics&list_uids=51762) | s28635 | GAAUGAUCCUGGGUAACAAtt | UUGUUACCCAGGAUCAUUCtt |
| NM_016530 | [RAB8B](http://www.ncbi.nlm.nih.gov/entrez/query.fcgi?db=gene&cmd=search&term=RAB8B) | [51762](http://www.ncbi.nlm.nih.gov/entrez/query.fcgi?db=gene&cmd=Retrieve&dopt=Graphics&list_uids=51762) | s28633 | GAAAGAUUCCGAACAAUCAtt | UGAUUGUUCGGAAUCUUUCct |
| NM_016530 | [RAB8B](http://www.ncbi.nlm.nih.gov/entrez/query.fcgi?db=gene&cmd=search&term=RAB8B) | [51762](http://www.ncbi.nlm.nih.gov/entrez/query.fcgi?db=gene&cmd=Retrieve&dopt=Graphics&list_uids=51762) | s28634 | CGAUAGAACUAGAUGGAAAtt | UUUCCAUCUAGUUCUAUCGtt |
| NM_004251 | [RAB9A](http://www.ncbi.nlm.nih.gov/entrez/query.fcgi?db=gene&cmd=search&term=RAB9A) | [9367](http://www.ncbi.nlm.nih.gov/entrez/query.fcgi?db=gene&cmd=Retrieve&dopt=Graphics&list_uids=9367) | s17917 | GGUCAGAUCAUUUGAUUCAtt | UGAAUCAAAUGAUCUGACCta |
| NM_004251 | [RAB9A](http://www.ncbi.nlm.nih.gov/entrez/query.fcgi?db=gene&cmd=search&term=RAB9A) | [9367](http://www.ncbi.nlm.nih.gov/entrez/query.fcgi?db=gene&cmd=Retrieve&dopt=Graphics&list_uids=9367) | s17916 | CCAGCUCUUCCAUACAAUAtt | UAUUGUAUGGAAGAGCUGGgt |
| NM_004251 | [RAB9A](http://www.ncbi.nlm.nih.gov/entrez/query.fcgi?db=gene&cmd=search&term=RAB9A) | [9367](http://www.ncbi.nlm.nih.gov/entrez/query.fcgi?db=gene&cmd=Retrieve&dopt=Graphics&list_uids=9367) | s17918 | GCUUCCAGAACUUAAGUAAtt | UUACUUAAGUUCUGGAAGCtt |
| NM_016370 | [RAB9B](http://www.ncbi.nlm.nih.gov/entrez/query.fcgi?db=gene&cmd=search&term=RAB9B) | [51209](http://www.ncbi.nlm.nih.gov/entrez/query.fcgi?db=gene&cmd=Retrieve&dopt=Graphics&list_uids=51209) | s27692 | GCUUAUGAACCGUUACGUAtt | UACGUAACGGUUCAUAAGCga |
| NM_016370 | [RAB9B](http://www.ncbi.nlm.nih.gov/entrez/query.fcgi?db=gene&cmd=search&term=RAB9B) | [51209](http://www.ncbi.nlm.nih.gov/entrez/query.fcgi?db=gene&cmd=Retrieve&dopt=Graphics&list_uids=51209) | s27691 | GGGUAACAAGGUAGACAAAtt | UUUGUCUACCUUGUUACCCag |
| NM_016370 | [RAB9B](http://www.ncbi.nlm.nih.gov/entrez/query.fcgi?db=gene&cmd=search&term=RAB9B) | [51209](http://www.ncbi.nlm.nih.gov/entrez/query.fcgi?db=gene&cmd=Retrieve&dopt=Graphics&list_uids=51209) | s27693 | AGAUGAUACUAAUGUGACAtt | UGUCACAUUAGUAUCAUCUtt |
| NM_007082 | [RABL2A](http://www.ncbi.nlm.nih.gov/entrez/query.fcgi?db=gene&cmd=search&term=RABL2A) | [11159](http://www.ncbi.nlm.nih.gov/entrez/query.fcgi?db=gene&cmd=Retrieve&dopt=Graphics&list_uids=11159) | s195089 | CGGUAGAUGGCAAGACCAUtt | AUGGUCUUGCCAUCUACCGtg |
| NM_007082 | [RABL2A](http://www.ncbi.nlm.nih.gov/entrez/query.fcgi?db=gene&cmd=search&term=RABL2A) | [11159](http://www.ncbi.nlm.nih.gov/entrez/query.fcgi?db=gene&cmd=Retrieve&dopt=Graphics&list_uids=11159) | s195090 | GUGUUUGAUAUACAGAGGAtt | UCCUCUGUAUAUCAAACACca |
| NM_007082 | [RABL2A](http://www.ncbi.nlm.nih.gov/entrez/query.fcgi?db=gene&cmd=search&term=RABL2A) | [11159](http://www.ncbi.nlm.nih.gov/entrez/query.fcgi?db=gene&cmd=Retrieve&dopt=Graphics&list_uids=11159) | s200490 | GGUGUUUGAUAUACAGAGGtt | CCUCUGUAUAUCAAACACCat |
| NM_007081 | [RABL2B](http://www.ncbi.nlm.nih.gov/entrez/query.fcgi?db=gene&cmd=search&term=RABL2B) | [11158](http://www.ncbi.nlm.nih.gov/entrez/query.fcgi?db=gene&cmd=Retrieve&dopt=Graphics&list_uids=11158) | s22011 | GCCUAGCUAUAGUUAGGAAtt | UUCCUAACUAUAGCUAGGCct |
| NM_007081 | [RABL2B](http://www.ncbi.nlm.nih.gov/entrez/query.fcgi?db=gene&cmd=search&term=RABL2B) | [11158](http://www.ncbi.nlm.nih.gov/entrez/query.fcgi?db=gene&cmd=Retrieve&dopt=Graphics&list_uids=11158) | s22012 | AAUCAUCUGUUUUUAAACAtt | UGUUUAAAAACAGAUGAUUgg |
| NM_007081 | [RABL2B](http://www.ncbi.nlm.nih.gov/entrez/query.fcgi?db=gene&cmd=search&term=RABL2B) | [11158](http://www.ncbi.nlm.nih.gov/entrez/query.fcgi?db=gene&cmd=Retrieve&dopt=Graphics&list_uids=11158) | s22013 | CCAGUCAUCUGGUUUGUUUtt | AAACAAACCAGAUGACUGGga |
| NM_173825 | [RABL3](http://www.ncbi.nlm.nih.gov/entrez/query.fcgi?db=gene&cmd=search&term=RABL3) | [285282](http://www.ncbi.nlm.nih.gov/entrez/query.fcgi?db=gene&cmd=Retrieve&dopt=Graphics&list_uids=285282) | s49875 | GUAUUAUUUUCGUACACGAtt | UCGUGUACGAAAAUAAUACca |
| NM_173825 | [RABL3](http://www.ncbi.nlm.nih.gov/entrez/query.fcgi?db=gene&cmd=search&term=RABL3) | [285282](http://www.ncbi.nlm.nih.gov/entrez/query.fcgi?db=gene&cmd=Retrieve&dopt=Graphics&list_uids=285282) | s49876 | GACCUACUACAUAGAAUUAtt | UAAUUCUAUGUAGUAGGUCtt |
| NM_173825 | [RABL3](http://www.ncbi.nlm.nih.gov/entrez/query.fcgi?db=gene&cmd=search&term=RABL3) | [285282](http://www.ncbi.nlm.nih.gov/entrez/query.fcgi?db=gene&cmd=Retrieve&dopt=Graphics&list_uids=285282) | s49874 | AGUUCAUGAUUACAAAGAAtt | UUCUUUGUAAUCAUGAACUct |
| NM_001077637 | [WTH3DI](http://www.ncbi.nlm.nih.gov/entrez/query.fcgi?db=gene&cmd=search&term=WTH3DI) | [150786](http://www.ncbi.nlm.nih.gov/entrez/query.fcgi?db=gene&cmd=Retrieve&dopt=Graphics&list_uids=150786) | s195786 | GCCAAACUAUUUCAGCGUAtt | UACGCUGAAAUAGUUUGGCtt |
| NM_001077637 | [WTH3DI](http://www.ncbi.nlm.nih.gov/entrez/query.fcgi?db=gene&cmd=search&term=WTH3DI) | [150786](http://www.ncbi.nlm.nih.gov/entrez/query.fcgi?db=gene&cmd=Retrieve&dopt=Graphics&list_uids=150786) | s195787 | GAUAAUGACGUUCUAAUGAtt | UCAUUAGAACGUCAUUAUCtt |
| NM_001077637 | [WTH3DI](http://www.ncbi.nlm.nih.gov/entrez/query.fcgi?db=gene&cmd=search&term=WTH3DI) | [150786](http://www.ncbi.nlm.nih.gov/entrez/query.fcgi?db=gene&cmd=Retrieve&dopt=Graphics&list_uids=150786) | s229454 | ACUAUUUCAGCGUAUUCCAtt | UGGAAUACGCUGAAAUAGUtt |
| NM_002866 | [RAB3A](http://www.ncbi.nlm.nih.gov/entrez/query.fcgi?db=gene&cmd=search&term=RAB3A) | [5864](http://www.ncbi.nlm.nih.gov/entrez/query.fcgi?db=gene&cmd=Retrieve&dopt=Graphics&list_uids=5864) | s11667 | CCAUCUAUCGCAACGACAAtt | UUGUCGUUGCGAUAGAUGGtc |
| NM_002866 | [RAB3A](http://www.ncbi.nlm.nih.gov/entrez/query.fcgi?db=gene&cmd=search&term=RAB3A) | [5864](http://www.ncbi.nlm.nih.gov/entrez/query.fcgi?db=gene&cmd=Retrieve&dopt=Graphics&list_uids=5864) | s11668 | CGACUACAUGUUCAAGAUUtt | AAUCUUGAACAUGUAGUCGaa |
| NM_002866 | [RAB3A](http://www.ncbi.nlm.nih.gov/entrez/query.fcgi?db=gene&cmd=search&term=RAB3A) | [5864](http://www.ncbi.nlm.nih.gov/entrez/query.fcgi?db=gene&cmd=Retrieve&dopt=Graphics&list_uids=5864) | s11666 | CCAUCACCACCGCAUACUAtt | UAGUAUGCGGUGGUGAUGGtc |
| **Signaling** |  |  |  |  |  |
| NM_024605 | [ARHGAP10](http://www.ncbi.nlm.nih.gov/entrez/query.fcgi?db=gene&cmd=search&term=ARHGAP10) | [79658](http://www.ncbi.nlm.nih.gov/entrez/query.fcgi?db=gene&cmd=Retrieve&dopt=Graphics&list_uids=79658) | s36029 | CACUAUUGCAUGUAUCGAAtt | UUCGAUACAUGCAAUAGUGtt |
| NM_024605 | [ARHGAP10](http://www.ncbi.nlm.nih.gov/entrez/query.fcgi?db=gene&cmd=search&term=ARHGAP10) | [79658](http://www.ncbi.nlm.nih.gov/entrez/query.fcgi?db=gene&cmd=Retrieve&dopt=Graphics&list_uids=79658) | s36030 | GAAUACACGGAAUCGAUUUtt | AAAUCGAUUCCGUGUAUUCtg |
| NM_024605 | [ARHGAP10](http://www.ncbi.nlm.nih.gov/entrez/query.fcgi?db=gene&cmd=search&term=ARHGAP10) | [79658](http://www.ncbi.nlm.nih.gov/entrez/query.fcgi?db=gene&cmd=Retrieve&dopt=Graphics&list_uids=79658) | s36028 | CAAACGAGCAAGUCAGUUUtt | AAACUGACUUGCUCGUUUGtg |
| NM_000118 | [ENG](http://www.ncbi.nlm.nih.gov/entrez/query.fcgi?db=gene&cmd=search&term=ENG) | [2022](http://www.ncbi.nlm.nih.gov/entrez/query.fcgi?db=gene&cmd=Retrieve&dopt=Graphics&list_uids=2022) | s4678 | GGACUGUCUUCAUGCGCUUtt | AAGCGCAUGAAGACAGUCCta |
| NM_000118 | [ENG](http://www.ncbi.nlm.nih.gov/entrez/query.fcgi?db=gene&cmd=search&term=ENG) | [2022](http://www.ncbi.nlm.nih.gov/entrez/query.fcgi?db=gene&cmd=Retrieve&dopt=Graphics&list_uids=2022) | s4679 | CAAGUAUGAUCAGCAAUGAtt | UCAUUGCUGAUCAUACUUGct |
| NM_000118 | [ENG](http://www.ncbi.nlm.nih.gov/entrez/query.fcgi?db=gene&cmd=search&term=ENG) | [2022](http://www.ncbi.nlm.nih.gov/entrez/query.fcgi?db=gene&cmd=Retrieve&dopt=Graphics&list_uids=2022) | s4677 | UGACCUGUCUGGUUGCACAtt | UGUGCAACCAGACAGGUCAgg |
| NM_013277 | [RACGAP1](http://www.ncbi.nlm.nih.gov/entrez/query.fcgi?db=gene&cmd=search&term=RACGAP1) | [29127](http://www.ncbi.nlm.nih.gov/entrez/query.fcgi?db=gene&cmd=Retrieve&dopt=Graphics&list_uids=29127) | s26552 | GCGAAAAGCUGGAACGACAtt | UGUCGUUCCAGCUUUUCGCag |
| NM_013277 | [RACGAP1](http://www.ncbi.nlm.nih.gov/entrez/query.fcgi?db=gene&cmd=search&term=RACGAP1) | [29127](http://www.ncbi.nlm.nih.gov/entrez/query.fcgi?db=gene&cmd=Retrieve&dopt=Graphics&list_uids=29127) | s26550 | CAGUGACUGUUCCCAAUGAtt | UCAUUGGGAACAGUCACUGta |
| NM_013277 | [RACGAP1](http://www.ncbi.nlm.nih.gov/entrez/query.fcgi?db=gene&cmd=search&term=RACGAP1) | [29127](http://www.ncbi.nlm.nih.gov/entrez/query.fcgi?db=gene&cmd=Retrieve&dopt=Graphics&list_uids=29127) | s26551 | CAACUAAGCGAGGAGCAAAtt | UUUGCUCCUCGCUUAGUUGaa |
| NM_004193 | [GBF1](http://www.ncbi.nlm.nih.gov/entrez/query.fcgi?db=gene&cmd=search&term=GBF1) | [8729](http://www.ncbi.nlm.nih.gov/entrez/query.fcgi?db=gene&cmd=Retrieve&dopt=Graphics&list_uids=8729) | s16633 | CAACCACAAUGUUCGUAAAtt | UUUACGAACAUUGUGGUUGtg |
| NM_004193 | [GBF1](http://www.ncbi.nlm.nih.gov/entrez/query.fcgi?db=gene&cmd=search&term=GBF1) | [8729](http://www.ncbi.nlm.nih.gov/entrez/query.fcgi?db=gene&cmd=Retrieve&dopt=Graphics&list_uids=8729) | s16635 | GAGCACUACUUGUACAUGAtt | UCAUGUACAAGUAGUGCUCgc |
| NM_004193 | [GBF1](http://www.ncbi.nlm.nih.gov/entrez/query.fcgi?db=gene&cmd=search&term=GBF1) | [8729](http://www.ncbi.nlm.nih.gov/entrez/query.fcgi?db=gene&cmd=Retrieve&dopt=Graphics&list_uids=8729) | s16634 | GCAUAGUUUCGGUCAUCUAtt | UAGAUGACCGAAACUAUGCag |
| NM_002890 | [RASA1](http://www.ncbi.nlm.nih.gov/entrez/query.fcgi?db=gene&cmd=search&term=RASA1) | [5921](http://www.ncbi.nlm.nih.gov/entrez/query.fcgi?db=gene&cmd=Retrieve&dopt=Graphics&list_uids=5921) | s11820 | CAUAGAUCACUAUCGAAAAtt | UUUUCGAUAGUGAUCUAUGat |
| NM_002890 | [RASA1](http://www.ncbi.nlm.nih.gov/entrez/query.fcgi?db=gene&cmd=search&term=RASA1) | [5921](http://www.ncbi.nlm.nih.gov/entrez/query.fcgi?db=gene&cmd=Retrieve&dopt=Graphics&list_uids=5921) | s11819 | GAAUCGUUGUUGUUAUGCAtt | UGCAUAACAACAACGAUUCaa |
| NM_002890 | [RASA1](http://www.ncbi.nlm.nih.gov/entrez/query.fcgi?db=gene&cmd=search&term=RASA1) | [5921](http://www.ncbi.nlm.nih.gov/entrez/query.fcgi?db=gene&cmd=Retrieve&dopt=Graphics&list_uids=5921) | s11821 | CCACGGAUGUUCAAUAUCAtt | UGAUAUUGAACAUCCGUGGat |
| NM_004296 | [RGS6](http://www.ncbi.nlm.nih.gov/entrez/query.fcgi?db=gene&cmd=search&term=RGS6) | [9628](http://www.ncbi.nlm.nih.gov/entrez/query.fcgi?db=gene&cmd=Retrieve&dopt=Graphics&list_uids=9628) | s18504 | GCGAGGAAGUGGGAAUUCAtt | UGAAUUCCCACUUCCUCGCaa |
| NM_004296 | [RGS6](http://www.ncbi.nlm.nih.gov/entrez/query.fcgi?db=gene&cmd=search&term=RGS6) | [9628](http://www.ncbi.nlm.nih.gov/entrez/query.fcgi?db=gene&cmd=Retrieve&dopt=Graphics&list_uids=9628) | s18505 | GGUCAAAUGCUUACCAGGAtt | UCCUGGUAAGCAUUUGACCgg |
| NM_004296 | [RGS6](http://www.ncbi.nlm.nih.gov/entrez/query.fcgi?db=gene&cmd=search&term=RGS6) | [9628](http://www.ncbi.nlm.nih.gov/entrez/query.fcgi?db=gene&cmd=Retrieve&dopt=Graphics&list_uids=9628) | s18503 | CAGUUGAAGCAAUACACUUtt | AAGUGUAUUGCUUCAACUGgg |
| NM_022771 | [TBC1D15](http://www.ncbi.nlm.nih.gov/entrez/query.fcgi?db=gene&cmd=search&term=TBC1D15) | [64786](http://www.ncbi.nlm.nih.gov/entrez/query.fcgi?db=gene&cmd=Retrieve&dopt=Graphics&list_uids=64786) | s34945 | CGGCAACUAUGAUAGGAUUtt | AAUCCUAUCAUAGUUGCCGta |
| NM_022771 | [TBC1D15](http://www.ncbi.nlm.nih.gov/entrez/query.fcgi?db=gene&cmd=search&term=TBC1D15) | [64786](http://www.ncbi.nlm.nih.gov/entrez/query.fcgi?db=gene&cmd=Retrieve&dopt=Graphics&list_uids=64786) | s34946 | GCAUUAGAUUCCUCUAGUAtt | UACUAGAGGAAUCUAAUGCat |
| NM_022771 | [TBC1D15](http://www.ncbi.nlm.nih.gov/entrez/query.fcgi?db=gene&cmd=search&term=TBC1D15) | [64786](http://www.ncbi.nlm.nih.gov/entrez/query.fcgi?db=gene&cmd=Retrieve&dopt=Graphics&list_uids=64786) | s34944 | GAAAGACCAAUGACCAAGAtt | UCUUGGUCAUUGGUCUUUCca |
| NM_014715 | [ARHGAP32](http://www.ncbi.nlm.nih.gov/entrez/query.fcgi?db=gene&cmd=search&term=) | [9743](http://www.ncbi.nlm.nih.gov/entrez/query.fcgi?db=gene&cmd=Retrieve&dopt=Graphics&list_uids=9743) | s18798 | GGCUGAUAAAAAUCCACGAtt | UCGUGGAUUUUUAUCAGCCtt |
| NM_014715 | [ARHGAP32](http://www.ncbi.nlm.nih.gov/entrez/query.fcgi?db=gene&cmd=search&term=) | [9743](http://www.ncbi.nlm.nih.gov/entrez/query.fcgi?db=gene&cmd=Retrieve&dopt=Graphics&list_uids=9743) | s18799 | GGCCUAUUUCCAAACCGAUtt | AUCGGUUUGGAAAUAGGCCtt |
| NM_014715 | [ARHGAP32](http://www.ncbi.nlm.nih.gov/entrez/query.fcgi?db=gene&cmd=search&term=) | [9743](http://www.ncbi.nlm.nih.gov/entrez/query.fcgi?db=gene&cmd=Retrieve&dopt=Graphics&list_uids=9743) | s18797 | GACAUUCCACCGUACCCUAtt | UAGGGUACGGUGGAAUGUCtt |
| NM_031407 | [HUWE1](http://www.ncbi.nlm.nih.gov/entrez/query.fcgi?db=gene&cmd=search&term=HUWE1) | [10075](http://www.ncbi.nlm.nih.gov/entrez/query.fcgi?db=gene&cmd=Retrieve&dopt=Graphics&list_uids=10075) | s19595 | CAUUGGAAAGUGCGAGUUAtt | UAACUCGCACUUUCCAAUGtt |
| NM_031407 | [HUWE1](http://www.ncbi.nlm.nih.gov/entrez/query.fcgi?db=gene&cmd=search&term=HUWE1) | [10075](http://www.ncbi.nlm.nih.gov/entrez/query.fcgi?db=gene&cmd=Retrieve&dopt=Graphics&list_uids=10075) | s19596 | CUGUGAGAGUGAUCGGGAAtt | UUCCCGAUCACUCUCACAGtt |
| NM_031407 | [HUWE1](http://www.ncbi.nlm.nih.gov/entrez/query.fcgi?db=gene&cmd=search&term=HUWE1) | [10075](http://www.ncbi.nlm.nih.gov/entrez/query.fcgi?db=gene&cmd=Retrieve&dopt=Graphics&list_uids=10075) | s19597 | GGUCUAAUCAUGCCGCAGAtt | UCUGCGGCAUGAUUAGACCtt |
| NM_014869 | [IQSEC1](http://www.ncbi.nlm.nih.gov/entrez/query.fcgi?db=gene&cmd=search&term=IQSEC1) | [9922](http://www.ncbi.nlm.nih.gov/entrez/query.fcgi?db=gene&cmd=Retrieve&dopt=Graphics&list_uids=9922) | s19245 | GGACUACACCAGCGAGAAAtt | UUUCUCGCUGGUGUAGUCCga |
| NM_014869 | [IQSEC1](http://www.ncbi.nlm.nih.gov/entrez/query.fcgi?db=gene&cmd=search&term=IQSEC1) | [9922](http://www.ncbi.nlm.nih.gov/entrez/query.fcgi?db=gene&cmd=Retrieve&dopt=Graphics&list_uids=9922) | s19246 | GAGCAGAUAUCAAAGUGUUtt | AACACUUUGAUAUCUGCUCcg |
| NM_014869 | [IQSEC1](http://www.ncbi.nlm.nih.gov/entrez/query.fcgi?db=gene&cmd=search&term=IQSEC1) | [9922](http://www.ncbi.nlm.nih.gov/entrez/query.fcgi?db=gene&cmd=Retrieve&dopt=Graphics&list_uids=9922) | s19247 | GGAUUGUGCUGUCCAACAUtt | AUGUUGGACAGCACAAUCCgg |
| NM_025072 | [PTGES2](http://www.ncbi.nlm.nih.gov/entrez/query.fcgi?db=gene&cmd=search&term=PTGES2) | [80142](http://www.ncbi.nlm.nih.gov/entrez/query.fcgi?db=gene&cmd=Retrieve&dopt=Graphics&list_uids=80142) | s36952 | GGCUCAUGCUCAACGAGAAtt | UUCUCGUUGAGCAUGAGCCag |
| NM_025072 | [PTGES2](http://www.ncbi.nlm.nih.gov/entrez/query.fcgi?db=gene&cmd=search&term=PTGES2) | [80142](http://www.ncbi.nlm.nih.gov/entrez/query.fcgi?db=gene&cmd=Retrieve&dopt=Graphics&list_uids=80142) | s36951 | GAUGCAUUCGAUGACCUGAtt | UCAGGUCAUCGAAUGCAUCca |
| NM_025072 | [PTGES2](http://www.ncbi.nlm.nih.gov/entrez/query.fcgi?db=gene&cmd=search&term=PTGES2) | [80142](http://www.ncbi.nlm.nih.gov/entrez/query.fcgi?db=gene&cmd=Retrieve&dopt=Graphics&list_uids=80142) | s36953 | GCAGGACGGUUUGUUUUCAtt | UGAAAACAAACCGUCCUGCtg |
| NM_006463 | [STAMBP](http://www.ncbi.nlm.nih.gov/entrez/query.fcgi?db=gene&cmd=search&term=STAMBP) | [10617](http://www.ncbi.nlm.nih.gov/entrez/query.fcgi?db=gene&cmd=Retrieve&dopt=Graphics&list_uids=10617) | s223141 | CGAUAUACCAAAGAAUAUAtt | UAUAUUCUUUGGUAUAUCGtt |
| NM_006463 | [STAMBP](http://www.ncbi.nlm.nih.gov/entrez/query.fcgi?db=gene&cmd=search&term=STAMBP) | [10617](http://www.ncbi.nlm.nih.gov/entrez/query.fcgi?db=gene&cmd=Retrieve&dopt=Graphics&list_uids=10617) | s223142 | CUGUCAUUCCUGAAAAGAAtt | UUCUUUUCAGGAAUGACAGca |
| NM_006463 | [STAMBP](http://www.ncbi.nlm.nih.gov/entrez/query.fcgi?db=gene&cmd=search&term=STAMBP) | [10617](http://www.ncbi.nlm.nih.gov/entrez/query.fcgi?db=gene&cmd=Retrieve&dopt=Graphics&list_uids=10617) | s20853 | GAGUUGAGAUUAUCCGAAUtt | AUUCGGAUAAUCUCAACUCca |
| NM_013254 | [TBK1](http://www.ncbi.nlm.nih.gov/entrez/query.fcgi?db=gene&cmd=search&term=TBK1) | [29110](http://www.ncbi.nlm.nih.gov/entrez/query.fcgi?db=gene&cmd=Retrieve&dopt=Graphics&list_uids=29110) | s761 | GAACGUAGAUUAGCUUAUAtt | UAUAAGCUAAUCUACGUUCtg |
| NM_013254 | [TBK1](http://www.ncbi.nlm.nih.gov/entrez/query.fcgi?db=gene&cmd=search&term=TBK1) | [29110](http://www.ncbi.nlm.nih.gov/entrez/query.fcgi?db=gene&cmd=Retrieve&dopt=Graphics&list_uids=29110) | s763 | GGCACAACAUUUCCCUAAAtt | UUUAGGGAAAUGUUGUGCCag |
| NM_013254 | [TBK1](http://www.ncbi.nlm.nih.gov/entrez/query.fcgi?db=gene&cmd=search&term=TBK1) | [29110](http://www.ncbi.nlm.nih.gov/entrez/query.fcgi?db=gene&cmd=Retrieve&dopt=Graphics&list_uids=29110) | s762 | GGAACCUCUGAAUACCAUAtt | UAUGGUAUUCAGAGGUUCCcg |
| NM_001012750 | [ABI1](http://www.ncbi.nlm.nih.gov/entrez/query.fcgi?db=gene&cmd=search&term=ABI1) | [10006](http://www.ncbi.nlm.nih.gov/entrez/query.fcgi?db=gene&cmd=Retrieve&dopt=Graphics&list_uids=10006) | s19444 | CUCUAGCUAGUGUUGCUUAtt | UAAGCAACACUAGCUAGAGat |
| NM_001012750 | [ABI1](http://www.ncbi.nlm.nih.gov/entrez/query.fcgi?db=gene&cmd=search&term=ABI1) | [10006](http://www.ncbi.nlm.nih.gov/entrez/query.fcgi?db=gene&cmd=Retrieve&dopt=Graphics&list_uids=10006) | s19446 | GGUAUAUUCGGAAACCUAUtt | AUAGGUUUCCGAAUAUACCtt |
| NM_001012750 | [ABI1](http://www.ncbi.nlm.nih.gov/entrez/query.fcgi?db=gene&cmd=search&term=ABI1) | [10006](http://www.ncbi.nlm.nih.gov/entrez/query.fcgi?db=gene&cmd=Retrieve&dopt=Graphics&list_uids=10006) | s19445 | CAACAGUUCCUAAUGACUAtt | UAGUCAUUAGGAACUGUUGgg |
| **ESCRT-I and -II** |  |  |  |  |  |
| NM_007241 | [SNF8](http://www.ncbi.nlm.nih.gov/entrez/query.fcgi?db=gene&cmd=search&term=SNF8) | [11267](http://www.ncbi.nlm.nih.gov/entrez/query.fcgi?db=gene&cmd=Retrieve&dopt=Graphics&list_uids=11267) | s22249 | CGAACUAGGUGUCCAAAUUtt | AAUUUGGACACCUAGUUCGta |
| NM_007241 | [SNF8](http://www.ncbi.nlm.nih.gov/entrez/query.fcgi?db=gene&cmd=search&term=SNF8) | [11267](http://www.ncbi.nlm.nih.gov/entrez/query.fcgi?db=gene&cmd=Retrieve&dopt=Graphics&list_uids=11267) | s22247 | CAACAGGUGUUGAAGGGAAtt | UUCCCUUCAACACCUGUUGat |
| NM_007241 | [SNF8](http://www.ncbi.nlm.nih.gov/entrez/query.fcgi?db=gene&cmd=search&term=SNF8) | [11267](http://www.ncbi.nlm.nih.gov/entrez/query.fcgi?db=gene&cmd=Retrieve&dopt=Graphics&list_uids=11267) | s22248 | GCCCAGAUGUCAAAGCAGUtt | ACUGCUUUGACAUCUGGGCta |
| NM_004712 | [HGS](http://www.ncbi.nlm.nih.gov/entrez/query.fcgi?db=gene&cmd=search&term=HGS) | [9146](http://www.ncbi.nlm.nih.gov/entrez/query.fcgi?db=gene&cmd=Retrieve&dopt=Graphics&list_uids=9146) | s17482 | CACGGUAUCUCAACCGGAAtt | UUCCGGUUGAGAUACCGUGcg |
| NM_004712 | [HGS](http://www.ncbi.nlm.nih.gov/entrez/query.fcgi?db=gene&cmd=search&term=HGS) | [9146](http://www.ncbi.nlm.nih.gov/entrez/query.fcgi?db=gene&cmd=Retrieve&dopt=Graphics&list_uids=9146) | s17481 | UGGAAUCUGUGGUAAAGAAtt | UUCUUUACCACAGAUUCCAtg |
| NM_004712 | [HGS](http://www.ncbi.nlm.nih.gov/entrez/query.fcgi?db=gene&cmd=search&term=HGS) | [9146](http://www.ncbi.nlm.nih.gov/entrez/query.fcgi?db=gene&cmd=Retrieve&dopt=Graphics&list_uids=9146) | s17480 | CGUCUUUCCAGAAUUCAAAtt | UUUGAAUUCUGGAAAGACGtg |
| NM_013374 | [PDCD6IP](http://www.ncbi.nlm.nih.gov/entrez/query.fcgi?db=gene&cmd=search&term=PDCD6IP) | [10015](http://www.ncbi.nlm.nih.gov/entrez/query.fcgi?db=gene&cmd=Retrieve&dopt=Graphics&list_uids=10015) | s19466 | GCUAUAAUCCUUAUGCGUAtt | UACGCAUAAGGAUUAUAGCcc |
| NM_013374 | [PDCD6IP](http://www.ncbi.nlm.nih.gov/entrez/query.fcgi?db=gene&cmd=search&term=PDCD6IP) | [10015](http://www.ncbi.nlm.nih.gov/entrez/query.fcgi?db=gene&cmd=Retrieve&dopt=Graphics&list_uids=10015) | s19465 | GAAGGAUGCUUUCGAUAAAtt | UUUAUCGAAAGCAUCCUUCca |
| NM_013374 | [PDCD6IP](http://www.ncbi.nlm.nih.gov/entrez/query.fcgi?db=gene&cmd=search&term=PDCD6IP) | [10015](http://www.ncbi.nlm.nih.gov/entrez/query.fcgi?db=gene&cmd=Retrieve&dopt=Graphics&list_uids=10015) | s19467 | GAAUUACUGCAACGAAAUAtt | UAUUUCGUUGCAGUAAUUCag |
| NM_003473 | [STAM](http://www.ncbi.nlm.nih.gov/entrez/query.fcgi?db=gene&cmd=search&term=STAM) | [8027](http://www.ncbi.nlm.nih.gov/entrez/query.fcgi?db=gene&cmd=Retrieve&dopt=Graphics&list_uids=8027) | s224851 | GCAACCAGCGAGAUGAAUAtt | UAUUCAUCUCGCUGGUUGCtt |
| NM_003473 | [STAM](http://www.ncbi.nlm.nih.gov/entrez/query.fcgi?db=gene&cmd=search&term=STAM) | [8027](http://www.ncbi.nlm.nih.gov/entrez/query.fcgi?db=gene&cmd=Retrieve&dopt=Graphics&list_uids=8027) | s224852 | GGCUUUGACUCUUCUAGGAtt | UCCUAGAAGAGUCAAAGCCtg |
| NM_003473 | [STAM](http://www.ncbi.nlm.nih.gov/entrez/query.fcgi?db=gene&cmd=search&term=STAM) | [8027](http://www.ncbi.nlm.nih.gov/entrez/query.fcgi?db=gene&cmd=Retrieve&dopt=Graphics&list_uids=8027) | s224853 | GUAAGCAACGUAUUAAAUAtt | UAUUUAAUACGUUGCUUACtt |
| NM_006292 | [TSG101](http://www.ncbi.nlm.nih.gov/entrez/query.fcgi?db=gene&cmd=search&term=TSG101) | [7251](http://www.ncbi.nlm.nih.gov/entrez/query.fcgi?db=gene&cmd=Retrieve&dopt=Graphics&list_uids=7251) | s14441 | GAGACCUAACUGUACGUGAtt | UCACGUACAGUUAGGUCUCtg |
| NM_006292 | [TSG101](http://www.ncbi.nlm.nih.gov/entrez/query.fcgi?db=gene&cmd=search&term=TSG101) | [7251](http://www.ncbi.nlm.nih.gov/entrez/query.fcgi?db=gene&cmd=Retrieve&dopt=Graphics&list_uids=7251) | s14440 | CUGUCAAUGUUAUUACUCUtt | AGAGUAAUAACAUUGACAGtt |
| NM_006292 | [TSG101](http://www.ncbi.nlm.nih.gov/entrez/query.fcgi?db=gene&cmd=search&term=TSG101) | [7251](http://www.ncbi.nlm.nih.gov/entrez/query.fcgi?db=gene&cmd=Retrieve&dopt=Graphics&list_uids=7251) | s14439 | GAAAAAGGGUCACCAGAAAtt | UUUCUGGUGACCCUUUUUCag |
| NM_016525 | [UBAP1](http://www.ncbi.nlm.nih.gov/entrez/query.fcgi?db=gene&cmd=search&term=UBAP1) | [51271](http://www.ncbi.nlm.nih.gov/entrez/query.fcgi?db=gene&cmd=Retrieve&dopt=Graphics&list_uids=51271) | s27814 | CUCAGUUAUUGGACAAUAAtt | UUAUUGUCCAAUAACUGAGcc |
| NM_016525 | [UBAP1](http://www.ncbi.nlm.nih.gov/entrez/query.fcgi?db=gene&cmd=search&term=UBAP1) | [51271](http://www.ncbi.nlm.nih.gov/entrez/query.fcgi?db=gene&cmd=Retrieve&dopt=Graphics&list_uids=51271) | s27813 | GGCUUUGAGCUGAAAGACAtt | UGUCUUUCAGCUCAAAGCCca |
| NM_016525 | [UBAP1](http://www.ncbi.nlm.nih.gov/entrez/query.fcgi?db=gene&cmd=search&term=UBAP1) | [51271](http://www.ncbi.nlm.nih.gov/entrez/query.fcgi?db=gene&cmd=Retrieve&dopt=Graphics&list_uids=51271) | s27812 | GCUGAGAAAUAUUCUGGUAtt | UACCAGAAUAUUUCUCAGCtc |
| NM_005154 | [USP8](http://www.ncbi.nlm.nih.gov/entrez/query.fcgi?db=gene&cmd=search&term=USP8) | [9101](http://www.ncbi.nlm.nih.gov/entrez/query.fcgi?db=gene&cmd=Retrieve&dopt=Graphics&list_uids=9101) | s17372 | GGAUGAUAUUAACAGGUCAtt | UGACCUGUUAAUAUCAUCCtg |
| NM_005154 | [USP8](http://www.ncbi.nlm.nih.gov/entrez/query.fcgi?db=gene&cmd=search&term=USP8) | [9101](http://www.ncbi.nlm.nih.gov/entrez/query.fcgi?db=gene&cmd=Retrieve&dopt=Graphics&list_uids=9101) | s17370 | GGAUACAGACGAUACCGAAtt | UUCGGUAUCGUCUGUAUCCtc |
| NM_005154 | [USP8](http://www.ncbi.nlm.nih.gov/entrez/query.fcgi?db=gene&cmd=search&term=USP8) | [9101](http://www.ncbi.nlm.nih.gov/entrez/query.fcgi?db=gene&cmd=Retrieve&dopt=Graphics&list_uids=9101) | s17371 | CUCCAACAGUUAAUCGGGAtt | UCCCGAUUAACUGUUGGAGtt |
| NM_032353 | [VPS25](http://www.ncbi.nlm.nih.gov/entrez/query.fcgi?db=gene&cmd=search&term=VPS25) | [84313](http://www.ncbi.nlm.nih.gov/entrez/query.fcgi?db=gene&cmd=Retrieve&dopt=Graphics&list_uids=84313) | s38895 | CGAUCCAGAUUGUAUUAGAtt | UCUAAUACAAUCUGGAUCGac |
| NM_032353 | [VPS25](http://www.ncbi.nlm.nih.gov/entrez/query.fcgi?db=gene&cmd=search&term=VPS25) | [84313](http://www.ncbi.nlm.nih.gov/entrez/query.fcgi?db=gene&cmd=Retrieve&dopt=Graphics&list_uids=84313) | s38894 | CGUCUUUACCCUGUAUGAAtt | UUCAUACAGGGUAAAGACGga |
| NM_032353 | [VPS25](http://www.ncbi.nlm.nih.gov/entrez/query.fcgi?db=gene&cmd=search&term=VPS25) | [84313](http://www.ncbi.nlm.nih.gov/entrez/query.fcgi?db=gene&cmd=Retrieve&dopt=Graphics&list_uids=84313) | s38896 | CCCACCCUUCUUUACGUUAtt | UAACGUAAAGAAGGGUGGGaa |
| NM_016208 | [VPS28](http://www.ncbi.nlm.nih.gov/entrez/query.fcgi?db=gene&cmd=search&term=VPS28) | [51160](http://www.ncbi.nlm.nih.gov/entrez/query.fcgi?db=gene&cmd=Retrieve&dopt=Graphics&list_uids=51160) | s27579 | GAAGUGAAGUUGUACAAGAtt | UCUUGUACAACUUCACUUCct |
| NM_016208 | [VPS28](http://www.ncbi.nlm.nih.gov/entrez/query.fcgi?db=gene&cmd=search&term=VPS28) | [51160](http://www.ncbi.nlm.nih.gov/entrez/query.fcgi?db=gene&cmd=Retrieve&dopt=Graphics&list_uids=51160) | s27578 | GGAACAAGCCGGAGCUGUAtt | UACAGCUCCGGCUUGUUCCca |
| NM_016208 | [VPS28](http://www.ncbi.nlm.nih.gov/entrez/query.fcgi?db=gene&cmd=search&term=VPS28) | [51160](http://www.ncbi.nlm.nih.gov/entrez/query.fcgi?db=gene&cmd=Retrieve&dopt=Graphics&list_uids=51160) | s27577 | AAUCAGCUCUAUUGACGAAtt | UUCGUCAAUAGAGCUGAUUtc |
| NM_016075 | [VPS36](http://www.ncbi.nlm.nih.gov/entrez/query.fcgi?db=gene&cmd=search&term=VPS36) | [51028](http://www.ncbi.nlm.nih.gov/entrez/query.fcgi?db=gene&cmd=Retrieve&dopt=Graphics&list_uids=51028) | s27277 | GCCCAUUCCAGAGUAGUAAtt | UUACUACUCUGGAAUGGGCca |
| NM_016075 | [VPS36](http://www.ncbi.nlm.nih.gov/entrez/query.fcgi?db=gene&cmd=search&term=VPS36) | [51028](http://www.ncbi.nlm.nih.gov/entrez/query.fcgi?db=gene&cmd=Retrieve&dopt=Graphics&list_uids=51028) | s27276 | CAGUUACCAGAGAAACCUAtt | UAGGUUUCUCUGGUAACUGgg |
| NM_016075 | [VPS36](http://www.ncbi.nlm.nih.gov/entrez/query.fcgi?db=gene&cmd=search&term=VPS36) | [51028](http://www.ncbi.nlm.nih.gov/entrez/query.fcgi?db=gene&cmd=Retrieve&dopt=Graphics&list_uids=51028) | s27278 | CCCAGUCAUUACAAACAAAtt | UUUGUUUGUAAUGACUGGGaa |
| **ESCRT-III** |  |  |  |  |  |
| NM_001083314 | [CHMP1A](http://www.ncbi.nlm.nih.gov/entrez/query.fcgi?db=gene&cmd=search&term=CHMP1A) | [5119](http://www.ncbi.nlm.nih.gov/entrez/query.fcgi?db=gene&cmd=Retrieve&dopt=Graphics&list_uids=5119) | s10142 | CGUCUUGCCUUUGUGCUGAtt | UCAGCACAAAGGCAAGACGcg |
| NM_001083314 | [CHMP1A](http://www.ncbi.nlm.nih.gov/entrez/query.fcgi?db=gene&cmd=search&term=CHMP1A) | [5119](http://www.ncbi.nlm.nih.gov/entrez/query.fcgi?db=gene&cmd=Retrieve&dopt=Graphics&list_uids=5119) | s10143 | GGUGUUGAGUUUCUGCAAAtt | UUUGCAGAAACUCAACACCag |
| NM_001083314 | [CHMP1A](http://www.ncbi.nlm.nih.gov/entrez/query.fcgi?db=gene&cmd=search&term=CHMP1A) | [5119](http://www.ncbi.nlm.nih.gov/entrez/query.fcgi?db=gene&cmd=Retrieve&dopt=Graphics&list_uids=5119) | s194678 | GCCUUAGGGUUGUUCCUGUtt | ACAGGAACAACCCUAAGGCca |
| NM_198426 | [CHMP2A](http://www.ncbi.nlm.nih.gov/entrez/query.fcgi?db=gene&cmd=search&term=CHMP2A) | [27243](http://www.ncbi.nlm.nih.gov/entrez/query.fcgi?db=gene&cmd=Retrieve&dopt=Graphics&list_uids=27243) | s26029 | GACUUAGCCUAACAGAUGAtt | UCAUCUGUUAGGCUAAGUCcc |
| NM_198426 | [CHMP2A](http://www.ncbi.nlm.nih.gov/entrez/query.fcgi?db=gene&cmd=search&term=CHMP2A) | [27243](http://www.ncbi.nlm.nih.gov/entrez/query.fcgi?db=gene&cmd=Retrieve&dopt=Graphics&list_uids=27243) | s26027 | AGAUGGAUGCUGUUCGCAUtt | AUGCGAACAGCAUCCAUCUgg |
| NM_198426 | [CHMP2A](http://www.ncbi.nlm.nih.gov/entrez/query.fcgi?db=gene&cmd=search&term=CHMP2A) | [27243](http://www.ncbi.nlm.nih.gov/entrez/query.fcgi?db=gene&cmd=Retrieve&dopt=Graphics&list_uids=27243) | s26028 | AGAAGAUCAUGAUGGAGUUtt | AACUCCAUCAUGAUCUUCUgg |
| NM_014169 | [CHMP4A](http://www.ncbi.nlm.nih.gov/entrez/query.fcgi?db=gene&cmd=search&term=CHMP4A) | [29082](http://www.ncbi.nlm.nih.gov/entrez/query.fcgi?db=gene&cmd=Retrieve&dopt=Graphics&list_uids=29082) | s26443 | CCCUCAGUCAAAUUGCCUAtt | UAGGCAAUUUGACUGAGGGtt |
| NM_014169 | [CHMP4A](http://www.ncbi.nlm.nih.gov/entrez/query.fcgi?db=gene&cmd=search&term=CHMP4A) | [29082](http://www.ncbi.nlm.nih.gov/entrez/query.fcgi?db=gene&cmd=Retrieve&dopt=Graphics&list_uids=29082) | s26441 | GGCUUUGCGGAGGAAGAAAtt | UUUCUUCCUCCGCAAAGCCtg |
| NM_014169 | [CHMP4A](http://www.ncbi.nlm.nih.gov/entrez/query.fcgi?db=gene&cmd=search&term=CHMP4A) | [29082](http://www.ncbi.nlm.nih.gov/entrez/query.fcgi?db=gene&cmd=Retrieve&dopt=Graphics&list_uids=29082) | s26442 | CAUGGACAUUGACAAGGUAtt | UACCUUGUCAAUGUCCAUGtc |
| NM_016410 | [CHMP5](http://www.ncbi.nlm.nih.gov/entrez/query.fcgi?db=gene&cmd=search&term=CHMP5) | [51510](http://www.ncbi.nlm.nih.gov/entrez/query.fcgi?db=gene&cmd=Retrieve&dopt=Graphics&list_uids=51510) | s224218 | AGAUUGAGGAUUUACAAGAtt | UCUUGUAAAUCCUCAAUCUgg |
| NM_016410 | [CHMP5](http://www.ncbi.nlm.nih.gov/entrez/query.fcgi?db=gene&cmd=search&term=CHMP5) | [51510](http://www.ncbi.nlm.nih.gov/entrez/query.fcgi?db=gene&cmd=Retrieve&dopt=Graphics&list_uids=51510) | s28237 | CUAUGAAACUGGGAGUAAAtt | UUUACUCCCAGUUUCAUAGca |
| NM_016410 | [CHMP5](http://www.ncbi.nlm.nih.gov/entrez/query.fcgi?db=gene&cmd=search&term=CHMP5) | [51510](http://www.ncbi.nlm.nih.gov/entrez/query.fcgi?db=gene&cmd=Retrieve&dopt=Graphics&list_uids=51510) | s28239 | GCCCAACAGUCAUUCAACAtt | UGUUGAAUGACUGUUGGGCaa |
| XM_001132548 | [CHMP6](http://www.ncbi.nlm.nih.gov/entrez/query.fcgi?db=gene&cmd=search&term=CHMP6) | [79643](http://www.ncbi.nlm.nih.gov/entrez/query.fcgi?db=gene&cmd=Retrieve&dopt=Graphics&list_uids=79643) | s35991 | GGAAAUGAGUGUCUGAACAtt | UGUUCAGACACUCAUUUCCaa |
| XM_001132548 | [CHMP6](http://www.ncbi.nlm.nih.gov/entrez/query.fcgi?db=gene&cmd=search&term=CHMP6) | [79643](http://www.ncbi.nlm.nih.gov/entrez/query.fcgi?db=gene&cmd=Retrieve&dopt=Graphics&list_uids=79643) | s35990 | CAAUCACUCAGGAACAAAUtt | AUUUGUUCCUGAGUGAUUGcg |
| XM_001132548 | [CHMP6](http://www.ncbi.nlm.nih.gov/entrez/query.fcgi?db=gene&cmd=search&term=CHMP6) | [79643](http://www.ncbi.nlm.nih.gov/entrez/query.fcgi?db=gene&cmd=Retrieve&dopt=Graphics&list_uids=79643) | s35989 | GAGUUCACCCAGAUCGAAAtt | UUUCGAUCUGGGUGAACUCaa |
| NM_014761 | [IST1](http://www.ncbi.nlm.nih.gov/entrez/query.fcgi?db=gene&cmd=search&term=IST1) | [9798](http://www.ncbi.nlm.nih.gov/entrez/query.fcgi?db=gene&cmd=Retrieve&dopt=Graphics&list_uids=9798) | s18939 | GGCCUUAUCCAGUCUAUGAtt | UCAUAGACUGGAUAAGGCCaa |
| NM_014761 | [IST1](http://www.ncbi.nlm.nih.gov/entrez/query.fcgi?db=gene&cmd=search&term=IST1) | [9798](http://www.ncbi.nlm.nih.gov/entrez/query.fcgi?db=gene&cmd=Retrieve&dopt=Graphics&list_uids=9798) | s18941 | GCAGAUAACUAUGACAACUtt | AGUUGUCAUAGUUAUCUGCag |
| NM_014761 | [IST1](http://www.ncbi.nlm.nih.gov/entrez/query.fcgi?db=gene&cmd=search&term=IST1) | [9798](http://www.ncbi.nlm.nih.gov/entrez/query.fcgi?db=gene&cmd=Retrieve&dopt=Graphics&list_uids=9798) | s18940 | GAGAGAUACCUGAUUGAAAtt | UUUCAAUCAGGUAUCUCUCca |
| NM_001005753 | [VPS24](http://www.ncbi.nlm.nih.gov/entrez/query.fcgi?db=gene&cmd=search&term=VPS24) | [51652](http://www.ncbi.nlm.nih.gov/entrez/query.fcgi?db=gene&cmd=Retrieve&dopt=Graphics&list_uids=51652) | s28473 | GCAUGGACGAUCAGGAAGAtt | UCUUCCUGAUCGUCCAUGCtt |
| NM_001005753 | [VPS24](http://www.ncbi.nlm.nih.gov/entrez/query.fcgi?db=gene&cmd=search&term=VPS24) | [51652](http://www.ncbi.nlm.nih.gov/entrez/query.fcgi?db=gene&cmd=Retrieve&dopt=Graphics&list_uids=51652) | s28474 | UGAGGGAGUUGUCCAAAGAtt | UCUUUGGACAACUCCCUCAtg |
| NM_001005753 | [VPS24](http://www.ncbi.nlm.nih.gov/entrez/query.fcgi?db=gene&cmd=search&term=VPS24) | [51652](http://www.ncbi.nlm.nih.gov/entrez/query.fcgi?db=gene&cmd=Retrieve&dopt=Graphics&list_uids=51652) | s28475 | GCUGGGAUCAUAGAGGAGAtt | UCUCCUCUAUGAUCCCAGCct |
| NM_013245 | [VPS4A](http://www.ncbi.nlm.nih.gov/entrez/query.fcgi?db=gene&cmd=search&term=VPS4A) | [27183](http://www.ncbi.nlm.nih.gov/entrez/query.fcgi?db=gene&cmd=Retrieve&dopt=Graphics&list_uids=27183) | s25967 | CGGAGUUCUUGGUCCAGAUtt | AUCUGGACCAAGAACUCCGtt |
| NM_013245 | [VPS4A](http://www.ncbi.nlm.nih.gov/entrez/query.fcgi?db=gene&cmd=search&term=VPS4A) | [27183](http://www.ncbi.nlm.nih.gov/entrez/query.fcgi?db=gene&cmd=Retrieve&dopt=Graphics&list_uids=27183) | s25966 | CCUCAGAUCUGAUGUCCAAtt | UUGGACAUCAGAUCUGAGGag |
| NM_013245 | [VPS4A](http://www.ncbi.nlm.nih.gov/entrez/query.fcgi?db=gene&cmd=search&term=VPS4A) | [27183](http://www.ncbi.nlm.nih.gov/entrez/query.fcgi?db=gene&cmd=Retrieve&dopt=Graphics&list_uids=27183) | s25968 | GAAGCUGAAGGAUUAUUUAtt | UAAAUAAUCCUUCAGCUUCtc |
| NM_016485 | [VTA1](http://www.ncbi.nlm.nih.gov/entrez/query.fcgi?db=gene&cmd=search&term=VTA1) | [51534](http://www.ncbi.nlm.nih.gov/entrez/query.fcgi?db=gene&cmd=Retrieve&dopt=Graphics&list_uids=51534) | s28286 | GGCUUAUUACUGUCGUUUAtt | UAAACGACAGUAAUAAGCCac |
| NM_016485 | [VTA1](http://www.ncbi.nlm.nih.gov/entrez/query.fcgi?db=gene&cmd=search&term=VTA1) | [51534](http://www.ncbi.nlm.nih.gov/entrez/query.fcgi?db=gene&cmd=Retrieve&dopt=Graphics&list_uids=51534) | s28287 | GCACAGGUGUAGCAAGUAAtt | UUACUUGCUACACCUGUGCtg |
| NM_016485 | [VTA1](http://www.ncbi.nlm.nih.gov/entrez/query.fcgi?db=gene&cmd=search&term=VTA1) | [51534](http://www.ncbi.nlm.nih.gov/entrez/query.fcgi?db=gene&cmd=Retrieve&dopt=Graphics&list_uids=51534) | s28285 | GAAUGAAGAUCGAUAGUAAtt | UUACUAUCGAUCUUCAUUCca |
| **Exocyst complex** |  |  |  |  |  |
| NM_178237 | [EXOC1](http://www.ncbi.nlm.nih.gov/entrez/query.fcgi?db=gene&cmd=search&term=EXOC1) | [55763](http://www.ncbi.nlm.nih.gov/entrez/query.fcgi?db=gene&cmd=Retrieve&dopt=Graphics&list_uids=55763) | s31452 | GCGAUUCAGUGAUUUGCGAtt | UCGCAAAUCACUGAAUCGCtg |
| NM_178237 | [EXOC1](http://www.ncbi.nlm.nih.gov/entrez/query.fcgi?db=gene&cmd=search&term=EXOC1) | [55763](http://www.ncbi.nlm.nih.gov/entrez/query.fcgi?db=gene&cmd=Retrieve&dopt=Graphics&list_uids=55763) | s31453 | GGACUUGCAGAAUCAAUCUtt | AGAUUGAUUCUGCAAGUCCag |
| NM_178237 | [EXOC1](http://www.ncbi.nlm.nih.gov/entrez/query.fcgi?db=gene&cmd=search&term=EXOC1) | [55763](http://www.ncbi.nlm.nih.gov/entrez/query.fcgi?db=gene&cmd=Retrieve&dopt=Graphics&list_uids=55763) | s31454 | GCAUACACCAAACUUAUCAtt | UGAUAAGUUUGGUGUAUGCtt |
| NM_018303 | [EXOC2](http://www.ncbi.nlm.nih.gov/entrez/query.fcgi?db=gene&cmd=search&term=EXOC2) | [55770](http://www.ncbi.nlm.nih.gov/entrez/query.fcgi?db=gene&cmd=Retrieve&dopt=Graphics&list_uids=55770) | s31470 | GACUUACUCAUGAAUCGUUtt | AACGAUUCAUGAGUAAGUCtt |
| NM_018303 | [EXOC2](http://www.ncbi.nlm.nih.gov/entrez/query.fcgi?db=gene&cmd=search&term=EXOC2) | [55770](http://www.ncbi.nlm.nih.gov/entrez/query.fcgi?db=gene&cmd=Retrieve&dopt=Graphics&list_uids=55770) | s31471 | GCCUGGUAUCUUAUAGAGAtt | UCUCUAUAAGAUACCAGGCtg |
| NM_018303 | [EXOC2](http://www.ncbi.nlm.nih.gov/entrez/query.fcgi?db=gene&cmd=search&term=EXOC2) | [55770](http://www.ncbi.nlm.nih.gov/entrez/query.fcgi?db=gene&cmd=Retrieve&dopt=Graphics&list_uids=55770) | s31472 | GCUCAUGCACAGUUGCAAAtt | UUUGCAACUGUGCAUGAGCtg |
| NM_001013848 | [EXOC6](http://www.ncbi.nlm.nih.gov/entrez/query.fcgi?db=gene&cmd=search&term=EXOC6) | [54536](http://www.ncbi.nlm.nih.gov/entrez/query.fcgi?db=gene&cmd=Retrieve&dopt=Graphics&list_uids=54536) | s29154 | GGUACUAUUCUGCCCUAAAtt | UUUAGGGCAGAAUAGUACCtt |
| NM_001013848 | [EXOC6](http://www.ncbi.nlm.nih.gov/entrez/query.fcgi?db=gene&cmd=search&term=EXOC6) | [54536](http://www.ncbi.nlm.nih.gov/entrez/query.fcgi?db=gene&cmd=Retrieve&dopt=Graphics&list_uids=54536) | s29156 | GAAGAUAUCAUUCGAUGUAtt | UACAUCGAAUGAUAUCUUCtg |
| NM_001013848 | [EXOC6](http://www.ncbi.nlm.nih.gov/entrez/query.fcgi?db=gene&cmd=search&term=EXOC6) | [54536](http://www.ncbi.nlm.nih.gov/entrez/query.fcgi?db=gene&cmd=Retrieve&dopt=Graphics&list_uids=54536) | s29155 | GGUUCUUUGUGGUAGAAGAtt | UCUUCUACCACAAAGAACCct |
| NM_015219 | [EXOC7](http://www.ncbi.nlm.nih.gov/entrez/query.fcgi?db=gene&cmd=search&term=EXOC7) | [23265](http://www.ncbi.nlm.nih.gov/entrez/query.fcgi?db=gene&cmd=Retrieve&dopt=Graphics&list_uids=23265) | s23433 | ACAUCAAGAAUGACCCGGAtt | UCCGGGUCAUUCUUGAUGUtg |
| NM_015219 | [EXOC7](http://www.ncbi.nlm.nih.gov/entrez/query.fcgi?db=gene&cmd=search&term=EXOC7) | [23265](http://www.ncbi.nlm.nih.gov/entrez/query.fcgi?db=gene&cmd=Retrieve&dopt=Graphics&list_uids=23265) | s23432 | AGAAGACCAUUGUCAAGGAtt | UCCUUGACAAUGGUCUUCUgg |
| NM_015219 | [EXOC7](http://www.ncbi.nlm.nih.gov/entrez/query.fcgi?db=gene&cmd=search&term=EXOC7) | [23265](http://www.ncbi.nlm.nih.gov/entrez/query.fcgi?db=gene&cmd=Retrieve&dopt=Graphics&list_uids=23265) | s23431 | CCAAGAACCCGGAGAAGUAtt | UACUUCUCCGGGUUCUUGGtg |
| NM_002881 | [RALB](http://www.ncbi.nlm.nih.gov/entrez/query.fcgi?db=gene&cmd=search&term=RALB) | [5899](http://www.ncbi.nlm.nih.gov/entrez/query.fcgi?db=gene&cmd=Retrieve&dopt=Graphics&list_uids=5899) | s11763 | CGAGAUAACUACUUUCGGAtt | UCCGAAAGUAGUUAUCUCGaa |
| NM_002881 | [RALB](http://www.ncbi.nlm.nih.gov/entrez/query.fcgi?db=gene&cmd=search&term=RALB) | [5899](http://www.ncbi.nlm.nih.gov/entrez/query.fcgi?db=gene&cmd=Retrieve&dopt=Graphics&list_uids=5899) | s11762 | CAUGAAUCCUUUACAGCAAtt | UUGCUGUAAAGGAUUCAUGtt |
| NM_002881 | [RALB](http://www.ncbi.nlm.nih.gov/entrez/query.fcgi?db=gene&cmd=search&term=RALB) | [5899](http://www.ncbi.nlm.nih.gov/entrez/query.fcgi?db=gene&cmd=Retrieve&dopt=Graphics&list_uids=5899) | s11761 | GUGUGAAGGCUGAAGAAGAtt | UCUUCUUCAGCCUUCACACgg |
| **Organelle biogenesis / lysosomes** |  |  |  |  |  |
| NM_032122 | [DTNBP1](http://www.ncbi.nlm.nih.gov/entrez/query.fcgi?db=gene&cmd=search&term=DTNBP1) | [84062](http://www.ncbi.nlm.nih.gov/entrez/query.fcgi?db=gene&cmd=Retrieve&dopt=Graphics&list_uids=84062) | s38426 | GUACUCUGCUGGAUUAGAAtt | UUCUAAUCCAGCAGAGUACtt |
| NM_032122 | [DTNBP1](http://www.ncbi.nlm.nih.gov/entrez/query.fcgi?db=gene&cmd=search&term=DTNBP1) | [84062](http://www.ncbi.nlm.nih.gov/entrez/query.fcgi?db=gene&cmd=Retrieve&dopt=Graphics&list_uids=84062) | s38427 | CAGCAAAUCUGACUCAUUUtt | AAAUGAGUCAGAUUUGCUGtc |
| NM_032122 | [DTNBP1](http://www.ncbi.nlm.nih.gov/entrez/query.fcgi?db=gene&cmd=search&term=DTNBP1) | [84062](http://www.ncbi.nlm.nih.gov/entrez/query.fcgi?db=gene&cmd=Retrieve&dopt=Graphics&list_uids=84062) | s38428 | CAGCUUUAAUCGCAGACUUtt | AAGUCUGCGAUUAAAGCUGgg |
| NM_000195 | [HPS1](http://www.ncbi.nlm.nih.gov/entrez/query.fcgi?db=gene&cmd=search&term=HPS1) | [3257](http://www.ncbi.nlm.nih.gov/entrez/query.fcgi?db=gene&cmd=Retrieve&dopt=Graphics&list_uids=3257) | s194532 | GCUGAAGUUCGGGCAGUCAtt | UGACUGCCCGAACUUCAGCcg |
| NM_000195 | [HPS1](http://www.ncbi.nlm.nih.gov/entrez/query.fcgi?db=gene&cmd=search&term=HPS1) | [3257](http://www.ncbi.nlm.nih.gov/entrez/query.fcgi?db=gene&cmd=Retrieve&dopt=Graphics&list_uids=3257) | s194533 | GGUCCUCUUCUACUGGACAtt | UGUCCAGUAGAAGAGGACCtc |
| NM_000195 | [HPS1](http://www.ncbi.nlm.nih.gov/entrez/query.fcgi?db=gene&cmd=search&term=HPS1) | [3257](http://www.ncbi.nlm.nih.gov/entrez/query.fcgi?db=gene&cmd=Retrieve&dopt=Graphics&list_uids=3257) | s194534 | GCUUCUCCACGGAAAAUGGtt | CCAUUUUCCGUGGAGAAGCag |
| NM_032383 | [HPS3](http://www.ncbi.nlm.nih.gov/entrez/query.fcgi?db=gene&cmd=search&term=HPS3) | [84343](http://www.ncbi.nlm.nih.gov/entrez/query.fcgi?db=gene&cmd=Retrieve&dopt=Graphics&list_uids=84343) | s38960 | GACUAUAGCAAUACCUAUAtt | UAUAGGUAUUGCUAUAGUCta |
| NM_032383 | HPS3 | 84343 | s38961 | CGAACAUCGGAGGAUCUGAtt | UCAGAUCCUCCGAUGUUCGca |
| NM_032383 | HPS3 | 84343 | s38959 | CAGUCAUUCCAUAUGCUAAtt | UUAGCAUAUGGAAUGACUGcc |
| NM_022081 | HPS4 | 89781 | s40114 | CUCCUACUCUUGUUCGUCUtt | AGACGAACAAGAGUAGGAGga |
| NM_022081 | HPS4 | 89781 | s40116 | CCUCAGUUCUAUUACGUUAtt | UAACGUAAUAGAACUGAGGtt |
| NM_022081 | HPS4 | 89781 | s40115 | GCGGUUUCUGGAUCAGCUAtt | UAGCUGAUCCAGAAACCGCtt |
| NM_181508 | HPS5 | 11234 | s22172 | GCUACCCGGUUGUAUGAAAtt | UUUCAUACAACCGGGUAGCat |
| NM_181508 | HPS5 | 11234 | s22173 | GGGACUCUUUAGGCAAUGAtt | UCAUUGCCUAAAGAGUCCCtt |
| NM_181508 | HPS5 | 11234 | s22174 | GGUUAAAGUUACUAGAUGAtt | UCAUCUAGUAACUUUAACCtt |
| NM_024747 | HPS6 | 79803 | s36367 | GCUGAGGACUGAGUUGAUAtt | UAUCAACUCAGUCCUCAGCag |
| NM_024747 | [HPS6](http://www.ncbi.nlm.nih.gov/entrez/query.fcgi?db=gene&cmd=search&term=HPS6) | [79803](http://www.ncbi.nlm.nih.gov/entrez/query.fcgi?db=gene&cmd=Retrieve&dopt=Graphics&list_uids=79803) | s36368 | GCUCAACACCGUUUUCCAAtt | UUGGAAAACGGUGUUGAGCtg |
| NM_024747 | [HPS6](http://www.ncbi.nlm.nih.gov/entrez/query.fcgi?db=gene&cmd=search&term=HPS6) | [79803](http://www.ncbi.nlm.nih.gov/entrez/query.fcgi?db=gene&cmd=Retrieve&dopt=Graphics&list_uids=79803) | s36366 | GGAAGGUCCUAAGUACAGAtt | UCUGUACUUAGGACCUUCCtc |
| NM_000081 | [LYST](http://www.ncbi.nlm.nih.gov/entrez/query.fcgi?db=gene&cmd=search&term=LYST) | [1130](http://www.ncbi.nlm.nih.gov/entrez/query.fcgi?db=gene&cmd=Retrieve&dopt=Graphics&list_uids=1130) | s3030 | GCAUUAGCACUGCGAGUUAtt | UAACUCGCAGUGCUAAUGCtt |
| NM_000081 | [LYST](http://www.ncbi.nlm.nih.gov/entrez/query.fcgi?db=gene&cmd=search&term=LYST) | [1130](http://www.ncbi.nlm.nih.gov/entrez/query.fcgi?db=gene&cmd=Retrieve&dopt=Graphics&list_uids=1130) | s3029 | GCAACGGUGUUUCAUCACAtt | UGUGAUGAAACACCGUUGCtt |
| NM_000081 | [LYST](http://www.ncbi.nlm.nih.gov/entrez/query.fcgi?db=gene&cmd=search&term=LYST) | [1130](http://www.ncbi.nlm.nih.gov/entrez/query.fcgi?db=gene&cmd=Retrieve&dopt=Graphics&list_uids=1130) | s3028 | CCAAAUGACUUACUCGAAAtt | UUUCGAGUAAGUCAUUUGGac |
| NM_005817 | [M6PRBP1](http://www.ncbi.nlm.nih.gov/entrez/query.fcgi?db=gene&cmd=search&term=M6PRBP1) | [10226](http://www.ncbi.nlm.nih.gov/entrez/query.fcgi?db=gene&cmd=Retrieve&dopt=Graphics&list_uids=10226) | s19952 | CCCUAUCAGUCAGUAGAGAtt | UCUCUACUGACUGAUAGGGtg |
| NM_005817 | [M6PRBP1](http://www.ncbi.nlm.nih.gov/entrez/query.fcgi?db=gene&cmd=search&term=M6PRBP1) | [10226](http://www.ncbi.nlm.nih.gov/entrez/query.fcgi?db=gene&cmd=Retrieve&dopt=Graphics&list_uids=10226) | s225424 | GAAUAUCAGCCUGUAGCUAtt | UAGCUACAGGCUGAUAUUCag |
| NM_005817 | [M6PRBP1](http://www.ncbi.nlm.nih.gov/entrez/query.fcgi?db=gene&cmd=search&term=M6PRBP1) | [10226](http://www.ncbi.nlm.nih.gov/entrez/query.fcgi?db=gene&cmd=Retrieve&dopt=Graphics&list_uids=10226) | s19954 | GCCUGAUGGAAACUGUCAAtt | UUGACAGUUUCCAUCAGGCtt |
| NM_000543 | [SMPD1](http://www.ncbi.nlm.nih.gov/entrez/query.fcgi?db=gene&cmd=search&term=SMPD1) | [6609](http://www.ncbi.nlm.nih.gov/entrez/query.fcgi?db=gene&cmd=Retrieve&dopt=Graphics&list_uids=6609) | s13169 | CUACCUACAUCGGCCUUAAtt | UUAAGGCCGAUGUAGGUAGtt |
| NM_000543 | [SMPD1](http://www.ncbi.nlm.nih.gov/entrez/query.fcgi?db=gene&cmd=search&term=SMPD1) | [6609](http://www.ncbi.nlm.nih.gov/entrez/query.fcgi?db=gene&cmd=Retrieve&dopt=Graphics&list_uids=6609) | s13168 | GCACACCUGUCAAUAGCUUtt | AAGCUAUUGACAGGUGUGCtt |
| NM_000543 | [SMPD1](http://www.ncbi.nlm.nih.gov/entrez/query.fcgi?db=gene&cmd=search&term=SMPD1) | [6609](http://www.ncbi.nlm.nih.gov/entrez/query.fcgi?db=gene&cmd=Retrieve&dopt=Graphics&list_uids=6609) | s13167 | UCACAGCACUUGUGAGGAAtt | UUCCUCACAAGUGCUGUGAcg |
| **Cell proliferation cell cycle** |  |  |  |  |  |
| NM_000077 | [CDKN2A](http://www.ncbi.nlm.nih.gov/entrez/query.fcgi?db=gene&cmd=search&term=CDKN2A) | [1029](http://www.ncbi.nlm.nih.gov/entrez/query.fcgi?db=gene&cmd=Retrieve&dopt=Graphics&list_uids=1029) | s218 | UGUCCUGCCUUUUAACGUAtt | UACGUUAAAAGGCAGGACAtt |
| NM_000077 | [CDKN2A](http://www.ncbi.nlm.nih.gov/entrez/query.fcgi?db=gene&cmd=search&term=CDKN2A) | [1029](http://www.ncbi.nlm.nih.gov/entrez/query.fcgi?db=gene&cmd=Retrieve&dopt=Graphics&list_uids=1029) | s216 | CUACCGUAAAUGUCCAUUUtt | AAAUGGACAUUUACGGUAGtg |
| NM_000077 | [CDKN2A](http://www.ncbi.nlm.nih.gov/entrez/query.fcgi?db=gene&cmd=search&term=CDKN2A) | [1029](http://www.ncbi.nlm.nih.gov/entrez/query.fcgi?db=gene&cmd=Retrieve&dopt=Graphics&list_uids=1029) | s217 | CCGUAAAUGUCCAUUUAUAtt | UAUAAAUGGACAUUUACGGta |
| **Heat Schock proteins** |  |  |  |  |  |
| NM_006597 | [HSPA8](http://www.ncbi.nlm.nih.gov/entrez/query.fcgi?db=gene&cmd=search&term=HSPA8) | [3312](http://www.ncbi.nlm.nih.gov/entrez/query.fcgi?db=gene&cmd=Retrieve&dopt=Graphics&list_uids=3312) | s6986 | GCUUACGGCUUAGACAAAAtt | UUUUGUCUAAGCCGUAAGCaa |
| NM_006597 | [HSPA8](http://www.ncbi.nlm.nih.gov/entrez/query.fcgi?db=gene&cmd=search&term=HSPA8) | [3312](http://www.ncbi.nlm.nih.gov/entrez/query.fcgi?db=gene&cmd=Retrieve&dopt=Graphics&list_uids=3312) | s6987 | CCUUUAUGGUGGUGAAUGAtt | UCAUUCACCACCAUAAAGGgc |
| NM_006597 | [HSPA8](http://www.ncbi.nlm.nih.gov/entrez/query.fcgi?db=gene&cmd=search&term=HSPA8) | [3312](http://www.ncbi.nlm.nih.gov/entrez/query.fcgi?db=gene&cmd=Retrieve&dopt=Graphics&list_uids=3312) | s6985 | GAGUUUAAGCGCAAGCAUAtt | UAUGCUUGCGCUUAAACUCag |
| **Adhesion receptor** |  |  |  |  |  |
| NM_002204 | [ITGA3](http://www.ncbi.nlm.nih.gov/entrez/query.fcgi?db=gene&cmd=search&term=ITGA3) | [3675](http://www.ncbi.nlm.nih.gov/entrez/query.fcgi?db=gene&cmd=Retrieve&dopt=Graphics&list_uids=3675) | s7543 | GGACUUAUCUGAGUAUAGUtt | ACUAUACUCAGAUAAGUCCca |
| NM_002204 | [ITGA3](http://www.ncbi.nlm.nih.gov/entrez/query.fcgi?db=gene&cmd=search&term=ITGA3) | [3675](http://www.ncbi.nlm.nih.gov/entrez/query.fcgi?db=gene&cmd=Retrieve&dopt=Graphics&list_uids=3675) | s7542 | CAACGUGACUGUGAAGGCAtt | UGCCUUCACAGUCACGUUGgt |
| NM_002204 | [ITGA3](http://www.ncbi.nlm.nih.gov/entrez/query.fcgi?db=gene&cmd=search&term=ITGA3) | [3675](http://www.ncbi.nlm.nih.gov/entrez/query.fcgi?db=gene&cmd=Retrieve&dopt=Graphics&list_uids=3675) | s7541 | GUAAAUCACCGGCUACAAAtt | UUUGUAGCCGGUGAUUUACca |
| **Caspases** |  |  |  |  |  |
| NM_001223 | [CASP1](http://www.ncbi.nlm.nih.gov/entrez/query.fcgi?db=gene&cmd=search&term=CASP1) | [834](http://www.ncbi.nlm.nih.gov/entrez/query.fcgi?db=gene&cmd=Retrieve&dopt=Graphics&list_uids=834) | s2407 | GGAAGACUCAUUGAACAUAtt | UAUGUUCAAUGAGUCUUCCaa |
| NM_001223 | [CASP1](http://www.ncbi.nlm.nih.gov/entrez/query.fcgi?db=gene&cmd=search&term=CASP1) | [834](http://www.ncbi.nlm.nih.gov/entrez/query.fcgi?db=gene&cmd=Retrieve&dopt=Graphics&list_uids=834) | s2409 | UGGAGGAAAUUUUCCGCAAtt | UUGCGGAAAAUUUCCUCCAca |
| NM_001223 | [CASP1](http://www.ncbi.nlm.nih.gov/entrez/query.fcgi?db=gene&cmd=search&term=CASP1) | [834](http://www.ncbi.nlm.nih.gov/entrez/query.fcgi?db=gene&cmd=Retrieve&dopt=Graphics&list_uids=834) | s2408 | CCACUGAAAGAGUGACUUUtt | AAAGUCACUCUUUCAGUGGtg |
| NM_033306 | [CASP4](http://www.ncbi.nlm.nih.gov/entrez/query.fcgi?db=gene&cmd=search&term=CASP4) | [837](http://www.ncbi.nlm.nih.gov/entrez/query.fcgi?db=gene&cmd=Retrieve&dopt=Graphics&list_uids=837) | s2414 | GAGACUAUGUAAAGAAAGAtt | UCUUUCUUUACAUAGUCUCag |
| NM_033306 | [CASP4](http://www.ncbi.nlm.nih.gov/entrez/query.fcgi?db=gene&cmd=search&term=CASP4) | [837](http://www.ncbi.nlm.nih.gov/entrez/query.fcgi?db=gene&cmd=Retrieve&dopt=Graphics&list_uids=837) | s2413 | GGAUGAAGGAGCUACUUGAtt | UCAAGUAGCUCCUUCAUCCct |
| NM_033306 | [CASP4](http://www.ncbi.nlm.nih.gov/entrez/query.fcgi?db=gene&cmd=search&term=CASP4) | [837](http://www.ncbi.nlm.nih.gov/entrez/query.fcgi?db=gene&cmd=Retrieve&dopt=Graphics&list_uids=837) | s2415 | GGGUCUGGACUAUAGUGUAtt | UACACUAUAGUCCAGACCCtc |
| NM_004347 | [CASP5](http://www.ncbi.nlm.nih.gov/entrez/query.fcgi?db=gene&cmd=search&term=CASP5) | [838](http://www.ncbi.nlm.nih.gov/entrez/query.fcgi?db=gene&cmd=Retrieve&dopt=Graphics&list_uids=838) | s229323 | CCUAGUACCUAAUACGGAUtt | AUCCGUAUUAGGUACUAGGgt |
| NM_004347 | [CASP5](http://www.ncbi.nlm.nih.gov/entrez/query.fcgi?db=gene&cmd=search&term=CASP5) | [838](http://www.ncbi.nlm.nih.gov/entrez/query.fcgi?db=gene&cmd=Retrieve&dopt=Graphics&list_uids=838) | s2417 | GAACUGCGCAUAAAAAGAAtt | UUCUUUUUAUGCGCAGUUCcg |
| NM_004347 | [CASP5](http://www.ncbi.nlm.nih.gov/entrez/query.fcgi?db=gene&cmd=search&term=CASP5) | [838](http://www.ncbi.nlm.nih.gov/entrez/query.fcgi?db=gene&cmd=Retrieve&dopt=Graphics&list_uids=838) | s229324 | ACGAUGUUCUGACAUUGAAtt | UUCAAUGUCAGAACAUCGUgt |
| **SEPTINS** |  |  |  |  |  |
| NM_006155 | [SEPT2](http://www.ncbi.nlm.nih.gov/entrez/query.fcgi?db=gene&cmd=search&term=SEPT2) (siRNA 3) | [4735](http://www.ncbi.nlm.nih.gov/entrez/query.fcgi?db=gene&cmd=Retrieve&dopt=Graphics&list_uids=4735) | s9420 | CAAUCAAGUUCACCGAAAAtt | UUUUCGGUGAACUUGAUUGgg |
| NM_006155 | [SEPT2](http://www.ncbi.nlm.nih.gov/entrez/query.fcgi?db=gene&cmd=search&term=SEPT2)  (siRNA 1) | [4735](http://www.ncbi.nlm.nih.gov/entrez/query.fcgi?db=gene&cmd=Retrieve&dopt=Graphics&list_uids=4735) | s9418 | GCCUAUUCCUAACUGAUCUtt | AGAUCAGUUAGGAAUAGGCtg |
| NM_006155 | [SEPT2](http://www.ncbi.nlm.nih.gov/entrez/query.fcgi?db=gene&cmd=search&term=SEPT2)  (siRNA 2) | [4735](http://www.ncbi.nlm.nih.gov/entrez/query.fcgi?db=gene&cmd=Retrieve&dopt=Graphics&list_uids=4735) | s9419 | GAAAAUCGACUCUCAUAAAtt | UUUAUGAGAGUCGAUUUUCct |
| NM_015129 | [SEPT6](http://www.ncbi.nlm.nih.gov/entrez/query.fcgi?db=gene&cmd=search&term=SEPT6) (siRNA 1) | [23157](http://www.ncbi.nlm.nih.gov/entrez/query.fcgi?db=gene&cmd=Retrieve&dopt=Graphics&list_uids=23157) | s250 | CGAAGAGUGAGCUAACAAAtt | UUUGUUAGCUCACUCUUCGaa |
| NM_015129 | [SEPT6](http://www.ncbi.nlm.nih.gov/entrez/query.fcgi?db=gene&cmd=search&term=SEPT6)  (siRNA 2) | [23157](http://www.ncbi.nlm.nih.gov/entrez/query.fcgi?db=gene&cmd=Retrieve&dopt=Graphics&list_uids=23157) | s251 | GUUUGACCGUCUGAAGAAAtt | UUUCUUCAGACGGUCAAACtt |
| NM_015129 | [SEPT6](http://www.ncbi.nlm.nih.gov/entrez/query.fcgi?db=gene&cmd=search&term=SEPT6)  (siRNA 3) | [23157](http://www.ncbi.nlm.nih.gov/entrez/query.fcgi?db=gene&cmd=Retrieve&dopt=Graphics&list_uids=23157) | s252 | GCACUGUGCAGGUUGAAAAtt | UUUUCAACCUGCACAGUGCcc |
| NM_001011553 | [SEPT7](http://www.ncbi.nlm.nih.gov/entrez/query.fcgi?db=gene&cmd=search&term=SEPT7) (siRNA 1) | [989](http://www.ncbi.nlm.nih.gov/entrez/query.fcgi?db=gene&cmd=Retrieve&dopt=Graphics&list_uids=989) | s2741 | GAAGGGAGCAUGUAGCUAAtt | UUAGCUACAUGCUCCCUUCtt |
| NM_001011553 | [SEPT7](http://www.ncbi.nlm.nih.gov/entrez/query.fcgi?db=gene&cmd=search&term=SEPT7)  (siRNA 2) | [989](http://www.ncbi.nlm.nih.gov/entrez/query.fcgi?db=gene&cmd=Retrieve&dopt=Graphics&list_uids=989) | s2742 | GGUGAAGAGAGGUUUUGAAtt | UUCAAAACCUCUCUUCACCga |
| NM_001011553 | [SEPT7](http://www.ncbi.nlm.nih.gov/entrez/query.fcgi?db=gene&cmd=search&term=SEPT7)  (siRNA 3) | [989](http://www.ncbi.nlm.nih.gov/entrez/query.fcgi?db=gene&cmd=Retrieve&dopt=Graphics&list_uids=989) | s2743 | GCCUGUUAUCGACUACAUUtt | AAUGUAGUCGAUAACAGGCtg |
| NM_006640 | [SEPT9](http://www.ncbi.nlm.nih.gov/entrez/query.fcgi?db=gene&cmd=search&term=SEPT9) (siRNA 2) | [10801](http://www.ncbi.nlm.nih.gov/entrez/query.fcgi?db=gene&cmd=Retrieve&dopt=Graphics&list_uids=10801) | s21223 | CAAUGACCAGUACGAGAAAtt | UUUCUCGUACUGGUCAUUGat |
| NM_006640 | [SEPT9](http://www.ncbi.nlm.nih.gov/entrez/query.fcgi?db=gene&cmd=search&term=SEPT9)  (siRNA 3) | [10801](http://www.ncbi.nlm.nih.gov/entrez/query.fcgi?db=gene&cmd=Retrieve&dopt=Graphics&list_uids=10801) | s21224 | CGGAUGAAGCUGACAGUGAtt | UCACUGUCAGCUUCAUCCGga |
| NM_006640 | SEPT9  (siRNA 1) | [10801](http://www.ncbi.nlm.nih.gov/entrez/query.fcgi?db=gene&cmd=Retrieve&dopt=Graphics&list_uids=10801) | s21222 | CGCACGAUAUUGAGGAGAAtt | UUCUCCUCAAUAUCGUGCGtg |
