## Supplemental Table 2 for "The Septin Cytoskeleton is Required for Plasma Membrane Repair"

**Supplemental Table 2:** Cells were enumerated pre- and post-kinetic in each well of the 96 well plates based on fluorescence microscopy imaging of H2BGFP-expressing HeLa cells and automated image analysis, as we previously described [32]. Cell numbers are the average of four full wells in each experimental condition and presented data are post-kinetic. Experimental samples are the average counts of cells targeted with specific siRNAs. Controls 1 to 3 are the counts of control cells plated at 100% (Control 1), 90% (Control 2), and 80% (Control 3) cell density and treated with non-targeting siRNA. In yellow are experimental conditions which were excluded due to the ratio of experimental sample to control outside of a 15% range to control cells.

| **Gene name** | **Experimental Sample** | **Control 1** | **Control 2** | **Control 3** | **Ratio to control 1** | **Ratio to control 2** | **Ratio to control 3** |
| --- | --- | --- | --- | --- | --- | --- | --- |
| ABI1 | 70764.00 | 66605.75 | 62758.00 | 50664.00 | 1.06 | 1.13 | 1.40 |
| ANXA1 | 72216.00 | 62653.75 | 57675.00 | 45848.25 | 1.15 | 1.25 | 1.58 |
| ANXA11 | 52925.50 | 63228.00 | 59137.00 | 45649.50 | 0.84 | 0.89 | 1.16 |
| ANXA2 | 57248.75 | 63228.00 | 59137.00 | 45649.50 | 0.91 | 0.97 | 1.25 |
| ANXA3 | 59129.25 | 63228.00 | 59137.00 | 45649.50 | 0.94 | 1.00 | 1.30 |
| ANXA4 | 55963.50 | 63228.00 | 59137.00 | 45649.50 | 0.89 | 0.95 | 1.23 |
| ANXA5 | 56709.00 | 63228.00 | 59137.00 | 45649.50 | 0.90 | 0.96 | 1.24 |
| ANXA6 | 58208.25 | 63228.00 | 59137.00 | 45649.50 | 0.92 | 0.98 | 1.28 |
| ANXA7 | 66643.00 | 63228.00 | 59137.00 | 45649.50 | 1.05 | 1.13 | 1.46 |
| AP1AR | 22220.25 | 42578.00 | 35172.75 | 24136.50 | 0.52 | 0.63 | 0.92 |
| AP1B1 | 49053.25 | 52565.00 | 41736.50 | 29078.25 | 0.93 | 1.18 | 1.69 |
| AP1G1 | 48396.50 | 52565.00 | 41736.50 | 29078.25 | 0.92 | 1.16 | 1.66 |
| AP1G2 | 54660.00 | 55655.00 | 42279.00 | 34164.00 | 0.98 | 1.29 | 1.60 |
| AP1GBP1 | 35386.25 | 42878.00 | 32106.00 | 21444.75 | 0.83 | 1.10 | 1.65 |
| AP1M1 | 55210.75 | 57920.00 | 43760.50 | 36524.75 | 0.95 | 1.26 | 1.51 |
| AP1M2 | 58143.50 | 57514.25 | 46520.25 | 32816.25 | 1.01 | 1.25 | 1.77 |
| AP1S1 | 58791.75 | 56792.75 | 44993.00 | 34582.00 | 1.04 | 1.31 | 1.70 |
| AP1S2 | 54855.75 | 55655.00 | 42279.00 | 34164.00 | 0.99 | 1.30 | 1.61 |
| AP1S3 | 42610.50 | 49892.75 | 38856.25 | 25878.75 | 0.85 | 1.10 | 1.65 |
| AP2A1 | 43123.75 | 52565.00 | 41736.50 | 29078.25 | 0.82 | 1.03 | 1.48 |
| AP2A2 | 48830.25 | 52565.00 | 41736.50 | 29078.25 | 0.93 | 1.17 | 1.68 |
| AP2B1 | 37666.00 | 52565.00 | 41736.50 | 29078.25 | 0.72 | 0.90 | 1.30 |
| AP2M1 | 49775.75 | 56792.75 | 44993.00 | 34582.00 | 0.88 | 1.11 | 1.44 |
| AP2S1 | 48398.75 | 56792.75 | 44993.00 | 34582.00 | 0.85 | 1.08 | 1.40 |
| AP3B2 | 53086.50 | 53274.50 | 38684.00 | 32412.00 | 1.00 | 1.37 | 1.64 |
| AP3D1 | 46804.25 | 57920.00 | 43760.50 | 36524.75 | 0.81 | 1.07 | 1.28 |
| AP3M1 | 45365.25 | 43884.00 | 36079.25 | 25806.00 | 1.03 | 1.26 | 1.76 |
| AP3M2 | 66676.75 | 57994.25 | 44638.50 | 35343.25 | 1.15 | 1.49 | 1.89 |
| AP3S1 | 56217.00 | 56792.75 | 44993.00 | 34582.00 | 0.99 | 1.25 | 1.63 |
| AP3S2 | 58271.75 | 57994.25 | 44638.50 | 35343.25 | 1.00 | 1.31 | 1.65 |
| AP4B1 | 55565.50 | 57994.25 | 44638.50 | 35343.25 | 0.96 | 1.24 | 1.57 |
| AP4E1 | 33776.50 | 42878.00 | 32106.00 | 21444.75 | 0.79 | 1.05 | 1.58 |
| AP4M1 | 57333.50 | 57920.00 | 43760.50 | 36524.75 | 0.99 | 1.31 | 1.57 |
| AP4S1 | 62732.50 | 57994.25 | 44638.50 | 35343.25 | 1.08 | 1.41 | 1.77 |
| ARF1 | 45798.00 | 52565.00 | 41736.50 | 29078.25 | 0.87 | 1.10 | 1.57 |
| ARHGAP10 | 51689.50 | 47050.25 | 38202.00 | 25381.50 | 1.10 | 1.35 | 2.04 |
| ARHGAP32 | 57833.00 | 57514.25 | 46520.25 | 32816.25 | 1.01 | 1.24 | 1.76 |
| CAPN1 | 52947.25 | 63925.50 | 60479.50 | 46606.00 | 0.83 | 0.88 | 1.14 |
| CAPN2 | 54240.00 | 63925.50 | 60479.50 | 46606.00 | 0.85 | 0.90 | 1.16 |
| CAPN3 | 48403.00 | 63925.50 | 60479.50 | 46606.00 | 0.76 | 0.80 | 1.04 |
| CASP1 | 63263.50 | 66605.75 | 62758.00 | 50664.00 | 0.95 | 1.01 | 1.25 |
| CASP4 | 63453.25 | 66605.75 | 62758.00 | 50664.00 | 0.95 | 1.01 | 1.25 |
| CASP5 | 66277.25 | 66605.75 | 62758.00 | 50664.00 | 1.00 | 1.06 | 1.31 |
| CAV1 | 42635.50 | 49721.75 | 44911.50 | 34100.75 | 0.86 | 0.95 | 1.25 |
| CAV2 | 52915.00 | 49721.75 | 44911.50 | 34100.75 | 1.06 | 1.18 | 1.55 |
| CAV3 | 49157.00 | 49721.75 | 44911.50 | 34100.75 | 0.99 | 1.09 | 1.44 |
| CDKN2A | 57647.00 | 49721.75 | 44911.50 | 34100.75 | 1.16 | 1.28 | 1.69 |
| CHMP1A | 51091.75 | 51691.75 | 40312.00 | 28852.25 | 0.99 | 1.27 | 1.77 |
| CHMP2A | 40361.25 | 51691.75 | 40312.00 | 28852.25 | 0.78 | 1.00 | 1.40 |
| CHMP4A | 42032.00 | 51175.25 | 41281.00 | 29996.75 | 0.82 | 1.02 | 1.40 |
| CHMP5 | 49792.25 | 51175.25 | 41281.00 | 29996.75 | 0.97 | 1.21 | 1.66 |
| CHMP6 | 32986.75 | 51175.25 | 41281.00 | 29996.75 | 0.64 | 0.80 | 1.10 |
| CLTA | 36398.50 | 56792.75 | 44993.00 | 34582.00 | 0.64 | 0.81 | 1.05 |
| CLTB | 30742.25 | 56792.75 | 44993.00 | 34582.00 | 0.54 | 0.68 | 0.89 |
| CLTC | 38204.50 | 56792.75 | 44993.00 | 34582.00 | 0.67 | 0.85 | 1.10 |
| COPA | 4677.25 | 56792.75 | 44993.00 | 34582.00 | 0.08 | 0.10 | 0.14 |
| COPB1 | 3921.00 | 56792.75 | 44993.00 | 34582.00 | 0.07 | 0.09 | 0.11 |
| COPB2 | 6562.00 | 57920.00 | 43760.50 | 36524.75 | 0.11 | 0.15 | 0.18 |
| COPE | 46532.25 | 42878.00 | 32106.00 | 21444.75 | 1.09 | 1.45 | 2.17 |
| COPG1 | 3717.50 | 42878.00 | 32106.00 | 21444.75 | 0.09 | 0.12 | 0.17 |
| COPG2 | 34455.75 | 43884.00 | 36079.25 | 25806.00 | 0.79 | 0.96 | 1.34 |
| COPZ1 | 5138.75 | 42878.00 | 32106.00 | 21444.75 | 0.12 | 0.16 | 0.24 |
| COPZ2 | 44888.75 | 42578.00 | 35172.75 | 24136.50 | 1.05 | 1.28 | 1.86 |
| DNM1 | 43126.75 | 56207.75 | 46626.75 | 36341.50 | 0.77 | 0.92 | 1.19 |
| DNM2 | 55099.00 | 56207.75 | 46626.75 | 36341.50 | 0.98 | 1.18 | 1.52 |
| DNM3 | 33060.50 | 43884.00 | 36079.25 | 25806.00 | 0.75 | 0.92 | 1.28 |
| DTNBP1 | 36182.75 | 47050.25 | 38202.00 | 25381.50 | 0.77 | 0.95 | 1.43 |
| ENG | 41537.00 | 56207.75 | 46626.75 | 36341.50 | 0.74 | 0.89 | 1.14 |
| EXOC1 | 39509.00 | 50124.75 | 38891.25 | 27570.50 | 0.79 | 1.02 | 1.43 |
| EXOC2 | 40777.75 | 50124.75 | 38891.25 | 27570.50 | 0.81 | 1.05 | 1.48 |
| EXOC6 | 76346.00 | 66605.75 | 62758.00 | 50664.00 | 1.15 | 1.22 | 1.51 |
| EXOC7 | 60498.50 | 66605.75 | 62758.00 | 50664.00 | 0.91 | 0.96 | 1.19 |
| FAM62B | 41668.00 | 42578.00 | 35172.75 | 24136.50 | 0.98 | 1.18 | 1.73 |
| FAM62C | 32040.75 | 47050.25 | 38202.00 | 25381.50 | 0.68 | 0.84 | 1.26 |
| FLOT1 | 49694.50 | 51691.75 | 40312.00 | 28852.25 | 0.96 | 1.23 | 1.72 |
| FLOT2 | 51340.50 | 51691.75 | 40312.00 | 28852.25 | 0.99 | 1.27 | 1.78 |
| GBF1 | 43678.25 | 55655.00 | 42279.00 | 34164.00 | 0.78 | 1.03 | 1.28 |
| HGS | 53314.50 | 50124.75 | 38891.25 | 27570.50 | 1.06 | 1.37 | 1.93 |
| HPS1 | 59815.25 | 56207.75 | 46626.75 | 36341.50 | 1.06 | 1.28 | 1.65 |
| HPS3 | 38763.25 | 49892.75 | 38856.25 | 25878.75 | 0.78 | 1.00 | 1.50 |
| HPS4 | 56717.00 | 49892.75 | 38856.25 | 25878.75 | 1.14 | 1.46 | 2.19 |
| HPS5 | 36842.75 | 42878.00 | 32106.00 | 21444.75 | 0.86 | 1.15 | 1.72 |
| HPS6 | 49820.00 | 47050.25 | 38202.00 | 25381.50 | 1.06 | 1.30 | 1.96 |
| HSPA8 | 37837.00 | 56207.75 | 46626.75 | 36341.50 | 0.67 | 0.81 | 1.04 |
| HUWE1 | 65393.25 | 57514.25 | 46520.25 | 32816.25 | 1.14 | 1.41 | 1.99 |
| IQSEC1 | 39731.75 | 57514.25 | 46520.25 | 32816.25 | 0.69 | 0.85 | 1.21 |
| IST1 | 32171.00 | 62653.75 | 57675.00 | 45848.25 | 0.51 | 0.56 | 0.70 |
| ITGA3 | 37957.50 | 56207.75 | 46626.75 | 36341.50 | 0.68 | 0.81 | 1.04 |
| LMAN1 | 41337.00 | 56207.75 | 46626.75 | 36341.50 | 0.74 | 0.89 | 1.14 |
| LYST | 59215.75 | 49721.75 | 44911.50 | 34100.75 | 1.19 | 1.32 | 1.74 |
| M6PRBP1 | 50137.00 | 57994.25 | 44638.50 | 35343.25 | 0.86 | 1.12 | 1.42 |
| MOCK | 58963.50 | 47314.50 | 41336.50 | 31515.75 | 1.25 | 1.43 | 1.87 |
| PDCD6IP | 48937.50 | 51691.75 | 40312.00 | 28852.25 | 0.95 | 1.21 | 1.70 |
| PREB | 48874.75 | 57514.25 | 46520.25 | 32816.25 | 0.85 | 1.05 | 1.49 |
| PTGES2 | 32883.25 | 47050.25 | 38202.00 | 25381.50 | 0.70 | 0.86 | 1.30 |
| RAB10 | 65858.75 | 61917.75 | 58930.00 | 48092.75 | 1.06 | 1.12 | 1.37 |
| RAB11A | 61056.00 | 60878.50 | 53757.00 | 41389.00 | 1.00 | 1.14 | 1.48 |
| RAB11B | 45236.25 | 60878.50 | 53757.00 | 41389.00 | 0.74 | 0.84 | 1.09 |
| RAB12 | 44255.25 | 45808.00 | 40718.50 | 27916.75 | 0.97 | 1.09 | 1.59 |
| RAB13 | 48530.75 | 60878.50 | 53757.00 | 41389.00 | 0.80 | 0.90 | 1.17 |
| RAB14 | 73877.00 | 66480.50 | 59920.00 | 48446.00 | 1.11 | 1.23 | 1.52 |
| RAB15 | 34751.75 | 52352.25 | 44761.25 | 31201.25 | 0.66 | 0.78 | 1.11 |
| RAB17 | 58758.25 | 65767.75 | 60563.00 | 45185.75 | 0.89 | 0.97 | 1.30 |
| RAB18 | 54308.75 | 64701.25 | 54309.25 | 46674.00 | 0.84 | 1.00 | 1.16 |
| RAB19 | 60179.75 | 57103.50 | 44491.00 | 33130.75 | 1.05 | 1.35 | 1.82 |
| RAB1A | 57550.25 | 57103.50 | 44491.00 | 33130.75 | 1.01 | 1.29 | 1.74 |
| RAB1B | 62437.00 | 65767.75 | 60563.00 | 45185.75 | 0.95 | 1.03 | 1.38 |
| RAB20 | 37240.25 | 65767.75 | 60563.00 | 45185.75 | 0.57 | 0.61 | 0.82 |
| RAB21 | 62376.50 | 64701.25 | 54309.25 | 46674.00 | 0.96 | 1.15 | 1.34 |
| RAB22A | 74857.00 | 65767.75 | 60563.00 | 45185.75 | 1.14 | 1.24 | 1.66 |
| RAB23 | 68000.00 | 66480.50 | 59920.00 | 48446.00 | 1.02 | 1.13 | 1.40 |
| RAB24 | 64457.25 | 66480.50 | 59920.00 | 48446.00 | 0.97 | 1.08 | 1.33 |
| RAB25 | 50509.00 | 65767.75 | 60563.00 | 45185.75 | 0.77 | 0.83 | 1.12 |
| RAB26 | 53849.00 | 64701.25 | 54309.25 | 46674.00 | 0.83 | 0.99 | 1.15 |
| RAB27A | 61918.50 | 60878.50 | 53757.00 | 41389.00 | 1.02 | 1.15 | 1.50 |
| RAB27B | 62867.00 | 60878.50 | 53757.00 | 41389.00 | 1.03 | 1.17 | 1.52 |
| RAB28 | 45958.25 | 61917.75 | 58930.00 | 48092.75 | 0.74 | 0.78 | 0.96 |
| RAB2A | 56695.00 | 57103.50 | 44491.00 | 33130.75 | 0.99 | 1.27 | 1.71 |
| RAB2B | 42268.50 | 45808.00 | 40718.50 | 27916.75 | 0.92 | 1.04 | 1.51 |
| RAB30 | 60998.50 | 66480.50 | 59920.00 | 48446.00 | 0.92 | 1.02 | 1.26 |
| RAB31 | 36568.50 | 61917.75 | 58930.00 | 48092.75 | 0.59 | 0.62 | 0.76 |
| RAB32 | 35772.75 | 61917.75 | 58930.00 | 48092.75 | 0.58 | 0.61 | 0.74 |
| RAB33B | 65686.00 | 65767.75 | 60563.00 | 45185.75 | 1.00 | 1.08 | 1.45 |
| RAB34 | 75312.25 | 65767.75 | 60563.00 | 45185.75 | 1.15 | 1.24 | 1.67 |
| RAB35 | 62890.25 | 61917.75 | 58930.00 | 48092.75 | 1.02 | 1.07 | 1.31 |
| RAB36 | 59004.00 | 61917.75 | 58930.00 | 48092.75 | 0.95 | 1.00 | 1.23 |
| RAB37 | 68635.25 | 64701.25 | 54309.25 | 46674.00 | 1.06 | 1.26 | 1.47 |
| RAB38 | 37015.50 | 64701.25 | 54309.25 | 46674.00 | 0.57 | 0.68 | 0.79 |
| RAB39A | 53017.75 | 66480.50 | 59920.00 | 48446.00 | 0.80 | 0.88 | 1.09 |
| RAB39B | 44288.00 | 45808.00 | 40718.50 | 27916.75 | 0.97 | 1.09 | 1.59 |
| RAB3A | 57389.00 | 56207.75 | 46626.75 | 36341.50 | 1.02 | 1.23 | 1.58 |
| RAB3B | 57073.25 | 57103.50 | 44491.00 | 33130.75 | 1.00 | 1.28 | 1.72 |
| RAB3C | 33703.75 | 45808.00 | 40718.50 | 27916.75 | 0.74 | 0.83 | 1.21 |
| RAB3D | 72404.25 | 61917.75 | 58930.00 | 48092.75 | 1.17 | 1.23 | 1.51 |
| RAB40A | 32480.00 | 45808.00 | 40718.50 | 27916.75 | 0.71 | 0.80 | 1.16 |
| RAB40AL | 51159.50 | 45808.00 | 40718.50 | 27916.75 | 1.12 | 1.26 | 1.83 |
| RAB40B | 56758.00 | 61917.75 | 58930.00 | 48092.75 | 0.92 | 0.96 | 1.18 |
| RAB40C | 68299.25 | 65767.75 | 60563.00 | 45185.75 | 1.04 | 1.13 | 1.51 |
| RAB41 | 53925.50 | 52352.25 | 44761.25 | 31201.25 | 1.03 | 1.20 | 1.73 |
| RAB42 | 39233.25 | 45808.00 | 40718.50 | 27916.75 | 0.86 | 0.96 | 1.41 |
| RAB43 | 45577.25 | 64701.25 | 54309.25 | 46674.00 | 0.70 | 0.84 | 0.98 |
| RAB44 | 49847.75 | 52352.25 | 44761.25 | 31201.25 | 0.95 | 1.11 | 1.60 |
| RAB4A | 61112.25 | 57103.50 | 44491.00 | 33130.75 | 1.07 | 1.37 | 1.84 |
| RAB4B | 57327.50 | 66480.50 | 59920.00 | 48446.00 | 0.86 | 0.96 | 1.18 |
| RAB5A | 56400.50 | 57103.50 | 44491.00 | 33130.75 | 0.99 | 1.27 | 1.70 |
| RAB5B | 58502.75 | 57103.50 | 44491.00 | 33130.75 | 1.02 | 1.31 | 1.77 |
| RAB5C | 47024.00 | 60878.50 | 53757.00 | 41389.00 | 0.77 | 0.87 | 1.14 |
| RAB6A | 66233.75 | 57103.50 | 44491.00 | 33130.75 | 1.16 | 1.49 | 2.00 |
| RAB6B | 66899.25 | 66480.50 | 59920.00 | 48446.00 | 1.01 | 1.12 | 1.38 |
| RAB6C | 27100.50 | 45808.00 | 40718.50 | 27916.75 | 0.59 | 0.67 | 0.97 |
| RAB7A | 67258.75 | 60878.50 | 53757.00 | 41389.00 | 1.10 | 1.25 | 1.63 |
| RAB7B | 55635.50 | 64701.25 | 54309.25 | 46674.00 | 0.86 | 1.02 | 1.19 |
| RAB7L1 | 64892.50 | 60878.50 | 53757.00 | 41389.00 | 1.07 | 1.21 | 1.57 |
| RAB8A | 56339.00 | 57103.50 | 44491.00 | 33130.75 | 0.99 | 1.27 | 1.70 |
| RAB8B | 75251.00 | 66480.50 | 59920.00 | 48446.00 | 1.13 | 1.26 | 1.55 |
| RAB9A | 25554.75 | 55655.00 | 42279.00 | 34164.00 | 0.46 | 0.60 | 0.75 |
| RAB9A | 47551.50 | 43884.00 | 36079.25 | 25806.00 | 1.08 | 1.32 | 1.84 |
| RAB9A | 56083.25 | 49721.75 | 44911.50 | 34100.75 | 1.13 | 1.25 | 1.64 |
| RAB9A | 59524.75 | 61917.75 | 58930.00 | 48092.75 | 0.96 | 1.01 | 1.24 |
| RAB9A | 73195.25 | 62653.75 | 57675.00 | 45848.25 | 1.17 | 1.27 | 1.60 |
| RAB9B | 46428.25 | 66480.50 | 59920.00 | 48446.00 | 0.70 | 0.77 | 0.96 |
| RABL2A | 52170.50 | 64701.25 | 54309.25 | 46674.00 | 0.81 | 0.96 | 1.12 |
| RABL2B | 63902.50 | 64701.25 | 54309.25 | 46674.00 | 0.99 | 1.18 | 1.37 |
| RABL3 | 35382.75 | 47314.50 | 41336.50 | 31515.75 | 0.75 | 0.86 | 1.12 |
| RABL6 | 37897.50 | 65767.75 | 60563.00 | 45185.75 | 0.58 | 0.63 | 0.84 |
| RACGAP1 | 19950.50 | 43884.00 | 36079.25 | 25806.00 | 0.45 | 0.55 | 0.77 |
| RALB | 55209.25 | 52352.25 | 44761.25 | 31201.25 | 1.05 | 1.23 | 1.77 |
| RASA1 | 40166.00 | 60268.75 | 47574.50 | 37832.50 | 0.67 | 0.84 | 1.06 |
| RGS6 | 42237.00 | 57514.25 | 46520.25 | 32816.25 | 0.73 | 0.91 | 1.29 |
| SAR1A | 29803.75 | 42578.00 | 35172.75 | 24136.50 | 0.70 | 0.85 | 1.23 |
| SAR1B | 21188.75 | 42578.00 | 35172.75 | 24136.50 | 0.50 | 0.60 | 0.88 |
| SEC23A | 55094.25 | 57994.25 | 44638.50 | 35343.25 | 0.95 | 1.23 | 1.56 |
| SEC24B | 66875.75 | 57994.25 | 44638.50 | 35343.25 | 1.15 | 1.50 | 1.89 |
| SEC31A | 44875.25 | 42878.00 | 32106.00 | 21444.75 | 1.05 | 1.40 | 2.09 |
| SEPT2 | 62488.25 | 63925.50 | 60479.50 | 46606.00 | 0.98 | 1.03 | 1.34 |
| SEPT6 | 59533.25 | 63925.50 | 60479.50 | 46606.00 | 0.93 | 0.98 | 1.28 |
| SEPT7 | 47174.75 | 63925.50 | 60479.50 | 46606.00 | 0.74 | 0.78 | 1.01 |
| SEPT9 | 60409.25 | 63925.50 | 60479.50 | 46606.00 | 0.94 | 1.00 | 1.30 |
| SMPD1 | 42559.00 | 61139.50 | 53217.50 | 42161.25 | 0.70 | 0.80 | 1.01 |
| SNAP23 | 54520.50 | 55655.00 | 42279.00 | 34164.00 | 0.98 | 1.29 | 1.60 |
| SNAP25 | 60602.75 | 60268.75 | 47574.50 | 37832.50 | 1.01 | 1.27 | 1.60 |
| SNAP29 | 46159.50 | 57514.25 | 46520.25 | 32816.25 | 0.80 | 0.99 | 1.41 |
| SNF8 | 44806.00 | 50124.75 | 38891.25 | 27570.50 | 0.89 | 1.15 | 1.63 |
| SNX26 | 50148.75 | 49892.75 | 38856.25 | 25878.75 | 1.01 | 1.29 | 1.94 |
| STAM | 50011.25 | 50124.75 | 38891.25 | 27570.50 | 1.00 | 1.29 | 1.81 |
| STAMBP | 53433.00 | 50124.75 | 38891.25 | 27570.50 | 1.07 | 1.37 | 1.94 |
| STX10 | 27113.50 | 55655.00 | 42279.00 | 34164.00 | 0.49 | 0.64 | 0.79 |
| STX11 | 44851.25 | 55655.00 | 42279.00 | 34164.00 | 0.81 | 1.06 | 1.31 |
| STX16 | 48000.25 | 55655.00 | 42279.00 | 34164.00 | 0.86 | 1.14 | 1.40 |
| STX17 | 36638.00 | 42578.00 | 35172.75 | 24136.50 | 0.86 | 1.04 | 1.52 |
| STX18 | 37983.25 | 42578.00 | 35172.75 | 24136.50 | 0.89 | 1.08 | 1.57 |
| STX19 | 43554.25 | 47314.50 | 41336.50 | 31515.75 | 0.92 | 1.05 | 1.38 |
| STX1A | 62259.75 | 60268.75 | 47574.50 | 37832.50 | 1.03 | 1.31 | 1.65 |
| STX1B | 46569.00 | 49892.75 | 38856.25 | 25878.75 | 0.93 | 1.20 | 1.80 |
| STX2 | 53822.25 | 56207.75 | 46626.75 | 36341.50 | 0.96 | 1.15 | 1.48 |
| STX3 | 51935.00 | 60268.75 | 47574.50 | 37832.50 | 0.86 | 1.09 | 1.37 |
| STX4 | 54329.75 | 60268.75 | 47574.50 | 37832.50 | 0.90 | 1.14 | 1.44 |
| STX5 | 49812.50 | 60268.75 | 47574.50 | 37832.50 | 0.83 | 1.05 | 1.32 |
| STX6 | 28479.00 | 57994.25 | 44638.50 | 35343.25 | 0.49 | 0.64 | 0.81 |
| STX7 | 52521.50 | 53274.50 | 38684.00 | 32412.00 | 0.99 | 1.36 | 1.62 |
| STX8 | 46878.00 | 57514.25 | 46520.25 | 32816.25 | 0.82 | 1.01 | 1.43 |
| STXBP1 | 61846.00 | 60268.75 | 47574.50 | 37832.50 | 1.03 | 1.30 | 1.63 |
| STXBP2 | 41864.75 | 60268.75 | 47574.50 | 37832.50 | 0.69 | 0.88 | 1.11 |
| STXBP3 | 57952.00 | 60268.75 | 47574.50 | 37832.50 | 0.96 | 1.22 | 1.53 |
| STXBP4 | 27984.50 | 47314.50 | 41336.50 | 31515.75 | 0.59 | 0.68 | 0.89 |
| STXBP5 | 52826.50 | 49892.75 | 38856.25 | 25878.75 | 1.06 | 1.36 | 2.04 |
| STXBP6 | 43523.25 | 43884.00 | 36079.25 | 25806.00 | 0.99 | 1.21 | 1.69 |
| SYT1 | 57324.00 | 53274.50 | 38684.00 | 32412.00 | 1.08 | 1.48 | 1.77 |
| SYT10 | 35536.75 | 47314.50 | 41336.50 | 31515.75 | 0.75 | 0.86 | 1.13 |
| SYT11 | 39626.25 | 42878.00 | 32106.00 | 21444.75 | 0.92 | 1.23 | 1.85 |
| SYT12 | 43325.25 | 49892.75 | 38856.25 | 25878.75 | 0.87 | 1.12 | 1.67 |
| SYT13 | 33130.75 | 42578.00 | 35172.75 | 24136.50 | 0.78 | 0.94 | 1.37 |
| SYT14 | 29443.50 | 47314.50 | 41336.50 | 31515.75 | 0.62 | 0.71 | 0.93 |
| SYT15 | 42325.00 | 47050.25 | 38202.00 | 25381.50 | 0.90 | 1.11 | 1.67 |
| SYT16 | 56463.75 | 47050.25 | 38202.00 | 25381.50 | 1.20 | 1.48 | 2.22 |
| SYT17 | 51798.75 | 42578.00 | 35172.75 | 24136.50 | 1.22 | 1.47 | 2.15 |
| SYT2 | 33262.50 | 49892.75 | 38856.25 | 25878.75 | 0.67 | 0.86 | 1.29 |
| SYT3 | 47648.25 | 47050.25 | 38202.00 | 25381.50 | 1.01 | 1.25 | 1.88 |
| SYT4 | 40783.00 | 53274.50 | 38684.00 | 32412.00 | 0.77 | 1.05 | 1.26 |
| SYT5 | 53814.50 | 53274.50 | 38684.00 | 32412.00 | 1.01 | 1.39 | 1.66 |
| SYT6 | 46602.25 | 47314.50 | 41336.50 | 31515.75 | 0.98 | 1.13 | 1.48 |
| SYT7 | 36296.75 | 57920.00 | 43760.50 | 36524.75 | 0.63 | 0.83 | 0.99 |
| SYT8 | 38557.50 | 49892.75 | 38856.25 | 25878.75 | 0.77 | 0.99 | 1.49 |
| SYT9 | 57769.00 | 47314.50 | 41336.50 | 31515.75 | 1.22 | 1.40 | 1.83 |
| TBC1D15 | 45577.75 | 47050.25 | 38202.00 | 25381.50 | 0.97 | 1.19 | 1.80 |
| TBK1 | 51424.00 | 66605.75 | 62758.00 | 50664.00 | 0.77 | 0.82 | 1.02 |
| TSG101 | 37753.50 | 51691.75 | 40312.00 | 28852.25 | 0.73 | 0.94 | 1.31 |
| TSNARE1 | 49088.50 | 50124.75 | 38891.25 | 27570.50 | 0.98 | 1.26 | 1.78 |
| UBAP1 | 45026.25 | 63925.50 | 60479.50 | 46606.00 | 0.70 | 0.74 | 0.97 |
| USP8 | 56902.50 | 50124.75 | 38891.25 | 27570.50 | 1.14 | 1.46 | 2.06 |
| VAMP1 | 55194.00 | 53274.50 | 38684.00 | 32412.00 | 1.04 | 1.43 | 1.70 |
| VAMP2 | 47581.25 | 53274.50 | 38684.00 | 32412.00 | 0.89 | 1.23 | 1.47 |
| VAMP3 | 65644.75 | 57514.25 | 46520.25 | 32816.25 | 1.14 | 1.41 | 2.00 |
| VAMP7 | 61808.00 | 53274.50 | 38684.00 | 32412.00 | 1.16 | 1.60 | 1.91 |
| VAMP8 | 40757.00 | 55655.00 | 42279.00 | 34164.00 | 0.73 | 0.96 | 1.19 |
| VAPB | 61712.00 | 57920.00 | 43760.50 | 36524.75 | 1.07 | 1.41 | 1.69 |
| VPS13B | 44162.75 | 47314.50 | 41336.50 | 31515.75 | 0.93 | 1.07 | 1.40 |
| VPS24 | 37102.50 | 51175.25 | 41281.00 | 29996.75 | 0.73 | 0.90 | 1.24 |
| VPS25 | 48719.00 | 51175.25 | 41281.00 | 29996.75 | 0.95 | 1.18 | 1.62 |
| VPS28 | 54631.75 | 66605.75 | 62758.00 | 50664.00 | 0.82 | 0.87 | 1.08 |
| VPS36 | 43263.75 | 51175.25 | 41281.00 | 29996.75 | 0.85 | 1.05 | 1.44 |
| VPS4A | 29368.75 | 51175.25 | 41281.00 | 29996.75 | 0.57 | 0.71 | 0.98 |
| VTA1 | 45647.00 | 51175.25 | 41281.00 | 29996.75 | 0.89 | 1.11 | 1.52 |
| VTI1B | 51879.50 | 51175.25 | 41281.00 | 29996.75 | 1.01 | 1.26 | 1.73 |
| WTH3DI | 43780.75 | 45808.00 | 40718.50 | 27916.75 | 0.96 | 1.08 | 1.57 |
