## Supplemental Table 3 for "The Septin Cytoskeleton is Required for Plasma Membrane Repair"

**Supplemental Table 3. Sequences used in this work**. The silencing sequences are highlighted in grey.

|  | **Oligonucleotides (shRNA)** |
| --- | --- |
| SEPTIN7  shRNA1* | 5’-CTAGCGGGAAGCTCAACAACGTATTT**CTCGAG**AAATACGTTGTTGAGCTTCCCTTTTT-3’  Sense  5’-AATTAAAAAGGGAAGCTCAACAACGTATTT**CTCGAG**AAATACGTTGTTGAGCTTCCCG-3’  antisense |
|  | **Sequencing Primers** |
| EZ-pLKO | 5’-GATTAGTGAACGGATCTCGACGG-3’ forward  5’-AACCCAGGGCTGCCTTGG-3’ reverse |

* SEPTIN 7 shRNA1 sequence obtained from the Genetic Perturbation Platform (Broad Institute).
